# Neural Networks and Foundation Models: Two Strategies for EEG-to-fMRI Prediction

**DOI:** 10.1101/2025.05.06.652346

**Authors:** Maël Donoso

## Abstract

Electroencephalography (EEG) and functional Magnetic Resonance Imaging (fMRI) are two widely used neuroimaging techniques, with complementary strengths and weaknesses. Predicting fMRI activity from EEG activity could give us the best of both worlds, and open new horizons for neuroscience research and neurotechnology applications. Here, we formulate this prediction objective both as a classification task (predicting whether the fMRI signal increases or decreases) and a regression task (predicting the value of this signal). We follow two distinct strategies: training classical machine learning and deep learning models (including MLP, CNN, RNN, and transformer) on an EEG-fMRI dataset, or leveraging the capabilities of pre-trained large language models (LLMs) and large multimodal models. We show that predicting fMRI activity from EEG activity is possible for the brain regions defined by the Harvard-Oxford cortical atlas, in the context of subjects performing a neurofeedback task. Interestingly, both strategies yield promising results, possibly highlighting two complementary paths for our prediction objective. Furthermore, a Chain-of-Thought approach demonstrates that LLMs can infer the cognitive functions associated with EEG data, and subsequently predict the fMRI data from these cognitive functions. The natural combination of the two strategies, i.e., fine-tuning an LLM on an EEG-fMRI dataset, is not straightforward and would certainly require further study. These findings could provide important insights for enhancing neural interfaces and advancing toward a multimodal foundation model for neuroscience, integrating EEG, fMRI, and possibly other neuroimaging modalities.

## 1 Introduction

Electroencephalography (EEG) and functional Magnetic Resonance Imaging (fMRI) are two widely used neuroimaging techniques for human brain activity investigation. While EEG provides a direct measure of the electrical activity of the brain by using electrodes placed on the scalp, fMRI measures the activity of the brain indirectly, by detecting changes in the cerebral blood flow and oxygen demand. As a consequence, EEG is typically used to record the activity of cortical areas, whereas fMRI can detect the activity of subcortical regions as well. EEG provides a high temporal resolution, and is a relatively simple and inexpensive neuroimaging technique, compatible with wearable devices. By contrast, fMRI offers a high spatial resolution, at the expense of a greater complexity and cost, and requires the subject to remain immobile inside the scanner. These complementary strengths and weaknesses motivated the emergence of EEG-fMRI research, in which both techniques are used simultaneously [1, 2].

### 1.1 Predicting fMRI from EEG

The accessibility of EEG-fMRI datasets allowed researchers to explore whether fMRI activity could be predicted from EEG activity. Early research demonstrated that blood oxygenation level dependent (BOLD) [3] fluctuations in the occipital cortex can be predicted from EEG data, using a linear combination of B-spline basis functions [4]. Beyond the occipital cortex, two early studies reported the successful prediction of amygdala activity, using classical machine learning models such as Ridge regression [5, 6]. Another study focused on connectivity, by applying a model based on sparse canonical correlation analysis to the prediction of the fMRI connectome from the EEG connectome at the scale of the entire brain [7].

More recently, deep learning models were applied to the prediction of fMRI activity from EEG activity, including convolutional neural networks [8], attentional graphs [9], autoencoders and generative adversarial networks [10]. Two preprints proposed novel neural network architectures with the objective to improve interpretability [11, 12]. Research also focused on addressing the hemodynamic delay of the BOLD signal [13], and on predicting the activity of specific brain regions such as the inferior frontal gyrus [14] and the ventral striatum [15], respectively associated with cognitive control and reward processes.

Very recently, a series of preprints proposed additional approaches, such as using sinusoidal representation networks [16], transformers [17, 18], and simpler neural networks inspired by the U-Net architecture [19]. Overall, current research consistently highlights that predicting fMRI activity from EEG activity could significantly enhance neuroimaging capabilities by giving us the best of both worlds, and open new horizons for neuroscience research and neurotechnology applications [9, 10, 11, 12, 13, 14, 15, 16, 17, 18, 19].

### 1.2 Neurofeedback

One of the neurotechnologies that might benefit the most from EEG-to-fMRI prediction is neurofeedback (NF). NF consists in providing real-time information to a subject about their own brain activity, and encouraging them to adapt their behavior according to this measure. The objective of a NF protocol is for the subject to learn self-regulation and reach a certain cognitive state, whether by increasing or decreasing the power of specific EEG frequency bands (EEG NF), the fMRI activity in specific brain regions (fMRI NF), or a combination of both (EEG-fMRI NF) [20]. A study demonstrated that fMRI NF scores and EEG-fMRI NF scores can be predicted from EEG data using a sparse regression model, and that this prediction adds significant information compared to EEG NF scores alone [21].

Since the prediction of fMRI activity from EEG activity is an emerging area of research, there is, to our knowledge, no systematic review identifying the most promising brain regions and cognitive processes that could be targeted for EEG-to-fMRI prediction. Our objective is not to provide such a systematic review. However, based on their importance for understanding human cognition, we speculate that the brain regions associated with valuation [22, 23], motivation [24, 25], exploration [26], decision-making [27, 28, 29], learning [30, 31], and reasoning [32], in particular, might be valuable candidates. Arguably, all these cognitive functions are engaged during a NF protocol, potentially highlighting the importance of EEG-fMRI NF datasets for future research. This perspective motivated us to select such a dataset for our experiments, although our conclusions do not depend on the fact that the EEG-fMRI data was acquired during a NF protocol.

### 1.3 Toward a multimodal foundation model

We argue that a model capable of predicting fMRI activity from EEG activity at the scale of the entire brain would meet the definition of a foundation model, since it would likely be trained on broad data and could be adapted to a wide range of tasks [33]. Recently, foundation models focusing on a single neuroimaging modality have been developed using self-supervised learning, whether for EEG [34, 35, 36, 37] or fMRI [38, 39, 40]. However, the model we envision would serve a different purpose. It would be a multimodal foundation model, capable of performing “neural translation” between several neuroimaging modalities. Advancing toward this multimodal foundation model could come with significant challenges, considering both the scarcity and heterogeneity of EEG-fMRI datasets.

Here, we suggest that these challenges might eventually be overcome using a combination of two strategies. The first, classical strategy is the supervised learning approach, which consists in training classical machine learning and deep learning models on an EEG-fMRI dataset. The second, novel strategy consists in directly leveraging the capabilities of pre-trained large language models (LLMs) and large multimodal models (LMMs) for this prediction objective. Indeed, LLMs such as Gemma- 2-2B-IT [41] and Llama-3.2-3B-Instruct [42], and LMMs such as PaliGemma2-3B-Mix-224 [43] are all pre-trained on extensive corpora of documents, which we assume should include a significant number of neuroscience articles and books, or other sources of knowledge on EEG and fMRI patterns. This extensive pre-training might help us to overcome the scarcity and heterogeneity of EEG-fMRI datasets, by effectively outsourcing a part of the problem. Foundation models are rapidly emerging as powerful instruments for accelerating scientific discovery [44, 45], including in neuroscience [46], and a very recent study explored the possibility of encoding EEG signals as LLM-compatible tokens [47]. However, to our knowledge, LLMs and LMMs have not yet been directly leveraged for the complex multimodal task of EEG-to-fMRI prediction.

### 1.5 Two strategies for EEG-to-fMRI prediction

Here, we evaluate these two strategies, i.e., training our own models or relying on pre-trained foundation models, on the same EEG-fMRI dataset. For both strategies, we formulate the prediction objective as a classification task: predicting whether the fMRI signal increases or decreases based on the EEG signal. For the first strategy, i.e., training our own models, we also add a regression task: predicting the value of the fMRI signal based on the EEG signal. Critically, leveraging LLMs or LMMs implies that both features and targets should be designed in order to be described by relevant keywords: EEG frequency bands “with names” instead of arbitrary spectral representations, fMRI brain regions “with names” instead of arbitrary voxel positions, and semantically meaningful classes (“increase”, “decrease”). Therefore, we do not leverage LLMs or LMMs for the regression task, whose targets are numerical.

Since neuroscience articles and books often associate EEG and fMRI patterns with cognitive functions, the keywords describing the latter might serve as intermediate representations between these two neuroimaging modalities. Following this idea, we hypothesize that LLMs could have the capability to infer the cognitive functions associated with EEG data, and subsequently predict the fMRI data from these cognitive functions. We test this hypothesis by implementing a Chain-of-Thought (CoT) [48] approach separating these two steps. Finally, we attempt the natural combination of the two strategies by fine-tuning an LLM on an EEG-fMRI dataset, in order to complement its pre-trained capabilities with additional, task-specific knowledge.

While potentially intriguing, the idea that a multimodal foundation model for EEG-to-fMRI prediction might be partially built upon existing language or vision-language foundation models is consistent with several emerging trends in AI research. In particular, it resonates with the growing interest in integrating foundation models with more specific models to address complex multimodal tasks [49, 50]. Furthermore, it aligns with the idea of using language as a universal interface between different data modalities, a strategy which proved to be successful, in a different field, to predict the evolution of proteins [51]. Overall, we believe that evaluating the capabilities of pre-trained foundation models, and comparing them with the performance of classical machine learning and deep learning models, is an important step that could open a new path for EEG-to-fMRI prediction.

## 2 Data

We use the publicly available EEG-fMRI NF dataset *A multi-modal human neuroimaging dataset for data integration: simultaneous EEG and fMRI acquisition during a motor imagery neurofeedback task: XP1* [52], which can be downloaded from OpenNeuro [53], an open repository for neuroimaging data. This dataset is the first published open-access bimodal NF dataset integrating EEG and fMRI, and was used in the EEG-fMRI NF study mentioned earlier [21]. The neuroimaging files are stored in Brain Imaging Data Structure (BIDS) format [54], and the dataset is released under the CC0 license. The dataset authors conducted a NF experiment in which 10 subjects (8 male, 2 female, median age = 27) were instructed to use EEG NF scores and fMRI NF scores to perform as well as possible in a motor imagery task (i.e., they needed to execute mentally a movement without any muscle activation). The experiment included six conditions, with alternating rest and task blocks within the conditions.

In our research, we focus on three conditions, which were completed in random order by the different subjects: 1) The eegfmriNF condition, corresponding to bimodal EEG-fMRI NF. 2) The eegNF condition, corresponding to unimodal EEG NF. 3) The fmriNF condition, corresponding to unimodal fMRI NF. For each condition, the dataset includes the raw fMRI data, the raw EEG data, and the EEG data preprocessed by the dataset authors. The fMRI data was acquired with a 3T MRI scanner using echo-planar imaging, with a repetition time of 2 seconds and a voxel size of 2× 2× 4 mm^3^. The EEG data was acquired with a 64-channel montage based on the extended 10–20 system, at a sampling rate of 5,000 Hz. The dataset authors subsequently resampled the EEG data to 200 Hz, and applied a low-pass filter at 50 Hz during their preprocessing. The acquisition of EEG data in an fMRI environment is technically complex and can result in high noise [52], making it particularly useful that the dataset authors included the preprocessed EEG data. However, since the preprocessed fMRI data is not included, it is necessary to perform some standard fMRI preprocessing steps before running the experiments.

## 3 Methods

### 3.1 Preprocessing

#### fMRI preprocessing

We preprocess the raw fMRI data using fMRIPrep [55], a robust preprocessing pipeline which automatically performs a series of standard fMRI preprocessing steps, such as motion correction, coregistration, and spatial normalization. In order to ensure subsequent compatibility with the Harvard-Oxford cortical atlas [56], we normalize the fMRI data using the MNI152Lin output space. We further preprocess the fMRI data using the NiBabel [57] and Nilearn [58] libraries. For each fMRI scan, we extract the average voxel values of our brain regions of interest, which are the regions defined in the Harvard-Oxford cortical atlas. Within this atlas, we use the maximum probability map with no threshold, assigning each voxel to the region with the highest probability, therefore ensuring full and non-overlapping cortical coverage. We remove a systematic drift of the BOLD signal, which tends to increase during an fMRI session. We also normalize the BOLD signal for each region by subtracting the mean and dividing by the standard deviation, and replace the outliers (STD > 3) with the value of the previous scan. The fMRIPrep preprocessing is documented in *Supplementary README*.*md*, and the details about the additional fMRI preprocessing can be found in *Supplementary Notebook 1*.

#### EEG preprocessing

We preprocess the EEG data using the MNE-Python [59] and YASA [60] libraries. We divide the EEG data into 2-second segments aligned with the fMRI scans. For each EEG segment, we compute the band powers for a series of frequency bands of interest: delta (1-4 Hz), theta (4-8 Hz), alpha (8-12 Hz), sigma (12-16 Hz), beta (16-30 Hz), and gamma (30-40 Hz). We normalize the band powers for each channel by subtracting the mean and dividing by the standard deviation, and replace the outliers (STD > 4) with the value of the previous segment. The details can be found in *Supplementary Notebook 2*.

#### Features and targets

For the classification task, the target is a binary label indicating whether the normalized BOLD signal is increasing or decreasing between two successive scans. For the regression task, the target is the normalized BOLD signal at a given scan. For both tasks, unless stated otherwise, the features consist of the normalized band powers computed at a given scan, along with those computed during the 5 preceding scans. This 5-scan sequence corresponds to 10 seconds, a duration that encompasses the peak of the hemodynamic response function [61], therefore improving our chances of capturing, in the EEG data, a trace of the events that influenced the fMRI data. Since the eegNF condition of one subject is missing, we remove this subject from all the experiments.

### 3.2 Machine learning models

We train a series of classical machine learning models using the Scikit-Learn [62] library. For the classification task, we select the following models: logistic regression, k-nearest neighbors (KNN), decision tree (DT), random forest (RF), support vector machine (SVM), and extreme gradient boosting (XGBoost) [63]. For the regression task, we select the following models: linear regression, KNN, DT, RF, SVM, and XGBoost. We evaluate the accuracy of the classification task, and the mean absolute error (MAE) of the regression task. For both tasks, we use the eegfmriNF condition as the train set and the fmriNF condition as the test set, individually for each subject. We fine-tune the number of neighbors for the KNN model, and the depth of the tree for the DT model, using grid search cross-validation. The details can be found in *Supplementary Notebooks 3-4*.

### 3.3 Deep learning models

We train a series of deep learning models using the TensorFlow [64] library. We select the following architectures: multi-layer perceptron (MLP), convolutional neural network (CNN) [65], recurrent neural network (RNN) with gated recurrent units [66], and transformer [67]. For the classification task, we use the binary cross-entropy loss function and evaluate the accuracy, whereas for the regression task, we use the mean squared error loss function and evaluate the MAE. For both tasks, we use the eegfmriNF condition as the train set, the eegNF condition as the validation set, and the fmriNF condition as the test set, individually for each subject. We train our models on the normalized band powers, except for the CNN model, used as a control, for which we rely directly on the EEG signal without band power extraction. We use the Adam [68] optimizer and the ReLU activation function for hidden layers. The details, including the exact architecture and number of trainable parameters for each model, can be found in *Supplementary Notebooks 5-8*.

### 3.4 Foundation models

#### Large language models

We leverage two LLMs, Gemma-2-2B-IT and Llama-3.2-3B-Instruct, using the Hugging Face Transformers [69] library. We focus on the classification task and the eegfmriNF condition, and since LLMs require significantly more computational resources than our classical machine learning and deep learning models, we select only a subset of our fMRI scans, band powers, and brain regions of interest. In order to further control the complexity of the prompts, we associate each fMRI brain region with a single EEG channel, which serves as its sole predictor. This association follows an ad hoc region-channel mapping, mainly based on electrode proximity to the brain region, and established, perhaps fittingly, with the help of an LLM. We query both models with prompts including a general context, the selected EEG channel, band powers, and brain region, the normalized band power values, and a description of the prediction task to perform. In this experiment and the following ones, we select the model parameters in order to ensure a relative variety of responses, and parse the model outputs by detecting the presence of inflected forms of the keywords “increase” or “decrease” (e.g., “increases”, “increased”, “increasing”) in the generated text. The cases where these keywords are missing, or where both keywords are present, are labeled as invalid predictions. We also ensure that the selected subset of brain regions spans at least a reasonable fraction of the cortex and a variety of cognitive functions. The details, including the region-channel mapping and the prompt structure, can be found in *Supplementary Notebook 9*.

#### Chain-of-Thought and fine-tuning

We conduct two additional experiments with Gemma-2-2B- IT. In the first experiment, we implement an intermediate reasoning step using a CoT approach. Specifically, we query the model with a first prompt to infer cognitive functions based on EEG data, then use a second prompt to infer fMRI data based on these cognitive functions. Both prompts also include a general context and a description of the prediction task to perform. In the second experiment, we fine-tune the model with parameter-efficient fine-tuning [70] on input-output pairs obtained from the fmriNF condition, before evaluating again its performance on the eegfmriNF condition. Fine-tuning is implemented by applying low-rank adaptation [71] to the query and value projection layers, and the model is optimized using the AdamW [72] optimizer. To improve stability and prevent erratic behavior (e.g., the model generating a long chain of “increase decrease increase decrease… “), the model is fine-tuned independently for each subject-region pair. The details can be found in *Supplementary Notebooks 10-11*.

#### Large multimodal model

We leverage one LMM, PaliGemma2-3B-Mix-224, using the Hugging Face Transformers library. We focus again on the classification task, the eegfmriNF condition, and a subset of our fMRI scans and brain regions of interest. However, instead of using multiple frequency bands and a single EEG channel, we take the opposite approach. Specifically, we create a topographic map using the MNE-Python library, displaying the beta band power (16-30 Hz) across all EEG channels. We prompt the model with this image, along with a general context, the selected brain region, and a description of the prediction task to perform. The details can be found in *Supplementary Notebook 12*.

### 3.5 Statistical tests

For each model, we compare the mean accuracy or MAE to a baseline, across all subject-region pairs (9 subjects *×*49 regions = 441 pairs). Since we do not assume normality, we perform a one-sided Wilcoxon signed-rank test using the SciPy [73] library, proceeding similarly for the classification and regression tasks (but searching for opposite effects: higher accuracy for classification, lower MAE for regression). For the foundation models, we also perform McNemar tests, pooling together the predictions from all subjects and brain regions, to compensate for the smaller sample size. Indeed, not only are the foundation models evaluated on a selection of our fMRI scans of interest, but missing or ambiguous predictions must be dynamically excluded, resulting in significantly fewer data points.

For the classical machine learning and deep learning models, the baselines for our statistical tests are relatively straightforward. For the classification task, we use as the baseline the constant prediction of an increase, which corresponds to the majority class, although the two classes are almost perfectly balanced (∼50% each). For the regression task, we use as the baseline the constant prediction of the mean value of the signal, which is zero due to the normalization. For the foundation models, given the lower number of fMRI scans and their more variable target distribution, we proceed differently. We use as the baseline the predictions of a model that randomly samples labels from the true target distribution of the selected fMRI scans, excluding the missing or ambiguous cases from this distribution. We repeat the sampling for 1,000 iterations (Wilcoxon tests) or 10,000 iterations (McNemar tests). The details can be found in *Supplementary Notebook 13*.

## 4 Results

### 4.1 Machine learning models

All the classical machine learning models for classification, i.e., the logistic regression, KNN, DT, RF, SVM, and XGBoost models, reach an accuracy higher than the baseline (p < 0.01 for KNN, p < 0.001 for the other models), as shown in Figure 1. Among the classical machine learning models for regression, only the RF (p < 0.05) and SVM (p < 0.001) models perform better than the baseline, as shown in Figure 2. The detailed results can be found in *Supplementary Notebook 14*.

**Figure 1:**
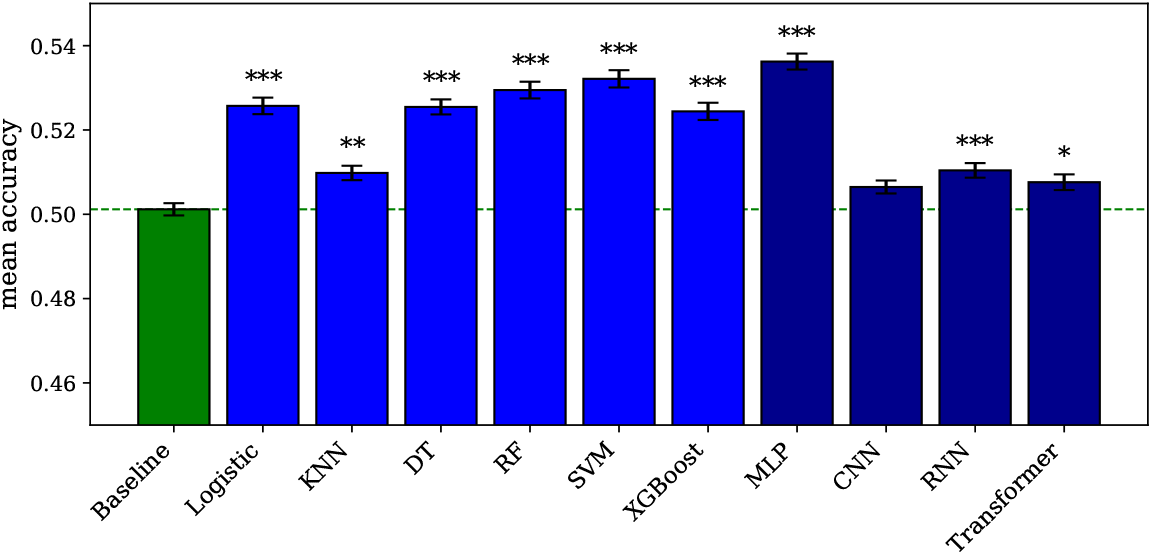
Machine learning and deep learning models for classification. The error bars represent the standard error of the mean across subject-region pairs. Significance is indicated by asterisks: * for *p <* 0.05, ** for *p <* 0.01, and *** for *p <* 0.001. The dashed line indicates the baseline level.

**Figure 2:**
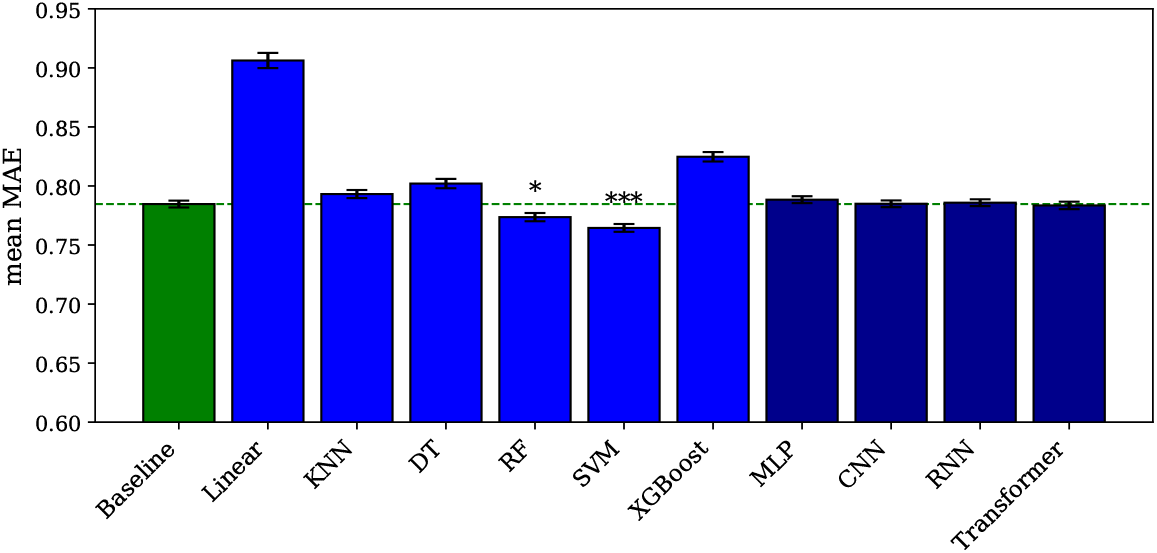
Machine learning and deep learning models for regression. The error bars represent the standard error of the mean across subject-region pairs. Significance is indicated by asterisks: * for *p <* 0.05, ** for *p <* 0.01, and *** for *p <* 0.001. The dashed line indicates the baseline level.

### 4.2 Deep learning models

Among the deep learning models for classification, the MLP (p < 0.001), RNN (p < 0.001), and transformer (p < 0.05) models reach an accuracy higher than the baseline, as shown in Figure 1. However, the training graphs of all models except MLP show signs of overfitting, with the validation loss often stagnating or even increasing over the epochs. Among the deep learning models for regression, no model performs better than the baseline, as shown in Figure 2. We also evaluate the number of floating-point operations (FLOPs) necessary for running the MLP (∼23M), CNN (∼51M), RNN (∼39M), and transformer (∼76M) models, to confirm that the higher performance of the MLP model for the classification task is not due to model complexity. The training graphs can be found in *Supplementary Notebooks 5-8*, and the detailed results in *Supplementary Notebook 14*.

### 4.3 Foundation models

Neither the LLMs nor the LMM perform better than the baseline when the predictions are evaluated using the one-sided Wilcoxon signed-rank test. However, when the predictions are evaluated using the less conservative McNemar test across multiple iterations, a more nuanced view emerges. For the different instances of the Gemma model, we observe a relatively low median p-value (median p = 0.12, 0.16, 0.15, 0.14) and a high proportion of statistically significant iterations (proportion p < 0.05 = 29%, 23%, 23%, 24%). The Llama model shows the opposite trend, with a relatively high median p-value (median p = 0.66) and a low proportion of statistically significant iterations (proportion p < 0.05 = 0.5%). The PaliGemma model shows an intermediate pattern (median p = 0.31, proportion p < 0.05 = 8.6%). The proportion of missing or ambiguous predictions varies across models, but is always less than 10%. The detailed results can be found in *Supplementary Notebooks 13-14*.

The visualization of the p-value distributions using histograms provides important insights as well. As shown in Figure 3, for the different instances of the Gemma model, we observe a right-skewed p-value distribution, with a mode near p = 0 and a long tail toward p = 1, suggesting that the Gemma model achieves small but reliable gains over the baseline, whether it is used with direct prompting or with a CoT approach. The Llama model shows almost the opposite trend, with a weakly left-skewed p-value distribution, suggesting an absence of improvement over the baseline. The PaliGemma model shows again an intermediate pattern, with a weakly right-skewed p-value distribution, suggesting a more ambiguous behavior. We also evaluate the number of FLOPs necessary for running the Gemma (∼3T), Llama (∼3.7T), and PaliGemma (∼1.6T) models, obtaining values that stand several orders of magnitude above those measured for the classical machine learning and deep learning models. Finally, we compare the performance of the Gemma model with and without fine-tuning, and observe that the difference is not statistically significant, whether the comparison is performed using the one-sided Wilcoxon signed-rank test (p = 0.42) or the McNemar test (p = 0.87).

**Figure 3:**
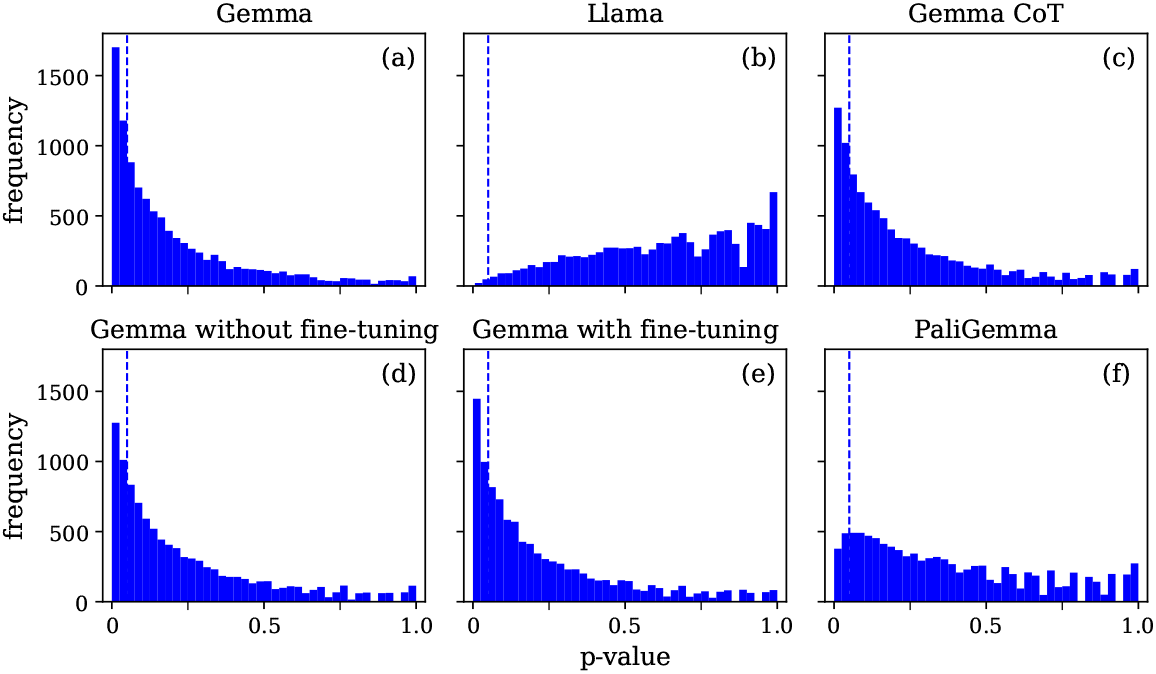
Distribution of p-values for the foundation models. (a) and (d) are both based on the Gemma model without fine-tuning, but differ by the subset of brain regions included. (d) and (e) include the same subset of brain regions. The dashed lines indicate the *p <* 0.05 threshold.

## 5 Discussion

The first strategy, which consists in training classical machine learning and deep learning models, proves to be the most successful. Interestingly, the MLP model is the best-performing model for the classification task, and the only deep learning model achieving a higher accuracy than the classical machine learning models. The superiority of the MLP model over the CNN, RNN, and transformer models is not due to model complexity, as confirmed by the number of FLOPs. Rather, it suggests that the relevant patterns for EEG-to-fMRI prediction might be globally distributed across space (EEG channels) and time (fMRI scans), therefore favoring feedforward neural network architectures which do not rely too strongly on spatial or temporal priors. We might expect that this global distribution of relevant features could also favor text-only LLMs over LMMs, since the latter typically rely on data with stronger structure, such as images. The superiority of the MLP model is also consistent with a very recent preprint suggesting that simple neural networks for EEG-to-fMRI prediction can outperform more complex ones [19]. Intriguingly, although the majority of our neural network architectures achieve significant results in the classification task, none of them performs better than the baseline in the regression task, whereas the RF and SVM models do. This suggests that the complexity threshold for a minimally working neural network might be higher for the regression task.

The second strategy, which consists in directly leveraging the capabilities of pre-trained foundation models, shows promising results as well. Although the observed effects are much smaller than for the classical machine learning and deep learning models, the Gemma model achieves statistically reliable gains over the baseline. Furthermore, our CoT approach demonstrates that the Gemma model can infer the cognitive functions associated with EEG data, and subsequently predict the fMRI data from these cognitive functions. This suggests the possibility that the performance of the Gemma model with direct prompting might be driven by a similar mechanism, with the model implicitly relying on cognitive functions as an intermediate reasoning step between EEG and fMRI, even in the absence of native CoT capabilities. The lower performance of the Llama and PaliGemma models might reflect their intrinsic limitation for this particular prediction task, or simply suggest that more exploration is needed to adapt the prompts and parameters for each foundation model. The absence of improvement observed after fine-tuning the Gemma model may seem intriguing, but while it is difficult to provide a definitive interpretation, a very recent study highlighted that the result of fine-tuning an LLM for a scientific problem depends both on the dataset and on the complexity of the question [74]. This research reported a weak predictive power for complex input variables and a very low number of training data points, which is precisely our situation. Overcoming this difficulty, and establishing a more successful fine-tuning strategy, would certainly require further study.

### Limitations

The EEG-fMRI dataset on which this research is based is limited in size, and unbalanced in terms of sex and age. Given the limited training set, the classical machine learning and deep learning models are evaluated across sessions, but not across subjects. The exact data on which the foundation models have been pre-trained, and in particular the number of neuroscience articles and books, or other sources of knowledge on EEG and fMRI patterns, is not publicly available. Running the foundation models requires significantly more computational resources than running the classical machine learning and deep learning models, which forces us to focus on a subset of our fMRI scans, band powers, and brain regions of interest, and to use an ad hoc region-channel mapping. Also because of this computational cost, we only evaluate small foundation models, which may be less performant than larger ones. In general, this research focuses on demonstrating the feasibility of the two strategies for EEG-to-fMRI prediction, and does not aim to achieve the best performance possible.

### Future directions

For the first strategy, a natural path forward would be to explore additional preprocessing pipelines and neural network architectures, and to experiment with more specific models, relying more strongly on our knowledge of the human brain. We would certainly benefit from larger EEG-fMRI datasets, and more balanced in terms of sex and age, if they become available in the future. Since the scarcity and heterogeneity of EEG-fMRI datasets is currently a significant obstacle, developing methods for integrating heterogeneous datasets (e.g., different EEG montages, fMRI scan durations, etc.) could also be highly valuable. For the second strategy, the possible future directions are relatively symmetrical. It would make sense to explore additional prompt engineering techniques and model parameters, and to experiment with foundation models specifically pre-trained on the neuroscience literature. We would certainly benefit from more advanced and general foundation models, which will likely become available in the future, in particular if these models have native CoT capabilities, potentially allowing them to explicitly use cognitive functions as an intermediate reasoning step between EEG and fMRI. Developing methods for successfully integrating the two strategies, for example by fine-tuning foundation models on existing EEG-fMRI datasets, could also be a promising path. Finally, both strategies could be extended beyond EEG and fMRI, to other neuroimaging modalities such as magnetoencephalography, bringing us a step closer to a multimodal foundation model for neuroscience.

## 5 Conclusion

In this research, we demonstrate the feasibility of predicting fMRI activity from EEG activity by following two distinct strategies: training classical machine learning and deep learning models on an EEG-fMRI dataset, or leveraging the capabilities of pre-trained foundation models. When this prediction objective is formulated as a classification task, the MLP model stands out as particularly effective, while the Gemma model achieves statistically reliable gains over the baseline, whether it is used with direct prompting or with a CoT approach. The latter case demonstrates that the Gemma model can infer the cognitive functions associated with EEG data, and subsequently predict the fMRI data from these cognitive functions. Although the observed effects are much smaller for the foundation models than for the classical machine learning and deep learning models, the possibility to leverage LLMs and LMMs for this task, and potentially to integrate the two strategies by fine-tuning foundation models on existing EEG-fMRI datasets, could open new horizons for EEG-to-fMRI prediction. As more advanced and general LLMs and LMMs continue to be developed, these models could become increasingly important tools for enhancing neural interfaces and advancing toward a multimodal foundation model for neuroscience.

### Reproducibility

The experiments in this research are based on the publicly available EEG-fMRI NF dataset *A multi-modal human neuroimaging dataset for data integration: simultaneous EEG and fMRI acquisition during a motor imagery neurofeedback task: XP1*, released under the CC0 license and accessible via OpenNeuro at https://openneuro.org/datasets/ds002336/versions/2.0.2. The Gemma, Llama, and PaliGemma models are also publicly available. The code is provided in fully commented Jupyter Notebooks (*Supplementary Notebooks 1-14*), designed to be run in a Conda environment specified by *Supplementary ENV*.*yml*. The fMRI preprocessing using fMRIPrep is documented in *Supplementary README*.*md*, and must be completed before running the experiments. On a standard personal computer (e.g., MacBook Pro), the fMRIPrep preprocessing takes around 24-36 hours, while the full execution of the code requires an additional 4-6 hours.

### Societal impact

This research does not introduce a new deployable system or asset, and is not expected to have an immediate societal impact. However, by contributing to the long-term objective of EEG-to-fMRI prediction, it could eventually support the development of enhanced neural interfaces, designed to achieve near-fMRI precision while retaining the affordability and wearability of EEG devices. While such neural interfaces remain hypothetical, the possibility to monitor cognitive processes at scale could have a profound impact in many domains, and would naturally require careful reflection and evaluation, considering the sensitive nature of human brain data.

## Additional Declarations

No competing interests reported.

## Notes

https://github.com/maeldonoso/neuropolis-x1

### Competing Interest Statement

The authors have declared no competing interest.

