## Supplementary material for "Neural Networks and Foundation Models: Two Strategies for EEG-to-fMRI Prediction": 01_fMRI_Preprocessing.ipynb

```
# %% [markdown]
# **Neural Networks and Foundation Models: Two Strategies for EEG-to-fMRI Prediction**
### **Supplementary Material**

# %% [markdown]
This project includes fourteen Notebooks:

***01: fMRI Preprocessing***

*02: EEG Preprocessing*

*03: Classification Models*

*04: Regression Models*

*05: Neural Networks*

*06: Convolutional Neural Networks*

*07: Recurrent Neural Networks*

*08: Transformers*

*09: Large Language Models*

*10: Large Language Model Chain-of-Thought*

*11: Large Language Model Fine-Tuning*

*12: Large Multimodal Model*

*13: Statistical Tests*

*14: Tables and Figures*

# %% [markdown]
### Notebook 01

# **fMRI Preprocessing**

# %% [markdown]
## **1. fMRI Data Extraction**

We **extract the fMRI data** from the brain images obtained after the fMRIPrep preprocessing step, using the neuroimaging library NiBabel. From each brain image, we extract a 4D NumPy array with dimensions (76, 95, 40, 210), where the fourth dimension corresponds to the number of scans, and the first three dimensions correspond to the three standard neurological views: sagittal (x), coronal (y), axial (z). In this 4D NumPy array, each value corresponds to the BOLD (Blood Oxygen Level Dependent) signal in a particular voxel of the brain, and for a particular scan. We plot some brain images through the middle section of the sagittal, coronal, and axial views, before and after applying the mask provided by fMRIPrep. 

# %% [code]
import numpy as np
import pandas as pd
import matplotlib.pyplot as plt
%matplotlib inline
import nibabel as nib
from nilearn import image, datasets
import os
import pickle

# %% [code]
# Define the fmriprep data path. 
fmriprep_data_path = '../fmriprep_data/'

# Define the preprocessed data path. 
preprocessed_data_path = '../preprocessed_data/'

# Define the results path. 
results_path = '../results/'

# %% [code]
# Define the list of subjects. 
subjects = ['sub-xp1' + str(x).zfill(2) for x in range(1, 11)]
print(subjects)

# %% [code]
# Define an example subject. 
subject = subjects[0]
subject

# %% [code]
# Load a NIFTI image using the NiBabel library, and display its size. 
img = nib.load(fmriprep_data_path + subject + '/func/' +  subject + '_task-eegfmriNF_space-MNI152Lin_desc-preproc_bold.nii.gz')
img.shape

# %% [code]
# Extract the image data, and display its size. 
image_data = img.get_fdata()
image_data.shape

# %% [code]
# Display the number of scans. 
nb_scans = image_data.shape[3]
nb_scans

# %% [code]
# Check that the image data is a NumPy array. 
type(image_data) == np.ndarray

# %% [code]
# Check that the image data is in float64 format. 
image_data.dtype == np.dtype(np.float64)

# %% [code]
# Display the total number of values for one 3D volume. 
np.prod(image_data[:, :, :, 0].shape)

# %% [code]
# Display the memory usage. 
print('Image size:', str(image_data.nbytes), 'bytes')

# %% [code]
# Display the metadata of the image. 
print(img.header)

# %% [code]
# Define a function to return the middle sections of an image data array. 
def middle_sections(image_data):
    return tuple((np.array(image_data[:, :, :].shape) / 2).astype(int))

# %% [code]
# Find the middle sections coordinates of the image data array. 
middle_coord = middle_sections(image_data[:, :, :, 0])
middle_coord

# %% [code]
# Define a function to display sections of an image through the sagittal, coronal, and axial views, using the middle sections coordinates. 
def display_fMRI_image(image_data, middle_coord, scan = 0):
    fig, axes = plt.subplots(nrows = 1, ncols = 3, figsize = (12, 5))

    # If the image_data array contains several scans, select the scan given as argument. 
    if len(image_data.shape) == 4:
        image_data = image_data[:, :, :, scan]

    axes[0].imshow(image_data[middle_coord[0], :, :].T, origin = 'lower')
    axes[0].set_title('Sagittal view')
    axes[1].imshow(image_data[:, middle_coord[1], :].T, origin = 'lower')
    axes[1].set_title('Coronal view')
    axes[2].imshow(image_data[:, :, middle_coord[2]], origin = 'lower')
    axes[2].set_title('Axial view');

# %% [code]
# Display sections of an image through the sagittal, coronal, and axial views, using the middle sections coordinates. 
display_fMRI_image(image_data, middle_coord)

# %% [code]
# Load the mask using the NiBabel library, and display its size. 
mask_img = nib.load(fmriprep_data_path + subject + '/func/' +  subject + '_task-eegfmriNF_space-MNI152Lin_desc-brain_mask.nii.gz')
mask_img.shape

# %% [code]
# Extract the image data, and display its size. 
mask_data = mask_img.get_fdata()
mask_data.shape

# %% [code]
# Display sections of an image through the sagittal, coronal, and axial views, using the middle sections coordinates. 
display_fMRI_image(mask_data, middle_coord)

# %% [code]
# Convert to Boolean. 
binary_mask_3d = mask_data.astype(bool)

# Convert the 3D binary mask into a 4D binary mask, by repeating the Boolean values along a 4th dimension. 
binary_mask_4d = np.repeat(binary_mask_3d[:, :, :, np.newaxis], nb_scans, axis = 3)
binary_mask_4d.shape

# %% [code]
# Transform the image data according to the binary mask. 
masked_image_data = image_data.copy()
masked_image_data[~binary_mask_4d] = np.nan
masked_image_data.shape

# %% [code]
# Display sections of an image through the sagittal, coronal, and axial views, using the middle sections coordinates. 
display_fMRI_image(masked_image_data, middle_coord)

# %% [code]
# Display the proportion of NaN values for all the scans. 
np.sum(np.isnan(masked_image_data)) / np.prod(masked_image_data.shape)

# %% [code]
# Display the difference between the two first scans, and between the first and the last scans. 
fig, axes = plt.subplots(nrows = 1, ncols = 2, figsize = (8, 5))
axes[0].imshow(masked_image_data[:, :, middle_coord[2], 0] - masked_image_data[:, :, middle_coord[2], 1], 
           origin = 'lower', cmap = plt.get_cmap('hot'))
axes[0].set_title('First scan - Second scan')
axes[1].imshow(masked_image_data[:, :, middle_coord[2], 0] - masked_image_data[:, :, middle_coord[2], (nb_scans - 1)], 
           origin = 'lower', cmap = plt.get_cmap('hot'))
axes[1].set_title('First scan - Last scan');

# %% [markdown]
## **2. Exploratory Data Analysis (EDA)**

We **explore the fMRI data** by comparing several brain images, and exploring the distribution of voxel values between subjects, between conditions, and through the three neurological views, focusing on our three conditions of interest: eegNF, fmriNF, eegfmriNF. Since the brain images are normalized using the MNI template, a (xi, yi, zi) voxel from one subject and one condition is comparable to a (xi, yi, zi) voxel of another subject or another condition, and we can perform elementwise comparisons. The fMRI data seems of good quality, with no obvious outlier. Nevertheless, we observe that the eegNF run is slightly shorter for subject sub-xp101, and missing for subject sub-xp102. 

# %% [code]
# Define a function to extract the masked image data for a particular subject, condition, and scan. 
def extract_masked_image_data(subject, condition, scan = 0):
    
    # Load the mask, extract the mask data, and convert to Boolean. 
    mask_img = nib.load(fmriprep_data_path + subject + '/func/' + subject + '_task-' + condition + '_space-MNI152Lin_desc-brain_mask.nii.gz')
    mask_data = mask_img.get_fdata()
    binary_mask_3d = mask_data.astype(bool)
    
    # Load an image, extract the image data, select a scan, and apply the mask. 
    img = nib.load(fmriprep_data_path + subject + '/func/' + subject + '_task-' + condition + '_space-MNI152Lin_desc-preproc_bold.nii.gz')
    image_data = img.get_fdata()
    masked_image_data = image_data[:, :, :, scan]
    masked_image_data[~binary_mask_3d] = np.nan

    # Return the masked image data. 
    return masked_image_data

# %% [code]
# Compare the 3D images between all subjects. 
fig, axes = plt.subplots(nrows = 4, ncols = 3, figsize = (12, 14))
fig.suptitle('All subjects', fontsize = 14, y = 0.92)
subject_index = 0

# Iterate through all subjects. 
for subject in subjects:

    # Extract the masked image data for the eegfmriNF condition and the first scan. 
    masked_image_data = extract_masked_image_data(subject, 'eegfmriNF')
    
    # Compute the row and column indexes. 
    row_index = subject_index // 3
    column_index = subject_index % 3
    
    # Display sections of an image through the axial view, using the middle sections coordinates. 
    axes[row_index, column_index].imshow(masked_image_data[:, :, middle_coord[2]], origin = 'lower')
    axes[row_index, column_index].set_title(subject);
    
    # Increment. 
    subject_index += 1

# Hide empty subplots. 
axes[3, 1].set_visible(False)
axes[3, 2].set_visible(False)

# %% [code]
# Compare the 3D images for the same subject between all conditions. 
fig, axes = plt.subplots(nrows = 2, ncols = 3, figsize = (12, 7))
fig.suptitle('All conditions', fontsize = 14, y = 0.95)
condition_index = 0

# Iterate through all conditions. 
for condition in ['eegNF', 'fmriNF', 'eegfmriNF', 'MIpre', 'MIpost', 'motorloc']:

    # Extract the masked image data for the third subject (for the first subject, MIpre and MIpost are missing) and the first scan. 
    masked_image_data = extract_masked_image_data('sub-xp103', condition)
    
    # Compute the row and column indexes. 
    row_index = condition_index // 3
    column_index = condition_index % 3
    
    # Display sections of an image through the axial view, using the middle sections coordinates. 
    axes[row_index, column_index].imshow(masked_image_data[:, :, middle_coord[2]], origin = 'lower')
    axes[row_index, column_index].set_title(condition);
    
    # Increment. 
    condition_index += 1

# %% [code]
# Load the brain images of interest and store them in dictionaries. 
dict_image_data = dict()
dict_masked_image_data = dict()
df_summary = pd.DataFrame()

# Iterate through all subjects. 
for subject in subjects:

    # Define the subject dictionaries. 
    dict_image_data[subject] = dict()
    dict_masked_image_data[subject] = dict()

    # Iterate through all conditions. 
    for condition in ['eegNF', 'fmriNF', 'eegfmriNF']:

        # Skip the current iteration for sub-xp102 and eegNF, since that run is missing. 
        if subject == 'sub-xp102' and condition == 'eegNF':
            df_summary.loc[subject, condition] = 'Missing'
            continue

        # Load the mask, extract the mask data, and convert to Boolean. 
        mask_img = nib.load(fmriprep_data_path + subject + '/func/' + subject + '_task-' + condition + '_space-MNI152Lin_desc-brain_mask.nii.gz')
        mask_data = mask_img.get_fdata()
        binary_mask_3d = mask_data.astype(bool)
        
        # Load an image, extract the image data, store the shape in the DataFrame, select the first scan, and apply the mask. 
        img = nib.load(fmriprep_data_path + subject + '/func/' + subject + '_task-' + condition + '_space-MNI152Lin_desc-preproc_bold.nii.gz')
        image_data = img.get_fdata()
        df_summary.loc[subject, condition] = str(image_data.shape)
        masked_image_data = image_data[:, :, :, 0].copy()
        masked_image_data[~binary_mask_3d] = np.nan

        # Store the images. 
        dict_image_data[subject][condition] = image_data[:, :, :, 0]
        dict_masked_image_data[subject][condition] = masked_image_data

# %% [code]
# Display the summary DataFrame. 
df_summary

# %% [code]
# Display the differences between subjects. 
fig, axes = plt.subplots(nrows = 10, ncols = 10, figsize = (12, 14))
fig.suptitle('Differences between subjects', fontsize = 14, y = 0.92)
subject_x_index = 0

# Iterate through all subjects. 
for subject_x in subjects:
    subject_y_index = 0

    # Iterate through all subjects. 
    for subject_y in subjects:
    
        # Retrieve the masked image data for the first scan. 
        masked_image_data_x = dict_masked_image_data[subject_x]['eegfmriNF']
        masked_image_data_y = dict_masked_image_data[subject_y]['eegfmriNF']

        # Display the differences between the two masked images. 
        axes[subject_x_index, subject_y_index].imshow(masked_image_data_x[:, :, middle_coord[2]] - masked_image_data_y[:, :, middle_coord[2]], 
                                                      origin = 'lower', cmap = plt.get_cmap('summer'))
        axes[subject_x_index, subject_y_index].set_title(str(subject_x_index + 1) + ' - ' + str(subject_y_index + 1))
        axes[subject_x_index, subject_y_index].tick_params(left = False, right = False, labelleft = False, 
                                                           labelbottom = False, bottom = False) 

        # Increment the subject index. 
        subject_y_index += 1

    # Increment the subject index. 
    subject_x_index += 1

# %% [code]
# Display the differences between conditions. 
fig, axes = plt.subplots(nrows = 3, ncols = 3, figsize = (12, 10))
fig.suptitle('Differences between conditions', fontsize = 14, y = 0.95)
condition_x_index = 0

# Iterate through all conditions. 
for condition_x in ['eegNF', 'fmriNF', 'eegfmriNF']:
    condition_y_index = 0

    # Iterate through all conditions. 
    for condition_y in ['eegNF', 'fmriNF', 'eegfmriNF']:
    
        # Retrieve the masked image data for the first scan. 
        masked_image_data_x = dict_masked_image_data['sub-xp101'][condition_x]
        masked_image_data_y = dict_masked_image_data['sub-xp101'][condition_y]

        # Display the differences between the two masked images. 
        axes[condition_x_index, condition_y_index].imshow(masked_image_data_x[:, :, middle_coord[2]] - masked_image_data_y[:, :, middle_coord[2]], 
                                                          origin = 'lower', cmap = plt.get_cmap('summer'))
        axes[condition_x_index, condition_y_index].set_title(condition_x + ' - ' + condition_y)
        axes[condition_x_index, condition_y_index].tick_params(left = False, right = False, labelleft = False, 
                                                               labelbottom = False, bottom = False) 

        # Increment the subject index. 
        condition_y_index += 1

    # Increment the subject index. 
    condition_x_index += 1

# %% [code]
# Display histograms and boxplots, differentiating by subject. 
fig, axes = plt.subplots(nrows = 10, ncols = 2, figsize = (12, 45))
fig.suptitle('Distribution of BOLD values across subjects', fontsize = 14, y = 0.9)
subject_index = 0

# Iterate through all subjects. 
for subject in subjects:
    condition_index = 0 

    # Iterate through all conditions. 
    for condition in ['eegNF', 'fmriNF', 'eegfmriNF']:

        # Skip the current iteration for sub-xp102 and eegNF, since that run is missing. 
        if subject == 'sub-xp102' and condition == 'eegNF':
            condition_index += 1
            continue

        # Retrieve the masked image data for the first scan. 
        masked_image_data = dict_masked_image_data[subject][condition]
        
        # Flatten the image and remove the NaN values. 
        flatten_image = masked_image_data.flatten()
        flatten_image = flatten_image[~np.isnan(flatten_image)]
        
        # Define colormap and select a color for the condition. 
        cm = plt.colormaps['Blues']
        c = cm(0.25 * condition_index + 0.25)
        
        # Display the data and set titles. 
        axes[subject_index, 0].hist(flatten_image, bins = 30, alpha = 0.5, color = c)
        axes[subject_index, 0].set_title(subject + ': all conditions')
        
        axes[subject_index, 1].boxplot(flatten_image, positions = [condition_index + 1], patch_artist = True,
                                       boxprops = dict(facecolor = c, color = c),
                                       capprops = dict(color = c),
                                       whiskerprops = dict(color = c),
                                       flierprops = dict(color = c, markeredgecolor = c),
                                       medianprops = dict(color = c),
                                       )
        axes[subject_index, 1].set_xlim(0, 4)
        axes[subject_index, 1].set_xticks([1, 2, 3])
        axes[subject_index, 1].set_xticklabels(['eegNF', 'fmriNF', 'eegfmriNF'])
        axes[subject_index, 1].set_title(subject + ': every condition')

        # Increment the condition index. 
        condition_index += 1

    # Increment the subject index. 
    subject_index += 1

# %% [code]
# Display histograms and boxplots, differentiating by condition. 
fig, axes = plt.subplots(nrows = 3, ncols = 2, figsize = (12, 13), gridspec_kw = {'width_ratios': [1, 1.5]})
fig.suptitle('Distribution of BOLD values across conditions', fontsize = 14, y = 0.95)
subject_index = 0

# Iterate through all subjects. 
for subject in subjects:
    condition_index = 0
    
    # Iterate through all conditions. 
    for condition in ['eegNF', 'fmriNF', 'eegfmriNF']:
        
        # Skip the current iteration for sub-xp102 and eegNF, since that run is missing. 
        if subject == 'sub-xp102' and condition == 'eegNF':
            condition_index += 1
            continue
        
        # Retrieve the masked image data for the first scan. 
        masked_image_data = dict_masked_image_data[subject][condition]
        
        # Flatten the image and remove the NaN values. 
        flatten_image = masked_image_data.flatten()
        flatten_image = flatten_image[~np.isnan(flatten_image)]
        
        # Define colormap and select a color for the subject. 
        cm = plt.colormaps['Blues']
        c = cm(0.07 * subject_index + 0.07)
        
        # Display the data and set titles. 
        axes[condition_index, 0].hist(flatten_image, bins = 30, alpha = 0.5, color = c)
        axes[condition_index, 0].set_title('All subjects: ' + condition)
        
        axes[condition_index, 1].boxplot(flatten_image, positions = [subject_index + 1], patch_artist = True,
                                         boxprops = dict(facecolor = c, color = c),
                                         capprops = dict(color = c),
                                         whiskerprops = dict(color = c),
                                         flierprops = dict(color = c, markeredgecolor = c),
                                         medianprops = dict(color = c),
                                         )
        axes[condition_index, 1].set_xlim(0, 11)
        axes[condition_index, 1].set_xticks(list(range(1, 11)))
        axes[condition_index, 1].set_xticklabels(range(1, 11))
        axes[condition_index, 1].set_title('Every subject: ' + condition)
        
        # Increment the condition index. 
        condition_index += 1

    # Increment the subject index. 
    subject_index += 1

# %% [code]
# Display statistics through the three neurological views, corresponding to the same condition on several subjects. 
fig, axes = plt.subplots(nrows = 3, ncols = 3, figsize = (12, 14))
fig.suptitle('Statistics across subjects', fontsize = 14, y = 0.94)
subject_index = 0

# Iterate through all subjects. 
for subject in subjects:
    
    # Retrieve the masked image data for the first scan. 
    masked_image_data = dict_masked_image_data[subject]['eegfmriNF']
    
    # Sagittal view. 
    sagittal_nonnan_value = []
    sagittal_sum_value = []
    sagittal_var_value = []
    for sagittal_index in range(0, masked_image_data.shape[0]):
        sagittal_nonnan_value.append(np.sum(~np.isnan(masked_image_data[sagittal_index, :, :])))
        sagittal_sum_value.append(np.nansum(masked_image_data[sagittal_index, :, :]))
        # Compute the variance only when the sum is different from zero. 
        if sagittal_sum_value[-1] != 0:
            sagittal_var_value.append(np.nanvar(masked_image_data[sagittal_index, :, :]))
        else:
            sagittal_var_value.append(0)
        
    # Coronal view. 
    coronal_nonnan_value = []
    coronal_sum_value = []
    coronal_var_value = []
    for coronal_index in range(0, masked_image_data.shape[1]):
        coronal_nonnan_value.append(np.sum(~np.isnan(masked_image_data[:, coronal_index, :])))
        coronal_sum_value.append(np.nansum(masked_image_data[:, coronal_index, :]))
        # Compute the variance only when the sum is different from zero. 
        if coronal_sum_value[-1] != 0:
            coronal_var_value.append(np.nanvar(masked_image_data[:, coronal_index, :]))
        else:
            coronal_var_value.append(0)
        
    # Axial view. 
    axial_nonnan_value = []
    axial_sum_value = []
    axial_var_value = []
    for axial_index in range(0, masked_image_data.shape[2]):
        axial_nonnan_value.append(np.sum(~np.isnan(masked_image_data[:, :, axial_index])))
        axial_sum_value.append(np.nansum(masked_image_data[:, :, axial_index]))
        # Compute the variance only when the sum is different from zero. 
        if axial_sum_value[-1] != 0:
            axial_var_value.append(np.nanvar(masked_image_data[:, :, axial_index]))
        else:
            axial_var_value.append(0)
        
    # Define colormap and select a color for the subject. 
    cm = plt.colormaps['Blues']
    c = cm(0.07 * subject_index + 0.07)
        
    # Display the data. 
    axes[0, 0].plot(sagittal_nonnan_value, color = c)
    axes[0, 1].plot(coronal_nonnan_value, color = c)
    axes[0, 2].plot(axial_nonnan_value, color = c)
    axes[1, 0].plot(sagittal_sum_value, color = c)
    axes[1, 1].plot(coronal_sum_value, color = c)
    axes[1, 2].plot(axial_sum_value, color = c)
    axes[2, 0].plot(sagittal_var_value, color = c)
    axes[2, 1].plot(coronal_var_value, color = c)
    axes[2, 2].plot(axial_var_value, color = c)

    # Use scientific notation for the variance graphs. 
    axes[2, 0].ticklabel_format(axis = 'y', style = 'sci', scilimits = (0, 0))
    axes[2, 1].ticklabel_format(axis = 'y', style = 'sci', scilimits = (0, 0))
    axes[2, 2].ticklabel_format(axis = 'y', style = 'sci', scilimits = (0, 0))
    
    # Set titles. 
    axes[0, 0].set_title('Sagittal sections: non-nan values')
    axes[0, 1].set_title('Coronal sections: non-nan values')
    axes[0, 2].set_title('Axial sections: non-nan values')
    axes[1, 0].set_title('Sagittal sections: sum of values')
    axes[1, 1].set_title('Coronal sections: sum of values')
    axes[1, 2].set_title('Axial sections: sum of values')
    axes[2, 0].set_title('Sagittal sections: variance of values')
    axes[2, 1].set_title('Coronal sections: variance of values')
    axes[2, 2].set_title('Axial sections: variance of values')

    # Increment the subject index. 
    subject_index += 1

# %% [code]
# Display statistics through the three neurological views, corresponding to several conditions on the same subject. 
fig, axes = plt.subplots(nrows = 3, ncols = 3, figsize = (12, 14))
fig.suptitle('Statistics across conditions', fontsize = 14, y = 0.94)
condition_index = 0

# Iterate through all conditions. 
for condition in ['eegNF', 'fmriNF', 'eegfmriNF']:
    
    # Retrieve the masked image data for the first scan. 
    masked_image_data = dict_masked_image_data['sub-xp101'][condition]
    
    # Sagittal view. 
    sagittal_nonnan_value = []
    sagittal_sum_value = []
    sagittal_var_value = []
    for sagittal_index in range(0, masked_image_data.shape[0]):
        sagittal_nonnan_value.append(np.sum(~np.isnan(masked_image_data[sagittal_index, :, :])))
        sagittal_sum_value.append(np.nansum(masked_image_data[sagittal_index, :, :]))
        # Compute the variance only when the sum is different from zero. 
        if sagittal_sum_value[-1] != 0:
            sagittal_var_value.append(np.nanvar(masked_image_data[sagittal_index, :, :]))
        else:
            sagittal_var_value.append(0)
        
    # Coronal view. 
    coronal_nonnan_value = []
    coronal_sum_value = []
    coronal_var_value = []
    for coronal_index in range(0, masked_image_data.shape[1]):
        coronal_nonnan_value.append(np.sum(~np.isnan(masked_image_data[:, coronal_index, :])))
        coronal_sum_value.append(np.nansum(masked_image_data[:, coronal_index, :]))
        # Compute the variance only when the sum is different from zero. 
        if coronal_sum_value[-1] != 0:
            coronal_var_value.append(np.nanvar(masked_image_data[:, coronal_index, :]))
        else:
            coronal_var_value.append(0)
        
    # Axial view. 
    axial_nonnan_value = []
    axial_sum_value = []
    axial_var_value = []
    for axial_index in range(0, masked_image_data.shape[2]):
        axial_nonnan_value.append(np.sum(~np.isnan(masked_image_data[:, :, axial_index])))
        axial_sum_value.append(np.nansum(masked_image_data[:, :, axial_index]))
        # Compute the variance only when the sum is different from zero. 
        if axial_sum_value[-1] != 0:
            axial_var_value.append(np.nanvar(masked_image_data[:, :, axial_index]))
        else:
            axial_var_value.append(0)
        
    # Define colormap and select a color for the subject. 
    cm = plt.colormaps['Blues']
    c = cm(0.25 * condition_index + 0.25)
        
    # Display the data. 
    axes[0, 0].plot(sagittal_nonnan_value, color = c)
    axes[0, 1].plot(coronal_nonnan_value, color = c)
    axes[0, 2].plot(axial_nonnan_value, color = c)
    axes[1, 0].plot(sagittal_sum_value, color = c)
    axes[1, 1].plot(coronal_sum_value, color = c)
    axes[1, 2].plot(axial_sum_value, color = c)
    axes[2, 0].plot(sagittal_var_value, color = c)
    axes[2, 1].plot(coronal_var_value, color = c)
    axes[2, 2].plot(axial_var_value, color = c)

    # Use scientific notation for the variance graphs. 
    axes[2, 0].ticklabel_format(axis = 'y', style = 'sci', scilimits = (0, 0))
    axes[2, 1].ticklabel_format(axis = 'y', style = 'sci', scilimits = (0, 0))
    axes[2, 2].ticklabel_format(axis = 'y', style = 'sci', scilimits = (0, 0))
    
    # Set titles. 
    axes[0, 0].set_title('Sagittal sections: non-nan values')
    axes[0, 1].set_title('Coronal sections: non-nan values')
    axes[0, 2].set_title('Axial sections: non-nan values')
    axes[1, 0].set_title('Sagittal sections: sum of values')
    axes[1, 1].set_title('Coronal sections: sum of values')
    axes[1, 2].set_title('Axial sections: sum of values')
    axes[2, 0].set_title('Sagittal sections: variance of values')
    axes[2, 1].set_title('Coronal sections: variance of values')
    axes[2, 2].set_title('Axial sections: variance of values')

    # Increment the condition index. 
    condition_index += 1

# %% [markdown]
## **3. Targets Extraction**

We **extract the targets** by averaging the voxel values within each brain region defined in the Harvard-Oxford cortical atlas. We use the maximum probability map with no threshold, meaning that each voxel is assigned to the region with the highest probability, ensuring full cortical coverage. 

# %% [code]
# Load and display informations about the Harvard-Oxford atlas for cortical regions. 
atlas = datasets.fetch_atlas_harvard_oxford('cort-maxprob-thr0-2mm', data_dir = None, symmetric_split = False, resume = True, verbose = 1)
atlas

# %% [code]
# Extract the atlas as a Nifti image. 
atlas_img = image.load_img(atlas.maps)

# Extract the atlas data, and display its size. 
atlas_data = atlas.maps.get_fdata()
atlas_data.shape

# %% [code]
# Find the middle sections coordinates of the atlas data array. 
middle_coord = middle_sections(atlas_data[:, :, :])
middle_coord

# %% [code]
# Display sections of the atlas through the sagittal, coronal, and axial views, using the middle sections coordinates. 
display_fMRI_image(atlas_data, middle_coord)

# %% [code]
# Retrieve and display the number of voxels for each brain region. 
unique, counts = np.unique(atlas_data, return_counts = True)
dict(zip(atlas.labels, counts))

# %% [code]
# Define an example subject. 
subject = subjects[0]
subject

# %% [code]
# Load a NIFTI image using the NiBabel library, and display its size. 
img = nib.load(fmriprep_data_path + subject + '/func/' +  subject + '_task-eegfmriNF_space-MNI152Lin_desc-preproc_bold.nii.gz')
img.shape

# %% [code]
# Resample the atlas to match the fMRI image dimensions. 
resampled_atlas = image.resample_to_img(atlas_img, img, interpolation = 'nearest', force_resample = True, copy_header = True)

# Extract the image data, and display its size. 
resampled_atlas_data = resampled_atlas.get_fdata()
resampled_atlas_data.shape

# %% [code]
# Find the middle sections coordinates of the image data array. 
middle_coord = middle_sections(resampled_atlas_data[:, :, :])
middle_coord

# %% [code]
# Display sections of the atlas through the sagittal, coronal, and axial views, using the middle sections coordinates. 
display_fMRI_image(resampled_atlas_data, middle_coord)

# %% [code]
# Store the first voxel of each brain region in a dictionary. 
dict_first_voxel_brain_region = dict()

# Iterate through all brain regions.
for brain_region_index in range(len(atlas['labels'])):

    # Display the brain region name. 
    print(str(brain_region_index) + ': ' + atlas['labels'][brain_region_index])

    # Find the first voxel of the brain region.
    brain_region_voxels = np.where(resampled_atlas_data == brain_region_index)
    combined_array = np.vstack((brain_region_voxels[0], brain_region_voxels[1], brain_region_voxels[2])).T
    brain_region_voxel = combined_array[0]

    # Store the first voxel of the brain region.
    dict_first_voxel_brain_region[brain_region_index] = brain_region_voxel
    print(brain_region_voxel)

# %% [code]
# Define a function to display the position of a voxel, with one image as background. 
def display_voxel_position(voxel_index, background_image, figure_title = ''):
    fig, axes = plt.subplots(nrows = 1, ncols = 3, figsize = (12, 4))
    fig.suptitle(figure_title, fontsize = 14, y = 0.85)

    # Sagittal view. 
    axes[0].imshow(background_image[voxel_index[0], :, :].T, origin = 'lower')
    axes[0].axhline(voxel_index[2], 0, background_image.shape[1], color = 'yellow') # Horizontal line through coronal view. 
    axes[0].axvline(voxel_index[1], 0, background_image.shape[2], color = 'yellow') # Vertical line through axial view. 
    axes[0].scatter(voxel_index[1], voxel_index[2], marker = 'o', c = 'white')
    axes[0].set_title('Sagittal view')

    # Coronal view. 
    axes[1].imshow(background_image[:, voxel_index[1], :].T, origin = 'lower')
    axes[1].axhline(voxel_index[2], 0, background_image.shape[0], color = 'yellow') # Horizontal line through sagittal view. 
    axes[1].axvline(voxel_index[0], 0, background_image.shape[2], color = 'yellow') # Vertical line through axial view. 
    axes[1].scatter(voxel_index[0], voxel_index[2], marker = 'o', c = 'white')
    axes[1].set_title('Coronal view')

    # Axial view. 
    axes[2].imshow(background_image[:, :, voxel_index[2]], origin = 'lower')
    axes[2].axhline(voxel_index[0], 0, background_image.shape[1], color = 'yellow') # Horizontal line through coronal view. 
    axes[2].axvline(voxel_index[1], 0, background_image.shape[0], color = 'yellow') # Vertical line through sagittal view. 
    axes[2].scatter(voxel_index[1], voxel_index[0], marker = 'o', c = 'white')
    axes[2].set_title('Axial view')
    plt.show();

# %% [code]
# Iterate through all brain regions.
for brain_region_index in range(len(atlas['labels'])):

    # Retrieve the position of the voxel. 
    first_voxel_position = dict_first_voxel_brain_region[brain_region_index]
    
    # Retrieve the atlas data as the background. 
    background_image = resampled_atlas_data

    # Display the position of the voxel. 
    display_voxel_position(first_voxel_position, background_image, figure_title = atlas['labels'][brain_region_index])
    

# %% [code]
# Define the number of scans and extract the target values. 
nb_scans = 200
dict_targets_values = dict()
dict_mean_values = dict()
dict_mean_values_inside_brain = dict()
dict_mean_values_outside_brain = dict()
multi_index = pd.MultiIndex.from_product([subjects, ['eegNF', 'fmriNF', 'eegfmriNF']], names = ['subject', 'condition'])
pd_proportion_voxels_inside_brain = pd.DataFrame(index = multi_index)

# Extract the atlas as a Nifti image. 
atlas_img = image.load_img(atlas.maps)

# Iterate through all subjects. 
for subject in subjects:

    # Define the subject dictionaries. 
    dict_targets_values[subject] = dict()
    dict_mean_values[subject] = dict()
    dict_mean_values_inside_brain[subject] = dict()
    dict_mean_values_outside_brain[subject] = dict()

    # Iterate through all conditions. 
    for condition in ['eegNF', 'fmriNF', 'eegfmriNF']:

        # Skip the current iteration for sub-xp102 and eegNF, since that run is missing. 
        if subject == 'sub-xp102' and condition == 'eegNF':
            continue

        # Load the image and extract the image data. 
        img = nib.load(fmriprep_data_path + subject + '/func/' + subject + '_task-' + condition + '_space-MNI152Lin_desc-preproc_bold.nii.gz')
        image_data = img.get_fdata()

        # Discard the first scan, as indicated in the article describing the dataset, and take the nb_scans following scans. 
        image_data = image_data[:, :, :, 1:(nb_scans + 1)] 
        
        # Load the mask, extract the mask data, convert to Boolean, and convert the 3D binary mask into a 4D binary mask. 
        mask_img = nib.load(fmriprep_data_path + subject + '/func/' + subject + '_task-' + condition + '_space-MNI152Lin_desc-brain_mask.nii.gz')
        mask_data = mask_img.get_fdata()
        binary_mask_3d = mask_data.astype(bool)
        binary_mask_4d = np.repeat(binary_mask_3d[:, :, :, np.newaxis], nb_scans, axis = 3)

        # Apply both the normal mask and the inverted mask. 
        image_data_inside_brain = image_data.copy()
        image_data_inside_brain[~binary_mask_4d] = np.nan
        image_data_outside_brain = image_data.copy()
        image_data_outside_brain[binary_mask_4d] = np.nan

        # Store the mean image values. 
        dict_mean_values[subject][condition] = np.mean(image_data, axis = (0, 1, 2))
        dict_mean_values_inside_brain[subject][condition] = np.nanmean(image_data_inside_brain, axis = (0, 1, 2))
        dict_mean_values_outside_brain[subject][condition] = np.nanmean(image_data_outside_brain, axis = (0, 1, 2))
        
        # Resample the atlas to match the fMRI image dimensions. 
        resampled_atlas = image.resample_to_img(atlas_img, img, interpolation = 'nearest', force_resample = True, copy_header = True)

        # Define the condition dictionary. 
        dict_targets_values[subject][condition] = dict()

        # Iterate through all brain regions.
        for brain_region_index in range(len(atlas['labels'])):

            # Extract the mean values for the current brain region.
            brain_region_values = np.mean(image_data[resampled_atlas_data == brain_region_index], axis = 0)

            # Store the mean values for the current brain region.
            dict_targets_values[subject][condition][brain_region_index] = brain_region_values

            # Extract the total number of voxels in the current brain region, and the number of voxels inside the brain mask.
            nb_voxels = np.sum(resampled_atlas_data == brain_region_index)
            nb_voxels_inside_brain = np.sum(np.logical_and(resampled_atlas_data == brain_region_index, binary_mask_3d))

            # Store the proportion of voxels inside the brain mask.
            pd_proportion_voxels_inside_brain.loc[(subject, condition), atlas['labels'][brain_region_index]] = nb_voxels_inside_brain / nb_voxels

# Display the proportion of voxels inside the brain mask for all subjects and conditions.
pd_proportion_voxels_inside_brain

# %% [code]
# Plot the target values by subject and condition. 
fig, axes = plt.subplots(nrows = 10, ncols = 3, figsize = (12, 38))
fig.suptitle('Target values', fontsize = 14, y = 0.9)
subject_index = 0

# Iterate through all subjects. 
for subject in subjects:
    condition_index = 0
    
    # Iterate through all conditions. 
    for condition in ['eegNF', 'fmriNF', 'eegfmriNF']:
        target_index = 0

        # Skip the current iteration for sub-xp102 and eegNF, since that run is missing. 
        if subject == 'sub-xp102' and condition == 'eegNF':
            axes[subject_index, condition_index].set_visible(False)
            condition_index += 1
            continue

        # Iterate through all brain regions.
        for brain_region_index in range(len(atlas['labels'])):
            
            # Define colormap and select a color. 
            cm = plt.colormaps['Blues']
            c = cm(0.02 * target_index + 0.05)
    
            # Retrieve and plot the target values. 
            target_values = dict_targets_values[subject][condition][brain_region_index]
            axes[subject_index, condition_index].plot(target_values, color = c)
            axes[subject_index, condition_index].set_title(subject + ' - ' + condition)

            # Increment the target index. 
            target_index += 1

        # Increment the condition index. 
        condition_index += 1

    # Increment the subject index. 
    subject_index += 1

# %% [code]
# Define an example subject and condition. 
subject = 'sub-xp101'
condition = 'eegfmriNF'

# Plot the targets values. 
plt.figure(figsize = (12, 5)) 
target_index = 0

# Iterate through all brain regions.
for brain_region_index in range(len(atlas['labels'])):
    
    # Define colormap and select a color. 
    cm = plt.colormaps['Blues']
    c = cm(0.02 * target_index + 0.05)

    # Retrieve and plot the targets values. 
    target_values = dict_targets_values[subject][condition][brain_region_index]
    plt.plot(target_values, color = c, label = brain_region_index)
    plt.title(subject + ' - ' + condition)

    # Increment the target index. 
    target_index += 1

# %% [markdown]
## **4. Target Engineering**

We observe that the mean BOLD signal inside the brain increases during sessions, a well-known effect in fMRI datasets. We perform **target engineering** by removing this effect, normalizing each target, and replacing the outliers (STD > 3) by the value of the previous scan. As the baseline for the machine learning and deep learning steps, we choose the mean value of the targets, which is now zero. 

# %% [code]
# Plot the mean values by subject and condition. 
fig, axes = plt.subplots(nrows = 10, ncols = 3, figsize = (12, 38))
fig.suptitle('Mean BOLD signal', fontsize = 14, y = 0.9)
subject_index = 0

# Iterate through all subjects. 
for subject in subjects:
    condition_index = 0
    
    # Iterate through all conditions. 
    for condition in ['eegNF', 'fmriNF', 'eegfmriNF']:

        # Skip the current iteration for sub-xp102 and eegNF, since that run is missing. 
        if subject == 'sub-xp102' and condition == 'eegNF':
            axes[subject_index, condition_index].set_visible(False)
            condition_index += 1
            continue

        # Define colormap. 
        cm = plt.colormaps['viridis']
            
        # Retrieve and plot the mean voxel values. 
        mean_values = dict_mean_values[subject][condition]
        axes[subject_index, condition_index].plot(mean_values, color = cm(0.25))
        axes[subject_index, condition_index].set_title(subject + ' - ' + condition)

        # Increment the condition index. 
        condition_index += 1

    # Increment the subject index. 
    subject_index += 1

# %% [code]
# Plot the mean values by subject and condition, for the voxels inside and outside the brain, centered. 
fig, axes = plt.subplots(nrows = 10, ncols = 3, figsize = (12, 38))
fig.suptitle('Mean BOLD signal, inside and outside brain, centered', fontsize = 14, y = 0.9)
subject_index = 0

# Iterate through all subjects. 
for subject in subjects:
    condition_index = 0
    
    # Iterate through all conditions. 
    for condition in ['eegNF', 'fmriNF', 'eegfmriNF']:

        # Skip the current iteration for sub-xp102 and eegNF, since that run is missing. 
        if subject == 'sub-xp102' and condition == 'eegNF':
            axes[subject_index, condition_index].set_visible(False)
            condition_index += 1
            continue

        # Define colormap. 
        cm = plt.colormaps['viridis']
            
        # Retrieve, center and plot the mean voxel values inside the brain. 
        mean_values_inside_brain = dict_mean_values_inside_brain[subject][condition]
        mean_values_inside_brain = mean_values_inside_brain - np.mean(mean_values_inside_brain)
        axes[subject_index, condition_index].plot(mean_values_inside_brain, color = cm(0.25), label = 'inside brain')

        # Retrieve, center and plot the mean voxel values outside the brain. 
        mean_values_outside_brain = dict_mean_values_outside_brain[subject][condition]
        mean_values_outside_brain = mean_values_outside_brain - np.mean(mean_values_outside_brain)
        axes[subject_index, condition_index].plot(mean_values_outside_brain, color = cm(0.5), label = 'outside brain')
        axes[subject_index, condition_index].set_title(subject + ' - ' + condition)
        axes[subject_index, condition_index].legend()

        # Increment the condition index. 
        condition_index += 1

    # Increment the subject index. 
    subject_index += 1
    

# %% [code]
# Define a dictionary to store the regression coefficients. 
dict_regression_coefficients = dict()

# Plot the mean values by subject and condition, for the voxels inside the brain, with a regression line. 
fig, axes = plt.subplots(nrows = 10, ncols = 3, figsize = (12, 38))
fig.suptitle('Mean BOLD signal, inside brain, with a regression line', fontsize = 14, y = 0.9)
subject_index = 0

# Iterate through all subjects. 
for subject in subjects:
    condition_index = 0

    # Define the subject dictionary. 
    dict_regression_coefficients[subject] = dict()
    
    # Iterate through all conditions. 
    for condition in ['eegNF', 'fmriNF', 'eegfmriNF']:

        # Skip the current iteration for sub-xp102 and eegNF, since that run is missing. 
        if subject == 'sub-xp102' and condition == 'eegNF':
            axes[subject_index, condition_index].set_visible(False)
            condition_index += 1
            continue

        # Define colormap. 
        cm = plt.colormaps['viridis']
            
        # Retrieve and plot the mean voxel values inside the brain. 
        mean_values_inside_brain = dict_mean_values_inside_brain[subject][condition]
        axes[subject_index, condition_index].plot(mean_values_inside_brain, color = cm(0.25), label = 'inside brain')

        # Compute and plot the regression line. 
        x_values = range(mean_values_inside_brain.shape[0])
        coefficients = np.polyfit(x_values, mean_values_inside_brain, 1)
        y_values = coefficients[0] * x_values + coefficients[1]
        axes[subject_index, condition_index].plot(y_values, color = cm(0.5), label = 'regression line')
        axes[subject_index, condition_index].set_title(subject + ' - ' + condition)
        axes[subject_index, condition_index].legend()

        # Store the regression coefficients in the dictionary. 
        dict_regression_coefficients[subject][condition] = coefficients

        # Increment the condition index. 
        condition_index += 1

    # Increment the subject index. 
    subject_index += 1
    

# %% [code]
# Define a dictionary to store the corrected targets. 
dict_targets_values_corrected = dict()

# Plot the target values by subject and condition. 
fig, axes = plt.subplots(nrows = 10, ncols = 3, figsize = (12, 38))
fig.suptitle('Target values, corrected', fontsize = 14, y = 0.9)
subject_index = 0

# Iterate through all subjects. 
for subject in subjects:
    condition_index = 0

    # Define the subject dictionary. 
    dict_targets_values_corrected[subject] = dict()
    
    # Iterate through all conditions. 
    for condition in ['eegNF', 'fmriNF', 'eegfmriNF']:
        target_index = 0

        # Skip the current iteration for sub-xp102 and eegNF, since that run is missing. 
        if subject == 'sub-xp102' and condition == 'eegNF':
            axes[subject_index, condition_index].set_visible(False)
            condition_index += 1
            continue

        # Define the condition dictionary. 
        dict_targets_values_corrected[subject][condition] = dict()

        # Iterate through all brain regions.
        for brain_region_index in range(len(atlas['labels'])):

            # Compute the corrected target values. 
            target_values = dict_targets_values[subject][condition][brain_region_index]
            coefficients = dict_regression_coefficients[subject][condition]
            x_values = range(target_values.shape[0])
            y_values = coefficients[0] * x_values + coefficients[1]
            target_values_corrected = target_values - y_values
            
            # Define colormap and select a color. 
            cm = plt.colormaps['Blues']
            c = cm(0.02 * target_index + 0.05)
    
            # Plot the corrected target values. 
            axes[subject_index, condition_index].plot(target_values_corrected, color = c)
            axes[subject_index, condition_index].set_title(subject + ' - ' + condition)

            # Store the corrected target values in the dictionary. 
            dict_targets_values_corrected[subject][condition][brain_region_index] = target_values_corrected

            # Increment the target index. 
            target_index += 1

        # Increment the condition index. 
        condition_index += 1

    # Increment the subject index. 
    subject_index += 1

# %% [code]
# Define a dictionary to store the normalized targets, and a DataFrame to store the outliers. 
dict_targets_values_normalized = dict()
df_outliers = pd.DataFrame()

# Plot the target values by subject and condition. 
fig, axes = plt.subplots(nrows = 10, ncols = 3, figsize = (12, 38))
fig.suptitle('Target values, normalized', fontsize = 14, y = 0.9)
subject_index = 0

# Iterate through all subjects. 
for subject in subjects:
    condition_index = 0

    # Define the subject dictionary. 
    dict_targets_values_normalized[subject] = dict()
    
    # Iterate through all conditions. 
    for condition in ['eegNF', 'fmriNF', 'eegfmriNF']:
        target_index = 0

        # Skip the current iteration for sub-xp102 and eegNF, since that run is missing. 
        if subject == 'sub-xp102' and condition == 'eegNF':
            axes[subject_index, condition_index].set_visible(False)
            condition_index += 1
            continue

        # Define the condition dictionary. 
        dict_targets_values_normalized[subject][condition] = dict()

        # Iterate through all brain regions.
        for brain_region_index in range(len(atlas['labels'])):

            # Compute the normalized target values. 
            target_values_corrected = dict_targets_values_corrected[subject][condition][brain_region_index]
            target_values_normalized = (target_values_corrected - np.mean(target_values_corrected)) / np.std(target_values_corrected)

            # Iterate through all scans and replace the outliers (STD > 3) by the value of the previous scan. 
            outliers_counter = 0
            for scan in range(nb_scans):
                if np.abs(target_values_normalized[scan]) > 3:
                    target_values_normalized[scan] = target_values_normalized[scan - 1]
                    outliers_counter += 1
            df_outliers.loc[subject + '/' + condition, 'subject'] = subject
            df_outliers.loc[subject + '/' + condition, 'condition'] = condition
            df_outliers.loc[subject + '/' + condition, atlas['labels'][brain_region_index]] = outliers_counter
            
            # Define colormap and select a color. 
            cm = plt.colormaps['Blues']
            c = cm(0.02 * target_index + 0.05)
    
            # Plot the normalized target values. 
            axes[subject_index, condition_index].plot(target_values_normalized, color = c, alpha = 0.5)
            axes[subject_index, condition_index].set_title(subject + ' - ' + condition)

            # Store the normalized target values in the dictionary. 
            dict_targets_values_normalized[subject][condition][atlas['labels'][brain_region_index]] = target_values_normalized

            # Increment the target index. 
            target_index += 1

        # Increment the condition index. 
        condition_index += 1

    # Increment the subject index. 
    subject_index += 1

# Change the indexes and the data type of the outliers DataFrame. 
df_outliers = df_outliers.set_index(['subject', 'condition'])
df_outliers = df_outliers.astype(int)

# %% [code]
# Display the outliers DataFrame. 
df_outliers

# %% [code]
# Display the number of outliers by brain region. 
df_outliers.sum()

# %% [code]
# Display the proportion of outliers by brain region. 
print(df_outliers.sum() / (df_outliers.shape[0] * nb_scans))

# %% [code]
# Store the baseline for all subjects. 
dict_baseline = dict()

# Iterate through all subjects. 
for subject in subjects:

    # Define the subject dictionary. 
    dict_baseline[subject] = dict()

    # Iterate through all conditions. 
    for condition in ['eegNF', 'fmriNF', 'eegfmriNF']:

        # Skip the current iteration for sub-xp102 and eegNF, since that run is missing. 
        if subject == 'sub-xp102' and condition == 'eegNF':
            continue

        # Define the condition dictionary. 
        dict_baseline[subject][condition] = dict()

        # Iterate through all brain regions.
        for brain_region_index in range(len(atlas['labels'])):

            # Store the baseline, which is now zero. 
            dict_baseline[subject][condition][atlas['labels'][brain_region_index]] = np.zeros((nb_scans))

# %% [markdown]
## **5. Preprocessed Targets**

We save the **preprocessed targets** in the appropriate directory. 

# %% [code]
# Create directories for the preprocessed data of the classification and regression tasks. 
if not os.path.isdir(preprocessed_data_path + 'classification/'):
    os.mkdir(preprocessed_data_path + 'classification/')
if not os.path.isdir(preprocessed_data_path + 'regression/'):
    os.mkdir(preprocessed_data_path + 'regression/')

# Create directories for the results of the classification and regression tasks. 
if not os.path.isdir(results_path + 'classification/'):
    os.mkdir(results_path + 'classification/')
if not os.path.isdir(results_path + 'regression/'):
    os.mkdir(results_path + 'regression/')

# Save the targets into a Pickle file. 
with open(preprocessed_data_path + 'dict_targets.p', 'wb') as file:
    pickle.dump(dict_targets_values_normalized, file)

# Save the baseline into a Pickle file. 
with open(preprocessed_data_path + 'dict_baseline.p', 'wb') as file:
    pickle.dump(dict_baseline, file)
```
