## Supplementary material for "Neural Networks and Foundation Models: Two Strategies for EEG-to-fMRI Prediction": 02_EEG_Preprocessing.ipynb

```
# %% [markdown]
# **Neural Networks and Foundation Models: Two Strategies for EEG-to-fMRI Prediction**
### **Supplementary Material**

# %% [markdown]
This project includes fourteen Notebooks:

*01: fMRI Preprocessing*

***02: EEG Preprocessing***

*03: Classification Models*

*04: Regression Models*

*05: Neural Networks*

*06: Convolutional Neural Networks*

*07: Recurrent Neural Networks*

*08: Transformers*

*09: Large Language Models*

*10: Large Language Model Chain-of-Thought*

*11: Large Language Model Fine-Tuning*

*12: Large Multimodal Model*

*13: Statistical Tests*

*14: Tables and Figures*

# %% [markdown]
### Notebook 02

# **EEG Preprocessing**

# %% [markdown]
## **1. EEG Data Extraction**

We **extract the EEG data** preprocessed by the authors of the dataset, using the neurophysiological library MNE. From each file, we extract a 2D NumPy array with dimensions (64, 80200), where the first dimension corresponds to the number of channels, and the second dimension to the number of time points. In this 2D NumPy array, each value corresponds to the voltage signal recorded at a particular electrode, and for a particular time point. We plot some data from the C1 channel, and observe some typical EEG fluctuations. 

# %% [code]
import os
os.environ['OUTDATED_IGNORE'] = '1'
import numpy as np
import pandas as pd
import matplotlib.pyplot as plt
%matplotlib inline
import mne
import yasa
import pickle

# %% [code]
# Define the raw data path. 
raw_data_path = '../ds002336-download/'

# Define the preprocessed data path. 
preprocessed_data_path = '../preprocessed_data/'

# %% [code]
# Define the list of subjects. 
subjects = ['sub-xp1' + str(x).zfill(2) for x in range(1, 11)]
print(subjects)

# %% [code]
# Define an example subject. 
subject = subjects[0]
subject

# %% [code]
# Define the path of the header file (.vhdr) for this subject, for the eegfmriNF condition. 
vhdr_path = raw_data_path + 'derivatives/' + subject + '/eeg_pp/' + subject + '_task-eegfmriNF_eeg_pp.vhdr'

# Load the EEG data using the MNE library, keeping error messages but suppressing warnings. 
channels_data = mne.io.read_raw_brainvision(vhdr_path, misc = ['ECG'], preload = True, verbose = 'ERROR')

# %% [code]
# Display the number of channels. 
len(channels_data.ch_names)

# %% [code]
# Display the channel names. 
print(channels_data.ch_names)

# %% [code]
# Identify the C1 channel index. 
c1_channel_index = channels_data.ch_names.index('C1')
c1_channel_index

# %% [code]
# Display information about the EEG data. 
channels_data.info

# %% [code]
# Retrieve the sampling frequency. 
sampling_frequency = channels_data.info['sfreq']
sampling_frequency

# %% [code]
# Display the time points. 
channels_data.times

# %% [code]
# Display the number of time points. 
channels_data.times.shape[0]

# %% [code]
# Compute the total length in seconds. 
channels_data.times.shape[0] / sampling_frequency

# %% [code]
# Check that the data is in float64 format. 
channels_data.times.dtype == np.dtype(np.float64)

# %% [code]
# Compute the number of unique time differences between two consecutive onsets, rounding up to the 10th decimal. 
time_differences = np.unique(np.unique(channels_data.times[1:] - channels_data.times[:-1]).round(10))
time_differences

# %% [code]
# Extract the EEG data as a NumPy array. 
np_channels_data = channels_data.get_data()
np_channels_data

# %% [code]
# Check that this is indeed a NumPy array. 
type(np_channels_data)

# %% [code]
# Display the size of the array. 
np_channels_data.shape

# %% [code]
# Plot the data for the C1 channel. 
plt.plot(np_channels_data[c1_channel_index, :], c = 'darkblue')
plt.title('C1 channel');

# %% [code]
# Define an arbitrary time interval, and compute the time points for this interval. 
time_interval = [10, 12]
starting_point, ending_point = int(sampling_frequency * time_interval[0]), int(sampling_frequency * time_interval[1])
print(str(ending_point - starting_point) + ' time points')

# %% [code]
# Plot the data for the C1 channel, focusing on a selection of time points. 
plt.plot(np_channels_data[c1_channel_index, starting_point:ending_point])
plt.title('C1 channel: selection of time points');

# %% [markdown]
## **2. Exploratory Data Analysis (EDA)**

We **explore the EEG data** by comparing the different channels. We compute the Power Spectral Density (PSD) for a selection of time points, for frequencies between 1 and 40 Hz, using both the Welch method and the multitaper method, before computing the bandpowers for a series of frequency bands of interest: delta (1-4 Hz), theta (4-8 Hz), alpha (8-12 Hz), sigma (12-16 Hz), beta (16-30 Hz), gamma (30-40 Hz). The EEG data shows some spikes that may need outlier removal. 

# %% [code]
# Compare all channels. 
fig, axes = plt.subplots(nrows = 16, ncols = 4, figsize = (12, 40))
fig.suptitle('EEG channels', fontsize = 14, y = 0.9)
channel_index = 0

# Iterate through all channels. 
for channel in channels_data.ch_names:

    # Compute the row and column indexes. 
    row_index = channel_index // 4
    column_index = channel_index % 4
    
    # Plot the data for the channel. 
    axes[row_index, column_index].plot(np_channels_data[channel_index, :], c = 'darkblue')
    axes[row_index, column_index].set_title(channel)
    axes[row_index, column_index].tick_params(left = False, right = False, labelleft = False, 
                                              labelbottom = False, bottom = False) 

    # Increment. 
    channel_index += 1


# %% [code]
# Compare all channels, focusing on a selection of time points. 
fig, axes = plt.subplots(nrows = 16, ncols = 4, figsize = (12, 40))
fig.suptitle('EEG channels: selection of time points', fontsize = 14, y = 0.9)
channel_index = 0

# Iterate through all channels. 
for channel in channels_data.ch_names:

    # Compute the row and column indexes. 
    row_index = channel_index // 4
    column_index = channel_index % 4
    
    # Plot the data for the channel. 
    axes[row_index, column_index].plot(np_channels_data[channel_index, starting_point:ending_point])
    axes[row_index, column_index].set_title(channel)
    axes[row_index, column_index].tick_params(left = False, right = False, labelleft = False, 
                                              labelbottom = False, bottom = False) 

    # Increment. 
    channel_index += 1


# %% [code]
# Compute the Power Spectral Density (PSD) for a selection of time points, for frequencies between 1 and 40 Hz, using the Welch method. 
psd_welch = channels_data.compute_psd(tmin = time_interval[0], tmax = time_interval[1], fmin = 1, fmax = 40, picks = 'eeg', method = 'welch')
psd_welch

# %% [code]
# Identify the new C1 channel index, since the ECG channel was dropped during the PSD computation. 
c1_channel_index_psd = psd_welch.ch_names.index('C1')
c1_channel_index_psd

# %% [code]
# Extract the PSD data as a NumPy array, then store and display its shape. 
np_psd_welch = psd_welch.get_data()
nb_channels_psd = np_psd_welch.shape[0]
nb_bins_psd = np_psd_welch.shape[1]
np_psd_welch.shape

# %% [code]
# Plot the PSD data for the C1 channel. 
plt.plot(np_psd_welch[c1_channel_index_psd, :])
plt.title('PSD with Welch method');

# %% [code]
# Extract the PSD frequencies as a NumPy array, and display its shape. 
np_psd_welch_freqs = psd_welch.freqs
np_psd_welch_freqs.shape

# %% [code]
# Display the frequencies. 
np_psd_welch_freqs

# %% [code]
# Compute the Power Spectral Density (PSD) for a selection of time points, for frequencies between 1 and 40 Hz, using the multitaper method. 
psd_multitaper = channels_data.compute_psd(tmin = time_interval[0], tmax = time_interval[1], fmin = 1, fmax = 40, picks = 'eeg', method = 'multitaper')
psd_multitaper

# %% [code]
# Extract the PSD data as a NumPy array, and display its shape. 
np_psd_multitaper = psd_multitaper.get_data()
np_psd_multitaper.shape

# %% [code]
# Plot the PSD data for the C1 channel, comparing the Welch method and the multitaper method. 
fig, axes = plt.subplots(nrows = 1, ncols = 2, figsize = (12, 5))
axes[0].plot(np_psd_welch[c1_channel_index_psd, :])
axes[0].set_title('PSD with Welch method')
axes[1].plot(np_psd_multitaper[c1_channel_index_psd, :])
axes[1].set_title('PSD with multitaper method');

# %% [code]
# Define the frequency bands of interest. 
frequency_bands = [(1, 4, 'Delta'), (4, 8, 'Theta'), (8, 12, 'Alpha'), (12, 16, 'Sigma'), (16, 30, 'Beta'), (30, 40, 'Gamma')]
frequency_bands

# %% [code]
# Using the YASA library, compute the bandpowers for a selection of time points, for our frequency bands of interest, using the Welch method. 
bandpowers = yasa.bandpower(np_channels_data[:, starting_point:ending_point], 
                            sf = sampling_frequency, ch_names = channels_data.ch_names, 
                            bands = frequency_bands, win_sec = 2)
bandpowers

# %% [code]
# Plot the bandpowers. 
frequency_bands_names = ['Delta', 'Theta', 'Alpha', 'Sigma', 'Beta', 'Gamma']
plt.bar(frequency_bands_names, bandpowers.iloc[c1_channel_index, 0:6])
plt.title('Bandpowers');

# %% [markdown]
## **3. Features Extraction**

We **extract the features** by selecting a scan duration that matches the fMRI data, and computing the bandpowers for each scan. We observe that the eegNF condition is missing for subject sub-xp102, and that the fmriNF and eegfmriNF conditions are slightly longer for subject sub-xp104. 

# %% [code]
# Define the number of scans, the scan duration in seconds, and the minimal and maximal frequencies for the PSD. 
nb_scans = 200
scan_duration = 2 # In seconds. 
fmin = 1 # In Hz. 
fmax = 40 # In Hz. 

# Define the features dictionary and the summary DataFrame. 
dict_features_bandpowers_values = dict()
df_summary = pd.DataFrame()

# Compute and display the sampling window, i.e., the number of data points during each scan duration. 
sampling_window = int(scan_duration * sampling_frequency)
sampling_window

# %% [code]
%%capture
# The magic command above hides all outputs from this cell. 

# Iterate through all subjects. 
for subject in subjects:

    # Display current subject. 
    print(subject)

    # Define the subject dictionary. 
    dict_features_bandpowers_values[subject] = dict()

    # Iterate through all conditions. 
    for condition in ['eegNF', 'fmriNF', 'eegfmriNF']:

        # Skip the current iteration for sub-xp102 and eegNF, since that run is missing. 
        if subject == 'sub-xp102' and condition == 'eegNF':
            df_summary.loc[subject, condition] = 'Missing'
            continue

        # Load and extract the EEG data. 
        vhdr_path = raw_data_path + 'derivatives/' + subject + '/eeg_pp/' + subject + '_task-' + condition + '_eeg_pp.vhdr'
        channels_data = mne.io.read_raw_brainvision(vhdr_path, misc = ['ECG'], preload = True, verbose = False)
        np_channels_data = channels_data.get_data()
        df_summary.loc[subject, condition] = str(np_channels_data.shape)

        # Define the condition dictionary, and the array to store the bandpowers values. 
        dict_features_bandpowers_values[subject][condition] = np.zeros((nb_channels_psd, len(frequency_bands), nb_scans))
        
        # Iterate through all scans. 
        for scan in range(nb_scans):
            
            # Compute the bandpowers using Welch method. 
            bandpowers = yasa.bandpower(np_channels_data[:, (scan * sampling_window):(scan * sampling_window) + sampling_window], 
                                        sf = sampling_frequency, ch_names = channels_data.ch_names, 
                                        bands = frequency_bands, win_sec = scan_duration)
                    
            # Store the bandpowers values. 
            dict_features_bandpowers_values[subject][condition][:, :, scan] = bandpowers.iloc[:, 0:6].drop('ECG')

# %% [code]
# Display the summary DataFrame. 
df_summary

# %% [markdown]
## **4. Feature Engineering**

We observe that different bandpowers have significantly different mean values, a well-known effect in EEG datasets. We perform **feature engineering** by normalizing the features, and replacing the outliers (STD > 4) by the value of the previous scan, while also keeping the bandpowers without normalization as an alternative features set. 

# %% [code]
# Plot the bandpowers values by subject and condition for the C1 channel. 
fig, axes = plt.subplots(nrows = 10, ncols = 3, figsize = (12, 38))
fig.suptitle('Bandpowers', fontsize = 14, y = 0.9)
subject_index = 0

# Iterate through all subjects. 
for subject in subjects:
    condition_index = 0
    
    # Iterate through all conditions. 
    for condition in ['eegNF', 'fmriNF', 'eegfmriNF']:
        bandpower_index = 0

        # Skip the current iteration for sub-xp102 and eegNF, since that run is missing. 
        if subject == 'sub-xp102' and condition == 'eegNF':
            axes[subject_index, condition_index].set_visible(False)
            condition_index += 1
            continue

        # Iterate through all bandpowers. 
        for bandpower in ['Delta', 'Theta', 'Alpha', 'Sigma', 'Beta', 'Gamma']:
            
            # Define colormap and select a color. 
            cm = plt.colormaps['turbo']
            c = cm(0.15 * bandpower_index + 0.15)
    
            # Retrieve and plot the bandpowers values for the C1 channel. The C1 channel position is shifted because we dropped the ECG channel. 
            bandpower_values = dict_features_bandpowers_values[subject][condition][c1_channel_index_psd, bandpower_index, :]
            axes[subject_index, condition_index].plot(bandpower_values, color = c)
            axes[subject_index, condition_index].set_title(subject + ' - ' + condition)

            # Increment the bandpower index. 
            bandpower_index += 1

        # Increment the condition index. 
        condition_index += 1

    # Increment the subject index. 
    subject_index += 1

# %% [code]
# Define a dictionary to store the normalized bandpowers, and a DataFrame to store the outliers. 
dict_features_bandpowers_normalized = dict()
df_outliers = pd.DataFrame()

# Iterate through all subjects. 
for subject in subjects:

    # Define the subject dictionary. 
    dict_features_bandpowers_normalized[subject] = dict()
    
    # Iterate through all conditions. 
    for condition in ['eegNF', 'fmriNF', 'eegfmriNF']:

        # Skip the current iteration for sub-xp102 and eegNF, since that run is missing. 
        if subject == 'sub-xp102' and condition == 'eegNF':
            continue

        # Compute the normalized bandpowers. 
        bandpowers = dict_features_bandpowers_values[subject][condition]
        bandpowers_mean = np.mean(bandpowers, axis = 2)
        bandpowers_std = np.std(bandpowers, axis = 2)
        bandpowers_normalized = (bandpowers - bandpowers_mean[:, :, np.newaxis]) / bandpowers_std[:, :, np.newaxis]

        # Iterate through all bandpowers. 
        for bandpower_index in range(dict_features_bandpowers_values[subject][condition].shape[1]):
             
             # Iterate through all channels. 
            for channel_index in range(dict_features_bandpowers_values[subject][condition].shape[0]):

                # Replace the outliers (STD > 4) by the value of the previous scan. 
                outliers_counter = 0
                for scan in range(nb_scans):
                    if np.abs(bandpowers_normalized[channel_index, bandpower_index, scan]) > 4:
                        bandpowers_normalized[channel_index, bandpower_index, scan] = bandpowers_normalized[channel_index, bandpower_index, scan - 1]
                        outliers_counter += 1
                    
            # Store the number of outliers in the DataFrame. 
            df_outliers.loc[subject + '/' + condition, 'subject'] = subject
            df_outliers.loc[subject + '/' + condition, 'condition'] = condition
            df_outliers.loc[subject + '/' + condition, frequency_bands_names[bandpower_index]] = outliers_counter

        # Store the normalized bandpowers. 
        dict_features_bandpowers_normalized[subject][condition] = bandpowers_normalized

# Change the indexes and the data type of the outliers DataFrame. 
df_outliers = df_outliers.set_index(['subject', 'condition'])
df_outliers = df_outliers.astype(int)

# %% [code]
# Display the outliers DataFrame. 
df_outliers

# %% [code]
# Display the number of outliers by frequency band. 
print(df_outliers.sum())

# %% [code]
# Plot the normalized bandpowers values by subject and condition for the C1 channel. 
fig, axes = plt.subplots(nrows = 10, ncols = 3, figsize = (12, 38))
fig.suptitle('Normalized bandpowers', fontsize = 14, y = 0.9)
subject_index = 0

# Iterate through all subjects. 
for subject in subjects:
    condition_index = 0
    
    # Iterate through all conditions. 
    for condition in ['eegNF', 'fmriNF', 'eegfmriNF']:
        bandpower_index = 0

        # Skip the current iteration for sub-xp102 and eegNF, since that run is missing. 
        if subject == 'sub-xp102' and condition == 'eegNF':
            axes[subject_index, condition_index].set_visible(False)
            condition_index += 1
            continue

        # Iterate through all bandpowers. 
        for bandpower in ['Delta', 'Theta', 'Alpha', 'Sigma', 'Beta', 'Gamma']:
            
            # Define colormap and select a color. 
            cm = plt.colormaps['turbo']
            c = cm(0.15 * bandpower_index + 0.15)
    
            # Retrieve and plot the normalized bandpowers values for the C1 channel. The C1 channel position is shifted because we dropped the ECG channel. 
            bandpower_values = dict_features_bandpowers_normalized[subject][condition][c1_channel_index_psd, bandpower_index, :]
            axes[subject_index, condition_index].plot(bandpower_values, color = c)
            axes[subject_index, condition_index].set_title(subject + ' - ' + condition)

            # Increment the bandpower index. 
            bandpower_index += 1

        # Increment the condition index. 
        condition_index += 1

    # Increment the subject index. 
    subject_index += 1

# %% [markdown]
## **5. Preprocessed Features**

We save the **preprocessed features** in the appropriate directory. 

# %% [code]
# Save the features into Pickle files. 
with open(preprocessed_data_path + 'dict_features_bandpowers.p', 'wb') as file:
    pickle.dump(dict_features_bandpowers_values, file)
with open(preprocessed_data_path + 'dict_features_bandpowers_normalized.p', 'wb') as file:
    pickle.dump(dict_features_bandpowers_normalized, file)
```
