## Supplementary material for "Neural Networks and Foundation Models: Two Strategies for EEG-to-fMRI Prediction": 04_Regression_Models.ipynb

```
# %% [markdown]
# **Neural Networks and Foundation Models: Two Strategies for EEG-to-fMRI Prediction**
### **Supplementary Material**

# %% [markdown]
This project includes fourteen Notebooks:

*01: fMRI Preprocessing*

*02: EEG Preprocessing*

*03: Classification Models*

***04: Regression Models***

*05: Neural Networks*

*06: Convolutional Neural Networks*

*07: Recurrent Neural Networks*

*08: Transformers*

*09: Large Language Models*

*10: Large Language Model Chain-of-Thought*

*11: Large Language Model Fine-Tuning*

*12: Large Multimodal Model*

*13: Statistical Tests*

*14: Tables and Figures*

# %% [markdown]
### Notebook 04

# **Regression Models**

# %% [markdown]
## **1. Linear Regression, Ridge, Lasso**

We run a **linear regression model** on all subjects, except sub-xp102 who has a missing condition, and use the eegfmriNF condition as the train set, and the eegNF and fmriNF conditions as the test set. We also run a linear regression with **Ridge** and **Lasso** regularizations, and use the R^2 as our metrics. 

# %% [code]
import numpy as np
import pandas as pd
import matplotlib.pyplot as plt
%matplotlib inline
import pickle
from sklearn.base import clone
from sklearn.pipeline import Pipeline
from sklearn.preprocessing import StandardScaler
from sklearn.model_selection import GridSearchCV
from sklearn.multioutput import MultiOutputRegressor
from sklearn.linear_model import LinearRegression, Ridge, Lasso
from sklearn.neighbors import KNeighborsRegressor
from sklearn.tree import DecisionTreeRegressor
from sklearn.ensemble import RandomForestRegressor
from sklearn.svm import SVR
import xgboost as xgb

# %% [code]
# Define the preprocessed data path. 
preprocessed_data_path = '../preprocessed_data/'

# Define the results path. 
results_path = '../results/'

# %% [code]
# Load the features, the targets, and the baseline. 
with open(preprocessed_data_path + 'dict_features_bandpowers_normalized.p', 'rb') as file:
    dict_features_bandpowers = pickle.load(file)
with open(preprocessed_data_path + 'dict_targets.p', 'rb') as file:
    dict_targets = pickle.load(file)
with open(preprocessed_data_path + 'dict_baseline.p', 'rb') as file:
    dict_baseline_original = pickle.load(file)

# %% [code]
# Define the list of subjects, removing sub-xp102 who has a missing condition. 
subjects = ['sub-xp1' + str(x).zfill(2) for x in range(1, 11)]
subjects.remove('sub-xp102')
subject = subjects[0]

# Retrieve the list of all targets. 
targets = list(dict_targets[subject]['eegfmriNF'].keys())

# Select a sequence length. 
selected_sequence_length = 5

# Retrieve the total number of scans. 
nb_scans_total = dict_targets[subject]['eegfmriNF']['Frontal Pole'].shape[0]

# %% [code]
# Define the dictionaries and the summary DataFrame for storing the features, targets, and baseline. 
dict_features_basis = dict()
dict_features_sequence = dict()
dict_targets_basis = dict()
dict_targets_sequence = dict()
dict_baseline_basis = dict()
dict_baseline_sequence = dict()
df_summary_global = pd.DataFrame()

# Iterate through all subjects. 
for subject in subjects:
        
    # Define the subject dictionaries. 
    dict_features_basis[subject] = dict()
    dict_features_sequence[subject] = dict()
    dict_targets_basis[subject] = dict()
    dict_targets_sequence[subject] = dict()
    dict_baseline_basis[subject] = dict()
    dict_baseline_sequence[subject] = dict()

    # Iterate through all conditions. 
    for condition in ['eegNF', 'fmriNF', 'eegfmriNF']:

        # Skip the current iteration for sub-xp102 and eegNF, since that run is missing. 
        if subject == 'sub-xp102' and condition == 'eegNF':
            continue
            
        # Define the general features array by stacking all the EEG channels. 
        X_basis = np.vstack(dict_features_bandpowers[subject][condition]).transpose()

        # Define the general targets array. 
        Y_basis = np.zeros((nb_scans_total, len(targets)))
        for target_index in range(len(targets)):
            Y_basis[:, target_index] = dict_targets[subject][condition][targets[target_index]]

        # Define the general baseline array. 
        Baseline_basis = np.zeros((nb_scans_total, len(targets)))
        for target_index in range(len(targets)):
            Baseline_basis[:, target_index] = dict_baseline_original[subject][condition][targets[target_index]]

        # Define the particular features array. 
        X_sequence = np.zeros((nb_scans_total - selected_sequence_length, X_basis.shape[1] * (selected_sequence_length + 1)))
        for scan in range(nb_scans_total - selected_sequence_length):
            X_sequence[scan, :] = X_basis[scan:(scan + selected_sequence_length + 1), :].ravel()

        # Define the particular targets array. 
        Y_sequence = Y_basis[selected_sequence_length:, :]

        # Define the particular baseline array. 
        Baseline_sequence = Baseline_basis[selected_sequence_length:, :]

        # Store the features, targets and baseline in the dictionaries. 
        dict_features_basis[subject][condition] = X_basis
        dict_features_sequence[subject][condition] = X_sequence
        dict_targets_basis[subject][condition] = Y_basis
        dict_targets_sequence[subject][condition] = Y_sequence
        dict_baseline_basis[subject][condition] = Baseline_basis
        dict_baseline_sequence[subject][condition] = Baseline_sequence

        # Store the shapes of the arrays in the DataFrame. 
        df_summary_global.loc[subject + '/' + condition, 'subject'] = subject
        df_summary_global.loc[subject + '/' + condition, 'condition'] = condition
        df_summary_global.loc[subject + '/' + condition, 'X_basis.shape'] = str(X_basis.shape)
        df_summary_global.loc[subject + '/' + condition, 'X_sequence.shape'] = str(X_sequence.shape)
        df_summary_global.loc[subject + '/' + condition, 'Y_basis.shape'] = str(Y_basis.shape)
        df_summary_global.loc[subject + '/' + condition, 'Y_sequence.shape'] = str(Y_sequence.shape)
        df_summary_global.loc[subject + '/' + condition, 'Baseline_basis.shape'] = str(Baseline_basis.shape)
        df_summary_global.loc[subject + '/' + condition, 'Baseline_sequence.shape'] = str(Baseline_sequence.shape)

# Change the indexes of the DataFrame. 
df_summary_global = df_summary_global.set_index(['subject', 'condition'])

# %% [code]
# Display the summary DataFrame. 
df_summary_global

# %% [code]
# Define the Mean Squared Error function. 
def MSE(y, y_pred):
    return np.mean(np.square(y - y_pred))

# %% [code]
# Define the Mean Absolute Error function. 
def MAE(y, y_pred):
    return np.mean(np.abs(y - y_pred))

# %% [code]
# Define the Residual Sum of Squares function. 
def RSS(y, y_pred):
    return np.sum(np.square(y - y_pred))

# %% [code]
# Define a function to run a regression model. 
def run_regression_model(RegressionModel, dict_features_sequence, dict_targets_sequence, dict_baseline_sequence, parameter_name):

    # Define the dictionaries and the summary DataFrame. 
    dict_models = dict()
    dict_predictions = dict()
    df_summary = pd.DataFrame()

    # TRAINING LOOP. Iterate through all subjects. 
    for subject in subjects:

        # Display current subject. 
        print(subject)

        # Clone the Scikit-Learn estimator. 
        PredictorBrain = clone(RegressionModel)

        # Retrieve the features and targets of the train set. 
        X_tr = dict_features_sequence[subject]['eegfmriNF']
        Y_tr = dict_targets_sequence[subject]['eegfmriNF']

        # Train the model. 
        PredictorBrain.fit(X_tr, Y_tr)

        # Store the trained model in the dictionary. 
        dict_models[subject] = PredictorBrain

    # PREDICTION LOOP. Iterate through all subjects. 
    for subject in subjects:

        # Define the subject dictionary. 
        dict_predictions[subject] = dict()

        # Iterate through all conditions. 
        for condition in ['eegNF', 'fmriNF']:

            # Skip the current iteration for sub-xp102 and eegNF, since that run is missing. 
            if subject == 'sub-xp102' and condition == 'eegNF':
                continue

            # Retrieve the features and targets of the test set, the baseline, and the trained model. 
            X_te = dict_features_sequence[subject][condition]
            Y_te = dict_targets_sequence[subject][condition]
            Baseline = dict_baseline_sequence[subject]['eegfmriNF']
            PredictorBrain = dict_models[subject]

            # Predict the target values. 
            Y_pred = PredictorBrain.predict(X_te)

            # Evaluate the performance metrics. 
            mse_baseline = MSE(Y_te, Baseline)
            mae_baseline = MAE(Y_te, Baseline)
            rss_baseline = RSS(Y_te, Baseline)
            mse_model = MSE(Y_te, Y_pred)
            mae_model = MAE(Y_te, Y_pred)
            rss_model = RSS(Y_te, Y_pred)
            r2_model = 1 - (rss_model / rss_baseline)

            # Store the performance metrics and the shapes of the arrays in the DataFrame. 
            df_summary.loc[subject + '/' + condition, 'subject'] = subject
            df_summary.loc[subject + '/' + condition, 'condition'] = condition
            df_summary.loc[subject + '/' + condition, 'mse_baseline'] = mse_baseline
            df_summary.loc[subject + '/' + condition, 'mae_baseline'] = mae_baseline
            df_summary.loc[subject + '/' + condition, 'rss_baseline'] = rss_baseline
            df_summary.loc[subject + '/' + condition, 'mse_model'] = mse_model
            df_summary.loc[subject + '/' + condition, 'mae_model'] = mae_model
            df_summary.loc[subject + '/' + condition, 'rss_model'] = rss_model
            df_summary.loc[subject + '/' + condition, 'r2_model'] = r2_model
            df_summary.loc[subject + '/' + condition, 'X_tr.shape'] = str(X_tr.shape)
            df_summary.loc[subject + '/' + condition, 'X_te.shape'] = str(X_te.shape)
            df_summary.loc[subject + '/' + condition, 'Y_tr.shape'] = str(Y_tr.shape)
            df_summary.loc[subject + '/' + condition, 'Y_te.shape'] = str(Y_te.shape)
            if parameter_name != '':
                df_summary.loc[subject + '/' + condition, parameter_name] = PredictorBrain.best_params_[parameter_name]

            # Store the predictions in the dictionary. 
            dict_predictions[subject][condition] = Y_pred

    # Change the indexes of the DataFrame. 
    df_summary = df_summary.set_index(['subject', 'condition'])
    
    return dict_models, dict_predictions, df_summary

# %% [code]
# Run the regression model. 
dict_models_lr, dict_predictions_lr, df_summary_lr = run_regression_model(LinearRegression(), 
                                                                          dict_features_sequence, 
                                                                          dict_targets_sequence, 
                                                                          dict_baseline_sequence, 
                                                                          parameter_name = '')

# %% [code]
# Display the results. 
print('Linear Regression')
df_summary_lr

# %% [code]
# Run the regression model. 
dict_models_ridge, dict_predictions_ridge, df_summary_ridge = run_regression_model(Ridge(), 
                                                                                   dict_features_sequence, 
                                                                                   dict_targets_sequence, 
                                                                                   dict_baseline_sequence, 
                                                                                   parameter_name = '')

# %% [code]
# Display the results. 
print('Ridge Regression')
df_summary_ridge

# %% [code]
# Run the regression model. 
dict_models_lasso, dict_predictions_lasso, df_summary_lasso = run_regression_model(Lasso(), 
                                                                                   dict_features_sequence, 
                                                                                   dict_targets_sequence, 
                                                                                   dict_baseline_sequence, 
                                                                                   parameter_name = '')

# %% [code]
# Display the results. 
print('Lasso Regression')
df_summary_lasso

# %% [markdown]
## **2. K-Nearest Neighbors (KNN)**

We run a **k-nearest neighbors (KNN) model** on all subjects, except sub-xp102 who has a missing condition, and use the eegfmriNF condition as the train set, and the eegNF and fmriNF conditions as the test set. We fine-tune the number of neighbors, and use the R^2 as our metrics. 

# %% [code]
# Define the set of values to explore, where k is the number of neighbors. 
k_values = np.arange(2, 40, 2)
k_values

# %% [code]
# Create pipeline. 
pipe_knn = Pipeline([
    ('scaler', StandardScaler()),
    ('knn', KNeighborsRegressor())
])

# Create cross-validation object. 
grid_knn = {
    'knn__n_neighbors': k_values
}
grid_cv_knn = GridSearchCV(pipe_knn, grid_knn, cv = 5, verbose = 0, n_jobs = -1)

# %% [code]
# Run the regression model. 
dict_models_knn, dict_predictions_knn, df_summary_knn = run_regression_model(grid_cv_knn, 
                                                                             dict_features_sequence, 
                                                                             dict_targets_sequence, 
                                                                             dict_baseline_sequence, 
                                                                             parameter_name = 'knn__n_neighbors')

# %% [code]
# Display the results. 
print('K-Nearest Neighbors')
df_summary_knn

# %% [markdown]
## **3. Decision Tree (DT)**

We run a **decision tree (DT) model** on all subjects, except sub-xp102 who has a missing condition, and use the eegfmriNF condition as the train set, and the eegNF and fmriNF conditions as the test set. We fine-tune the depth of the tree, and use the R^2 as our metrics. 

# %% [code]
# Define the set of values to explore, where d is the depth of the tree. 
d_values = np.arange(1, 20, 1)
d_values

# %% [code]
# Create pipeline. 
pipe_dt = Pipeline([
    ('scaler', None),
    ('dt', DecisionTreeRegressor())
])

# Create cross-validation object. 
grid_dt = {
    'dt__max_depth': d_values
}
grid_cv_dt = GridSearchCV(pipe_dt, grid_dt, cv = 5, verbose = 0, n_jobs = -1)

# %% [code]
# Run the regression model. 
dict_models_dt, dict_predictions_dt, df_summary_dt = run_regression_model(grid_cv_dt, 
                                                                          dict_features_sequence, 
                                                                          dict_targets_sequence, 
                                                                          dict_baseline_sequence, 
                                                                          parameter_name = 'dt__max_depth')

# %% [code]
# Display the results. 
print('Decision Tree')
df_summary_dt

# %% [markdown]
## **4. Random Forest (RF)**

We run a **random forest (RF) model** on all subjects, except sub-xp102 who has a missing condition, and use the eegfmriNF condition as the train set, and the eegNF and fmriNF conditions as the test set. We use the R^2 as our metrics. 

# %% [code]
# Create pipeline. 
pipe_rf = Pipeline([
    ('scaler', None),
    ('rf', RandomForestRegressor())
])

# %% [code]
# Run the regression model. 
dict_models_rf, dict_predictions_rf, df_summary_rf = run_regression_model(pipe_rf, 
                                                                          dict_features_sequence, 
                                                                          dict_targets_sequence, 
                                                                          dict_baseline_sequence, 
                                                                          parameter_name = '')

# %% [code]
# Display the results. 
print('Random Forests')
df_summary_rf

# %% [markdown]
## **5. Support Vector Machine (SVM)**

We run a **support vector machine (SVM) model** on all subjects, except sub-xp102 who has a missing condition, and use the eegfmriNF condition as the train set, and the eegNF and fmriNF conditions as the test set. We use the R^2 as our metrics. 

# %% [code]
# Create pipeline. 
pipe_svm = Pipeline([
    ('scaler', StandardScaler()),
    ('svm', MultiOutputRegressor(SVR()))
])

# %% [code]
# Run the regression model. 
dict_models_svm, dict_predictions_svm, df_summary_svm = run_regression_model(pipe_svm, 
                                                                             dict_features_sequence, 
                                                                             dict_targets_sequence, 
                                                                             dict_baseline_sequence, 
                                                                             parameter_name = '')

# %% [code]
# Display the results. 
print('Support Vector Machines')
df_summary_svm

# %% [markdown]
## **6. Extreme Gradient Boosting (XGBoost)**

We run an **extreme gradient boosting (XGBoost) model** on all subjects, except sub-xp102 who has a missing condition, and use the eegfmriNF condition as the train set, and the eegNF and fmriNF conditions as the test set. We use the R^2 as our metrics. 

# %% [code]
# Create pipeline. 
pipe_xgb = Pipeline([
    ('scaler', StandardScaler()),
    ('xgb', xgb.XGBRegressor())
])

# %% [code]
# Run the regression model. 
dict_models_xgb, dict_predictions_xgb, df_summary_xgb = run_regression_model(pipe_xgb, 
                                                                             dict_features_sequence, 
                                                                             dict_targets_sequence, 
                                                                             dict_baseline_sequence, 
                                                                             parameter_name = '')

# %% [code]
# Display the results. 
print('XGBoost')
df_summary_xgb

# %% [markdown]
## **7. Results**

# %% [code]
# Retrieve the average MAE and the standard error of the MAE. 
mae = [df_summary_lr['mae_baseline'].mean(), 
       df_summary_lr['mae_model'].mean(), 
       df_summary_ridge['mae_model'].mean(), 
       df_summary_lasso['mae_model'].mean(), 
       df_summary_knn['mae_model'].mean(), 
       df_summary_dt['mae_model'].mean(), 
       df_summary_rf['mae_model'].mean(), 
       df_summary_svm['mae_model'].mean(),
       df_summary_xgb['mae_model'].mean()]
mae_se = [np.std(df_summary_lr['mae_baseline']) / np.sqrt(df_summary_lr['mae_baseline'].shape[0]), 
          np.std(df_summary_lr['mae_model']) / np.sqrt(df_summary_lr['mae_model'].shape[0]), 
          np.std(df_summary_ridge['mae_model']) / np.sqrt(df_summary_ridge['mae_model'].shape[0]), 
          np.std(df_summary_lasso['mae_model']) / np.sqrt(df_summary_lasso['mae_model'].shape[0]), 
          np.std(df_summary_knn['mae_model']) / np.sqrt(df_summary_knn['mae_model'].shape[0]), 
          np.std(df_summary_dt['mae_model']) / np.sqrt(df_summary_dt['mae_model'].shape[0]), 
          np.std(df_summary_rf['mae_model']) / np.sqrt(df_summary_rf['mae_model'].shape[0]), 
          np.std(df_summary_svm['mae_model']) / np.sqrt(df_summary_svm['mae_model'].shape[0]),
          np.std(df_summary_xgb['mae_model']) / np.sqrt(df_summary_xgb['mae_model'].shape[0])]

# Display the average MAE and its standard error in a bar plot. 
fig, axes = plt.subplots(nrows = 1, ncols = 1, figsize = (10, 4))
axes.bar(range(9), mae, width = 0.6, color = ['green', 'blue', 'blue', 'blue', 'blue', 'blue', 'blue', 'blue', 'blue', 'blue'])
axes.errorbar(range(9), mae, yerr = mae_se, fmt = 'None', color = 'black')
axes.axhline(mae[0], 0, 1, color = 'green', linestyle = 'dashed', alpha = 0.5)
axes.set_xticks(range(9), labels = ['Baseline', 'LR', 'Ridge', 'Lasso', 'KNN', 'DT', 'RF', 'SVM', 'XGB'])
axes.set_ylabel('Average MAE')
axes.set_title('Regression models');

# %% [code]
# Save the features into Pickle files. 
with open(preprocessed_data_path + 'regression/dict_features_regression_basis.p', 'wb') as file:
    pickle.dump(dict_features_basis, file)
with open(preprocessed_data_path + 'regression/dict_features_regression_sequence.p', 'wb') as file:
    pickle.dump(dict_features_sequence, file)
    
# Save the targets into Pickle files. 
with open(preprocessed_data_path + 'regression/dict_targets_regression_basis.p', 'wb') as file:
    pickle.dump(dict_targets_basis, file)
with open(preprocessed_data_path + 'regression/dict_targets_regression_sequence.p', 'wb') as file:
    pickle.dump(dict_targets_sequence, file)

# Save the baselines into Pickle files. 
with open(preprocessed_data_path + 'regression/dict_baseline_regression_basis.p', 'wb') as file:
    pickle.dump(dict_baseline_basis, file)
with open(preprocessed_data_path + 'regression/dict_baseline_regression_sequence.p', 'wb') as file:
    pickle.dump(dict_baseline_sequence, file)

# Save the models into Pickle files. 
with open(results_path + 'regression/dict_models_lr.p', 'wb') as file:
    pickle.dump(dict_models_lr, file)
with open(results_path + 'regression/dict_models_ridge.p', 'wb') as file:
    pickle.dump(dict_models_ridge, file)
with open(results_path + 'regression/dict_models_lasso.p', 'wb') as file:
    pickle.dump(dict_models_lasso, file)
with open(results_path + 'regression/dict_models_knn.p', 'wb') as file:
    pickle.dump(dict_models_knn, file)
with open(results_path + 'regression/dict_models_dt.p', 'wb') as file:
    pickle.dump(dict_models_dt, file)
with open(results_path + 'regression/dict_models_rf.p', 'wb') as file:
    pickle.dump(dict_models_rf, file)
with open(results_path + 'regression/dict_models_svm.p', 'wb') as file:
    pickle.dump(dict_models_svm, file)
with open(results_path + 'regression/dict_models_xgb.p', 'wb') as file:
    pickle.dump(dict_models_xgb, file)

# Save the predictions into Pickle files. 
with open(results_path + 'regression/dict_predictions_lr.p', 'wb') as file:
    pickle.dump(dict_predictions_lr, file)
with open(results_path + 'regression/dict_predictions_ridge.p', 'wb') as file:
    pickle.dump(dict_predictions_ridge, file)
with open(results_path + 'regression/dict_predictions_lasso.p', 'wb') as file:
    pickle.dump(dict_predictions_lasso, file)
with open(results_path + 'regression/dict_predictions_knn.p', 'wb') as file:
    pickle.dump(dict_predictions_knn, file)
with open(results_path + 'regression/dict_predictions_dt.p', 'wb') as file:
    pickle.dump(dict_predictions_dt, file)
with open(results_path + 'regression/dict_predictions_rf.p', 'wb') as file:
    pickle.dump(dict_predictions_rf, file)
with open(results_path + 'regression/dict_predictions_svm.p', 'wb') as file:
    pickle.dump(dict_predictions_svm, file)
with open(results_path + 'regression/dict_predictions_xgb.p', 'wb') as file:
    pickle.dump(dict_predictions_xgb, file)
```
