## Supplementary material for "Neural Networks and Foundation Models: Two Strategies for EEG-to-fMRI Prediction": 06_Convolutional_Neural_Networks.ipynb

```
# %% [markdown]
# **Neural Networks and Foundation Models: Two Strategies for EEG-to-fMRI Prediction**
### **Supplementary Material**

# %% [markdown]
This project includes fourteen Notebooks:

*01: fMRI Preprocessing*

*02: EEG Preprocessing*

*03: Classification Models*

*04: Regression Models*

*05: Neural Networks*

***06: Convolutional Neural Networks***

*07: Recurrent Neural Networks*

*08: Transformers*

*09: Large Language Models*

*10: Large Language Model Chain-of-Thought*

*11: Large Language Model Fine-Tuning*

*12: Large Multimodal Model*

*13: Statistical Tests*

*14: Tables and Figures*

# %% [markdown]
### Notebook 06

# **Convolutional Neural Networks**

# %% [markdown]
## **1. Classification**

We define a **convolutional neural network model** for a classification task, using the **Adam optimizer** and the **binary cross-entropy loss function**. Our objective is to predict whether the fMRI signal increases or decreases in our brain regions of interest, based directly on the raw EEG signal instead of the EEG bandpowers. We run the model on all subjects, except sub-xp102 who has a missing condition, and use the eegfmriNF condition as the train set, the eegNF condition as the validation set, and the fmriNF condition as the test set. 

# %% [code]
import numpy as np
import pandas as pd
import matplotlib.pyplot as plt
%matplotlib inline
import pickle
import mne
from sklearn.preprocessing import StandardScaler
import tensorflow as tf
import os
import logging

# %% [code]
# Define the raw data path. 
raw_data_path = '../ds002336-download/'

# Define the preprocessed data path. 
preprocessed_data_path = '../preprocessed_data/'

# Define the results path. 
results_path = '../results/'

# %% [code]
# Load the targets and the baseline for the classification task. 
with open(preprocessed_data_path + 'dict_targets.p', 'rb') as file:
    dict_targets = pickle.load(file)
with open(preprocessed_data_path + 'classification/dict_targets_classification_sequence.p', 'rb') as file:
    dict_targets_sequence_class = pickle.load(file)
with open(preprocessed_data_path + 'classification/dict_baseline_classification_sequence.p', 'rb') as file:
    dict_baseline_sequence_class = pickle.load(file)

# %% [code]
# Define the list of subjects, removing sub-xp102 who has a missing condition. 
subjects = ['sub-xp1' + str(x).zfill(2) for x in range(1, 11)]
subjects.remove('sub-xp102')
subject = subjects[0]

# Retrieve the list of all targets. 
targets = list(dict_targets[subject]['eegfmriNF'].keys())

# Select a sequence length. 
selected_sequence_length = 5

# Retrieve the total number of scans. 
nb_scans_total = dict_targets[subject]['eegfmriNF']['Frontal Pole'].shape[0]

# Define the batch size for the neural networks models. 
batch_size = 32

# Define a dictionary to store the FLOPS for each model. 
dict_flops = dict()

# %% [code]
# Define the path of the header file (.vhdr) for an example subject, for the eegfmriNF condition. 
vhdr_path = raw_data_path + 'derivatives/' + subject + '/eeg_pp/' + subject + '_task-eegfmriNF_eeg_pp.vhdr'

# Load the EEG data using the MNE library. 
channels_data = mne.io.read_raw_brainvision(vhdr_path, misc = ['ECG'], preload = True, verbose = False)

# %% [code]
# Retrieve and display the number of channels. 
nb_channels = len(channels_data.ch_names)
nb_channels

# %% [code]
# Retrieve and display the sampling frequency. 
sampling_frequency = channels_data.info['sfreq']
sampling_frequency

# %% [code]
# Define the number of scans, and the scan duration in seconds. 
nb_scans = 200
scan_duration = 2

# Compute and display the sampling window, i.e., the number of data points during each scan duration. 
sampling_window = int(scan_duration * sampling_frequency)
sampling_window

# %% [code]
# Compute and display the sequence window, i.e., the number of data points during each sequence duration (including the current scan). 
sequence_window = (sampling_window * selected_sequence_length) + sampling_window
sequence_window

# %% [code]
# Define the dictionaries and the summary DataFrame for storing the features. 
dict_features_basis = dict()
dict_features_sequence_class = dict()
dict_features_sequence_reg = dict()
df_summary_global = pd.DataFrame()

# %% [code]
%%capture
# The magic command above hides all outputs from this cell. 

# Iterate through all subjects. 
for subject in subjects:
        
    # Define the subject dictionaries. 
    dict_features_basis[subject] = dict()
    dict_features_sequence_class[subject] = dict()
    dict_features_sequence_reg[subject] = dict()

    # Iterate through all conditions. 
    for condition in ['eegNF', 'fmriNF', 'eegfmriNF']:

        # Skip the current iteration for sub-xp102 and eegNF, since that run is missing. 
        if subject == 'sub-xp102' and condition == 'eegNF':
            continue

        # Load and extract the EEG data in the general features array (2D). Dimensions: data points, channels. 
        vhdr_path = raw_data_path + 'derivatives/' + subject + '/eeg_pp/' + subject + '_task-' + condition + '_eeg_pp.vhdr'
        channels_data = mne.io.read_raw_brainvision(vhdr_path, misc = ['ECG'], preload = True, verbose = False)
        np_channels_data = channels_data.get_data()
        X_basis = np_channels_data.reshape((np_channels_data.shape[1], np_channels_data.shape[0]))

        # Define the particular features array (3D). Dimensions: scans, data points, channels. 
        X_sequence = np.zeros((nb_scans_total - selected_sequence_length, sequence_window, X_basis.shape[1]))
        for scan in range(nb_scans_total - selected_sequence_length):
            X_sequence[scan, :, :] = X_basis[(scan * sampling_window):((scan * sampling_window) + sequence_window), :]

        # Store the features in the dictionaries. 
        dict_features_basis[subject][condition] = X_basis
        dict_features_sequence_class[subject][condition] = X_sequence[1:, :, :]
        dict_features_sequence_reg[subject][condition] = X_sequence

        # Store the shapes of the arrays in the DataFrame. 
        df_summary_global.loc[subject + '/' + condition, 'subject'] = subject
        df_summary_global.loc[subject + '/' + condition, 'condition'] = condition
        df_summary_global.loc[subject + '/' + condition, 'X_basis.shape'] = str(X_basis.shape)
        df_summary_global.loc[subject + '/' + condition, 'dict_features_sequence_class.shape'] = str(X_sequence[1:, :, :].shape)
        df_summary_global.loc[subject + '/' + condition, 'dict_features_sequence_reg.shape'] = str(X_sequence.shape)

# Change the indexes of the DataFrame. 
df_summary_global = df_summary_global.set_index(['subject', 'condition'])

# %% [code]
# Display the summary DataFrame. 
df_summary_global

# %% [code]
# Define a function to run a convolutional neural network model for a classification task. 
def run_cnn_classification(NeuralNetworkModel, dict_features_sequence, dict_targets_sequence, dict_baseline_sequence):

    # Define the dictionaries and the summary DataFrame. 
    dict_models = dict()
    dict_history = dict()
    dict_predictions = dict()
    df_summary = pd.DataFrame()

    # Iterate through all subjects. 
    for subject in subjects:

        # Display current subject. 
        print(subject)

        # Clone the Neural Network model. 
        PredictorBrain = tf.keras.models.clone_model(NeuralNetworkModel)

        # Retrieve the features of the train set, validation set, and test set. 
        X_tr = dict_features_sequence[subject]['eegfmriNF']
        X_va = dict_features_sequence[subject]['eegNF']
        X_te = dict_features_sequence[subject]['fmriNF']
        
        # Retrieve the targets of the train set, validation set, and test set. 
        Y_tr = dict_targets_sequence[subject]['eegfmriNF']
        Y_va = dict_targets_sequence[subject]['eegNF']
        Y_te = dict_targets_sequence[subject]['fmriNF']

        # Retrieve the baseline of the train set and determine the baseline accuracy. 
        Baseline = dict_baseline_sequence[subject]['eegfmriNF']
        accuracy_baseline = np.mean(Baseline == Y_te)

        # Rescale features variables. 
        X_tr_standardized = np.zeros(X_tr.shape)
        X_va_standardized = np.zeros(X_va.shape)
        X_te_standardized = np.zeros(X_te.shape)
        for channel_index in range(nb_channels):
            scaler = StandardScaler()
            X_tr_standardized[:, channel_index, :] = scaler.fit_transform(X_tr[:, channel_index, :])
            X_va_standardized[:, channel_index, :] = scaler.transform(X_va[:, channel_index, :])
            X_te_standardized[:, channel_index, :] = scaler.transform(X_te[:, channel_index, :])

        # End training when accuracy stops improving. 
        early_stopping = tf.keras.callbacks.EarlyStopping(monitor = 'val_loss', patience = 6, 
                                                          restore_best_weights = True)

        # Define the loss function, optimizer, and metrics to track during training. 
        PredictorBrain.compile(optimizer = tf.keras.optimizers.Adam(learning_rate = 0.001), 
                               loss = 'binary_crossentropy', metrics = ['binary_accuracy'])

        # Fit the model. 
        NeuralNetworkHistory = PredictorBrain.fit(
            x = X_tr_standardized, y = Y_tr,
            validation_data = (X_va_standardized, Y_va), 
            batch_size = batch_size, epochs = 30,
            shuffle = True, callbacks = [early_stopping], verbose = 0
        )

        # Evaluate the test accuracy. 
        (loss_test, accuracy_test) = PredictorBrain.evaluate(X_te_standardized, Y_te, batch_size = batch_size)
        print(f'test loss: {loss_test:.4f}, test accuracy: {accuracy_test:.4f}')

        # Compute the predictions from the neural network. 
        Y_pred = PredictorBrain.predict(X_te_standardized)

        # Convert the probabilities to binary predictions. 
        Y_pred_binary = (Y_pred > 0.5).astype(int)

        # Store the performance metrics and the shapes of the arrays in the DataFrame. 
        df_summary.loc[subject, 'loss_train'] = NeuralNetworkHistory.history['loss'][-1]
        df_summary.loc[subject, 'loss_valid'] = NeuralNetworkHistory.history['val_loss'][-1]
        df_summary.loc[subject, 'loss_test'] = loss_test
        df_summary.loc[subject, 'accuracy_train'] = NeuralNetworkHistory.history['binary_accuracy'][-1]
        df_summary.loc[subject, 'accuracy_valid'] = NeuralNetworkHistory.history['val_binary_accuracy'][-1]
        df_summary.loc[subject, 'accuracy_test'] = accuracy_test
        df_summary.loc[subject, 'accuracy_baseline'] = accuracy_baseline
        df_summary.loc[subject, 'above_baseline'] = str(accuracy_test > accuracy_baseline)
        df_summary.loc[subject, 'X_tr.shape'] = str(X_tr_standardized.shape)
        df_summary.loc[subject, 'X_va.shape'] = str(X_va_standardized.shape)
        df_summary.loc[subject, 'X_te.shape'] = str(X_te_standardized.shape)
        df_summary.loc[subject, 'Y_tr.shape'] = str(Y_tr.shape)
        df_summary.loc[subject, 'Y_va.shape'] = str(Y_va.shape)
        df_summary.loc[subject, 'Y_te.shape'] = str(Y_te.shape)

        # Store the trained model, the model history, and the predictions in the dictionary. 
        dict_models[subject] = PredictorBrain
        dict_history[subject] = NeuralNetworkHistory
        dict_predictions[subject] = Y_pred_binary
    
    return dict_models, dict_history, dict_predictions, df_summary

# %% [code]
# Define a function to plot the results of the convolutional neural network training. 
def plot_cnn_classification(dict_history_class, title):
    
    # Display the graph. 
    fig, axes = plt.subplots(nrows = 9, ncols = 2, figsize = (12, 52))
    fig.suptitle(title, fontsize = 14, y = 0.895)
    subject_index = 0

    # Iterate through all subjects. 
    for subject in subjects:

        # Retrieve the history. 
        NeuralNetworkHistory = dict_history_class[subject]

        # Plot loss values. 
        axes[subject_index, 0].plot(NeuralNetworkHistory.history['loss'], label = 'train loss', color = 'blue')
        axes[subject_index, 0].plot(NeuralNetworkHistory.history['val_loss'], label = 'val loss', color = 'green')
        axes[subject_index, 0].set_title(str(subject) + ' valid. loss: {:.3f} (mean last 3)'.format(
            np.mean(NeuralNetworkHistory.history['val_loss'][-3:]) # Take the mean of the last three values. 
        ))
        axes[subject_index, 0].set_xlabel('Epoch')
        axes[subject_index, 0].set_ylabel('Loss value')
        axes[subject_index, 0].legend()

        # Plot accuracy values. 
        axes[subject_index, 1].plot(NeuralNetworkHistory.history['binary_accuracy'], label = 'train acc', color = 'blue')
        axes[subject_index, 1].plot(NeuralNetworkHistory.history['val_binary_accuracy'], label = 'val acc', color = 'green')
        axes[subject_index, 1].set_title(str(subject) + ' valid. accuracy: {:.3f} (mean last 3)'.format(
            np.mean(NeuralNetworkHistory.history['val_binary_accuracy'][-3:]) # Take the mean of the last three values. 
        ))
        axes[subject_index, 1].set_xlabel('Epoch')
        axes[subject_index, 1].set_ylabel('Accuracy')
        axes[subject_index, 1].legend();

        # Increment. 
        subject_index += 1

# %% [code]
# Define the neural network model. 
tf.random.set_seed(0)
NeuralNetworkModel = tf.keras.Sequential()
input_dim = dict_features_sequence_class[subject]['eegfmriNF'].shape[1:]

# Input layer. 
NeuralNetworkModel.add(tf.keras.layers.InputLayer(shape = input_dim))

# 1D convolutional layer. 
NeuralNetworkModel.add(tf.keras.layers.Conv1D(
    filters = 16, 
    kernel_size = 8,
    activation = tf.keras.activations.relu, 
    data_format = 'channels_last'
    ))
NeuralNetworkModel.add(tf.keras.layers.MaxPool1D(data_format = 'channels_last'))
NeuralNetworkModel.add(tf.keras.layers.Dropout(0.5))

# 1D convolutional layer. 
NeuralNetworkModel.add(tf.keras.layers.Conv1D(
    filters = 32, 
    kernel_size = 8, 
    activation = tf.keras.activations.relu, 
    data_format = 'channels_last'
    ))
NeuralNetworkModel.add(tf.keras.layers.MaxPool1D(data_format = 'channels_last'))
NeuralNetworkModel.add(tf.keras.layers.Flatten(data_format = 'channels_last'))
NeuralNetworkModel.add(tf.keras.layers.Dropout(0.5))

# Output layer. 
NeuralNetworkModel.add(tf.keras.layers.Dense(
    units = len(targets), activation = tf.keras.activations.sigmoid,
    kernel_initializer = tf.keras.initializers.VarianceScaling(scale = 1.0, seed = 0)))

# Print network summary. 
NeuralNetworkModel.summary()

# %% [code]
# Run the convolutional neural network model. 
dict_models_class, dict_history_class, dict_predictions_class, df_summary_class = run_cnn_classification(NeuralNetworkModel, 
                                                                                                         dict_features_sequence_class, 
                                                                                                         dict_targets_sequence_class, 
                                                                                                         dict_baseline_sequence_class)

# %% [code]
# Display the results. 
print('Convolutional Neural Networks: Classification')
df_summary_class

# %% [code]
# Display the results. 
plot_cnn_classification(dict_history_class, title = 'Convolutional Neural Networks: Classification')

# %% [code]
# Define a function to measure the FLOPs of a model. 
def get_flops(model, input_shape):

    # Retrieve the graph of the concrete function. 
    concrete_func = tf.function(model).get_concrete_function(tf.TensorSpec(input_shape, tf.float32))
    graph = concrete_func.graph

    # Measure the FLOPs. 
    run_meta = tf.compat.v1.RunMetadata()
    opts = tf.compat.v1.profiler.ProfileOptionBuilder.float_operation()
    flops = tf.compat.v1.profiler.profile(graph = graph,
                                          run_meta = run_meta,
                                          cmd = 'op',
                                          options = opts)

    return flops.total_float_ops

# %% [code]
# Suppress TensorFlow warnings. 
tf.get_logger().setLevel(logging.ERROR)

# Measure the FLOPs of the model. 
input_shape = (1, *input_dim)  # Batch size = 1. Input dimension is entered with argument unpacking. 
flops = get_flops(NeuralNetworkModel, input_shape)

# %% [code]
# Store and display the number of FLOPs. 
dict_flops['Convolutional Neural Networks: Classification'] = flops
print('FLOPs per inference:', flops)

# %% [code]
# Retrieve the average accuracy and the standard error of the accuracy. 
acc = [df_summary_class['accuracy_baseline'].mean(), 
       df_summary_class['accuracy_train'].mean(), 
       df_summary_class['accuracy_valid'].mean(), 
       df_summary_class['accuracy_test'].mean()]
acc_se = [np.std(df_summary_class['accuracy_baseline']) / np.sqrt(df_summary_class['accuracy_baseline'].shape[0]), 
          np.std(df_summary_class['accuracy_train']) / np.sqrt(df_summary_class['accuracy_train'].shape[0]), 
          np.std(df_summary_class['accuracy_valid']) / np.sqrt(df_summary_class['accuracy_valid'].shape[0]), 
          np.std(df_summary_class['accuracy_test']) / np.sqrt(df_summary_class['accuracy_test'].shape[0])]

# Display the average accuracy and its standard error in a bar plot. 
fig, axes = plt.subplots(nrows = 1, ncols = 1, figsize = (6, 4))
axes.bar(range(4), acc, width = 0.6, color = ['green', 'blue', 'blue', 'blue'])
axes.errorbar(range(4), acc, yerr = acc_se, fmt = 'None', color = 'black')
axes.axhline(acc[0], 0, 1, color = 'green', linestyle = 'dashed', alpha = 0.5)
axes.set_xticks(range(4), labels = ['Baseline', 'Train', 'Valid', 'Test'])
axes.set_ylabel('Average accuracy')
axes.set_title('Convolutional Neural Networks: Classification');

# %% [markdown]
## **2. Regression**

We define a **convolutional neural network model** for a regression task, using the **Adam optimizer** and the **mean squared error loss function**. Our objective is to predict the value of the fMRI signal in our brain regions of interest, based directly on the raw EEG signal instead of the EEG bandpowers. We run the model on all subjects, except sub-xp102 who has a missing condition, and use the eegfmriNF condition as the train set, the eegNF condition as the validation set, and the fmriNF condition as the test set. 

# %% [code]
# Load the targets and the baseline for the regression task. 
with open(preprocessed_data_path + 'regression/dict_targets_regression_sequence.p', 'rb') as file:
    dict_targets_sequence_reg = pickle.load(file)
with open(preprocessed_data_path + 'regression/dict_baseline_regression_sequence.p', 'rb') as file:
    dict_baseline_sequence_reg = pickle.load(file)

# %% [code]
# Define the Mean Squared Error function. 
def MSE(y, y_pred):
    return np.mean(np.square(y - y_pred))

# %% [code]
# Define the Mean Absolute Error function. 
def MAE(y, y_pred):
    return np.mean(np.abs(y - y_pred))

# %% [code]
# Define the Residual Sum of Squares function. 
def RSS(y, y_pred):
    return np.sum(np.square(y - y_pred))

# %% [code]
# Define a function to run a convolutional neural network model for a regression task. 
def run_cnn_regression(NeuralNetworkModel, dict_features_sequence, dict_targets_sequence, dict_baseline_sequence):

    # Define the dictionaries and the summary DataFrame. 
    dict_models = dict()
    dict_history = dict()
    dict_predictions = dict()
    df_summary = pd.DataFrame()

    # Iterate through all subjects. 
    for subject in subjects:

        # Display current subject. 
        print(subject)

        # Clone the Neural Network model. 
        PredictorBrain = tf.keras.models.clone_model(NeuralNetworkModel)

        # Retrieve the features of the train set, validation set, and test set. 
        X_tr = dict_features_sequence[subject]['eegfmriNF']
        X_va = dict_features_sequence[subject]['eegNF']
        X_te = dict_features_sequence[subject]['fmriNF']

        # Retrieve the targets of the train set, validation set, and test set. 
        Y_tr = dict_targets_sequence[subject]['eegfmriNF']
        Y_va = dict_targets_sequence[subject]['eegNF']
        Y_te = dict_targets_sequence[subject]['fmriNF']

        # Retrieve the baseline of the train set. 
        Baseline = dict_baseline_sequence[subject]['eegfmriNF']

        # Rescale features variables. 
        X_tr_standardized = np.zeros(X_tr.shape)
        X_va_standardized = np.zeros(X_va.shape)
        X_te_standardized = np.zeros(X_te.shape)
        for channel_index in range(nb_channels):
            scaler = StandardScaler()
            X_tr_standardized[:, channel_index, :] = scaler.fit_transform(X_tr[:, channel_index, :])
            X_va_standardized[:, channel_index, :] = scaler.transform(X_va[:, channel_index, :])
            X_te_standardized[:, channel_index, :] = scaler.transform(X_te[:, channel_index, :])

        # End training when accuracy stops improving. 
        early_stopping = tf.keras.callbacks.EarlyStopping(monitor = 'val_loss', patience = 6, 
                                                          restore_best_weights = True)

        # Define the loss function, optimizer, and metrics to track during training. 
        optimizer = tf.keras.optimizers.Adam(learning_rate = 0.005, beta_1 = 0.9)
        PredictorBrain.compile(optimizer = optimizer, loss = 'mse', metrics = [tf.keras.metrics.MeanAbsoluteError()])

        # Fit the model. 
        NeuralNetworkHistory = PredictorBrain.fit(
            x = X_tr_standardized, y = Y_tr,
            validation_data = (X_va_standardized, Y_va), 
            batch_size = batch_size, epochs = 30, 
            shuffle = True, callbacks = [early_stopping], verbose = 0
        )

        # Evaluate the test MSE and MAE. 
        (mse_test, mae_test) = PredictorBrain.evaluate(X_te_standardized, Y_te, batch_size = batch_size)
        print(f'test MSE: {mse_test:.4f}, test MAE: {mae_test:.4f}')

        # Compute the predictions from the neural network. 
        Y_pred = PredictorBrain.predict(X_te_standardized)

        # Evaluate the performance metrics. 
        mse_baseline = MSE(Y_te, Baseline)
        mae_baseline = MAE(Y_te, Baseline)
        rss_baseline = RSS(Y_te, Baseline)
        mse_model = MSE(Y_te, Y_pred)
        mae_model = MAE(Y_te, Y_pred)
        rss_model = RSS(Y_te, Y_pred)
        r2_model = 1 - (rss_model / rss_baseline)

        # Store the performance metrics and the shapes of the arrays in the DataFrame. 
        df_summary.loc[subject, 'mse_train'] = NeuralNetworkHistory.history['loss'][-1]
        df_summary.loc[subject, 'mse_valid'] = NeuralNetworkHistory.history['val_loss'][-1]
        df_summary.loc[subject, 'mse_test'] = mse_test
        df_summary.loc[subject, 'mae_train'] = NeuralNetworkHistory.history['mean_absolute_error'][-1]
        df_summary.loc[subject, 'mae_valid'] = NeuralNetworkHistory.history['val_mean_absolute_error'][-1]
        df_summary.loc[subject, 'mae_test'] = mae_test
        df_summary.loc[subject, 'mae_baseline'] = mae_baseline
        df_summary.loc[subject, 'r2_model'] = r2_model
        df_summary.loc[subject, 'X_tr.shape'] = str(X_tr_standardized.shape)
        df_summary.loc[subject, 'X_va.shape'] = str(X_va_standardized.shape)
        df_summary.loc[subject, 'X_te.shape'] = str(X_te_standardized.shape)
        df_summary.loc[subject, 'Y_tr.shape'] = str(Y_tr.shape)
        df_summary.loc[subject, 'Y_va.shape'] = str(Y_va.shape)
        df_summary.loc[subject, 'Y_te.shape'] = str(Y_te.shape)

        # Store the trained model, the model history, and the predictions in the dictionary. 
        dict_models[subject] = PredictorBrain
        dict_history[subject] = NeuralNetworkHistory
        dict_predictions[subject] = Y_pred
    
    return dict_models, dict_history, dict_predictions, df_summary

# %% [code]
# Define a function to plot the results of the neural network training. 
def plot_cnn_regression(dict_history_reg, title):
    
    # Display the graph. 
    fig, axes = plt.subplots(nrows = 9, ncols = 2, figsize = (12, 52))
    fig.suptitle(title, fontsize = 14, y = 0.895)
    subject_index = 0

    # Iterate through all subjects. 
    for subject in subjects:

        # Retrieve the history. 
        NeuralNetworkHistory = dict_history_reg[subject]

        # Plot MSE values. 
        axes[subject_index, 0].plot(NeuralNetworkHistory.history['loss'], label = 'train MSE', color = 'blue')
        axes[subject_index, 0].plot(NeuralNetworkHistory.history['val_loss'], label = 'val MSE', color = 'green')
        axes[subject_index, 0].set_title(str(subject) + ' valid. MSE: {:.3f} (mean last 3)'.format(
            np.mean(NeuralNetworkHistory.history['val_loss'][-3:]) # Take the mean of the last three values. 
        ))
        axes[subject_index, 0].set_xlabel('Epoch')
        axes[subject_index, 0].set_ylabel('MSE')
        axes[subject_index, 0].legend()

        # Plot MAE values. 
        axes[subject_index, 1].plot(NeuralNetworkHistory.history['mean_absolute_error'], label = 'train MAE', color = 'blue')
        axes[subject_index, 1].plot(NeuralNetworkHistory.history['val_mean_absolute_error'], label = 'val MAE', color = 'green')
        axes[subject_index, 1].set_title(str(subject) + ' valid. MAE: {:.3f} (mean last 3)'.format(
            np.mean(NeuralNetworkHistory.history['val_mean_absolute_error'][-3:]) # Take the mean of the last three values. 
        ))
        axes[subject_index, 1].set_xlabel('Epoch')
        axes[subject_index, 1].set_ylabel('MAE')
        axes[subject_index, 1].legend();

        # Increment. 
        subject_index += 1

# %% [code]
# Define the neural network model. 
tf.random.set_seed(0)
NeuralNetworkModel = tf.keras.Sequential()
input_dim = dict_features_sequence_reg[subject]['eegfmriNF'].shape[1:]

# Input layer. 
NeuralNetworkModel.add(tf.keras.layers.InputLayer(shape = input_dim))

# 1D convolutional layer. 
NeuralNetworkModel.add(tf.keras.layers.Conv1D(
    filters = 16, 
    kernel_size = 8,
    activation = tf.keras.activations.relu, 
    data_format = 'channels_last'
    ))
NeuralNetworkModel.add(tf.keras.layers.MaxPool1D(data_format = 'channels_last'))
NeuralNetworkModel.add(tf.keras.layers.Dropout(0.5))

# 1D convolutional layer. 
NeuralNetworkModel.add(tf.keras.layers.Conv1D(
    filters = 32, 
    kernel_size = 8, 
    activation = tf.keras.activations.relu, 
    data_format = 'channels_last'
    ))
NeuralNetworkModel.add(tf.keras.layers.MaxPool1D(data_format = 'channels_last'))
NeuralNetworkModel.add(tf.keras.layers.Flatten(data_format = 'channels_last'))
NeuralNetworkModel.add(tf.keras.layers.Dropout(0.5))

# Output layer. 
NeuralNetworkModel.add(tf.keras.layers.Dense(
    units = dict_targets_sequence_reg[subject]['eegfmriNF'].shape[1], 
    ))

# Print network summary. 
NeuralNetworkModel.summary()

# %% [code]
# Run the convolutional neural network model. 
dict_models_reg, dict_history_reg, dict_predictions_reg, df_summary_reg = run_cnn_regression(NeuralNetworkModel, 
                                                                                             dict_features_sequence_reg, 
                                                                                             dict_targets_sequence_reg, 
                                                                                             dict_baseline_sequence_reg)

# %% [code]
# Display the results. 
print('Convolutional Neural Networks: Regression')
df_summary_reg

# %% [code]
# Display the results. 
plot_cnn_regression(dict_history_reg, title = 'Convolutional Neural Networks: Regression')

# %% [code]
# Measure the FLOPs of the model. 
input_shape = (1, *input_dim)  # Batch size = 1. Input dimension is entered with argument unpacking. 
flops = get_flops(NeuralNetworkModel, input_shape)

# %% [code]
# Store and display the number of FLOPs. 
dict_flops['Convolutional Neural Networks: Regression'] = flops
print('FLOPs per inference:', flops)

# %% [code]
# Retrieve the average MAE and the standard error of the MAE. 
mae = [df_summary_reg['mae_baseline'].mean(), 
       df_summary_reg['mae_train'].mean(), 
       df_summary_reg['mae_valid'].mean(), 
       df_summary_reg['mae_test'].mean()]
mae_se = [np.std(df_summary_reg['mae_baseline']) / np.sqrt(df_summary_reg['mae_baseline'].shape[0]), 
          np.std(df_summary_reg['mae_train']) / np.sqrt(df_summary_reg['mae_train'].shape[0]), 
          np.std(df_summary_reg['mae_valid']) / np.sqrt(df_summary_reg['mae_valid'].shape[0]), 
          np.std(df_summary_reg['mae_test']) / np.sqrt(df_summary_reg['mae_test'].shape[0])]

# Display the average MAE and its standard error in a bar plot. 
fig, axes = plt.subplots(nrows = 1, ncols = 1, figsize = (6, 4))
axes.bar(range(4), mae, width = 0.6, color = ['green', 'blue', 'blue', 'blue'])
axes.errorbar(range(4), mae, yerr = mae_se, fmt = 'None', color = 'black')
axes.axhline(mae[0], 0, 1, color = 'green', linestyle = 'dashed', alpha = 0.5)
axes.set_xticks(range(4), labels = ['Baseline', 'Train', 'Valid', 'Test'])
axes.set_ylabel('Average MAE')
axes.set_title('Convolutional Neural Networks: Regression');

# %% [code]
# Select the subject with the best predictions. 
subject = subjects[np.argmax(df_summary_reg['r2_model'])]

# Plot the true values, the predictions, and the baseline for all the targets. 
fig, axes = plt.subplots(nrows = 17, ncols = 3, figsize = (12, 60))
fig.suptitle('Convolutional Neural Networks: Regression', fontsize = 14, y = 0.9)

# Iterate through all brain regions. 
for brain_region_index in range(len(targets)):

    # Retrieve the true values, the predictions, and the baseline for the selected subject and sequence length. 
    Y_te = dict_targets_sequence_reg[subject]['fmriNF'][:, brain_region_index]
    Y_pred = dict_predictions_reg[subject][:, brain_region_index]
    Baseline = dict_baseline_sequence_reg[subject]['eegfmriNF'][:, brain_region_index]

    # Compute the row and column indexes. 
    row_index = brain_region_index // 3
    column_index = brain_region_index % 3

    # Plot the true values, the predictions, and the baseline. 
    axes[row_index, column_index].plot(Y_te,  color = 'green', label = 'true values')
    axes[row_index, column_index].plot(Y_pred,  color = 'blue', label = 'predictions')
    axes[row_index, column_index].plot(Baseline,  color = 'orange', label = 'baseline')
    axes[row_index, column_index].legend()
    axes[row_index, column_index].set_title(targets[brain_region_index])

# Hide empty subplots. 
axes[5, 2].set_visible(False)
axes[16, 1].set_visible(False)
axes[16, 2].set_visible(False)

# %% [markdown]
## **3. Results**

# %% [code]
# Create directories for the results. 
if not os.path.isdir(results_path + 'classification/convolutional_neural_networks/'):
    os.mkdir(results_path + 'classification/convolutional_neural_networks/')
if not os.path.isdir(results_path + 'regression/convolutional_neural_networks/'):
    os.mkdir(results_path + 'regression/convolutional_neural_networks/')

# Save the models in the native Keras format. 
for subject in subjects:
    dict_models_class[subject].save(results_path + 'classification/convolutional_neural_networks/' + subject + '.keras')
    dict_models_reg[subject].save(results_path + 'regression/convolutional_neural_networks/' + subject + '.keras')

# Save the predictions into Pickle files. 
with open(results_path + 'classification/convolutional_neural_networks/dict_predictions_convolutional_neural_networks_class.p', 'wb') as file:
    pickle.dump(dict_predictions_class, file)
with open(results_path + 'regression/convolutional_neural_networks/dict_predictions_convolutional_neural_networks_reg.p', 'wb') as file:
    pickle.dump(dict_predictions_reg, file)

# Save the summaries into Pickle files. 
with open(results_path + 'classification/convolutional_neural_networks/df_summary_convolutional_neural_networks_class.p', 'wb') as file:
    pickle.dump(df_summary_class, file)
with open(results_path + 'regression/convolutional_neural_networks/df_summary_convolutional_neural_networks_reg.p', 'wb') as file:
    pickle.dump(df_summary_reg, file)

# Save the FLOPs into a Pickle file. 
with open(results_path + 'flops_convolutional_neural_networks.p', 'wb') as file:
    pickle.dump(dict_flops, file)
```
