## Supplementary material for "Neural Networks and Foundation Models: Two Strategies for EEG-to-fMRI Prediction": 07_Recurrent_Neural_Networks.ipynb

```
# %% [markdown]
# **Neural Networks and Foundation Models: Two Strategies for EEG-to-fMRI Prediction**
### **Supplementary Material**

# %% [markdown]
This project includes fourteen Notebooks:

*01: fMRI Preprocessing*

*02: EEG Preprocessing*

*03: Classification Models*

*04: Regression Models*

*05: Neural Networks*

*06: Convolutional Neural Networks*

***07: Recurrent Neural Networks***

*08: Transformers*

*09: Large Language Models*

*10: Large Language Model Chain-of-Thought*

*11: Large Language Model Fine-Tuning*

*12: Large Multimodal Model*

*13: Statistical Tests*

*14: Tables and Figures*

# %% [markdown]
### Notebook 07

# **Recurrent Neural Networks**

# %% [markdown]
## **1. Classification**

We define a **recurrent neural network model** for a classification task, using the **Adam optimizer** and the **binary cross-entropy loss function**. Our objective is to predict whether the fMRI signal increases or decreases in our brain regions of interest. We run the model on all subjects, except sub-xp102 who has a missing condition, and use the eegfmriNF condition as the train set, the eegNF condition as the validation set, and the fmriNF condition as the test set. 

# %% [code]
import numpy as np
import pandas as pd
import matplotlib.pyplot as plt
%matplotlib inline
import pickle
from sklearn.preprocessing import StandardScaler
import tensorflow as tf
import os
import logging

# %% [code]
# Define the raw data path. 
raw_data_path = '../ds002336-download/'

# Define the preprocessed data path. 
preprocessed_data_path = '../preprocessed_data/'

# Define the results path. 
results_path = '../results/'

# %% [code]
# Load the targets. 
with open(preprocessed_data_path + 'dict_targets.p', 'rb') as file:
    dict_targets = pickle.load(file)

# Load the features, the targets, and the baseline for the classification task. 
with open(preprocessed_data_path + 'classification/dict_features_classification_basis.p', 'rb') as file:
    dict_features_basis_class = pickle.load(file)
with open(preprocessed_data_path + 'classification/dict_features_classification_sequence.p', 'rb') as file:
    dict_features_sequence_class = pickle.load(file)
with open(preprocessed_data_path + 'classification/dict_targets_classification_basis.p', 'rb') as file:
    dict_targets_basis_class = pickle.load(file)
with open(preprocessed_data_path + 'classification/dict_baseline_classification_basis.p', 'rb') as file:
    dict_baseline_basis_class = pickle.load(file)

# %% [code]
# Define the list of subjects, removing sub-xp102 who has a missing condition. 
subjects = ['sub-xp1' + str(x).zfill(2) for x in range(1, 11)]
subjects.remove('sub-xp102')
subject = subjects[0]

# Retrieve the list of all targets. 
targets = list(dict_targets[subject]['eegfmriNF'].keys())

# Select a sequence length. 
selected_sequence_length = 5

# Retrieve the number of scans. 
nb_scans_class = dict_targets[subject]['eegfmriNF']['Frontal Pole'].shape[0] - 1
nb_scans_reg = dict_targets[subject]['eegfmriNF']['Frontal Pole'].shape[0]

# Define the batch size for the neural networks models. 
batch_size = 32

# Define a dictionary to store the FLOPS for each model. 
dict_flops = dict()

# %% [code]
# Define a function to run a recurrent neural network model for a classification task. 
def run_rnn_classification(NeuralNetworkModel, dict_features_basis, dict_targets_basis, dict_baseline_basis):

    # Define the dictionaries and the summary DataFrame. 
    dict_models = dict()
    dict_history = dict()
    dict_predictions = dict()
    df_summary = pd.DataFrame()

    # Iterate through all subjects. 
    for subject in subjects:

        # Display current subject. 
        print(subject)

        # Clone the Neural Network model. 
        PredictorBrain = tf.keras.models.clone_model(NeuralNetworkModel)

        # Retrieve the general features of the train set, validation set, and test set. 
        X_tr_basis = dict_features_basis[subject]['eegfmriNF']
        X_va_basis = dict_features_basis[subject]['eegNF']
        X_te_basis = dict_features_basis[subject]['fmriNF']

        # Rescale features variables. 
        scaler = StandardScaler()
        X_tr_basis_standardized = scaler.fit_transform(X_tr_basis)
        X_va_basis_standardized = scaler.transform(X_va_basis)
        X_te_basis_standardized = scaler.transform(X_te_basis)

        # Define the particular features arrays. 
        X_tr_standardized = np.zeros((nb_scans_class - selected_sequence_length, selected_sequence_length + 1, X_tr_basis_standardized.shape[1]))
        X_va_standardized = np.zeros((nb_scans_class - selected_sequence_length, selected_sequence_length + 1, X_va_basis_standardized.shape[1]))
        X_te_standardized = np.zeros((nb_scans_class - selected_sequence_length, selected_sequence_length + 1, X_te_basis_standardized.shape[1]))
        for scan in range(nb_scans_class - selected_sequence_length):
            for time_step in range(selected_sequence_length + 1):
                X_tr_standardized[scan, time_step, :] = X_tr_basis_standardized[scan + time_step, :].ravel()
                X_va_standardized[scan, time_step, :] = X_va_basis_standardized[scan + time_step, :].ravel()
                X_te_standardized[scan, time_step, :] = X_te_basis_standardized[scan + time_step, :].ravel()

        # Retrieve the targets of the train set, validation set, and test set. 
        Y_tr_basis = dict_targets_basis[subject]['eegfmriNF']
        Y_va_basis = dict_targets_basis[subject]['eegNF']
        Y_te_basis = dict_targets_basis[subject]['fmriNF']

        # Define the particular targets arrays. 
        Y_tr = Y_tr_basis[selected_sequence_length:, :]
        Y_va = Y_va_basis[selected_sequence_length:, :]
        Y_te = Y_te_basis[selected_sequence_length:, :]

        # Retrieve the baseline of the train set and determine the baseline accuracy. 
        Baseline = dict_baseline_basis[subject]['eegfmriNF'][selected_sequence_length:, :]
        accuracy_baseline = np.mean(Baseline == Y_te)

        # End training when accuracy stops improving. 
        early_stopping = tf.keras.callbacks.EarlyStopping(monitor = 'val_loss', patience = 6, 
                                                          restore_best_weights = True)

        # Define the loss function, optimizer, and metrics to track during training. 
        PredictorBrain.compile(optimizer = tf.keras.optimizers.Adam(learning_rate = 0.001), 
                               loss = 'binary_crossentropy', metrics = ['binary_accuracy'])

        # Fit the model. 
        NeuralNetworkHistory = PredictorBrain.fit(
            x = X_tr_standardized, y = Y_tr,
            validation_data = (X_va_standardized, Y_va), 
            batch_size = batch_size, epochs = 50,
            shuffle = True, callbacks = [early_stopping], verbose = 0
        )

        # Evaluate the test accuracy. 
        (loss_test, accuracy_test) = PredictorBrain.evaluate(X_te_standardized, Y_te, batch_size = batch_size)
        print(f'test loss: {loss_test:.4f}, test accuracy: {accuracy_test:.4f}')

        # Compute the predictions from the neural network. 
        Y_pred = PredictorBrain.predict(X_te_standardized)

        # Convert the probabilities to binary predictions. 
        Y_pred_binary = (Y_pred > 0.5).astype(int)

        # Store the performance metrics and the shapes of the arrays in the DataFrame. 
        df_summary.loc[subject, 'loss_train'] = NeuralNetworkHistory.history['loss'][-1]
        df_summary.loc[subject, 'loss_valid'] = NeuralNetworkHistory.history['val_loss'][-1]
        df_summary.loc[subject, 'loss_test'] = loss_test
        df_summary.loc[subject, 'accuracy_train'] = NeuralNetworkHistory.history['binary_accuracy'][-1]
        df_summary.loc[subject, 'accuracy_valid'] = NeuralNetworkHistory.history['val_binary_accuracy'][-1]
        df_summary.loc[subject, 'accuracy_test'] = accuracy_test
        df_summary.loc[subject, 'accuracy_baseline'] = accuracy_baseline
        df_summary.loc[subject, 'above_baseline'] = str(accuracy_test > accuracy_baseline)
        df_summary.loc[subject, 'X_tr.shape'] = str(X_tr_standardized.shape)
        df_summary.loc[subject, 'X_va.shape'] = str(X_va_standardized.shape)
        df_summary.loc[subject, 'X_te.shape'] = str(X_te_standardized.shape)
        df_summary.loc[subject, 'Y_tr.shape'] = str(Y_tr.shape)
        df_summary.loc[subject, 'Y_va.shape'] = str(Y_va.shape)
        df_summary.loc[subject, 'Y_te.shape'] = str(Y_te.shape)

        # Store the trained model, the model history, and the predictions in the dictionary. 
        dict_models[subject] = PredictorBrain
        dict_history[subject] = NeuralNetworkHistory
        dict_predictions[subject] = Y_pred_binary
    
    return dict_models, dict_history, dict_predictions, df_summary

# %% [code]
# Define a function to plot the results of the recurrent neural network training. 
def plot_rnn_classification(dict_history_class, title):
    
    # Display the graph. 
    fig, axes = plt.subplots(nrows = 9, ncols = 2, figsize = (12, 52))
    fig.suptitle(title, fontsize = 14, y = 0.895)
    subject_index = 0

    # Iterate through all subjects. 
    for subject in subjects:

        # Retrieve the history. 
        NeuralNetworkHistory = dict_history_class[subject]

        # Plot loss values. 
        axes[subject_index, 0].plot(NeuralNetworkHistory.history['loss'], label = 'train loss', color = 'blue')
        axes[subject_index, 0].plot(NeuralNetworkHistory.history['val_loss'], label = 'val loss', color = 'green')
        axes[subject_index, 0].set_title(str(subject) + ' valid. loss: {:.3f} (mean last 3)'.format(
            np.mean(NeuralNetworkHistory.history['val_loss'][-3:]) # Take the mean of the last three values. 
        ))
        axes[subject_index, 0].set_xlabel('Epoch')
        axes[subject_index, 0].set_ylabel('Loss value')
        axes[subject_index, 0].legend()

        # Plot accuracy values. 
        axes[subject_index, 1].plot(NeuralNetworkHistory.history['binary_accuracy'], label = 'train acc', color = 'blue')
        axes[subject_index, 1].plot(NeuralNetworkHistory.history['val_binary_accuracy'], label = 'val acc', color = 'green')
        axes[subject_index, 1].set_title(str(subject) + ' valid. accuracy: {:.3f} (mean last 3)'.format(
            np.mean(NeuralNetworkHistory.history['val_binary_accuracy'][-3:]) # Take the mean of the last three values. 
        ))
        axes[subject_index, 1].set_xlabel('Epoch')
        axes[subject_index, 1].set_ylabel('Accuracy')
        axes[subject_index, 1].legend();

        # Increment. 
        subject_index += 1

# %% [code]
# Define the neural network model. 
tf.random.set_seed(0)
NeuralNetworkModel = tf.keras.Sequential()
input_dim = (dict_features_sequence_class[subject]['eegfmriNF'].shape[0], # Number of time steps. 
             int(dict_features_sequence_class[subject]['eegfmriNF'].shape[1] / (selected_sequence_length + 1))) # Number of features per time step. 

# Input layer. 
NeuralNetworkModel.add(tf.keras.layers.InputLayer(shape = input_dim))

# GRU layer. 
NeuralNetworkModel.add(tf.keras.layers.GRU(
    units = 2048, activation = tf.keras.activations.relu, recurrent_dropout = 0.3, 
    return_sequences = True
    ))

# GRU layer. 
NeuralNetworkModel.add(tf.keras.layers.GRU(
    units = 512, activation = tf.keras.activations.relu, recurrent_dropout = 0.3, 
    return_sequences = True
    ))

# GRU layer. 
NeuralNetworkModel.add(tf.keras.layers.GRU(
    units = 256, activation = tf.keras.activations.relu, recurrent_dropout = 0.3, 
    return_sequences = True
    ))

# GRU layer. 
NeuralNetworkModel.add(tf.keras.layers.GRU(
    units = 64, activation = tf.keras.activations.relu, recurrent_dropout = 0.3
    ))
NeuralNetworkModel.add(tf.keras.layers.Dropout(0.5))

# Output layer. 
NeuralNetworkModel.add(tf.keras.layers.Dense(
    units = len(targets), activation = tf.keras.activations.sigmoid,
    kernel_initializer = tf.keras.initializers.VarianceScaling(scale = 1.0, seed = 0)))

# Print network summary. 
NeuralNetworkModel.summary()

# %% [code]
# Run the recurrent neural network model. 
dict_models_class, dict_history_class, dict_predictions_class, df_summary_class = run_rnn_classification(NeuralNetworkModel, 
                                                                                                         dict_features_basis_class, 
                                                                                                         dict_targets_basis_class, 
                                                                                                         dict_baseline_basis_class)

# %% [code]
# Display the results. 
print('Recurrent Neural Networks: Classification')
df_summary_class

# %% [code]
# Display the results. 
plot_rnn_classification(dict_history_class, title = 'Recurrent Neural Networks: Classification')

# %% [code]
# Define a function to measure the FLOPs of a model. 
def get_flops(model, input_shape):

    # Retrieve the graph of the concrete function. 
    concrete_func = tf.function(model).get_concrete_function(tf.TensorSpec(input_shape, tf.float32))
    graph = concrete_func.graph

    # Measure the FLOPs. 
    run_meta = tf.compat.v1.RunMetadata()
    opts = tf.compat.v1.profiler.ProfileOptionBuilder.float_operation()
    flops = tf.compat.v1.profiler.profile(graph = graph,
                                          run_meta = run_meta,
                                          cmd = 'op',
                                          options = opts)

    return flops.total_float_ops

# %% [code]
# Suppress TensorFlow warnings. 
tf.get_logger().setLevel(logging.ERROR)

# Measure the FLOPs of the model. 
input_shape = (1, *input_dim)  # Batch size = 1. Input dimension is entered with argument unpacking. 
flops = get_flops(NeuralNetworkModel, input_shape)

# %% [code]
# Store and display the number of FLOPs. 
dict_flops['Recurrent Neural Networks: Classification'] = flops
print('FLOPs per inference:', flops)

# %% [code]
# Retrieve the average accuracy and the standard error of the accuracy. 
acc = [df_summary_class['accuracy_baseline'].mean(), 
       df_summary_class['accuracy_train'].mean(), 
       df_summary_class['accuracy_valid'].mean(), 
       df_summary_class['accuracy_test'].mean()]
acc_se = [np.std(df_summary_class['accuracy_baseline']) / np.sqrt(df_summary_class['accuracy_baseline'].shape[0]), 
          np.std(df_summary_class['accuracy_train']) / np.sqrt(df_summary_class['accuracy_train'].shape[0]), 
          np.std(df_summary_class['accuracy_valid']) / np.sqrt(df_summary_class['accuracy_valid'].shape[0]), 
          np.std(df_summary_class['accuracy_test']) / np.sqrt(df_summary_class['accuracy_test'].shape[0])]

# Display the average accuracy and its standard error in a bar plot. 
fig, axes = plt.subplots(nrows = 1, ncols = 1, figsize = (6, 4))
axes.bar(range(4), acc, width = 0.6, color = ['green', 'blue', 'blue', 'blue'])
axes.errorbar(range(4), acc, yerr = acc_se, fmt = 'None', color = 'black')
axes.axhline(acc[0], 0, 1, color = 'green', linestyle = 'dashed', alpha = 0.5)
axes.set_xticks(range(4), labels = ['Baseline', 'Train', 'Valid', 'Test'])
axes.set_ylabel('Average accuracy')
axes.set_title('Recurrent Neural Networks: Classification');

# %% [markdown]
## **2. Regression**

We define a **recurrent neural network model** for a regression task, using the **Adam optimizer** and the **mean squared error loss function**. Our objective is to predict the value of the fMRI signal in our brain regions of interest. We run the model on all subjects, except sub-xp102 who has a missing condition, and use the eegfmriNF condition as the train set, the eegNF condition as the validation set, and the fmriNF condition as the test set. 

# %% [code]
# Load the features, the targets, and the baseline for the regression task. 
with open(preprocessed_data_path + 'regression/dict_features_regression_basis.p', 'rb') as file:
    dict_features_basis_reg = pickle.load(file)
with open(preprocessed_data_path + 'regression/dict_targets_regression_basis.p', 'rb') as file:
    dict_targets_basis_reg = pickle.load(file)
with open(preprocessed_data_path + 'regression/dict_targets_regression_sequence.p', 'rb') as file:
    dict_targets_sequence_reg = pickle.load(file)
with open(preprocessed_data_path + 'regression/dict_baseline_regression_basis.p', 'rb') as file:
    dict_baseline_basis_reg = pickle.load(file)
with open(preprocessed_data_path + 'regression/dict_baseline_regression_sequence.p', 'rb') as file:
    dict_baseline_sequence_reg = pickle.load(file)

# %% [code]
# Define the Mean Squared Error function. 
def MSE(y, y_pred):
    return np.mean(np.square(y - y_pred))

# %% [code]
# Define the Mean Absolute Error function. 
def MAE(y, y_pred):
    return np.mean(np.abs(y - y_pred))

# %% [code]
# Define the Residual Sum of Squares function. 
def RSS(y, y_pred):
    return np.sum(np.square(y - y_pred))

# %% [code]
# Define a function to run a recurrent neural network model for a regression task. 
def run_rnn_regression(NeuralNetworkModel, dict_features_basis, dict_targets_basis, dict_baseline_basis):

    # Define the dictionaries and the summary DataFrame. 
    dict_models = dict()
    dict_history = dict()
    dict_predictions = dict()
    df_summary = pd.DataFrame()

    # Iterate through all subjects. 
    for subject in subjects:

        # Display current subject. 
        print(subject)

        # Clone the Neural Network model. 
        PredictorBrain = tf.keras.models.clone_model(NeuralNetworkModel)

        # Retrieve the general features of the train set, validation set, and test set. 
        X_tr_basis = dict_features_basis[subject]['eegfmriNF']
        X_va_basis = dict_features_basis[subject]['eegNF']
        X_te_basis = dict_features_basis[subject]['fmriNF']

        # Rescale features variables. 
        scaler = StandardScaler()
        X_tr_basis_standardized = scaler.fit_transform(X_tr_basis)
        X_va_basis_standardized = scaler.transform(X_va_basis)
        X_te_basis_standardized = scaler.transform(X_te_basis)

        # Define the particular features arrays. 
        X_tr_standardized = np.zeros((nb_scans_reg - selected_sequence_length, selected_sequence_length + 1, X_tr_basis_standardized.shape[1]))
        X_va_standardized = np.zeros((nb_scans_reg - selected_sequence_length, selected_sequence_length + 1, X_va_basis_standardized.shape[1]))
        X_te_standardized = np.zeros((nb_scans_reg - selected_sequence_length, selected_sequence_length + 1, X_te_basis_standardized.shape[1]))
        for scan in range(nb_scans_reg - selected_sequence_length):
            for time_step in range(selected_sequence_length + 1):
                X_tr_standardized[scan, time_step, :] = X_tr_basis_standardized[scan + time_step, :].ravel()
                X_va_standardized[scan, time_step, :] = X_va_basis_standardized[scan + time_step, :].ravel()
                X_te_standardized[scan, time_step, :] = X_te_basis_standardized[scan + time_step, :].ravel()

        # Retrieve the targets of the train set, validation set, and test set. 
        Y_tr_basis = dict_targets_basis[subject]['eegfmriNF']
        Y_va_basis = dict_targets_basis[subject]['eegNF']
        Y_te_basis = dict_targets_basis[subject]['fmriNF']

        # Define the particular targets arrays. 
        Y_tr = Y_tr_basis[selected_sequence_length:, :]
        Y_va = Y_va_basis[selected_sequence_length:, :]
        Y_te = Y_te_basis[selected_sequence_length:, :]

        # Retrieve the baseline of the train set and determine the baseline accuracy. 
        Baseline = dict_baseline_basis[subject]['eegfmriNF'][selected_sequence_length:, :]

        # End training when accuracy stops improving. 
        early_stopping = tf.keras.callbacks.EarlyStopping(monitor = 'val_loss', patience = 6, 
                                                          restore_best_weights = True)

        # Define the loss function, optimizer, and metrics to track during training. 
        optimizer = tf.keras.optimizers.Adam(learning_rate = 0.005, beta_1 = 0.9)
        PredictorBrain.compile(optimizer = optimizer, loss = 'mse', metrics = [tf.keras.metrics.MeanAbsoluteError()])

        # Fit the model. 
        NeuralNetworkHistory = PredictorBrain.fit(
            x = X_tr_standardized, y = Y_tr,
            validation_data = (X_va_standardized, Y_va), 
            batch_size = batch_size, epochs = 50,
            shuffle = True, callbacks = [early_stopping], verbose = 0
        )

        # Evaluate the test MSE and MAE. 
        (mse_test, mae_test) = PredictorBrain.evaluate(X_te_standardized, Y_te, batch_size = batch_size)
        print(f'test MSE: {mse_test:.4f}, test MAE: {mae_test:.4f}')

        # Compute the predictions from the neural network. 
        Y_pred = PredictorBrain.predict(X_te_standardized)

        # Evaluate the performance metrics. 
        mse_baseline = MSE(Y_te, Baseline)
        mae_baseline = MAE(Y_te, Baseline)
        rss_baseline = RSS(Y_te, Baseline)
        mse_model = MSE(Y_te, Y_pred)
        mae_model = MAE(Y_te, Y_pred)
        rss_model = RSS(Y_te, Y_pred)
        r2_model = 1 - (rss_model / rss_baseline)

        # Store the performance metrics and the shapes of the arrays in the DataFrame. 
        df_summary.loc[subject, 'mse_train'] = NeuralNetworkHistory.history['loss'][-1]
        df_summary.loc[subject, 'mse_valid'] = NeuralNetworkHistory.history['val_loss'][-1]
        df_summary.loc[subject, 'mse_test'] = mse_test
        df_summary.loc[subject, 'mae_train'] = NeuralNetworkHistory.history['mean_absolute_error'][-1]
        df_summary.loc[subject, 'mae_valid'] = NeuralNetworkHistory.history['val_mean_absolute_error'][-1]
        df_summary.loc[subject, 'mae_test'] = mae_test
        df_summary.loc[subject, 'mae_baseline'] = mae_baseline
        df_summary.loc[subject, 'r2_model'] = r2_model
        df_summary.loc[subject, 'X_tr.shape'] = str(X_tr_standardized.shape)
        df_summary.loc[subject, 'X_va.shape'] = str(X_va_standardized.shape)
        df_summary.loc[subject, 'X_te.shape'] = str(X_te_standardized.shape)
        df_summary.loc[subject, 'Y_tr.shape'] = str(Y_tr.shape)
        df_summary.loc[subject, 'Y_va.shape'] = str(Y_va.shape)
        df_summary.loc[subject, 'Y_te.shape'] = str(Y_te.shape)

        # Store the trained model, the model history, and the predictions in the dictionary. 
        dict_models[subject] = PredictorBrain
        dict_history[subject] = NeuralNetworkHistory
        dict_predictions[subject] = Y_pred
    
    return dict_models, dict_history, dict_predictions, df_summary

# %% [code]
# Define a function to plot the results of the recurrent neural network training. 
def plot_rnn_regression(dict_history_reg, title):
    
    # Display the graph. 
    fig, axes = plt.subplots(nrows = 9, ncols = 2, figsize = (12, 52))
    fig.suptitle(title, fontsize = 14, y = 0.895)
    subject_index = 0

    # Iterate through all subjects. 
    for subject in subjects:

        # Retrieve the history. 
        NeuralNetworkHistory = dict_history_reg[subject]

        # Plot MSE values. 
        axes[subject_index, 0].plot(NeuralNetworkHistory.history['loss'], label = 'train MSE', color = 'blue')
        axes[subject_index, 0].plot(NeuralNetworkHistory.history['val_loss'], label = 'val MSE', color = 'green')
        axes[subject_index, 0].set_title(str(subject) + ' valid. MSE: {:.3f} (mean last 3)'.format(
            np.mean(NeuralNetworkHistory.history['val_loss'][-3:]) # Take the mean of the last three values. 
        ))
        axes[subject_index, 0].set_xlabel('Epoch')
        axes[subject_index, 0].set_ylabel('MSE')
        axes[subject_index, 0].legend()

        # Plot MAE values. 
        axes[subject_index, 1].plot(NeuralNetworkHistory.history['mean_absolute_error'], label = 'train MAE', color = 'blue')
        axes[subject_index, 1].plot(NeuralNetworkHistory.history['val_mean_absolute_error'], label = 'val MAE', color = 'green')
        axes[subject_index, 1].set_title(str(subject) + ' valid. MAE: {:.3f} (mean last 3)'.format(
            np.mean(NeuralNetworkHistory.history['val_mean_absolute_error'][-3:]) # Take the mean of the last three values. 
        ))
        axes[subject_index, 1].set_xlabel('Epoch')
        axes[subject_index, 1].set_ylabel('MAE')
        axes[subject_index, 1].legend();

        # Increment. 
        subject_index += 1

# %% [code]
# Define the neural network model. 
tf.random.set_seed(0)
NeuralNetworkModel = tf.keras.Sequential()
input_dim = (dict_features_sequence_class[subject]['eegfmriNF'].shape[0], # Number of time steps. 
             int(dict_features_sequence_class[subject]['eegfmriNF'].shape[1] / (selected_sequence_length + 1))) # Number of features per time step. 

# Input layer. 
NeuralNetworkModel.add(tf.keras.layers.InputLayer(shape = input_dim))

# GRU layer. 
NeuralNetworkModel.add(tf.keras.layers.GRU(
    units = 2048, activation = tf.keras.activations.relu, recurrent_dropout = 0.3, 
    return_sequences = True
    ))

# GRU layer. 
NeuralNetworkModel.add(tf.keras.layers.GRU(
    units = 512, activation = tf.keras.activations.relu, recurrent_dropout = 0.3, 
    return_sequences = True
    ))

# GRU layer. 
NeuralNetworkModel.add(tf.keras.layers.GRU(
    units = 256, activation = tf.keras.activations.relu, recurrent_dropout = 0.3, 
    return_sequences = True
    ))

# GRU layer. 
NeuralNetworkModel.add(tf.keras.layers.GRU(
    units = 64, activation = tf.keras.activations.relu, recurrent_dropout = 0.3
    ))
NeuralNetworkModel.add(tf.keras.layers.Dropout(0.5))

# Output layer. 
NeuralNetworkModel.add(tf.keras.layers.Dense(
    units = len(targets),
    ))

# Print network summary. 
NeuralNetworkModel.summary()

# %% [code]
# Run the recurrent neural network model. 
dict_models_reg, dict_history_reg, dict_predictions_reg, df_summary_reg = run_rnn_regression(NeuralNetworkModel, 
                                                                                             dict_features_basis_reg, 
                                                                                             dict_targets_basis_reg, 
                                                                                             dict_baseline_basis_reg)

# %% [code]
# Display the results. 
print('Recurrent Neural Networks: Regression')
df_summary_reg

# %% [code]
# Display the results. 
plot_rnn_regression(dict_history_reg, title = 'Recurrent Neural Networks: Regression')

# %% [code]
# Measure the FLOPs of the model. 
input_shape = (1, *input_dim)  # Batch size = 1. Input dimension is entered with argument unpacking. 
flops = get_flops(NeuralNetworkModel, input_shape)

# %% [code]
# Store and display the number of FLOPs. 
dict_flops['Recurrent Neural Networks: Regression'] = flops
print('FLOPs per inference:', flops)

# %% [code]
# Retrieve the average MAE and the standard error of the MAE. 
mae = [df_summary_reg['mae_baseline'].mean(), 
       df_summary_reg['mae_train'].mean(), 
       df_summary_reg['mae_valid'].mean(), 
       df_summary_reg['mae_test'].mean()]
mae_se = [np.std(df_summary_reg['mae_baseline']) / np.sqrt(df_summary_reg['mae_baseline'].shape[0]), 
          np.std(df_summary_reg['mae_train']) / np.sqrt(df_summary_reg['mae_train'].shape[0]), 
          np.std(df_summary_reg['mae_valid']) / np.sqrt(df_summary_reg['mae_valid'].shape[0]), 
          np.std(df_summary_reg['mae_test']) / np.sqrt(df_summary_reg['mae_test'].shape[0])]

# Display the average MAE and its standard error in a bar plot. 
fig, axes = plt.subplots(nrows = 1, ncols = 1, figsize = (6, 4))
axes.bar(range(4), mae, width = 0.6, color = ['green', 'blue', 'blue', 'blue'])
axes.errorbar(range(4), mae, yerr = mae_se, fmt = 'None', color = 'black')
axes.axhline(mae[0], 0, 1, color = 'green', linestyle = 'dashed', alpha = 0.5)
axes.set_xticks(range(4), labels = ['Baseline', 'Train', 'Valid', 'Test'])
axes.set_ylabel('Average MAE')
axes.set_title('Recurrent Neural Networks: Regression');

# %% [code]
# Select the subject with the best predictions. 
subject = subjects[np.argmax(df_summary_reg['r2_model'])]

# Plot the true values, the predictions, and the baseline for all the targets. 
fig, axes = plt.subplots(nrows = 17, ncols = 3, figsize = (12, 60))
fig.suptitle('Recurrent Neural Networks: Regression', fontsize = 14, y = 0.9)

# Iterate through all brain regions. 
for brain_region_index in range(len(targets)):

    # Retrieve the true values, the predictions, and the baseline for the selected subject and sequence length. 
    Y_te = dict_targets_sequence_reg[subject]['fmriNF'][:, brain_region_index]
    Y_pred = dict_predictions_reg[subject][:, brain_region_index]
    Baseline = dict_baseline_sequence_reg[subject]['eegfmriNF'][:, brain_region_index]

    # Compute the row and column indexes. 
    row_index = brain_region_index // 3
    column_index = brain_region_index % 3

    # Plot the true values, the predictions, and the baseline. 
    axes[row_index, column_index].plot(Y_te,  color = 'green', label = 'true values')
    axes[row_index, column_index].plot(Y_pred,  color = 'blue', label = 'predictions')
    axes[row_index, column_index].plot(Baseline,  color = 'orange', label = 'baseline')
    axes[row_index, column_index].legend()
    axes[row_index, column_index].set_title(targets[brain_region_index])

# Hide empty subplots. 
axes[5, 2].set_visible(False)
axes[16, 1].set_visible(False)
axes[16, 2].set_visible(False)

# %% [markdown]
## **3. Results**

# %% [code]
# Create directories for the results. 
if not os.path.isdir(results_path + 'classification/recurrent_neural_networks/'):
    os.mkdir(results_path + 'classification/recurrent_neural_networks/')
if not os.path.isdir(results_path + 'regression/recurrent_neural_networks/'):
    os.mkdir(results_path + 'regression/recurrent_neural_networks/')

# Save the models in the native Keras format. 
for subject in subjects:
    dict_models_class[subject].save(results_path + 'classification/recurrent_neural_networks/' + subject + '.keras')
    dict_models_reg[subject].save(results_path + 'regression/recurrent_neural_networks/' + subject + '.keras')

# Save the predictions into Pickle files. 
with open(results_path + 'classification/recurrent_neural_networks/dict_predictions_recurrent_neural_networks_class.p', 'wb') as file:
    pickle.dump(dict_predictions_class, file)
with open(results_path + 'regression/recurrent_neural_networks/dict_predictions_recurrent_neural_networks_reg.p', 'wb') as file:
    pickle.dump(dict_predictions_reg, file)

# Save the summaries into Pickle files. 
with open(results_path + 'classification/recurrent_neural_networks/df_summary_recurrent_neural_networks_class.p', 'wb') as file:
    pickle.dump(df_summary_class, file)
with open(results_path + 'regression/recurrent_neural_networks/df_summary_recurrent_neural_networks_reg.p', 'wb') as file:
    pickle.dump(df_summary_reg, file)

# Save the FLOPs into a Pickle file. 
with open(results_path + 'flops_recurrent_neural_networks.p', 'wb') as file:
    pickle.dump(dict_flops, file)
```
