## Supplementary material for "Neural Networks and Foundation Models: Two Strategies for EEG-to-fMRI Prediction": 09_Large_Language_Models.ipynb

```
# %% [markdown]
# **Neural Networks and Foundation Models: Two Strategies for EEG-to-fMRI Prediction**
### **Supplementary Material**

# %% [markdown]
This project includes fourteen Notebooks:

*01: fMRI Preprocessing*

*02: EEG Preprocessing*

*03: Classification Models*

*04: Regression Models*

*05: Neural Networks*

*06: Convolutional Neural Networks*

*07: Recurrent Neural Networks*

*08: Transformers*

***09: Large Language Models***

*10: Large Language Model Chain-of-Thought*

*11: Large Language Model Fine-Tuning*

*12: Large Multimodal Model*

*13: Statistical Tests*

*14: Tables and Figures*

# %% [markdown]
### Notebook 09

# **Large Language Models**

# %% [markdown]
## **1. Region-Channel Mapping**

We define a **region-channel mapping**, associating each brain region to the EEG channel whose signal is most likely to provide relevant information regarding its fMRI activity. This ad hoc mapping is based, in particular, on electrode proximity to the brain region. 

# %% [code]
import numpy as np
import pandas as pd
import matplotlib.pyplot as plt
%matplotlib inline
import pickle
import mne
import torch
from transformers import AutoModelForCausalLM, AutoTokenizer, logging as hf_logging
from fvcore.nn import FlopCountAnalysis
import logging

# %% [code]
# Define the raw data path. 
raw_data_path = '../ds002336-download/'

# Define the preprocessed data path. 
preprocessed_data_path = '../preprocessed_data/'

# Define the results path. 
results_path = '../results/'

# %% [code]
# Load the features and the targets for the classification task. 
with open(preprocessed_data_path + 'dict_targets.p', 'rb') as file:
    dict_targets = pickle.load(file)
with open(preprocessed_data_path + 'classification/dict_features_classification_basis.p', 'rb') as file:
    dict_features_basis_class = pickle.load(file)
with open(preprocessed_data_path + 'classification/dict_targets_classification_basis.p', 'rb') as file:
    dict_targets_basis_class = pickle.load(file)

# %% [code]
# Define the list of subjects, removing sub-xp102 who has a missing condition. 
subjects = ['sub-xp1' + str(x).zfill(2) for x in range(1, 11)]
subjects.remove('sub-xp102')
subject = subjects[0]

# Retrieve and display the total number of scans. 
nb_scans_total = dict_targets[subject]['eegfmriNF']['Frontal Pole'].shape[0]
nb_scans_total

# %% [code]
# Retrieve the list of all targets. 
targets = list(dict_targets[subject]['eegfmriNF'].keys())

# Display the brain region names and the number of brain regions. 
print(targets)
print(len(targets), 'brain regions')

# %% [code]
# Define the bandpower names. 
bandpower_names = ['Delta (1-4 Hz)', 'Theta (4-8 Hz)', 'Alpha (8-12 Hz)', 'Sigma (12-16 Hz)', 'Beta (16-30 Hz)', 'Gamma (30-40 Hz)']

# Define the path of the header file (.vhdr) for this subject, for the eegfmriNF condition. 
vhdr_path = raw_data_path + 'derivatives/' + subject + '/eeg_pp/' + subject + '_task-eegfmriNF_eeg_pp.vhdr'

# Load the EEG data using the MNE library, keeping error messages but suppressing warnings. 
channels_data = mne.io.read_raw_brainvision(vhdr_path, misc = ['ECG'], preload = True, verbose = 'ERROR')

# Remove the ECG channel from the channel names. 
channel_names = channels_data.ch_names
channel_names.remove('ECG')

# Display the channel names and the number of channels. 
print(channel_names)
print(len(channel_names), 'channels')

# %% [code]
# Define dictionaries to store the brain regions, sequences of fMRI scans, and FLOPs for each model. 
dict_brain_regions = dict()
dict_fmri_scans = dict()
dict_flops = dict()

# %% [code]
# Define for each brain region the EEG channel whose signal is most likely to provide relevant information regarding fMRI activity. 
region_channel_mapping = {
    'Frontal Pole': 'Fpz',
    'Insular Cortex': 'T7',
    'Superior Frontal Gyrus': 'Fz',
    'Middle Frontal Gyrus': 'F3',
    'Inferior Frontal Gyrus, pars triangularis': 'F7',
    'Inferior Frontal Gyrus, pars opercularis': 'F7',
    'Precentral Gyrus': 'C3',
    'Temporal Pole': 'FT9',
    'Superior Temporal Gyrus, anterior division': 'FT7',
    'Superior Temporal Gyrus, posterior division': 'T7',
    'Middle Temporal Gyrus, anterior division': 'T7',
    'Middle Temporal Gyrus, posterior division': 'T7',
    'Middle Temporal Gyrus, temporooccipital part': 'P7',
    'Inferior Temporal Gyrus, anterior division': 'T7',
    'Inferior Temporal Gyrus, posterior division': 'P7',
    'Inferior Temporal Gyrus, temporooccipital part': 'PO7',
    'Postcentral Gyrus': 'C3',
    'Superior Parietal Lobule': 'Pz',
    'Supramarginal Gyrus, anterior division': 'CP3',
    'Supramarginal Gyrus, posterior division': 'P3',
    'Angular Gyrus': 'P3',
    'Lateral Occipital Cortex, superior division': 'PO3',
    'Lateral Occipital Cortex, inferior division': 'O1',
    'Intracalcarine Cortex': 'Oz',
    'Frontal Medial Cortex': 'Fz',
    'Juxtapositional Lobule Cortex (formerly Supplementary Motor Cortex)': 'Cz',
    'Subcallosal Cortex': 'Fpz',
    'Paracingulate Gyrus': 'Fz',
    'Cingulate Gyrus, anterior division': 'Fz',
    'Cingulate Gyrus, posterior division': 'Pz',
    'Precuneous Cortex': 'Pz',
    'Cuneal Cortex': 'Oz',
    'Frontal Orbital Cortex': 'Fp1',
    'Parahippocampal Gyrus, anterior division': 'FT9',
    'Parahippocampal Gyrus, posterior division': 'TP9',
    'Lingual Gyrus': 'Oz',
    'Temporal Fusiform Cortex, anterior division': 'FT9',
    'Temporal Fusiform Cortex, posterior division': 'TP9',
    'Temporal Occipital Fusiform Cortex': 'PO7',
    'Occipital Fusiform Gyrus': 'PO7',
    'Frontal Opercular Cortex': 'FC5',
    'Central Opercular Cortex': 'C3',
    'Parietal Opercular Cortex': 'CP3',
    'Planum Polare': 'FT7',
    "Heschl's Gyrus (includes H1 and H2)": 'T7',
    'Planum Temporale': 'TP7',
    'Supracalcarine Cortex': 'Oz',
    'Occipital Pole': 'Oz'
}

# %% [markdown]
## **2. Gemma**

We use the **Gemma-2-2B-IT large language model**, and select its hyperparameters in order to ensure a relative variety of responses. We define a prompt where we provide a sequence of EEG bandpower values, and ask the model to predict the corresponding fMRI activity. We run the large language model on the eegfmriNF condition of all subjects, for a selection of brain regions spanning both a substantial fraction of the brain and a variety of cognitive functions. In the bar plot presenting the results, an indicative baseline is given for visualization purposes, but the real baseline depends both on the selected fMRI scans and the distribution of missing or ambiguous predictions. 

# %% [code]
# Define the LLM to use. 
model_name = 'google/gemma-2-2b-it'

# Load the tokenizer and the model. 
tokenizer_gemma = AutoTokenizer.from_pretrained(model_name)
model_gemma = AutoModelForCausalLM.from_pretrained(model_name, torch_dtype = torch.float16, device_map = 'auto')

# %% [code]
# Define a function to create a LLM prompt with a sequence of bandpower values. 
def prompt_llm(X, fmri_scan_index, sequence_length, selected_bandpowers, selected_eeg_channel, selected_brain_region):

    # Retrieve the sequence of features. 
    X_sequence = X[(fmri_scan_index - sequence_length):fmri_scan_index, :]

    # Create the prompt. 
    bandpowers_results = ', '.join(
        f"{bandpower_names[bandpower_index]}: {X_sequence[:, bandpower_index]}" for bandpower_index in selected_bandpowers
    )
    prompt = (
        f"A human subject participated in a neuroscience experiment where EEG data and fMRI data were recorded simultaneously. "
        f"The EEG band powers were measured every two seconds at electrode {selected_eeg_channel}, "
        f"with the following results: {bandpowers_results}. "
        f"The fMRI BOLD signal was measured in the {selected_brain_region} during the last two seconds. "
        f"Given the EEG data, is the fMRI signal in this brain region likely increasing or decreasing during these last two seconds? "
        f"Base your answer on your general knowledge in neuroscience, EEG research, and fMRI research. "
        f"Since this is a time series, you might need to take into account the hemodynamic response function (HRF), "
        f"and the fact that after an event, the fMRI response is delayed compared to the EEG response. "
        f"Please answer with only one word: Increasing or Decreasing. Just give your best prediction, without any explanation."
    )

    return prompt

# %% [code]
# Define a function to run a LLM. 
def run_llm(prompt, tokenizer, model):

    # Run the LLM. 
    inputs = tokenizer(prompt, return_tensors = 'pt').to(model.device)
    output = model.generate(**inputs, 
                            max_new_tokens = 20, 
                            temperature = 1.2, 
                            top_p = 0.9, 
                            top_k = 50, 
                            do_sample = True, 
                            repetition_penalty = 1)

    # Decode the generated text, remove the original prompt, and return the answer. 
    generated_text = tokenizer.decode(output[0], skip_special_tokens = True)
    answer_only = generated_text[len(prompt):].strip()
    
    return answer_only

# %% [code]
# Define a function to run a LLM multiple times. 
def run_llm_series(X, Y, fmri_scans, sequence_length, selected_bandpowers, selected_eeg_channel, selected_brain_region, tokenizer, model):

    # Create the NumPy arrays to store the true values and predictions.     
    Y_true = np.zeros(len(fmri_scans))
    Y_pred = np.zeros(len(fmri_scans))
    answers = []
    counter = 0

    # Iterate through all selected fMRI scans. 
    for fmri_scan_index in fmri_scans:

        # Create the LLM prompt. 
        prompt = prompt_llm(X, 
                            fmri_scan_index = fmri_scan_index, 
                            sequence_length = sequence_length, 
                            selected_bandpowers = selected_bandpowers, 
                            selected_eeg_channel = selected_eeg_channel, 
                            selected_brain_region = selected_brain_region)
        
        # Run the LLM and store the result. 
        answer_only = run_llm(prompt, tokenizer, model)
        answers.append(answer_only) # For debugging purposes. 

        # Analyze the answer. 
        increase_detected = 'increas' in answer_only.lower()
        decrease_detected = 'decreas' in answer_only.lower()
        if increase_detected and not decrease_detected:
            Y_pred[counter] = 1
        elif decrease_detected and not increase_detected:
            Y_pred[counter] = 0
        else:
            Y_pred[counter] = -1

        # Store the true value.
        Y_true[counter] = Y[fmri_scan_index]

        # Increment. 
        counter += 1

    return Y_true, Y_pred

# %% [code]
# Select a subject and a condition. 
subject = 'sub-xp101'
condition = 'eegfmriNF'

# Select a fMRI brain region and retrieve its corresponding EEG channel. 
selected_brain_region = 'Frontal Pole'
selected_brain_region_index = list(targets).index(selected_brain_region)
selected_eeg_channel = region_channel_mapping[selected_brain_region]
selected_eeg_channel_index = channel_names.index(selected_eeg_channel)

# Display the fMRI brain region and its corresponding EEG channel. 
print('Brain region:', selected_brain_region)
print('Brain region index:', int(selected_brain_region_index))
print('EEG channel:', selected_eeg_channel)
print('EEG channel index:', int(selected_eeg_channel_index))

# %% [code]
# Retrieve the features for the selected subject, condition, and EEG channel. 
X = dict_features_basis_class[subject][condition][:, (selected_eeg_channel_index * len(bandpower_names)):(selected_eeg_channel_index * len(bandpower_names)) + len(bandpower_names)]

# Retrieve the targets for the selected subject, condition, and fMRI brain region. 
Y = dict_targets_basis_class[subject][condition][:, selected_brain_region_index]

# Select the sequence length and the bandpowers. 
sequence_length = 5
selected_bandpowers = [0, 1, 2, 4]

# Create the LLM prompt. 
prompt = prompt_llm(X, 
                    fmri_scan_index = 5, 
                    sequence_length = sequence_length, 
                    selected_bandpowers = selected_bandpowers, 
                    selected_eeg_channel = selected_eeg_channel, 
                    selected_brain_region = selected_brain_region)
prompt

# %% [code]
# Run the LLM once, and display the result. 
answer_only = run_llm(prompt, tokenizer_gemma, model_gemma)
answer_only

# %% [code]
# Run the LLM multiple times. 
fmri_scans = range(5, (nb_scans_total - 1), 10)
Y_true, Y_pred = run_llm_series(X, Y, fmri_scans, sequence_length, selected_bandpowers, selected_eeg_channel, selected_brain_region, tokenizer_gemma, model_gemma)

# Compute the average accuracy. 
accuracy_array = Y_pred[Y_pred != -1] == Y_true[Y_pred != -1]
accuracy_sequence = np.sum(accuracy_array) / accuracy_array.shape[0]
accuracy_sequence

# %% [code]
# Select a condition and a list of brain regions. 
condition = 'eegfmriNF'
brain_regions = ['Frontal Pole', # High-order cognitive functions, but sometimes noisy EEG signal. 
                 'Insular Cortex', # Integrative functions. 
                 'Inferior Frontal Gyrus, pars triangularis', # Language and executive functions. 
                 'Precentral Gyrus', # Primary motor cortex. 
                 'Superior Temporal Gyrus, posterior division', # Auditory functions. 
                 'Superior Parietal Lobule', # Attention functions. 
                 'Angular Gyrus', # Default-mode network. 
                 'Occipital Pole'] # Primary visual cortex. 

# Select a sequence of fMRI scans. 
fmri_scans = range(5, (nb_scans_total - 1), 10) # Extensive search.
#fmri_scans = range(5, (nb_scans_total - 1), 100) # Quick search for debugging.

# Store the brain regions and the sequence of fMRI scans in dictionaries. 
dict_brain_regions['Large Language Models: Gemma'] = brain_regions
dict_fmri_scans['Large Language Models: Gemma'] = fmri_scans

# Define a dictionary to store the predictions and a Pandas DataFrame to store the accuracy. 
dict_predictions_gemma = dict()
accuracy_summary_gemma = pd.DataFrame(columns = brain_regions)

# Iterate through all brain regions. 
for selected_brain_region in brain_regions:

    # Display the current brain region. 
    print(selected_brain_region)

    # Retrieve the EEG channel corresponding to the brain region. 
    selected_brain_region_index = list(targets).index(selected_brain_region)
    selected_eeg_channel = region_channel_mapping[selected_brain_region]
    selected_eeg_channel_index = channel_names.index(selected_eeg_channel)

    # Define a dictionary to store the predictions for this brain region. 
    dict_predictions_gemma[selected_brain_region] = dict()

    # Iterate through all subjects. 
    for subject in subjects:
        
        # Retrieve the features for the selected subject, condition, and EEG channel. 
        X = dict_features_basis_class[subject][condition][:, (selected_eeg_channel_index * len(bandpower_names)):(selected_eeg_channel_index * len(bandpower_names)) + len(bandpower_names)]

        # Retrieve the targets for the selected subject, condition, and fMRI brain region. 
        Y = dict_targets_basis_class[subject][condition][:, selected_brain_region_index]

        # Run the LLM multiple times. 
        Y_true, Y_pred = run_llm_series(X, Y, fmri_scans, sequence_length, selected_bandpowers, selected_eeg_channel, selected_brain_region, tokenizer_gemma, model_gemma)

        # Store the predictions. 
        dict_predictions_gemma[selected_brain_region][subject] = Y_pred

        # Compute and store the average accuracy. 
        accuracy_array = Y_pred[Y_pred != -1] == Y_true[Y_pred != -1]
        accuracy_sequence = np.sum(accuracy_array) / accuracy_array.shape[0]
        accuracy_summary_gemma.loc[subject, selected_brain_region] = accuracy_sequence

        # Display the number of ambiguous predictions. 
        print(subject + ': ' + str(np.sum(Y_pred == -1)), 'ambiguous prediction(s) out of', len(Y_pred), '-', str(np.sum(Y_pred == -1) / len(Y_pred) * 100), '%')

# %% [code]
# Set the logging level to suppress warnings. 
logging.getLogger('fvcore.nn.jit_handles').setLevel(logging.ERROR)
logging.getLogger('fvcore.nn.jit_analysis').setLevel(logging.ERROR)

# Store current device. 
device_before = next(model_gemma.parameters()).device
print('Original device of the model:', device_before)

# Define a class wrapping the model. 
# We output only logits (Tensor data type) for compatibility with FlopCountAnalysis. 
class WrappedModel(torch.nn.Module):
    def __init__(self, model):
        super().__init__()
        self.model = model
    def forward(self, input_ids):
        output = self.model(input_ids)
        return output.logits

# Move the model to CPU and prepare dummy input for FLOPs analysis. 
model_gemma_cpu = WrappedModel(model_gemma).to('cpu')
dummy_input = (torch.ones((1, len(prompt)), dtype = torch.long),) # batch_size = 1, seq_len = prompt length
print('Prompt length:', len(prompt))

# Measure the FLOPs. 
flops = FlopCountAnalysis(model_gemma_cpu, dummy_input)
print(f'Total FLOPs: {flops.total():,}')
print('FLOPs by operator:', flops.by_operator())

# Move the model back to its original device. 
model_gemma.to(device_before)
print('Model moved back to:', device_before)

# %% [code]
# Store and display the number of FLOPs. 
dict_flops['Large Language Models: Gemma'] = flops.total()
print('FLOPs per inference:', flops.total())

# %% [code]
# Display the accuracies per subject and brain region. 
accuracy_summary_gemma

# %% [code]
# Retrieve the average accuracy across subjects, and the standard error of the accuracy across subjects. 
accuracy_mean = list(accuracy_summary_gemma.mean().values)
accuracy_mean.insert(0, 0.5) # Insert a baseline average of 0.5. 
accuracy_se = list(accuracy_summary_gemma.std().values / np.sqrt(len(subjects)))
accuracy_se.insert(0, 0) # Insert a baseline standard error of 0. 

# Append the total average accuracy and its standard error. 
nb_brain_regions = accuracy_summary_gemma.shape[1]
accuracy_mean.append(np.mean(accuracy_mean[1:]))
accuracy_se.append(np.std(accuracy_mean[1:]) / (np.sqrt(len(subjects) * nb_brain_regions)))

# Create the labels. 
labels = brain_regions.copy()
labels.insert(0, 'Baseline')
labels.append('Total')

# Display the average accuracy and its standard error in a bar plot. 
fig, axes = plt.subplots(nrows = 1, ncols = 1, figsize = (10, 4))
axes.bar(range(nb_brain_regions + 2), accuracy_mean, width = 0.6, color = ['green', *['blue'] * nb_brain_regions, 'mediumblue'])
axes.errorbar(range(nb_brain_regions + 2), accuracy_mean, yerr = accuracy_se, fmt = 'None', color = 'black')
axes.axhline(accuracy_mean[0], 0, 1, color = 'green', linestyle = 'dashed', alpha = 0.5)
axes.set_xticks(range(nb_brain_regions + 2), labels = labels)
axes.set_xticklabels(labels, rotation = 45, ha = 'right')
axes.set_ylabel('Average accuracy across subjects')
axes.set_title('Gemma-2-2B-IT');

# %% [markdown]
## **3. Llama**

We use the **Llama-3.2-3B-Instruct large language model**, selecting the same hyperparameters as for the Gemma-2-2B-IT model, and defining the prompt in the same way. We run the large language model on the eegfmriNF condition of all subjects, for the same selection of brain regions. In the bar plot presenting the results, an indicative baseline is given for visualization purposes, but the real baseline depends both on the selected fMRI scans and the distribution of missing or ambiguous predictions. 

# %% [code]
# Define the LLM to use. 
model_name = 'meta-llama/Llama-3.2-3B-Instruct'

# Load the tokenizer and the model. 
tokenizer_llama = AutoTokenizer.from_pretrained(model_name)
model_llama = AutoModelForCausalLM.from_pretrained(model_name, torch_dtype = torch.float16, device_map = 'auto')

# %% [code]
# Select a subject and a condition. 
subject = 'sub-xp101'
condition = 'eegfmriNF'

# Select a fMRI brain region and retrieve its corresponding EEG channel. 
selected_brain_region = 'Frontal Pole'
selected_brain_region_index = list(targets).index(selected_brain_region)
selected_eeg_channel = region_channel_mapping[selected_brain_region]
selected_eeg_channel_index = channel_names.index(selected_eeg_channel)

# Display the fMRI brain region and its corresponding EEG channel. 
print('Brain region:', selected_brain_region)
print('Brain region index:', int(selected_brain_region_index))
print('EEG channel:', selected_eeg_channel)
print('EEG channel index:', int(selected_eeg_channel_index))

# %% [code]
# Retrieve the features for the selected subject, condition, and EEG channel. 
X = dict_features_basis_class[subject][condition][:, (selected_eeg_channel_index * len(bandpower_names)):(selected_eeg_channel_index * len(bandpower_names)) + len(bandpower_names)]

# Retrieve the targets for the selected subject, condition, and fMRI brain region. 
Y = dict_targets_basis_class[subject][condition][:, selected_brain_region_index]

# Select the sequence length and the bandpowers. 
sequence_length = 5
selected_bandpowers = [0, 1, 2, 4]

# Create the LLM prompt. 
prompt = prompt_llm(X, 
                    fmri_scan_index = 5, 
                    sequence_length = sequence_length, 
                    selected_bandpowers = selected_bandpowers, 
                    selected_eeg_channel = selected_eeg_channel, 
                    selected_brain_region = selected_brain_region)
prompt

# %% [code]
# Suppress the transformers library logging. 
hf_logging.set_verbosity_error()

# Run the LLM once, and display the result. 
answer_only = run_llm(prompt, tokenizer_llama, model_llama)
answer_only

# %% [code]
# Run the LLM multiple times. 
fmri_scans = range(5, (nb_scans_total - 1), 10)
Y_true, Y_pred = run_llm_series(X, Y, fmri_scans, sequence_length, selected_bandpowers, selected_eeg_channel, selected_brain_region, tokenizer_llama, model_llama)

# Compute the average accuracy. 
accuracy_array = Y_pred[Y_pred != -1] == Y_true[Y_pred != -1]
accuracy_sequence = np.sum(accuracy_array) / accuracy_array.shape[0]
accuracy_sequence

# %% [code]
# Select a condition and a list of brain regions. 
condition = 'eegfmriNF'
brain_regions = ['Frontal Pole', # High-order cognitive functions, but sometimes noisy EEG signal. 
                 'Insular Cortex', # Integrative functions. 
                 'Inferior Frontal Gyrus, pars triangularis', # Language and executive functions. 
                 'Precentral Gyrus', # Primary motor cortex. 
                 'Superior Temporal Gyrus, posterior division', # Auditory functions. 
                 'Superior Parietal Lobule', # Attention functions. 
                 'Angular Gyrus', # Default-mode network. 
                 'Occipital Pole'] # Primary visual cortex. 

# Select a sequence of fMRI scans. 
fmri_scans = range(5, (nb_scans_total - 1), 10) # Extensive search.
#fmri_scans = range(5, (nb_scans_total - 1), 100) # Quick search for debugging.

# Store the brain regions and the sequence of fMRI scans in dictionaries. 
dict_brain_regions['Large Language Models: Llama'] = brain_regions
dict_fmri_scans['Large Language Models: Llama'] = fmri_scans

# Define a dictionary to store the predictions and a Pandas DataFrame to store the accuracy. 
dict_predictions_llama = dict()
accuracy_summary_llama = pd.DataFrame(columns = brain_regions)

# Iterate through all brain regions. 
for selected_brain_region in brain_regions:

    # Display the current brain region. 
    print(selected_brain_region)

    # Retrieve the EEG channel corresponding to the brain region. 
    selected_brain_region_index = list(targets).index(selected_brain_region)
    selected_eeg_channel = region_channel_mapping[selected_brain_region]
    selected_eeg_channel_index = channel_names.index(selected_eeg_channel)

    # Define a dictionary to store the predictions for this brain region. 
    dict_predictions_llama[selected_brain_region] = dict()

    # Iterate through all subjects. 
    for subject in subjects:
        
        # Retrieve the features for the selected subject, condition, and EEG channel. 
        X = dict_features_basis_class[subject][condition][:, (selected_eeg_channel_index * len(bandpower_names)):(selected_eeg_channel_index * len(bandpower_names)) + len(bandpower_names)]

        # Retrieve the targets for the selected subject, condition, and fMRI brain region. 
        Y = dict_targets_basis_class[subject][condition][:, selected_brain_region_index]

        # Run the LLM multiple times. 
        Y_true, Y_pred = run_llm_series(X, Y, fmri_scans, sequence_length, selected_bandpowers, selected_eeg_channel, selected_brain_region, tokenizer_llama, model_llama)

        # Store the predictions. 
        dict_predictions_llama[selected_brain_region][subject] = Y_pred
        
        # Compute and store the average accuracy. 
        accuracy_array = Y_pred[Y_pred != -1] == Y_true[Y_pred != -1]
        accuracy_sequence = np.sum(accuracy_array) / accuracy_array.shape[0]
        accuracy_summary_llama.loc[subject, selected_brain_region] = accuracy_sequence

        # Display the number of ambiguous predictions. 
        print(subject + ': ' + str(np.sum(Y_pred == -1)), 'ambiguous prediction(s) out of', len(Y_pred), '-', str(np.sum(Y_pred == -1) / len(Y_pred) * 100), '%')

# %% [code]
# Store current device. 
device_before = next(model_llama.parameters()).device
print('Original device of the model:', device_before)

# Move the model to CPU and prepare dummy input for FLOPs analysis. 
model_llama_cpu = WrappedModel(model_llama).to('cpu')
dummy_input = (torch.ones((1, len(prompt)), dtype = torch.long),) # batch_size = 1, seq_len = prompt length
print('Prompt length:', len(prompt))

# Measure the FLOPs. 
flops = FlopCountAnalysis(model_llama_cpu, dummy_input)
print(f'Total FLOPs: {flops.total():,}')
print('FLOPs by operator:', flops.by_operator())

# Move the model back to its original device. 
model_llama.to(device_before)
print('Model moved back to:', device_before)

# %% [code]
# Store and display the number of FLOPs. 
dict_flops['Large Language Models: Llama'] = flops.total()
print('FLOPs per inference:', flops.total())

# %% [code]
# Display the accuracies per subject and brain region. 
accuracy_summary_llama

# %% [code]
# Retrieve the average accuracy across subjects, and the standard error of the accuracy across subjects. 
accuracy_mean = list(accuracy_summary_llama.mean().values)
accuracy_mean.insert(0, 0.5) # Insert a baseline average of 0.5. 
accuracy_se = list(accuracy_summary_llama.std().values / np.sqrt(len(subjects)))
accuracy_se.insert(0, 0) # Insert a baseline standard error of 0. 

# Append the total average accuracy and its standard error. 
nb_brain_regions = accuracy_summary_llama.shape[1]
accuracy_mean.append(np.mean(accuracy_mean[1:]))
accuracy_se.append(np.std(accuracy_mean[1:]) / (np.sqrt(len(subjects) * nb_brain_regions)))

# Create the labels. 
labels = brain_regions.copy()
labels.insert(0, 'Baseline')
labels.append('Total')

# Display the average accuracy and its standard error in a bar plot. 
fig, axes = plt.subplots(nrows = 1, ncols = 1, figsize = (10, 4))
axes.bar(range(nb_brain_regions + 2), accuracy_mean, width = 0.6, color = ['green', *['blue'] * nb_brain_regions, 'mediumblue'])
axes.errorbar(range(nb_brain_regions + 2), accuracy_mean, yerr = accuracy_se, fmt = 'None', color = 'black')
axes.axhline(accuracy_mean[0], 0, 1, color = 'green', linestyle = 'dashed', alpha = 0.5)
axes.set_xticks(range(nb_brain_regions + 2), labels = labels)
axes.set_xticklabels(labels, rotation = 45, ha = 'right')
axes.set_ylabel('Average accuracy across subjects')
axes.set_title('Llama-3.2-3B-Instruct');

# %% [markdown]
## **4. Results**

# %% [code]
# Save the region-channel mapping into a Pickle file. 
with open(results_path + 'region_channel_mapping.p', 'wb') as file:
    pickle.dump(region_channel_mapping, file)

# Save the predictions into Pickle files. 
with open(results_path + 'classification/dict_predictions_gemma.p', 'wb') as file:
    pickle.dump(dict_predictions_gemma, file)
with open(results_path + 'classification/dict_predictions_llama.p', 'wb') as file:
    pickle.dump(dict_predictions_llama, file)

# Save the summaries into Pickle files. 
with open(results_path + 'classification/df_summary_gemma.p', 'wb') as file:
    pickle.dump(accuracy_summary_gemma, file)
with open(results_path + 'classification/df_summary_llama.p', 'wb') as file:
    pickle.dump(accuracy_summary_llama, file)

# Save the brain regions, sequences of fMRI scans, and FLOPs into Pickle files. 
with open(results_path + 'brain_regions_large_language_models.p', 'wb') as file:
    pickle.dump(dict_brain_regions, file)
with open(results_path + 'fmri_scans_large_language_models.p', 'wb') as file:
    pickle.dump(dict_fmri_scans, file)
with open(results_path + 'flops_large_language_models.p', 'wb') as file:
    pickle.dump(dict_flops, file)
```
