## Supplementary material for "Neural Networks and Foundation Models: Two Strategies for EEG-to-fMRI Prediction": 10_Large_Language_Model_Chain-of-Thought.ipynb

```
# %% [markdown]
# **Neural Networks and Foundation Models: Two Strategies for EEG-to-fMRI Prediction**
### **Supplementary Material**

# %% [markdown]
This project includes fourteen Notebooks:

*01: fMRI Preprocessing*

*02: EEG Preprocessing*

*03: Classification Models*

*04: Regression Models*

*05: Neural Networks*

*06: Convolutional Neural Networks*

*07: Recurrent Neural Networks*

*08: Transformers*

*09: Large Language Models*

***10: Large Language Model Chain-of-Thought***

*11: Large Language Model Fine-Tuning*

*12: Large Multimodal Model*

*13: Statistical Tests*

*14: Tables and Figures*

# %% [markdown]
### Notebook 10

# **Large Language Model Chain-of-Thought**

# %% [markdown]
## **1. Region-Channel Mapping**

We retrieve the **region-channel mapping**, associating each brain region to the EEG channel whose signal is most likely to provide relevant information regarding its fMRI activity. 

# %% [code]
import numpy as np
import pandas as pd
import matplotlib.pyplot as plt
%matplotlib inline
import pickle
import mne
import torch
from transformers import AutoModelForCausalLM, AutoTokenizer
from fvcore.nn import FlopCountAnalysis
import logging

# %% [code]
# Define the raw data path. 
raw_data_path = '../ds002336-download/'

# Define the preprocessed data path. 
preprocessed_data_path = '../preprocessed_data/'

# Define the results path. 
results_path = '../results/'

# %% [code]
# Load the features and the targets for the classification task. 
with open(preprocessed_data_path + 'dict_targets.p', 'rb') as file:
    dict_targets = pickle.load(file)
with open(preprocessed_data_path + 'classification/dict_features_classification_basis.p', 'rb') as file:
    dict_features_basis_class = pickle.load(file)
with open(preprocessed_data_path + 'classification/dict_targets_classification_basis.p', 'rb') as file:
    dict_targets_basis_class = pickle.load(file)

# Load the region-channel mapping. 
with open(results_path + 'region_channel_mapping.p', 'rb') as file:
    region_channel_mapping = pickle.load(file)

# %% [code]
# Define the list of subjects, removing sub-xp102 who has a missing condition. 
subjects = ['sub-xp1' + str(x).zfill(2) for x in range(1, 11)]
subjects.remove('sub-xp102')
subject = subjects[0]

# Retrieve and display the total number of scans. 
nb_scans_total = dict_targets[subject]['eegfmriNF']['Frontal Pole'].shape[0]
nb_scans_total

# %% [code]
# Retrieve the list of all targets. 
targets = list(dict_targets[subject]['eegfmriNF'].keys())

# Display the brain region names and the number of brain regions. 
print(targets)
print(len(targets), 'brain regions')

# %% [code]
# Define the bandpower names. 
bandpower_names = ['Delta (1-4 Hz)', 'Theta (4-8 Hz)', 'Alpha (8-12 Hz)', 'Sigma (12-16 Hz)', 'Beta (16-30 Hz)', 'Gamma (30-40 Hz)']

# Define the path of the header file (.vhdr) for this subject, for the eegfmriNF condition. 
vhdr_path = raw_data_path + 'derivatives/' + subject + '/eeg_pp/' + subject + '_task-eegfmriNF_eeg_pp.vhdr'

# Load the EEG data using the MNE library, keeping error messages but suppressing warnings. 
channels_data = mne.io.read_raw_brainvision(vhdr_path, misc = ['ECG'], preload = True, verbose = 'ERROR')

# Remove the ECG channel from the channel names. 
channel_names = channels_data.ch_names
channel_names.remove('ECG')

# Display the channel names and the number of channels. 
print(channel_names)
print(len(channel_names), 'channels')

# %% [code]
# Define dictionaries to store the brain regions, sequences of fMRI scans, and FLOPs for each model. 
dict_brain_regions = dict()
dict_fmri_scans = dict()
dict_flops = dict()

# %% [code]
# Display the region-channel mapping. 
region_channel_mapping

# %% [markdown]
## **2. Gemma Chain-of-Thought**

We use the **Gemma-2-2B-IT large language model** with a Chain-of-Thought approach, and select its hyperparameters in order to ensure a relative variety of responses. We define two prompts, one to predict cognitive functions based on EEG data, and the other to predict fMRI data based on cognitive functions. We run the large language model on the eegfmriNF condition of all subjects, for a selection of brain regions spanning both a reasonable fraction of the brain and a variety of cognitive functions. In the bar plot presenting the results, an indicative baseline is given for visualization purposes, but the real baseline depends both on the selected fMRI scans and the distribution of missing or ambiguous predictions. 

# %% [code]
# Define the LLM to use. 
model_name = 'google/gemma-2-2b-it'

# Load the tokenizer and the model. 
tokenizer_gemma = AutoTokenizer.from_pretrained(model_name)
model_gemma = AutoModelForCausalLM.from_pretrained(model_name, torch_dtype = torch.float16, device_map = 'auto')

# %% [code]
# Define a function to create the first LLM prompt, from EEG to cognitive functions. 
def prompt_llm_eeg_to_cognition(X, fmri_scan_index, sequence_length, selected_bandpowers, selected_eeg_channel):

    # Retrieve the sequence of features. 
    X_sequence = X[(fmri_scan_index - sequence_length):fmri_scan_index, :]

    # Create the prompt. 
    bandpowers_results = ', '.join(
        f"{bandpower_names[bandpower_index]}: {X_sequence[:, bandpower_index]}" for bandpower_index in selected_bandpowers
    )
    prompt = (
        f"A human subject participated in a neuroscience experiment where EEG data and fMRI data were recorded simultaneously. "
        f"The EEG band powers were measured every two seconds at electrode {selected_eeg_channel}, "
        f"with the following results: {bandpowers_results}. "
        f"Given the EEG data, which cognitive functions is the subject likely engaging in? "
        f"Base your answer on your general knowledge in neuroscience and EEG research. "
        f"Please answer with only a list of cognitive functions. Just give your best prediction, without any explanation."
    )

    return prompt

# %% [code]
# Define a function to create the second LLM prompt, from cognitive functions to fMRI. 
def prompt_llm_cognition_to_fmri(cognitive_functions, selected_brain_region):

    # Create the prompt. 
    prompt = (
        f"A human subject participated in a neuroscience experiment where EEG data and fMRI data were recorded simultaneously. "
        f"This subject experienced the following cognitive functions: {cognitive_functions}. "
        f"The fMRI BOLD signal was measured in the {selected_brain_region}. "
        f"Given the cognitive functions, is the fMRI signal in this brain region likely increasing or decreasing? "
        f"Base your answer on your general knowledge in neuroscience and fMRI research. "
        f"Please answer with only one word: Increasing or Decreasing. Just give your best prediction, without any explanation."
    )

    return prompt

# %% [code]
# Define a function to run a LLM. 
def run_llm(prompt, tokenizer, model):

    # Run the LLM. 
    inputs = tokenizer(prompt, return_tensors = 'pt').to(model.device)
    output = model.generate(**inputs, 
                            max_new_tokens = 20, 
                            temperature = 1.2, 
                            top_p = 0.9, 
                            top_k = 50, 
                            do_sample = True, 
                            repetition_penalty = 1)

    # Decode the generated text, remove the original prompt, and return the answer. 
    generated_text = tokenizer.decode(output[0], skip_special_tokens = True)
    answer_only = generated_text[len(prompt):].strip()
    
    return answer_only

# %% [code]
# Define a function to run a LLM multiple times with a Chain-of-Thought approach. 
def run_llm_series(X, Y, fmri_scans, sequence_length, selected_bandpowers, selected_eeg_channel, selected_brain_region, tokenizer, model):

    # Create the NumPy arrays to store the true values and predictions.     
    Y_true = np.zeros(len(fmri_scans))
    Y_pred = np.zeros(len(fmri_scans))
    answers_cognitive_functions = []
    answers_fmri_predictions = []
    counter = 0

    # Iterate through all selected fMRI scans. 
    for fmri_scan_index in fmri_scans:

        # Create the first LLM prompt. 
        prompt_eeg_to_cognition = prompt_llm_eeg_to_cognition(X, 
                                                              fmri_scan_index = fmri_scan_index, 
                                                              sequence_length = sequence_length, 
                                                              selected_bandpowers = selected_bandpowers, 
                                                              selected_eeg_channel = selected_eeg_channel)
        
        # Run the LLM a first time, and store the result. 
        cognitive_functions = run_llm(prompt_eeg_to_cognition, tokenizer, model)
        answers_cognitive_functions.append(cognitive_functions) # For debugging purposes. 

        # Create the second LLM prompt. 
        prompt_cognition_to_fmri = prompt_llm_cognition_to_fmri(cognitive_functions, 
                                                                selected_brain_region = selected_brain_region)
        
        # Run the LLM a second time, and store the result. 
        fmri_prediction = run_llm(prompt_cognition_to_fmri, tokenizer, model)
        answers_fmri_predictions.append(fmri_prediction) # For debugging purposes. 

        # Analyze the answer. 
        increase_detected = 'increas' in fmri_prediction.lower()
        decrease_detected = 'decreas' in fmri_prediction.lower()
        if increase_detected and not decrease_detected:
            Y_pred[counter] = 1
        elif decrease_detected and not increase_detected:
            Y_pred[counter] = 0
        else:
            Y_pred[counter] = -1

        # Store the true value.
        Y_true[counter] = Y[fmri_scan_index]

        # Increment. 
        counter += 1

    return Y_true, Y_pred

# %% [code]
# Select a subject and a condition. 
subject = 'sub-xp101'
condition = 'eegfmriNF'

# Select a fMRI brain region and retrieve its corresponding EEG channel. 
selected_brain_region = 'Frontal Pole'
selected_brain_region_index = list(targets).index(selected_brain_region)
selected_eeg_channel = region_channel_mapping[selected_brain_region]
selected_eeg_channel_index = channel_names.index(selected_eeg_channel)

# Display the fMRI brain region and its corresponding EEG channel. 
print('Brain region:', selected_brain_region)
print('Brain region index:', int(selected_brain_region_index))
print('EEG channel:', selected_eeg_channel)
print('EEG channel index:', int(selected_eeg_channel_index))

# %% [code]
# Retrieve the features for the selected subject, condition, and EEG channel. 
X = dict_features_basis_class[subject][condition][:, (selected_eeg_channel_index * len(bandpower_names)):(selected_eeg_channel_index * len(bandpower_names)) + len(bandpower_names)]

# Retrieve the targets for the selected subject, condition, and fMRI brain region. 
Y = dict_targets_basis_class[subject][condition][:, selected_brain_region_index]

# Select the sequence length and the bandpowers. 
sequence_length = 5
selected_bandpowers = [0, 1, 2, 4]

# %% [code]
# Create the first LLM prompt. 
prompt_eeg_to_cognition = prompt_llm_eeg_to_cognition(X, 
                                                      fmri_scan_index = 5, 
                                                      sequence_length = sequence_length, 
                                                      selected_bandpowers = selected_bandpowers, 
                                                      selected_eeg_channel = selected_eeg_channel)
prompt_eeg_to_cognition

# %% [code]
# Run the LLM, and display the result. 
cognitive_functions = run_llm(prompt_eeg_to_cognition, tokenizer_gemma, model_gemma)
cognitive_functions

# %% [code]
# Create the second LLM prompt. 
prompt_cognition_to_fmri = prompt_llm_cognition_to_fmri(cognitive_functions, 
                                                        selected_brain_region = selected_brain_region)
prompt_cognition_to_fmri

# %% [code]
# Run the LLM, and display the result. 
fmri_prediction = run_llm(prompt_cognition_to_fmri, tokenizer_gemma, model_gemma)
fmri_prediction

# %% [code]
# Run the LLM multiple times. 
fmri_scans = range(5, (nb_scans_total - 1), 10)
Y_true, Y_pred = run_llm_series(X, Y, fmri_scans, sequence_length, selected_bandpowers, selected_eeg_channel, selected_brain_region, tokenizer_gemma, model_gemma)

# Compute the average accuracy. 
accuracy_array = Y_pred[Y_pred != -1] == Y_true[Y_pred != -1]
accuracy_sequence = np.sum(accuracy_array) / accuracy_array.shape[0]
accuracy_sequence

# %% [code]
# Select a condition and a list of brain regions. 
condition = 'eegfmriNF'
brain_regions = ['Frontal Pole', # High-order cognitive functions, but sometimes noisy EEG signal. 
                 'Precentral Gyrus', # Primary motor cortex. 
                 'Superior Temporal Gyrus, posterior division', # Auditory functions. 
                 'Superior Parietal Lobule', # Attention functions. 
                 'Occipital Pole'] # Primary visual cortex. 

# Select a sequence of fMRI scans. 
fmri_scans = range(5, (nb_scans_total - 1), 10) # Extensive search.
#fmri_scans = range(5, (nb_scans_total - 1), 100) # Quick search for debugging.

# Store the brain regions and the sequence of fMRI scans in dictionaries. 
dict_brain_regions['Large Language Model Chain-of-Thought'] = brain_regions
dict_fmri_scans['Large Language Model Chain-of-Thought'] = fmri_scans

# Define a dictionary to store the predictions and a Pandas DataFrame to store the accuracy. 
dict_predictions_gemma_CoT = dict()
accuracy_summary_gemma_CoT = pd.DataFrame(columns = brain_regions)

# Iterate through all brain regions. 
for selected_brain_region in brain_regions:

    # Display the current brain region. 
    print(selected_brain_region)

    # Retrieve the EEG channel corresponding to the brain region. 
    selected_brain_region_index = list(targets).index(selected_brain_region)
    selected_eeg_channel = region_channel_mapping[selected_brain_region]
    selected_eeg_channel_index = channel_names.index(selected_eeg_channel)

    # Define a dictionary to store the predictions for this brain region. 
    dict_predictions_gemma_CoT[selected_brain_region] = dict()

    # Iterate through all subjects. 
    for subject in subjects:
        
        # Retrieve the features for the selected subject, condition, and EEG channel. 
        X = dict_features_basis_class[subject][condition][:, (selected_eeg_channel_index * len(bandpower_names)):(selected_eeg_channel_index * len(bandpower_names)) + len(bandpower_names)]

        # Retrieve the targets for the selected subject, condition, and fMRI brain region. 
        Y = dict_targets_basis_class[subject][condition][:, selected_brain_region_index]

        # Run the LLM multiple times. 
        Y_true, Y_pred = run_llm_series(X, Y, fmri_scans, sequence_length, selected_bandpowers, selected_eeg_channel, selected_brain_region, tokenizer_gemma, model_gemma)

        # Store the predictions. 
        dict_predictions_gemma_CoT[selected_brain_region][subject] = Y_pred
        
        # Compute and store the average accuracy. 
        accuracy_array = Y_pred[Y_pred != -1] == Y_true[Y_pred != -1]
        accuracy_sequence = np.sum(accuracy_array) / accuracy_array.shape[0]
        accuracy_summary_gemma_CoT.loc[subject, selected_brain_region] = accuracy_sequence

        # Display the number of ambiguous predictions. 
        print(subject + ': ' + str(np.sum(Y_pred == -1)), 'ambiguous prediction(s) out of', len(Y_pred), '-', str(np.sum(Y_pred == -1) / len(Y_pred) * 100), '%')

# %% [code]
# Set the logging level to suppress warnings. 
logging.getLogger('fvcore.nn.jit_handles').setLevel(logging.ERROR)
logging.getLogger('fvcore.nn.jit_analysis').setLevel(logging.ERROR)

# Store current device. 
device_before = next(model_gemma.parameters()).device
print('Original device of the model:', device_before)

# Define a class wrapping the model. 
# We output only logits (Tensor data type) for compatibility with FlopCountAnalysis. 
class WrappedModel(torch.nn.Module):
    def __init__(self, model):
        super().__init__()
        self.model = model
    def forward(self, input_ids):
        output = self.model(input_ids)
        return output.logits

# Move the model to CPU. 
model_gemma_cpu = WrappedModel(model_gemma).to('cpu')

# Prepare dummy input for FLOPs analysis. 
print('==================')
print('STEP 1: EEG to Cognitive Functions')
dummy_input = (torch.ones((1, len(prompt_eeg_to_cognition)), dtype = torch.long),) # batch_size = 1, seq_len = prompt length
print('Prompt length:', len(prompt_eeg_to_cognition))

# Measure the FLOPs. 
flops_eeg_to_cognition = FlopCountAnalysis(model_gemma_cpu, dummy_input)
print(f'Total FLOPs: {flops_eeg_to_cognition.total():,}')
print('FLOPs by operator:', flops_eeg_to_cognition.by_operator())

# Prepare dummy input for FLOPs analysis. 
print('==================')
print('STEP 2: Cognitive Functions to fMRI')
dummy_input = (torch.ones((1, len(prompt_cognition_to_fmri)), dtype = torch.long),) # batch_size = 1, seq_len = prompt length
print('Prompt length:', len(prompt_cognition_to_fmri))

# Measure the FLOPs. 
flops_cognition_to_fmri = FlopCountAnalysis(model_gemma_cpu, dummy_input)
print(f'Total FLOPs: {flops_cognition_to_fmri.total():,}')
print('FLOPs by operator:', flops_cognition_to_fmri.by_operator())

# Move the model back to its original device. 
model_gemma.to(device_before)
print('Model moved back to:', device_before)

# %% [code]
# Store and display the number of FLOPs. 
dict_flops['Large Language Model Chain-of-Thought: EEG to Cognitive Functions'] = flops_eeg_to_cognition.total()
dict_flops['Large Language Model Chain-of-Thought: Cognitive Functions to fMRI'] = flops_cognition_to_fmri.total()
print('FLOPs per inference, EEG to cognitive functions:', flops_eeg_to_cognition.total())
print('FLOPs per inference, cognitive functions to fMRI:', flops_cognition_to_fmri.total())

# %% [code]
# Display the accuracies per subject and brain region. 
accuracy_summary_gemma_CoT

# %% [code]
# Retrieve the average accuracy across subjects, and the standard error of the accuracy across subjects. 
accuracy_mean = list(accuracy_summary_gemma_CoT.mean().values)
accuracy_mean.insert(0, 0.5) # Insert a baseline average of 0.5. 
accuracy_se = list(accuracy_summary_gemma_CoT.std().values / np.sqrt(len(subjects)))
accuracy_se.insert(0, 0) # Insert a baseline standard error of 0. 

# Append the total average accuracy and its standard error. 
nb_brain_regions = accuracy_summary_gemma_CoT.shape[1]
accuracy_mean.append(np.mean(accuracy_mean[1:]))
accuracy_se.append(np.std(accuracy_mean[1:]) / (np.sqrt(len(subjects) * nb_brain_regions)))

# Create the labels. 
labels = brain_regions.copy()
labels.insert(0, 'Baseline')
labels.append('Total')

# Display the average accuracy and its standard error in a bar plot. 
fig, axes = plt.subplots(nrows = 1, ncols = 1, figsize = (10, 4))
axes.bar(range(nb_brain_regions + 2), accuracy_mean, width = 0.6, color = ['green', *['blue'] * nb_brain_regions, 'mediumblue'])
axes.errorbar(range(nb_brain_regions + 2), accuracy_mean, yerr = accuracy_se, fmt = 'None', color = 'black')
axes.axhline(accuracy_mean[0], 0, 1, color = 'green', linestyle = 'dashed', alpha = 0.5)
axes.set_xticks(range(nb_brain_regions + 2), labels = labels)
axes.set_xticklabels(labels, rotation = 45, ha = 'right')
axes.set_ylabel('Average accuracy across subjects')
axes.set_title('Gemma-2-2B-IT with Chain-of-Thought');

# %% [markdown]
## **3. Results**

# %% [code]
# Save the predictions into a Pickle file. 
with open(results_path + 'classification/dict_predictions_gemma_CoT.p', 'wb') as file:
    pickle.dump(dict_predictions_gemma_CoT, file)

# Save the summary into a Pickle file. 
with open(results_path + 'classification/df_summary_gemma_CoT.p', 'wb') as file:
    pickle.dump(accuracy_summary_gemma_CoT, file)

# Save the brain regions, sequences of fMRI scans, and FLOPs into Pickle files. 
with open(results_path + 'brain_regions_large_language_model_CoT.p', 'wb') as file:
    pickle.dump(dict_brain_regions, file)
with open(results_path + 'fmri_scans_large_language_model_CoT.p', 'wb') as file:
    pickle.dump(dict_fmri_scans, file)
with open(results_path + 'flops_large_language_model_CoT.p', 'wb') as file:
    pickle.dump(dict_flops, file)
```
