## Supplementary material for "Neural Networks and Foundation Models: Two Strategies for EEG-to-fMRI Prediction": 11_Large_Language_Model_Fine-Tuning.ipynb

```
# %% [markdown]
# **Neural Networks and Foundation Models: Two Strategies for EEG-to-fMRI Prediction**
### **Supplementary Material**

# %% [markdown]
This project includes fourteen Notebooks:

*01: fMRI Preprocessing*

*02: EEG Preprocessing*

*03: Classification Models*

*04: Regression Models*

*05: Neural Networks*

*06: Convolutional Neural Networks*

*07: Recurrent Neural Networks*

*08: Transformers*

*09: Large Language Models*

*10: Large Language Model Chain-of-Thought*

***11: Large Language Model Fine-Tuning***

*12: Large Multimodal Model*

*13: Statistical Tests*

*14: Tables and Figures*

# %% [markdown]
### Notebook 11

# **Large Language Model Fine-Tuning**

# %% [markdown]
## **1. Region-Channel Mapping**

We retrieve the **region-channel mapping**, associating each brain region to the EEG channel whose signal is most likely to provide relevant information regarding its fMRI activity. 

# %% [code]
import numpy as np
import pandas as pd
import matplotlib.pyplot as plt
%matplotlib inline
import pickle
import mne
import torch
from transformers import AutoModelForCausalLM, AutoTokenizer, TrainingArguments, DataCollatorForSeq2Seq, Trainer
import datasets
from peft import LoraConfig, get_peft_model, PeftModel
from fvcore.nn import FlopCountAnalysis
import logging
from copy import deepcopy
import gc

# %% [code]
# Define the raw data path. 
raw_data_path = '../ds002336-download/'

# Define the preprocessed data path. 
preprocessed_data_path = '../preprocessed_data/'

# Define the fine-tuned models path. 
finetuned_models_path = '../fine-tuned_models/'

# Define the results path. 
results_path = '../results/'

# %% [code]
# Load the features and the targets for the classification task. 
with open(preprocessed_data_path + 'dict_targets.p', 'rb') as file:
    dict_targets = pickle.load(file)
with open(preprocessed_data_path + 'classification/dict_features_classification_basis.p', 'rb') as file:
    dict_features_basis_class = pickle.load(file)
with open(preprocessed_data_path + 'classification/dict_targets_classification_basis.p', 'rb') as file:
    dict_targets_basis_class = pickle.load(file)

# Load the region-channel mapping. 
with open(results_path + 'region_channel_mapping.p', 'rb') as file:
    region_channel_mapping = pickle.load(file)

# %% [code]
# Define the list of subjects, removing sub-xp102 who has a missing condition. 
subjects = ['sub-xp1' + str(x).zfill(2) for x in range(1, 11)]
subjects.remove('sub-xp102')
subject = subjects[0]

# Retrieve and display the total number of scans. 
nb_scans_total = dict_targets[subject]['eegfmriNF']['Frontal Pole'].shape[0]
nb_scans_total

# %% [code]
# Retrieve the list of all targets. 
targets = list(dict_targets[subject]['eegfmriNF'].keys())

# Display the brain region names and the number of brain regions. 
print(targets)
print(len(targets), 'brain regions')

# %% [code]
# Define the bandpower names. 
bandpower_names = ['Delta (1-4 Hz)', 'Theta (4-8 Hz)', 'Alpha (8-12 Hz)', 'Sigma (12-16 Hz)', 'Beta (16-30 Hz)', 'Gamma (30-40 Hz)']

# Define the path of the header file (.vhdr) for this subject, for the eegfmriNF condition. 
vhdr_path = raw_data_path + 'derivatives/' + subject + '/eeg_pp/' + subject + '_task-eegfmriNF_eeg_pp.vhdr'

# Load the EEG data using the MNE library, keeping error messages but suppressing warnings. 
channels_data = mne.io.read_raw_brainvision(vhdr_path, misc = ['ECG'], preload = True, verbose = 'ERROR')

# Remove the ECG channel from the channel names. 
channel_names = channels_data.ch_names
channel_names.remove('ECG')

# Display the channel names and the number of channels. 
print(channel_names)
print(len(channel_names), 'channels')

# %% [code]
# Define dictionaries to store the brain regions, sequences of fMRI scans, and FLOPs for each model. 
dict_brain_regions = dict()
dict_fmri_scans = dict()
dict_flops = dict()

# %% [code]
# Display the region-channel mapping. 
region_channel_mapping

# %% [markdown]
## **2. Gemma**

We use the **Gemma-2-2B-IT large language model**, and select its hyperparameters in order to ensure a relative variety of responses. We define a prompt where we provide a sequence of EEG bandpower values, and ask the model to predict the corresponding fMRI activity. We run the large language model on the eegfmriNF condition of all subjects, for a selection of brain regions spanning both a reasonable fraction of the brain and a variety of cognitive functions. In the bar plot presenting the results, an indicative baseline is given for visualization purposes, but the real baseline depends both on the selected fMRI scans and the distribution of missing or ambiguous predictions. 

# %% [code]
# Define the LLM to use. 
model_name = 'google/gemma-2-2b-it'

# Load the tokenizer and the model. 
tokenizer_gemma = AutoTokenizer.from_pretrained(model_name)
model_gemma = AutoModelForCausalLM.from_pretrained(model_name, torch_dtype = torch.float16, device_map = 'auto', attn_implementation = 'eager')

# %% [code]
# Define a function to create a LLM prompt with a sequence of bandpower values. 
def prompt_llm(X, fmri_scan_index, sequence_length, selected_bandpowers, selected_eeg_channel, selected_brain_region):

    # Retrieve the sequence of features. 
    X_sequence = X[(fmri_scan_index - sequence_length):fmri_scan_index, :]

    # Create the prompt. 
    bandpowers_results = ', '.join(
        f"{bandpower_names[bandpower_index]}: {X_sequence[:, bandpower_index]}" for bandpower_index in selected_bandpowers
    )
    prompt = (
        f"A human subject participated in a neuroscience experiment where EEG data and fMRI data were recorded simultaneously. "
        f"The EEG band powers were measured every two seconds at electrode {selected_eeg_channel}, "
        f"with the following results: {bandpowers_results}. "
        f"The fMRI BOLD signal was measured in the {selected_brain_region} during the last two seconds. "
        f"Given the EEG data, is the fMRI signal in this brain region likely increasing or decreasing during these last two seconds? "
        f"Base your answer on your general knowledge in neuroscience, EEG research, and fMRI research. "
        f"Since this is a time series, you might need to take into account the hemodynamic response function (HRF), "
        f"and the fact that after an event, the fMRI response is delayed compared to the EEG response. "
        f"Please answer with only one word: Increasing or Decreasing. Just give your best prediction, without any explanation."
    )

    return prompt

# %% [code]
# Define a function to run a LLM. 
def run_llm(prompt, tokenizer, model):

    # Run the LLM. 
    inputs = tokenizer(prompt, return_tensors = 'pt').to(model.device)
    output = model.generate(**inputs, 
                            max_new_tokens = 20, 
                            temperature = 1.2, 
                            top_p = 0.9, 
                            top_k = 50, 
                            do_sample = True, 
                            repetition_penalty = 1)

    # Decode the generated text, remove the original prompt, and return the answer. 
    generated_text = tokenizer.decode(output[0], skip_special_tokens = True)
    answer_only = generated_text[len(prompt):].strip()
    
    return answer_only

# %% [code]
# Define a function to run a LLM multiple times. 
def run_llm_series(X, Y, fmri_scans, sequence_length, selected_bandpowers, selected_eeg_channel, selected_brain_region, tokenizer, model):

    # Create the NumPy arrays to store the true values and predictions.     
    Y_true = np.zeros(len(fmri_scans))
    Y_pred = np.zeros(len(fmri_scans))
    answers = []
    counter = 0

    # Iterate through all selected fMRI scans. 
    for fmri_scan_index in fmri_scans:

        # Create the LLM prompt. 
        prompt = prompt_llm(X, 
                            fmri_scan_index = fmri_scan_index, 
                            sequence_length = sequence_length, 
                            selected_bandpowers = selected_bandpowers, 
                            selected_eeg_channel = selected_eeg_channel, 
                            selected_brain_region = selected_brain_region)
        
        # Run the LLM and store the result. 
        answer_only = run_llm(prompt, tokenizer, model)
        answers.append(answer_only) # For debugging purposes. 

        # Analyze the answer. 
        increase_detected = 'increas' in answer_only.lower()
        decrease_detected = 'decreas' in answer_only.lower()
        if increase_detected and not decrease_detected:
            Y_pred[counter] = 1
        elif decrease_detected and not increase_detected:
            Y_pred[counter] = 0
        else:
            Y_pred[counter] = -1

        # Store the true value.
        Y_true[counter] = Y[fmri_scan_index]

        # Increment. 
        counter += 1

    return Y_true, Y_pred

# %% [code]
# Select a subject and a condition. 
subject = 'sub-xp101'
condition = 'eegfmriNF'

# Select a fMRI brain region and retrieve its corresponding EEG channel. 
selected_brain_region = 'Frontal Pole'
selected_brain_region_index = list(targets).index(selected_brain_region)
selected_eeg_channel = region_channel_mapping[selected_brain_region]
selected_eeg_channel_index = channel_names.index(selected_eeg_channel)

# Display the fMRI brain region and its corresponding EEG channel. 
print('Brain region:', selected_brain_region)
print('Brain region index:', int(selected_brain_region_index))
print('EEG channel:', selected_eeg_channel)
print('EEG channel index:', int(selected_eeg_channel_index))

# %% [code]
# Retrieve the features for the selected subject, condition, and EEG channel. 
X = dict_features_basis_class[subject][condition][:, (selected_eeg_channel_index * len(bandpower_names)):(selected_eeg_channel_index * len(bandpower_names)) + len(bandpower_names)]

# Retrieve the targets for the selected subject, condition, and fMRI brain region. 
Y = dict_targets_basis_class[subject][condition][:, selected_brain_region_index]

# Select the sequence length and the bandpowers. 
sequence_length = 5
selected_bandpowers = [0, 1, 2, 4]

# Create the LLM prompt. 
prompt = prompt_llm(X, 
                    fmri_scan_index = 5, 
                    sequence_length = sequence_length, 
                    selected_bandpowers = selected_bandpowers, 
                    selected_eeg_channel = selected_eeg_channel, 
                    selected_brain_region = selected_brain_region)
prompt

# %% [code]
# Run the LLM once, and display the result. 
answer_only = run_llm(prompt, tokenizer_gemma, model_gemma)
answer_only

# %% [code]
# Select the sequence length, bandpowers, condition, and brain regions. 
sequence_length = 5
selected_bandpowers = [0, 1, 2, 4]
condition = 'eegfmriNF'
brain_regions = ['Frontal Pole', # High-order cognitive functions, but sometimes noisy EEG signal. 
                 'Precentral Gyrus', # Primary motor cortex. 
                 'Superior Temporal Gyrus, posterior division', # Auditory functions. 
                 'Superior Parietal Lobule', # Attention functions. 
                 'Occipital Pole'] # Primary visual cortex. 

# Select a sequence of fMRI scans. 
fmri_scans = range(5, (nb_scans_total - 1), 10) # Extensive search.
#fmri_scans = range(5, (nb_scans_total - 1), 100) # Quick search for debugging.

# Store the brain regions and the sequence of fMRI scans in dictionaries. 
dict_brain_regions['Large Language Model Fine-Tuning: Without Fine-Tuning'] = brain_regions
dict_fmri_scans['Large Language Model Fine-Tuning: Without Fine-Tuning'] = fmri_scans

# Define a dictionary to store the predictions and a Pandas DataFrame to store the accuracy. 
dict_predictions_gemma_no_finetuning = dict()
accuracy_summary_gemma_no_finetuning = pd.DataFrame(columns = brain_regions)

# Iterate through all brain regions. 
for selected_brain_region in brain_regions:

    # Display the current brain region. 
    print(selected_brain_region)

    # Retrieve the EEG channel corresponding to the brain region. 
    selected_brain_region_index = list(targets).index(selected_brain_region)
    selected_eeg_channel = region_channel_mapping[selected_brain_region]
    selected_eeg_channel_index = channel_names.index(selected_eeg_channel)

    # Define a dictionary to store the predictions for this brain region. 
    dict_predictions_gemma_no_finetuning[selected_brain_region] = dict()

    # Iterate through all subjects. 
    for subject in subjects:
        
        # Retrieve the features for the selected subject, condition, and EEG channel. 
        X = dict_features_basis_class[subject][condition][:, (selected_eeg_channel_index * len(bandpower_names)):(selected_eeg_channel_index * len(bandpower_names)) + len(bandpower_names)]

        # Retrieve the targets for the selected subject, condition, and fMRI brain region. 
        Y = dict_targets_basis_class[subject][condition][:, selected_brain_region_index]

        # Run the LLM multiple times. 
        Y_true, Y_pred = run_llm_series(X, Y, fmri_scans, sequence_length, selected_bandpowers, selected_eeg_channel, selected_brain_region, tokenizer_gemma, model_gemma)

        # Store the predictions. 
        dict_predictions_gemma_no_finetuning[selected_brain_region][subject] = Y_pred
        
        # Compute and store the average accuracy. 
        accuracy_array = Y_pred[Y_pred != -1] == Y_true[Y_pred != -1]
        accuracy_sequence = np.sum(accuracy_array) / accuracy_array.shape[0]
        accuracy_summary_gemma_no_finetuning.loc[subject, selected_brain_region] = accuracy_sequence

        # Display the number of ambiguous predictions. 
        print(subject + ': ' + str(np.sum(Y_pred == -1)), 'ambiguous prediction(s) out of', len(Y_pred), '-', str(np.sum(Y_pred == -1) / len(Y_pred) * 100), '%')

# %% [code]
# Set the logging level to suppress warnings. 
logging.getLogger('fvcore.nn.jit_handles').setLevel(logging.ERROR)
logging.getLogger('fvcore.nn.jit_analysis').setLevel(logging.ERROR)

# Store current device. 
device_before = next(model_gemma.parameters()).device
print('Original device of the model:', device_before)

# Define a class wrapping the model. 
# We output only logits (Tensor data type) for compatibility with FlopCountAnalysis. 
class WrappedModel(torch.nn.Module):
    def __init__(self, model):
        super().__init__()
        self.model = model
    def forward(self, input_ids):
        output = self.model(input_ids)
        return output.logits

# Move the model to CPU and prepare dummy input for FLOPs analysis. 
model_gemma_cpu = WrappedModel(model_gemma).to('cpu')
dummy_input = (torch.ones((1, len(prompt)), dtype = torch.long),) # batch_size = 1, seq_len = prompt length
print('Prompt length:', len(prompt))

# Measure the FLOPs. 
flops = FlopCountAnalysis(model_gemma_cpu, dummy_input)
print(f'Total FLOPs: {flops.total():,}')
print('FLOPs by operator:', flops.by_operator())

# Move the model back to its original device. 
model_gemma.to(device_before)
print('Model moved back to:', device_before)

# %% [code]
# Store and display the number of FLOPs. 
dict_flops['Large Language Model Fine-Tuning: Without Fine-Tuning'] = flops.total()
print('FLOPs per inference:', flops.total())

# %% [code]
# Display the accuracies per subject and brain region. 
accuracy_summary_gemma_no_finetuning

# %% [code]
# Retrieve the average accuracy across subjects, and the standard error of the accuracy across subjects. 
accuracy_mean = list(accuracy_summary_gemma_no_finetuning.mean().values)
accuracy_mean.insert(0, 0.5) # Insert a baseline average of 0.5. 
accuracy_se = list(accuracy_summary_gemma_no_finetuning.std().values / np.sqrt(len(subjects)))
accuracy_se.insert(0, 0) # Insert a baseline standard error of 0. 

# Append the total average accuracy and its standard error. 
nb_brain_regions = accuracy_summary_gemma_no_finetuning.shape[1]
accuracy_mean.append(np.mean(accuracy_mean[1:]))
accuracy_se.append(np.std(accuracy_mean[1:]) / (np.sqrt(len(subjects) * nb_brain_regions)))

# Create the labels. 
labels = brain_regions.copy()
labels.insert(0, 'Baseline')
labels.append('Total')

# Display the average accuracy and its standard error in a bar plot. 
fig, axes = plt.subplots(nrows = 1, ncols = 1, figsize = (10, 4))
axes.bar(range(nb_brain_regions + 2), accuracy_mean, width = 0.6, color = ['green', *['blue'] * nb_brain_regions, 'mediumblue'])
axes.errorbar(range(nb_brain_regions + 2), accuracy_mean, yerr = accuracy_se, fmt = 'None', color = 'black')
axes.axhline(accuracy_mean[0], 0, 1, color = 'green', linestyle = 'dashed', alpha = 0.5)
axes.set_xticks(range(nb_brain_regions + 2), labels = labels)
axes.set_xticklabels(labels, rotation = 45, ha = 'right')
axes.set_ylabel('Average accuracy across subjects')
axes.set_title('Gemma-2-2B-IT without fine-tuning');

# %% [markdown]
## **3. Gemma Fine-Tuning**

We **fine-tune** a separate instance of the **Gemma-2-2B-IT large language model** for each brain region of each subject, using input-output pairs obtained from the fmriNF session. This strategy turns out to give better results than fine-tuning either a single instance for all subjects and brain regions, or one instance per subject. However, since we are working with very few examples and binary targets ("Increasing" or "Decreasing"), the gradient updates can be very noisy. In this situation, the model can behave unexpectedly, producing empty outputs or oscillating between the two targets. We use gradient accumulation to ensure a more stable convergence, and a low learning rate to avoid catastrophic forgetting. Once the model is fine-tuned, we generate predictions for the eegfmriNF condition, and compare them with the predictions of the original model. In the bar plot presenting the results, an indicative baseline is given for visualization purposes, but the real baseline depends both on the selected fMRI scans and the distribution of missing or ambiguous predictions. 

# %% [code]
# Define the tokenize function. 
def tokenize_function(example):

    # Tokenize inputs and labels. 
    input_encodings = tokenizer_gemma(example['input'], truncation = True, padding = 'max_length', max_length = 512)
    label_encodings = tokenizer_gemma(example['output'], add_special_tokens = False, truncation = True, padding = 'max_length', max_length = 512)
    
    # Replace pad tokens with -100 in order to ignore them in the loss calculation, and remove the BOS token. 
    labels = label_encodings['input_ids']
    labels = [label if label != tokenizer_gemma.pad_token_id else -100 for label in labels]
    
    # Return formatted data. 
    return {
        'input_ids': input_encodings['input_ids'],
        'attention_mask': input_encodings['attention_mask'],
        'labels': labels
    }

# %% [code]
# Define a function to replace pad tokens with ignore tokens. 
def replace_pad_with_ignore(labels, pad_token_id, ignore_index = -100):
    return [ignore_index if token == pad_token_id else token for token in labels]

# %% [code]
# Select the sequence length, bandpowers, condition, and brain regions. 
sequence_length = 5
selected_bandpowers = [0, 1, 2, 4]
condition = 'fmriNF'
brain_regions = ['Frontal Pole', # High-order cognitive functions, but sometimes noisy EEG signal. 
                 'Precentral Gyrus', # Primary motor cortex. 
                 'Superior Temporal Gyrus, posterior division', # Auditory functions. 
                 'Superior Parietal Lobule', # Attention functions. 
                 'Occipital Pole'] # Primary visual cortex. 

# Select a sequence of fMRI scans. 
fmri_scans = range(5, (nb_scans_total - 1), 20) # Extensive search. Do not select more scans than this to avoid overfitting issues. 
#fmri_scans = range(5, (nb_scans_total - 1), 100) # Quick search for debugging.

# Define a dictionary to store the fine-tuned models. Warning: this dictionary is very large and requires a lot of memory. 
#finetuned_models = dict() # For debugging. 

# Iterate through all brain regions. 
for selected_brain_region in brain_regions:

    # Display the current brain region. 
    print(selected_brain_region)

    # Retrieve the EEG channel corresponding to the brain region. 
    selected_brain_region_index = list(targets).index(selected_brain_region)
    selected_eeg_channel = region_channel_mapping[selected_brain_region]
    selected_eeg_channel_index = channel_names.index(selected_eeg_channel)

    # Define a dictionary at the level of brain regions. Warning: this dictionary is very large and requires a lot of memory. 
    #finetuned_models[selected_brain_region] = dict() # For debugging. 

    # Iterate through all subjects. 
    for subject in subjects:

        # Display the current subject. 
        print(subject)

        # Define the train data list for fine-tuning. 
        train_data = []

        # PROMPT GENERATION ==========================================================================================
                                
        # Retrieve the features for the selected subject, condition, and EEG channel. 
        X = dict_features_basis_class[subject][condition][:, (selected_eeg_channel_index * len(bandpower_names)):(selected_eeg_channel_index * len(bandpower_names)) + len(bandpower_names)]

        # Retrieve the targets for the selected subject, condition, and fMRI brain region. 
        Y = dict_targets_basis_class[subject][condition][:, selected_brain_region_index]

        # Iterate through all selected fMRI scans. 
        for fmri_scan_index in fmri_scans:

            # Create the LLM prompt. 
            prompt = prompt_llm(X, 
                                fmri_scan_index = fmri_scan_index, 
                                sequence_length = sequence_length, 
                                selected_bandpowers = selected_bandpowers, 
                                selected_eeg_channel = selected_eeg_channel, 
                                selected_brain_region = selected_brain_region)
            
            # Retrieve the correct answer. 
            if Y[fmri_scan_index] == 1:
                answer = 'increasing'
            else:
                answer = 'decreasing'

            # Append the prompt and the answer to the train data list. 
            train_data.append({'input': prompt, 'output': answer})

        # FINE-TUNING =================================================================================================
        
        # Tokenize the dataset, and replace pad tokens by ignore tokens. 
        datasets.disable_progress_bar()
        dataset = datasets.Dataset.from_list(train_data)
        dataset = dataset.map(tokenize_function, batched = True)
        dataset = dataset.map(lambda example: {'labels': replace_pad_with_ignore(example['labels'], pad_token_id = 0)})

        # Define the LoRA configuration. 
        lora_config = LoraConfig(
            r = 16, lora_alpha = 32, target_modules = ['q_proj', 'v_proj'], lora_dropout = 0.1, bias = 'none', task_type = 'CAUSAL_LM'
        )

        # Create a new instance of the model with the LoRA configuration. 
        model_copy = deepcopy(model_gemma)
        model_gemma_finetuned = get_peft_model(model_copy, lora_config)

        # Define the training arguments. 
        training_args = TrainingArguments(
            output_dir = finetuned_models_path, 
            per_device_train_batch_size = 1, 
            gradient_accumulation_steps = 8, 
            learning_rate = 1e-5, 
            weight_decay = 0.01, 
            num_train_epochs = 5, 
            save_strategy = 'no', 
            logging_strategy = 'no', 
            disable_tqdm = True, 
            label_names = ['labels']
        )

        # Define the data collator. 
        data_collator = DataCollatorForSeq2Seq(tokenizer_gemma, padding = True, label_pad_token_id = -100)

        # Define the trainer. 
        trainer = Trainer(
            model = model_gemma_finetuned, 
            args = training_args, 
            train_dataset = dataset, 
            data_collator = data_collator
        )

        # Fine-tune the model. 
        trainer.train()

        # Define the fine-tuned model path and save the model. 
        finetuned_model_path = finetuned_models_path + 'gemma-2-2b-it_' + subject + '_' + selected_brain_region.replace(' ', '_')
        trainer.save_model(finetuned_model_path)

        # Store the fine-tuned model in the dictionary. Warning: this dictionary is very large and requires a lot of memory. 
        #finetuned_models[selected_brain_region][subject] = model_gemma_finetuned # For debugging. 

        # Delete the model and run garbage collection. 
        del model_copy
        del model_gemma_finetuned
        gc.collect()

# %% [code]
# Display the Dataset object. 
dataset

# %% [code]
# Display an example of input and output. 
print('Input:', dataset['input'][0])
print('Output:', dataset['output'][0])

# %% [code]
# Display the first and last input tokens. 
print('The first input tokens should be zero given the left padding. The last input tokens should be non-zero.')
print('\nFirst input tokens:', dataset['input_ids'][0][0:10])
print('Last input tokens:', dataset['input_ids'][0][-10:])

# %% [code]
# Plot the attention mask values. 
print('The attention mask values should be zero for the padding tokens and one for the non-padding tokens.')
plt.figure(figsize=(5, 3))
plt.plot(dataset['attention_mask'][0]);

# %% [code]
# Display the label values. 
print('The label values should be -100 (the ignore token) for all positions except the last one.')
pd.Series(dataset['labels'][0]).value_counts()

# %% [code]
# Display the last label value. 
last_label = dataset['labels'][0][-1]
print('Last label:', last_label)
print('Last label decoded:', tokenizer_gemma.decode([last_label]))

# %% [code]
# Select a subject and a condition. 
subject = 'sub-xp101'
condition = 'eegfmriNF'

# Select a fMRI brain region and retrieve its corresponding EEG channel. 
selected_brain_region = 'Frontal Pole'
selected_brain_region_index = list(targets).index(selected_brain_region)
selected_eeg_channel = region_channel_mapping[selected_brain_region]
selected_eeg_channel_index = channel_names.index(selected_eeg_channel)

# Display the fMRI brain region and its corresponding EEG channel. 
print('Brain region:', selected_brain_region)
print('Brain region index:', int(selected_brain_region_index))
print('EEG channel:', selected_eeg_channel)
print('EEG channel index:', int(selected_eeg_channel_index))

# %% [code]
# Retrieve the features for the selected subject, condition, and EEG channel. 
X = dict_features_basis_class[subject][condition][:, (selected_eeg_channel_index * len(bandpower_names)):(selected_eeg_channel_index * len(bandpower_names)) + len(bandpower_names)]

# Retrieve the targets for the selected subject, condition, and fMRI brain region. 
Y = dict_targets_basis_class[subject][condition][:, selected_brain_region_index]

# Select the sequence length and the bandpowers. 
sequence_length = 5
selected_bandpowers = [0, 1, 2, 4]

# Create the LLM prompt. 
prompt = prompt_llm(X, 
                    fmri_scan_index = 5, 
                    sequence_length = sequence_length, 
                    selected_bandpowers = selected_bandpowers, 
                    selected_eeg_channel = selected_eeg_channel, 
                    selected_brain_region = selected_brain_region)
prompt

# %% [code]
# Define the fine-tuned model path. 
finetuned_model_path = finetuned_models_path + 'gemma-2-2b-it_' + subject + '_' + selected_brain_region.replace(' ', '_')

# Copy the base model. 
model_copy = deepcopy(model_gemma)

# Load the fine-tuned model. 
model_gemma_finetuned = PeftModel.from_pretrained(model_copy, finetuned_model_path)

# Run the LLM once. 
answer_only = run_llm(prompt, tokenizer_gemma, model_gemma_finetuned)

# Delete the model and run garbage collection. 
del model_copy
del model_gemma_finetuned
gc.collect()

# Display the result. 
answer_only

# %% [code]
# Select the sequence length, bandpowers, condition, and brain regions. 
sequence_length = 5
selected_bandpowers = [0, 1, 2, 4]
condition = 'eegfmriNF'
brain_regions = ['Frontal Pole', # High-order cognitive functions, but sometimes noisy EEG signal. 
                 'Precentral Gyrus', # Primary motor cortex. 
                 'Superior Temporal Gyrus, posterior division', # Auditory functions. 
                 'Superior Parietal Lobule', # Attention functions. 
                 'Occipital Pole'] # Primary visual cortex. 

# Select a sequence of fMRI scans. 
fmri_scans = range(5, (nb_scans_total - 1), 10) # Extensive search.
#fmri_scans = range(5, (nb_scans_total - 1), 100) # Quick search for debugging.

# Store the brain regions and the sequence of fMRI scans in dictionaries. 
dict_brain_regions['Large Language Model Fine-Tuning: With Fine-Tuning'] = brain_regions
dict_fmri_scans['Large Language Model Fine-Tuning: With Fine-Tuning'] = fmri_scans

# Define a dictionary to store the predictions and a Pandas DataFrame to store the accuracy. 
dict_predictions_gemma_finetuning = dict()
accuracy_summary_gemma_finetuning = pd.DataFrame(columns = brain_regions)

# Iterate through all brain regions. 
for selected_brain_region in brain_regions:

    # Display the current brain region. 
    print(selected_brain_region)

    # Retrieve the EEG channel corresponding to the brain region. 
    selected_brain_region_index = list(targets).index(selected_brain_region)
    selected_eeg_channel = region_channel_mapping[selected_brain_region]
    selected_eeg_channel_index = channel_names.index(selected_eeg_channel)

    # Define a dictionary to store the predictions for this brain region. 
    dict_predictions_gemma_finetuning[selected_brain_region] = dict()

    # Iterate through all subjects. 
    for subject in subjects:

        # Retrieve the features for the selected subject, condition, and EEG channel. 
        X = dict_features_basis_class[subject][condition][:, (selected_eeg_channel_index * len(bandpower_names)):(selected_eeg_channel_index * len(bandpower_names)) + len(bandpower_names)]

        # Retrieve the targets for the selected subject, condition, and fMRI brain region. 
        Y = dict_targets_basis_class[subject][condition][:, selected_brain_region_index]

        # Define the fine-tuned model path. 
        finetuned_model_path = finetuned_models_path + 'gemma-2-2b-it_' + subject + '_' + selected_brain_region.replace(' ', '_')

        # Copy the base model. 
        model_copy = deepcopy(model_gemma)

        # Load the fine-tuned model. 
        model_gemma_finetuned = PeftModel.from_pretrained(model_copy, finetuned_model_path)

        # Run the LLM multiple times. 
        #model_gemma_finetuned = finetuned_models[selected_brain_region][subject] # For debugging. 
        Y_true, Y_pred = run_llm_series(X, Y, fmri_scans, sequence_length, selected_bandpowers, selected_eeg_channel, selected_brain_region, tokenizer_gemma, model_gemma_finetuned)

        # Store the predictions. 
        dict_predictions_gemma_finetuning[selected_brain_region][subject] = Y_pred
        
        # Compute and store the average accuracy. 
        accuracy_array = Y_pred[Y_pred != -1] == Y_true[Y_pred != -1]
        accuracy_sequence = np.sum(accuracy_array) / accuracy_array.shape[0]
        accuracy_summary_gemma_finetuning.loc[subject, selected_brain_region] = accuracy_sequence

        # Display the number of ambiguous predictions. 
        print(subject + ': ' + str(np.sum(Y_pred == -1)), 'ambiguous prediction(s) out of', len(Y_pred), '-', str(np.sum(Y_pred == -1) / len(Y_pred) * 100), '%')

        # Delete the model and run garbage collection. 
        del model_copy
        del model_gemma_finetuned
        gc.collect()

# %% [code]
# Store current device. 
device_before = next(model_gemma.parameters()).device
print('Original device of the model:', device_before)

# Move the model to CPU and prepare dummy input for FLOPs analysis. 
model_gemma_cpu = WrappedModel(model_gemma).to('cpu')
dummy_input = (torch.ones((1, len(prompt)), dtype = torch.long),) # batch_size = 1, seq_len = prompt length
print('Prompt length:', len(prompt))

# Measure the FLOPs. 
flops = FlopCountAnalysis(model_gemma_cpu, dummy_input)
print(f'Total FLOPs: {flops.total():,}')
print('FLOPs by operator:', flops.by_operator())

# Move the model back to its original device. 
model_gemma.to(device_before)
print('Model moved back to:', device_before)

# %% [code]
# Store and display the number of FLOPs. 
dict_flops['Large Language Model Fine-Tuning: With Fine-Tuning'] = flops.total()
print('FLOPs per inference:', flops.total())

# %% [code]
# Display the accuracies per subject and brain region. 
accuracy_summary_gemma_finetuning

# %% [code]
# Retrieve the average accuracy across subjects, and the standard error of the accuracy across subjects. 
accuracy_mean = list(accuracy_summary_gemma_finetuning.mean().values)
accuracy_mean.insert(0, 0.5) # Insert a baseline average of 0.5. 
accuracy_se = list(accuracy_summary_gemma_finetuning.std().values / np.sqrt(len(subjects)))
accuracy_se.insert(0, 0) # Insert a baseline standard error of 0. 

# Append the total average accuracy and its standard error. 
nb_brain_regions = accuracy_summary_gemma_finetuning.shape[1]
accuracy_mean.append(np.mean(accuracy_mean[1:]))
accuracy_se.append(np.std(accuracy_mean[1:]) / (np.sqrt(len(subjects) * nb_brain_regions)))

# Create the labels. 
labels = brain_regions.copy()
labels.insert(0, 'Baseline')
labels.append('Total')

# Display the average accuracy and its standard error in a bar plot. 
fig, axes = plt.subplots(nrows = 1, ncols = 1, figsize = (10, 4))
axes.bar(range(nb_brain_regions + 2), accuracy_mean, width = 0.6, color = ['green', *['blue'] * nb_brain_regions, 'mediumblue'])
axes.errorbar(range(nb_brain_regions + 2), accuracy_mean, yerr = accuracy_se, fmt = 'None', color = 'black')
axes.axhline(accuracy_mean[0], 0, 1, color = 'green', linestyle = 'dashed', alpha = 0.5)
axes.set_xticks(range(nb_brain_regions + 2), labels = labels)
axes.set_xticklabels(labels, rotation = 45, ha = 'right')
axes.set_ylabel('Average accuracy across subjects')
axes.set_title('Gemma-2-2B-IT with fine-tuning');

# %% [code]
# Retrieve the average accuracy across subjects, and the standard error of the accuracy across subjects.
accuracy_mean_no_finetuning = list(accuracy_summary_gemma_no_finetuning.mean().values)
accuracy_mean_finetuning = list(accuracy_summary_gemma_finetuning.mean().values)
accuracy_se_no_finetuning = list(accuracy_summary_gemma_no_finetuning.std().values / np.sqrt(len(subjects)))
accuracy_se_finetuning = list(accuracy_summary_gemma_finetuning.std().values / np.sqrt(len(subjects)))

# Append the total average accuracy and its standard error.
nb_brain_regions = accuracy_summary_gemma_no_finetuning.shape[1]
accuracy_mean_no_finetuning.append(np.mean(accuracy_mean_no_finetuning))
accuracy_mean_finetuning.append(np.mean(accuracy_mean_finetuning))
accuracy_se_no_finetuning.append(np.std(accuracy_mean_no_finetuning) / (np.sqrt(len(subjects) * nb_brain_regions)))
accuracy_se_finetuning.append(np.std(accuracy_mean_finetuning) / (np.sqrt(len(subjects) * nb_brain_regions)))

# Create the labels.
labels = brain_regions.copy()
labels.append('Total')

# Define the bar width and positions.
bar_width = 0.4
index = np.arange(nb_brain_regions + 1)

# Display the average accuracy and its standard error in a grouped bar plot.
fig, axes = plt.subplots(nrows = 1, ncols = 1, figsize = (10, 5))
axes.bar(index - bar_width/2, accuracy_mean_no_finetuning, bar_width, color = [*['cornflowerblue'] * nb_brain_regions, 'royalblue'], label = 'No fine-tuning')
axes.bar(index + bar_width/2, accuracy_mean_finetuning, bar_width, color = [*['blue'] * nb_brain_regions, 'mediumblue'], label = 'Fine-tuning')
axes.errorbar(index - bar_width/2, accuracy_mean_no_finetuning, yerr = accuracy_se_no_finetuning, fmt = 'None', color = 'black')
axes.errorbar(index + bar_width/2, accuracy_mean_finetuning, yerr = accuracy_se_finetuning, fmt = 'None', color = 'black')
axes.axhline(accuracy_mean[0], 0, 1, color = 'green', linestyle = 'dashed', alpha = 0.5)
axes.set_xticks(index)
axes.set_xticklabels(labels, rotation = 45, ha = 'right')
axes.set_ylabel('Average accuracy across subjects')
axes.set_title('Gemma-2-2B-IT: no fine-tuning vs. fine-tuning')
axes.legend();

# %% [markdown]
## **4. Results**

# %% [code]
# Save the predictions into Pickle files. 
with open(results_path + 'classification/dict_predictions_gemma_no_finetuning.p', 'wb') as file:
    pickle.dump(dict_predictions_gemma_no_finetuning, file)
with open(results_path + 'classification/dict_predictions_gemma_finetuning.p', 'wb') as file:
    pickle.dump(dict_predictions_gemma_finetuning, file)

# Save the summaries into Pickle files. 
with open(results_path + 'classification/df_summary_gemma_no_finetuning.p', 'wb') as file:
    pickle.dump(accuracy_summary_gemma_no_finetuning, file)
with open(results_path + 'classification/df_summary_gemma_finetuning.p', 'wb') as file:
    pickle.dump(accuracy_summary_gemma_finetuning, file)

# Save the brain regions, sequences of fMRI scans, and FLOPs into Pickle files. 
with open(results_path + 'brain_regions_large_language_model_fine-tuning.p', 'wb') as file:
    pickle.dump(dict_brain_regions, file)
with open(results_path + 'fmri_scans_large_language_model_fine-tuning.p', 'wb') as file:
    pickle.dump(dict_fmri_scans, file)
with open(results_path + 'flops_large_language_model_fine-tuning.p', 'wb') as file:
    pickle.dump(dict_flops, file)
```
