## Supplementary material for "Neural Networks and Foundation Models: Two Strategies for EEG-to-fMRI Prediction": 12_Large_Multimodal_Model.ipynb

```
# %% [markdown]
# **Neural Networks and Foundation Models: Two Strategies for EEG-to-fMRI Prediction**
### **Supplementary Material**

# %% [markdown]
This project includes fourteen Notebooks:

*01: fMRI Preprocessing*

*02: EEG Preprocessing*

*03: Classification Models*

*04: Regression Models*

*05: Neural Networks*

*06: Convolutional Neural Networks*

*07: Recurrent Neural Networks*

*08: Transformers*

*09: Large Language Models*

*10: Large Language Model Chain-of-Thought*

*11: Large Language Model Fine-Tuning*

***12: Large Multimodal Model***

*13: Statistical Tests*

*14: Tables and Figures*

# %% [markdown]
### Notebook 12

# **Large Multimodal Model**

# %% [markdown]
## **1. Topographic Map**

We retrieve the EEG montage, and create a **topographic map** showing the beta bandpower across all EEG channels. 

# %% [code]
import numpy as np
import pandas as pd
import matplotlib.pyplot as plt
%matplotlib inline
import mne
import pickle
import io
from PIL import Image
import torch
from transformers import AutoProcessor, PaliGemmaForConditionalGeneration
from fvcore.nn import FlopCountAnalysis
import logging

# %% [code]
# Define the raw data path. 
raw_data_path = '../ds002336-download/'

# Define the preprocessed data path. 
preprocessed_data_path = '../preprocessed_data/'

# Define the results path. 
results_path = '../results/'

# %% [code]
# Load the features and the targets for the classification task. 
with open(preprocessed_data_path + 'dict_targets.p', 'rb') as file:
    dict_targets = pickle.load(file)
with open(preprocessed_data_path + 'dict_features_bandpowers_normalized.p', 'rb') as file:
    dict_features_bandpowers = pickle.load(file)
with open(preprocessed_data_path + 'classification/dict_targets_classification_basis.p', 'rb') as file:
    dict_targets_basis_class = pickle.load(file)

# %% [code]
# Define the list of subjects, removing sub-xp102 who has a missing condition. 
subjects = ['sub-xp1' + str(x).zfill(2) for x in range(1, 11)]
subjects.remove('sub-xp102')
subject = subjects[0]

# Retrieve and display the total number of scans. 
nb_scans_total = dict_targets[subject]['eegfmriNF']['Frontal Pole'].shape[0]
nb_scans_total

# %% [code]
# Retrieve the list of all targets. 
targets = list(dict_targets[subject]['eegfmriNF'].keys())

# Display the brain region names and the number of brain regions. 
print(targets)
print(len(targets), 'brain regions')

# %% [code]
# Define the bandpower names. 
bandpower_names = ['Delta (1-4 Hz)', 'Theta (4-8 Hz)', 'Alpha (8-12 Hz)', 'Sigma (12-16 Hz)', 'Beta (16-30 Hz)', 'Gamma (30-40 Hz)']

# Define the path of the header file (.vhdr) for this subject, for the eegfmriNF condition. 
vhdr_path = raw_data_path + 'derivatives/' + subject + '/eeg_pp/' + subject + '_task-eegfmriNF_eeg_pp.vhdr'

# Load the EEG data using the MNE library, keeping error messages but suppressing warnings. 
channels_data = mne.io.read_raw_brainvision(vhdr_path, misc = ['ECG'], preload = True, verbose = 'ERROR')

# Retrieve and display the EEG montage. 
montage = channels_data.get_montage()
montage

# %% [code]
# Retrieve and display the sampling frequency. 
sampling_frequency = channels_data.info['sfreq']
sampling_frequency

# %% [code]
# Remove the ECG channel from the channel names. 
channel_names = channels_data.ch_names
channel_names.remove('ECG')

# Display the channel names and the number of channels. 
print(channel_names)
print(len(channel_names), 'channels')

# %% [code]
# Select a condition, a bandpower index, and a scan. 
condition = 'eegfmriNF'
selected_bandpower_index = 4
selected_scan = 10

# Retrieve the bandpower data. 
bandpower_data = dict_features_bandpowers[subject][condition][:, selected_bandpower_index, selected_scan]
bandpower_data.shape

# %% [code]
# Create the measurement information object. 
info = mne.create_info(ch_names = montage.ch_names, sfreq = sampling_frequency, ch_types = 'eeg')
info.set_montage(montage)

# %% [code]
# Define a function to create a topographic map image. 
def create_topomap_image(bandpower_data, info):
    
    # Create the evoked potential array. 
    evoked = mne.EvokedArray(bandpower_data[:, np.newaxis], info)

    # Generate the topographic map, and save it into a BytesIO buffer. 
    fig = evoked.plot_topomap(times = [0], scalings = 1, cmap = 'coolwarm', contours = 0, show = False)
    buffer = io.BytesIO()
    fig.savefig(buffer, format = 'png', bbox_inches = 'tight')
    buffer.seek(0) # Reset the pointer to the beginning of the stream. 

    # Convert the figure to a PIL Image, then close the figure to free memory. 
    topomap_image = Image.open(buffer)
    plt.close(fig)

    # Resize the topographic map, and ensure it has RGB channels. 
    topomap_image = topomap_image.resize((224, 224), Image.Resampling.LANCZOS)
    topomap_image = topomap_image.convert('RGB')
    
    return topomap_image

# %% [code]
# Create and display a topographic map image. 
topomap_image = create_topomap_image(bandpower_data, info)
topomap_image

# %% [code]
# Define dictionaries to store the brain regions, sequences of fMRI scans, and FLOPs. 
dict_brain_regions = dict()
dict_fmri_scans = dict()
dict_flops = dict()

# %% [markdown]
## **2. PaliGemma**

We use the **PaliGemma2-3B-Mix-224 large multimodal model**, and select its hyperparameters in order to ensure a relative variety of responses. We define a prompt where we provide a topographic map of the beta bandpower, and ask the model to predict the corresponding fMRI activity. We run the large multimodal model on the eegfmriNF condition of all subjects, for a selection of brain regions spanning both a substantial fraction of the brain and a variety of cognitive functions. In the bar plot presenting the results, an indicative baseline is given for visualization purposes, but the real baseline depends both on the selected fMRI scans and the distribution of missing or ambiguous predictions. 

# %% [code]
# Define the LMM to use. 
model_name = 'google/paligemma2-3b-mix-224'

# Load the processor and the model. 
processor_paligemma = AutoProcessor.from_pretrained(model_name, use_fast = True)
model_paligemma = PaliGemmaForConditionalGeneration.from_pretrained(model_name)

# %% [code]
# Define a function to create a LMM prompt. 
def prompt_lmm(selected_bandpower, selected_brain_region):

    # Create the prompt. 
    prompt = (
        f"<image>A human subject participated in a neuroscience experiment where EEG data and fMRI data were recorded simultaneously. "
        f"This EEG topographic map shows the brain activity pattern observed for the band power {selected_bandpower}. "
        f"The fMRI BOLD signal was measured in the {selected_brain_region} four seconds after that. "
        f"Given the EEG data, is the fMRI signal in this brain region likely increasing or decreasing? "
        f"Base your answer on your general knowledge in neuroscience, EEG research, and fMRI research. "
        f"Please answer with only one word: Increasing or Decreasing. Just give your best prediction, without any explanation."
    )

    return prompt

# %% [code]
# Define a function to run a LMM. 
def run_lmm(prompt, image, processor, model):

    # Run the LMM. 
    inputs = processor(images = image, text = prompt, return_tensors = 'pt').to(model.device)
    output = model.generate(**inputs, 
                            max_new_tokens = 20, 
                            temperature = 1.2, 
                            top_p = 0.9, 
                            top_k = 50, 
                            do_sample = True, 
                            repetition_penalty = 1)

    # Decode the generated text, remove the original prompt, and return the answer. 
    generated_text = processor.decode(output[0], skip_special_tokens = True)
    answer_only = generated_text[len(prompt[7:]):].strip()
    
    return answer_only

# %% [code]
# Define a function to run a LMM multiple times. 
def run_lmm_series(X, Y, fmri_scans, selected_bandpower_index, selected_brain_region, info, processor, model):

    # Create the NumPy arrays to store the true values and predictions.     
    Y_true = np.zeros(len(fmri_scans))
    Y_pred = np.zeros(len(fmri_scans))
    answers = []
    counter = 0

    # Iterate through all selected fMRI scans. 
    for fmri_scan_index in fmri_scans:

        # Create the topographic map image. 
        topomap_image = create_topomap_image(X[:, fmri_scan_index], info)

        # Create the LMM prompt. 
        selected_bandpower = bandpower_names[selected_bandpower_index]
        prompt = prompt_lmm(selected_bandpower, selected_brain_region)
        
        # Run the LMM and store the result. 
        answer_only = run_lmm(prompt, topomap_image, processor, model)
        answers.append(answer_only) # For debugging purposes. 

        # Analyze the answer. 
        increase_detected = 'increas' in answer_only.lower()
        decrease_detected = 'decreas' in answer_only.lower()
        if increase_detected and not decrease_detected:
            Y_pred[counter] = 1
        elif decrease_detected and not increase_detected:
            Y_pred[counter] = 0
        else:
            Y_pred[counter] = -1

        # Store the true value, which is the fMRI signal two scans later, at the peak of the hemodynamic response. 
        Y_true[counter] = Y[fmri_scan_index + 2]

        # Increment. 
        counter += 1

    return Y_true, Y_pred

# %% [code]
# Select a subject, a condition, and a scan. 
subject = 'sub-xp101'
condition = 'eegfmriNF'
selected_scan = 5

# Select a fMRI brain region and an EEG bandpower. 
selected_brain_region = 'Frontal Pole'
selected_brain_region_index = list(targets).index(selected_brain_region)
selected_bandpower_index = 4
selected_bandpower = bandpower_names[selected_bandpower_index]

# Display the fMRI brain region and the EEG bandpower. 
print('Brain region:', selected_brain_region)
print('Brain region index:', int(selected_brain_region_index))
print('EEG bandpower:', selected_bandpower)

# %% [code]
# Retrieve the bandpower data. 
bandpower_data = dict_features_bandpowers[subject][condition][:, selected_bandpower_index, selected_scan]

# Create and display the topographic map image. 
topomap_image = create_topomap_image(bandpower_data, info)
topomap_image

# %% [code]
# Create the LMM prompt. 
prompt = prompt_lmm(selected_bandpower, selected_brain_region)
prompt[7:]

# %% [code]
# Run the LMM once, and display the result. 
answer_only = run_lmm(prompt, topomap_image, processor_paligemma, model_paligemma)
answer_only

# %% [code]
# Retrieve the features for the selected subject, condition, and EEG bandpower. 
X = dict_features_bandpowers[subject][condition][:, selected_bandpower_index, 1:]

# Retrieve the targets for the selected subject, condition, and fMRI brain region. 
Y = dict_targets_basis_class[subject][condition][:, selected_brain_region_index]

# Display the shape of the features and the targets.
print('X shape:', X.shape)
print('Y shape:', Y.shape)

# %% [code]
# Run the LMM multiple times. 
fmri_scans = range(5, (nb_scans_total - 3), 20)
Y_true, Y_pred = run_lmm_series(X, Y, fmri_scans, selected_bandpower_index, selected_brain_region, info, processor_paligemma, model_paligemma)

# Compute the average accuracy. 
accuracy_array = Y_pred[Y_pred != -1] == Y_true[Y_pred != -1]
accuracy_sequence = np.sum(accuracy_array) / accuracy_array.shape[0]
accuracy_sequence

# %% [code]
# Select a condition, a bandpower index, and a list of brain regions. 
condition = 'eegfmriNF'
selected_bandpower_index = 4
brain_regions = ['Frontal Pole', # High-order cognitive functions, but sometimes noisy EEG signal. 
                 'Insular Cortex', # Integrative functions. 
                 'Inferior Frontal Gyrus, pars triangularis', # Language and executive functions. 
                 'Precentral Gyrus', # Primary motor cortex. 
                 'Superior Temporal Gyrus, posterior division', # Auditory functions. 
                 'Superior Parietal Lobule', # Attention functions. 
                 'Angular Gyrus', # Default-mode network. 
                 'Occipital Pole'] # Primary visual cortex. 

# Select a sequence of fMRI scans. 
fmri_scans = range(5, (nb_scans_total - 1), 20) # Extensive search.
#fmri_scans = range(5, (nb_scans_total - 1), 100) # Quick search for debugging.

# Store the brain regions and the sequence of fMRI scans in dictionaries. 
dict_brain_regions['Large Multimodal Model'] = brain_regions
dict_fmri_scans['Large Multimodal Model'] = fmri_scans

# Define a dictionary to store the predictions and a Pandas DataFrame to store the accuracy. 
dict_predictions_paligemma = dict()
accuracy_summary_paligemma = pd.DataFrame(columns = brain_regions)

# Retrieve the selected bandpower. 
selected_bandpower = bandpower_names[selected_bandpower_index]

# Iterate through all brain regions. 
for selected_brain_region in brain_regions:

    # Display the current brain region. 
    print(selected_brain_region)

    # Define a dictionary to store the predictions for this brain region. 
    dict_predictions_paligemma[selected_brain_region] = dict()

    # Iterate through all subjects. 
    for subject in subjects:

        # Retrieve the features for the selected subject, condition, and EEG bandpower. 
        X = dict_features_bandpowers[subject][condition][:, selected_bandpower_index, 1:]

        # Retrieve the targets for the selected subject, condition, and fMRI brain region. 
        Y = dict_targets_basis_class[subject][condition][:, selected_brain_region_index]

        # Run the LMM multiple times. 
        Y_true, Y_pred = run_lmm_series(X, Y, fmri_scans, selected_bandpower_index, selected_brain_region, info, processor_paligemma, model_paligemma)

        # Store the predictions. 
        dict_predictions_paligemma[selected_brain_region][subject] = Y_pred

        # Compute and store the average accuracy. 
        accuracy_array = Y_pred[Y_pred != -1] == Y_true[Y_pred != -1]
        accuracy_sequence = np.sum(accuracy_array) / accuracy_array.shape[0]
        accuracy_summary_paligemma.loc[subject, selected_brain_region] = accuracy_sequence

        # Display the number of ambiguous predictions. 
        print(subject + ': ' + str(np.sum(Y_pred == -1)), 'ambiguous prediction(s) out of', len(Y_pred), '-', str(np.sum(Y_pred == -1) / len(Y_pred) * 100), '%')

# %% [code]
# Set the logging level to suppress warnings. 
logging.getLogger('fvcore.nn.jit_handles').setLevel(logging.ERROR)
logging.getLogger('fvcore.nn.jit_analysis').setLevel(logging.ERROR)

# Store current device. 
device_before = next(model_paligemma.parameters()).device
print('Original device of the model:', device_before)

# Define a class wrapping the model. 
# We output only logits (Tensor data type) for compatibility with FlopCountAnalysis. 
class WrappedModel(torch.nn.Module):
    def __init__(self, model):
        super().__init__()
        self.model = model
    def forward(self, input_ids):
        output = self.model(input_ids)
        return output.logits

# Move the model to CPU and prepare dummy input for FLOPs analysis. 
model_paligemma_cpu = WrappedModel(model_paligemma).to('cpu')
dummy_input = (torch.ones((1, len(prompt)), dtype = torch.long),) # batch_size = 1, seq_len = prompt length
print('Prompt length:', len(prompt))

# Measure the FLOPs. 
flops = FlopCountAnalysis(model_paligemma_cpu, dummy_input)
print(f'Total FLOPs: {flops.total():,}')
print('FLOPs by operator:', flops.by_operator())

# Move the model back to its original device. 
model_paligemma.to(device_before)
print('Model moved back to:', device_before)

# %% [code]
# Store and display the number of FLOPs. 
dict_flops['Large Multimodal Model'] = flops.total()
print('FLOPs per inference:', flops.total())

# %% [code]
# Display the accuracies per subject and brain region. 
accuracy_summary_paligemma

# %% [code]
# Retrieve the average accuracy across subjects, and the standard error of the accuracy across subjects. 
accuracy_mean = list(accuracy_summary_paligemma.mean().values)
accuracy_mean.insert(0, 0.5) # Insert a baseline average of 0.5. 
accuracy_se = list(accuracy_summary_paligemma.std().values / np.sqrt(len(subjects)))
accuracy_se.insert(0, 0) # Insert a baseline standard error of 0. 

# Append the total average accuracy and its standard error. 
nb_brain_regions = accuracy_summary_paligemma.shape[1]
accuracy_mean.append(np.mean(accuracy_mean[1:]))
accuracy_se.append(np.std(accuracy_mean[1:]) / (np.sqrt(len(subjects) * nb_brain_regions)))

# Create the labels. 
labels = brain_regions.copy()
labels.insert(0, 'Baseline')
labels.append('Total')

# Display the average accuracy and its standard error in a bar plot. 
fig, axes = plt.subplots(nrows = 1, ncols = 1, figsize = (10, 4))
axes.bar(range(nb_brain_regions + 2), accuracy_mean, width = 0.6, color = ['green', *['blue'] * nb_brain_regions, 'mediumblue'])
axes.errorbar(range(nb_brain_regions + 2), accuracy_mean, yerr = accuracy_se, fmt = 'None', color = 'black')
axes.axhline(accuracy_mean[0], 0, 1, color = 'green', linestyle = 'dashed', alpha = 0.5)
axes.set_xticks(range(nb_brain_regions + 2), labels = labels)
axes.set_xticklabels(labels, rotation = 45, ha = 'right')
axes.set_ylabel('Average accuracy across subjects')
axes.set_title('PaliGemma2-3B-Mix-224');

# %% [markdown]
## **3. Results**

# %% [code]
# Save the predictions into a Pickle file. 
with open(results_path + 'classification/dict_predictions_paligemma.p', 'wb') as file:
    pickle.dump(dict_predictions_paligemma, file)

# Save the summary into a Pickle file. 
with open(results_path + 'classification/df_summary_paligemma.p', 'wb') as file:
    pickle.dump(accuracy_summary_paligemma, file)

# Save the brain regions, sequences of fMRI scans, and FLOPs into Pickle files. 
with open(results_path + 'brain_regions_large_multimodal_model.p', 'wb') as file:
    pickle.dump(dict_brain_regions, file)
with open(results_path + 'fmri_scans_large_multimodal_model.p', 'wb') as file:
    pickle.dump(dict_fmri_scans, file)
with open(results_path + 'flops_large_multimodal_model.p', 'wb') as file:
    pickle.dump(dict_flops, file)
```
