## Supplementary material for "Neural Networks and Foundation Models: Two Strategies for EEG-to-fMRI Prediction": 13_Statistical_Tests.ipynb

```
# %% [markdown]
# **Neural Networks and Foundation Models: Two Strategies for EEG-to-fMRI Prediction**
### **Supplementary Material**

# %% [markdown]
This project includes fourteen Notebooks:

*01: fMRI Preprocessing*

*02: EEG Preprocessing*

*03: Classification Models*

*04: Regression Models*

*05: Neural Networks*

*06: Convolutional Neural Networks*

*07: Recurrent Neural Networks*

*08: Transformers*

*09: Large Language Models*

*10: Large Language Model Chain-of-Thought*

*11: Large Language Model Fine-Tuning*

*12: Large Multimodal Model*

***13: Statistical Tests***

*14: Tables and Figures*

# %% [markdown]
### Notebook 13

# **Statistical Tests**

# %% [markdown]
## **1. Machine Learning**

We perform **Wilcoxon signed-rank tests** on the results of the machine learning models for the classification and regression tasks. 

# %% [code]
import numpy as np
import pandas as pd
import matplotlib.pyplot as plt
%matplotlib inline
import pickle
import scipy
import pingouin
from statsmodels.stats.contingency_tables import mcnemar

# %% [code]
# Define the preprocessed data path. 
preprocessed_data_path = '../preprocessed_data/'

# Define the results path. 
results_path = '../results/'

# %% [code]
# Load the targets. 
with open(preprocessed_data_path + 'dict_targets.p', 'rb') as file:
    dict_targets = pickle.load(file)
with open(preprocessed_data_path + 'classification/dict_targets_classification_sequence.p', 'rb') as file:
    dict_targets_class = pickle.load(file)
with open(preprocessed_data_path + 'regression/dict_targets_regression_sequence.p', 'rb') as file:
    dict_targets_reg = pickle.load(file)
with open(preprocessed_data_path + 'classification/dict_targets_classification_basis.p', 'rb') as file:
    dict_targets_foundation_models = pickle.load(file)

# Load the machine learning results for the classification task. 
with open(results_path + 'classification/dict_predictions_lr.p', 'rb') as file:
    dict_predictions_lr_class = pickle.load(file)
with open(results_path + 'classification/dict_predictions_knn.p', 'rb') as file:
    dict_predictions_knn_class = pickle.load(file)
with open(results_path + 'classification/dict_predictions_dt.p', 'rb') as file:
    dict_predictions_dt_class = pickle.load(file)
with open(results_path + 'classification/dict_predictions_rf.p', 'rb') as file:
    dict_predictions_rf_class = pickle.load(file)
with open(results_path + 'classification/dict_predictions_svm.p', 'rb') as file:
    dict_predictions_svm_class = pickle.load(file)
with open(results_path + 'classification/dict_predictions_xgb.p', 'rb') as file:
    dict_predictions_xgb_class = pickle.load(file)

# %% [code]
# Define the list of subjects, removing sub-xp102 who has a missing condition. 
subjects = ['sub-xp1' + str(x).zfill(2) for x in range(1, 11)]
subjects.remove('sub-xp102')
subject = subjects[0]

# Retrieve and display the brain region names and the number of brain regions. 
brain_regions = list(dict_targets[subject]['eegfmriNF'].keys())
print(brain_regions)
print(len(brain_regions), 'brain regions')

# %% [code]
# Define a function to create the DataFrame for the statistical tests. 
def create_df(dict_targets, dict_predictions, brain_regions, score_type, model_type):

    # Define Pandas DataFrames to store the results. 
    df = pd.DataFrame(columns = ['subject', 'brain_region', 'model_' + score_type, 'baseline_' + score_type])
    counter = 0

    # Iterate through all subjects. 
    for subject in subjects:

        # Iterate through all brain regions. 
        for brain_region_index in range(len(brain_regions)):

            # Fill the DataFrame with the subject and brain region. 
            df.loc[counter, 'subject'] = subject
            df.loc[counter, 'brain_region'] = brain_regions[brain_region_index]

            # Retrieve the targets and predictions values for the current subject and brain region. 
            targets_values = dict_targets[subject]['fmriNF'][:, brain_region_index]
            if model_type == 'machine_learning':
                predictions_values = dict_predictions[subject]['fmriNF'][:, brain_region_index]
            elif model_type == 'deep_learning':
                predictions_values = dict_predictions[subject][:, brain_region_index]

            # For the classification task, compute the accuracy. 
            if score_type == 'accuracy':
                model_score = np.mean(targets_values == predictions_values)
                baseline_score = np.mean(targets_values == 1)
                
            # For the regression task, compute the mean absolute error (MAE).
            elif score_type == 'MAE':
                model_score = np.mean(np.abs(targets_values - predictions_values))
                baseline_score = np.mean(np.abs(targets_values - np.mean(targets_values)))

            # Fill the DataFrame with the model score and baseline score. 
            df.loc[counter, 'model_' + score_type] = model_score
            df.loc[counter, 'baseline_' + score_type] = baseline_score

            # Increment. 
            counter += 1

    # Return the DataFrame. 
    return df

# %% [code]
# Example: Create and display the DataFrame for the logistic regression model. 
df_lr_class = create_df(dict_targets_class, dict_predictions_lr_class, brain_regions, 'accuracy', 'machine_learning')
df_lr_class

# %% [code]
# Define a function to perform a Wilcoxon signed-rank test. 
def wilcoxon_test(df, score_type):

    # Compute the mean and standard deviation of the model and the baseline. 
    mean_model_score = df['model_' + score_type].mean()
    mean_baseline_score = df['baseline_' + score_type].mean()
    std_model_score = df['model_' + score_type].std()
    std_baseline_score = df['baseline_' + score_type].std()

    # Perform the Wilcoxon signed-rank test using SciPy. 
    x = df['model_' + score_type].astype(float).values
    y = df['baseline_' + score_type].astype(float).values
    if score_type == 'accuracy':
        wilcoxon_stat, p_value = scipy.stats.wilcoxon(x, y, alternative = 'greater')
    elif score_type == 'MAE':
        wilcoxon_stat, p_value = scipy.stats.wilcoxon(x, y, alternative = 'less')

    # Compute the confidence interval of the mean. 
    res = scipy.stats.bootstrap((df['model_' + score_type],), np.mean, confidence_level = 0.95)
    ci_low, ci_high = res.confidence_interval

    # Store the statistics in a DataFrame. 
    statistics_scipy = pd.DataFrame([{
        'N': len(df),
        'mean_model_' + score_type: mean_model_score,
        'mean_baseline_' + score_type: mean_baseline_score,
        'std_model_' + score_type: std_model_score,
        'std_baseline_' + score_type: std_baseline_score,
        'wilcoxon_stat': wilcoxon_stat,
        'p_value': p_value,
        'ci_low': ci_low, 
        'ci_high': ci_high
    }])

    # Perform the same test using Pingouin to obtain the RBC (Rank-Biserial Correlation) and CLES (Common Language Effect Size). 
    if score_type == 'accuracy':
        statistics_pingouin = pingouin.wilcoxon(x, y, alternative = 'greater')
    elif score_type == 'MAE':
        statistics_pingouin = pingouin.wilcoxon(x, y, alternative = 'less')
    
    # Return the statistics. 
    return statistics_scipy, statistics_pingouin

# %% [code]
# Perform the Wilcoxon signed-rank test. 
statistics_scipy_lr_class, statistics_pingouin_lr_class = wilcoxon_test(df_lr_class, 'accuracy')

# %% [code]
# Display the SciPy statistics. 
statistics_scipy_lr_class

# %% [code]
# Display the Pingouin statistics. 
statistics_pingouin_lr_class

# %% [code]
# Define a function to create a summary DataFrame. 
def create_summary_df(model_names, score_type):
    
    # Create the summary DataFrame. 
    df_summary = pd.DataFrame(index = model_names, 
                              columns = ['N', 
                                         'mean model ' + score_type, 
                                         'mean baseline ' + score_type, 
                                         'STD model ' + score_type,
                                         'STD baseline ' + score_type,
                                         'Wilcoxon W statistic', 
                                         'p-value', 
                                         'CI (lower)',
                                         'CI (upper)', 
                                         'RBC', 
                                         'CLES'])
    
    return df_summary

# %% [code]
# Define a function to store statistics in the summary DataFrame. 
def store_statistics_in_summary_df(df_summary, model_names, model_index, statistics_scipy, statistics_pingouin, score_type):

    # Store the statistics in the summary DataFrame. 
    df_summary.loc[model_names[model_index], 'N'] = statistics_scipy['N'].values[0]
    df_summary.loc[model_names[model_index], 'mean model ' + score_type] = statistics_scipy['mean_model_' + score_type].values[0]
    df_summary.loc[model_names[model_index], 'mean baseline ' + score_type] = statistics_scipy['mean_baseline_' + score_type].values[0]
    df_summary.loc[model_names[model_index], 'STD model ' + score_type] = statistics_scipy['std_model_' + score_type].values[0]
    df_summary.loc[model_names[model_index], 'STD baseline ' + score_type] = statistics_scipy['std_baseline_' + score_type].values[0]
    df_summary.loc[model_names[model_index], 'Wilcoxon W statistic'] = statistics_scipy['wilcoxon_stat'].values[0]
    df_summary.loc[model_names[model_index], 'p-value'] = statistics_scipy['p_value'].values[0]
    df_summary.loc[model_names[model_index], 'CI (lower)'] = statistics_scipy['ci_low'].values[0]
    df_summary.loc[model_names[model_index], 'CI (upper)'] = statistics_scipy['ci_high'].values[0]
    df_summary.loc[model_names[model_index], 'RBC'] = statistics_pingouin['RBC'].values[0]
    df_summary.loc[model_names[model_index], 'CLES'] = statistics_pingouin['CLES'].values[0]

    # Ensure the p-value column is numeric, then format it in scientific notation. 
    df_summary['p-value'] = pd.to_numeric(df_summary['p-value'], errors = 'coerce')
    df_summary['p-value'] = df_summary['p-value'].apply(lambda x: f'{x:.2e}')

    return df_summary

# %% [code]
# Define a function to perform a Wilcoxon signed-rank test on a series of models. 
def run_wilcoxon_tests(model_names, dict_targets, list_dict_predictions, brain_regions, score_type, model_type):

    # Create a summary DataFrame to store the statistics for all models. 
    df_summary = create_summary_df(model_names, score_type)

    # Iterate through all models. 
    for model_index in range(len(model_names)):

        # Create the DataFrame and perform the Wilcoxon signed-rank test. 
        dict_predictions = list_dict_predictions[model_index]
        df = create_df(dict_targets, dict_predictions, brain_regions, score_type, model_type)
        statistics_scipy, statistics_pingouin = wilcoxon_test(df, score_type)

        # Store the statistics in the summary DataFrame. 
        df_summary = store_statistics_in_summary_df(df_summary, model_names, model_index, statistics_scipy, statistics_pingouin, score_type)
        
    # Return the summary DataFrame. 
    return df_summary

# %% [code]
# Define the machine learning models for classification. 
model_names = ['Logistic Regression', 'K-Nearest Neighbors', 'Decision Tree', 'Random Forest', 'Support Vector Machine', 'XGBoost']
predictions_ml_class = [dict_predictions_lr_class, 
                        dict_predictions_knn_class, 
                        dict_predictions_dt_class, 
                        dict_predictions_rf_class, 
                        dict_predictions_svm_class, 
                        dict_predictions_xgb_class]

# Run the Wilcoxon signed-rank tests for all models. 
df_summary_ml_class = run_wilcoxon_tests(model_names, dict_targets_class, predictions_ml_class, brain_regions, 'accuracy', 'machine_learning')
df_summary_ml_class

# %% [code]
# Load the machine learning results for the regression task. 
with open(results_path + 'regression/dict_predictions_lr.p', 'rb') as file:
    dict_predictions_lr_reg = pickle.load(file)
with open(results_path + 'regression/dict_predictions_ridge.p', 'rb') as file:
    dict_predictions_ridge_reg = pickle.load(file)
with open(results_path + 'regression/dict_predictions_lasso.p', 'rb') as file:
    dict_predictions_lasso_reg = pickle.load(file)
with open(results_path + 'regression/dict_predictions_knn.p', 'rb') as file:
    dict_predictions_knn_reg = pickle.load(file)
with open(results_path + 'regression/dict_predictions_dt.p', 'rb') as file:
    dict_predictions_dt_reg = pickle.load(file)
with open(results_path + 'regression/dict_predictions_rf.p', 'rb') as file:
    dict_predictions_rf_reg = pickle.load(file)
with open(results_path + 'regression/dict_predictions_svm.p', 'rb') as file:
    dict_predictions_svm_reg = pickle.load(file)
with open(results_path + 'regression/dict_predictions_xgb.p', 'rb') as file:
    dict_predictions_xgb_reg = pickle.load(file)

# %% [code]
# Example: Create and display the DataFrame for the support vector machine (SVM) model. 
df_svm_reg = create_df(dict_targets_reg, dict_predictions_svm_reg, brain_regions, 'MAE', 'machine_learning')
df_svm_reg

# %% [code]
# Perform Wilcoxon signed-rank test. 
statistics_scipy_svm_reg, statistics_pingouin_svm_reg = wilcoxon_test(df_svm_reg, 'MAE')

# %% [code]
# Display the SciPy statistics. 
statistics_scipy_svm_reg

# %% [code]
# Display the Pingouin statistics. 
statistics_pingouin_svm_reg

# %% [code]
# Define the machine learning models for regression. 
model_names = ['Linear Regression', 'Ridge Regression', 'Lasso Regression', 'K-Nearest Neighbors', 'Decision Tree', 'Random Forest', 'Support Vector Machine', 'XGBoost']
predictions_ml_reg = [dict_predictions_lr_reg,
                      dict_predictions_ridge_reg, 
                      dict_predictions_lasso_reg, 
                      dict_predictions_knn_reg, 
                      dict_predictions_dt_reg, 
                      dict_predictions_rf_reg, 
                      dict_predictions_svm_reg, 
                      dict_predictions_xgb_reg]

# Run the Wilcoxon signed-rank tests for all models. 
df_summary_ml_reg = run_wilcoxon_tests(model_names, dict_targets_reg, predictions_ml_reg, brain_regions, 'MAE', 'machine_learning')
df_summary_ml_reg

# %% [markdown]
## **2. Deep Learning**

We perform **Wilcoxon signed-rank tests** on the results of the deep learning models for the classification and regression tasks. 

# %% [code]
# Load the deep learning results for the classification task. 
with open(results_path + 'classification/neural_networks/dict_predictions_neural_networks_class.p', 'rb') as file:
    dict_predictions_neural_networks_class = pickle.load(file)
with open(results_path + 'classification/convolutional_neural_networks/dict_predictions_convolutional_neural_networks_class.p', 'rb') as file:
    dict_predictions_convolutional_neural_networks_class = pickle.load(file)
with open(results_path + 'classification/recurrent_neural_networks/dict_predictions_recurrent_neural_networks_class.p', 'rb') as file:
    dict_predictions_recurrent_neural_networks_class = pickle.load(file)
with open(results_path + 'classification/transformers/dict_predictions_transformers_class.p', 'rb') as file:
    dict_predictions_transformers_class = pickle.load(file)

# %% [code]
# Example: Create and display the DataFrame for the neural network model. 
df_neural_networks_class = create_df(dict_targets_class, dict_predictions_neural_networks_class, brain_regions, 'accuracy', 'deep_learning')
df_neural_networks_class

# %% [code]
# Perform Wilcoxon signed-rank test. 
statistics_scipy_neural_networks_class, statistics_pingouin_neural_networks_class = wilcoxon_test(df_neural_networks_class, 'accuracy')

# %% [code]
# Display the SciPy statistics. 
statistics_scipy_neural_networks_class

# %% [code]
# Display the Pingouin statistics. 
statistics_pingouin_neural_networks_class

# %% [code]
# Define the deep learning models for classification. 
model_names = ['Multi-Layer Perceptron', 'Convolutional Neural Network', 'Recurrent Neural Network', 'Transformer']
predictions_dl_class = [dict_predictions_neural_networks_class, 
                        dict_predictions_convolutional_neural_networks_class, 
                        dict_predictions_recurrent_neural_networks_class, 
                        dict_predictions_transformers_class]

# Run the Wilcoxon signed-rank tests for all models. 
df_summary_dl_class = run_wilcoxon_tests(model_names, dict_targets_class, predictions_dl_class, brain_regions, 'accuracy', 'deep_learning')
df_summary_dl_class

# %% [code]
# Load the deep learning results for the regression task. 
with open(results_path + 'regression/neural_networks/dict_predictions_neural_networks_reg.p', 'rb') as file:
    dict_predictions_neural_networks_reg = pickle.load(file)
with open(results_path + 'regression/convolutional_neural_networks/dict_predictions_convolutional_neural_networks_reg.p', 'rb') as file:
    dict_predictions_convolutional_neural_networks_reg = pickle.load(file)
with open(results_path + 'regression/recurrent_neural_networks/dict_predictions_recurrent_neural_networks_reg.p', 'rb') as file:
    dict_predictions_recurrent_neural_networks_reg = pickle.load(file)
with open(results_path + 'regression/transformers/dict_predictions_transformers_reg.p', 'rb') as file:
    dict_predictions_transformers_reg = pickle.load(file)

# %% [code]
# Define the deep learning models for regression. 
model_names = ['Multi-Layer Perceptron', 'Convolutional Neural Network', 'Recurrent Neural Network', 'Transformer']
predictions_dl_reg = [dict_predictions_neural_networks_reg, 
                      dict_predictions_convolutional_neural_networks_reg, 
                      dict_predictions_recurrent_neural_networks_reg, 
                      dict_predictions_transformers_reg]

# Run the Wilcoxon signed-rank tests for all models. 
df_summary_dl_reg = run_wilcoxon_tests(model_names, dict_targets_reg, predictions_dl_reg, brain_regions, 'MAE', 'deep_learning')
df_summary_dl_reg

# %% [code]
# Load the FLOPs. 
with open(results_path + 'flops_neural_networks.p', 'rb') as file:
    dict_flops_neural_networks = pickle.load(file)
with open(results_path + 'flops_convolutional_neural_networks.p', 'rb') as file:
    dict_flops_convolutional_neural_networks = pickle.load(file)
with open(results_path + 'flops_recurrent_neural_networks.p', 'rb') as file:
    dict_flops_recurrent_neural_networks = pickle.load(file)
with open(results_path + 'flops_transformers.p', 'rb') as file:
    dict_flops_transformers = pickle.load(file)

# %% [code]
# Store the FLOPs in a DataFrame. 
df_flops_deep_learning = pd.DataFrame({
    'FLOPs for classification task': {
        'Multi-Layer Perceptron': list(dict_flops_neural_networks.values())[0],
        'Convolutional Neural Network': list(dict_flops_convolutional_neural_networks.values())[0],
        'Recurrent Neural Network': list(dict_flops_recurrent_neural_networks.values())[0],
        'Transformer': list(dict_flops_transformers.values())[0]
    },
    'FLOPs for regression task': {
        'Multi-Layer Perceptron': list(dict_flops_neural_networks.values())[1],
        'Convolutional Neural Network': list(dict_flops_convolutional_neural_networks.values())[1],
        'Recurrent Neural Network': list(dict_flops_recurrent_neural_networks.values())[1],
        'Transformer': list(dict_flops_transformers.values())[1]
    }
})

# Display the DataFrame. 
df_flops_deep_learning

# %% [markdown]
## **3. Foundation Models (Wilcoxon Tests)**

We perform **Wilcoxon signed-rank tests** on the results of the foundation models. 

# %% [code]
# Load the foundation models results. 
with open(results_path + 'classification/dict_predictions_gemma.p', 'rb') as file:
    dict_predictions_gemma = pickle.load(file)
with open(results_path + 'classification/dict_predictions_llama.p', 'rb') as file:
    dict_predictions_llama = pickle.load(file)
with open(results_path + 'classification/dict_predictions_gemma_CoT.p', 'rb') as file:
    dict_predictions_gemma_CoT = pickle.load(file)
with open(results_path + 'classification/dict_predictions_gemma_no_finetuning.p', 'rb') as file:
    dict_predictions_gemma_no_finetuning = pickle.load(file)
with open(results_path + 'classification/dict_predictions_gemma_finetuning.p', 'rb') as file:
    dict_predictions_gemma_finetuning = pickle.load(file)
with open(results_path + 'classification/dict_predictions_paligemma.p', 'rb') as file:
    dict_predictions_paligemma = pickle.load(file)

# Load the brain regions. 
with open(results_path + 'brain_regions_large_language_models.p', 'rb') as file:
    dict_brain_regions_large_language_models = pickle.load(file)
with open(results_path + 'brain_regions_large_language_model_CoT.p', 'rb') as file:
    dict_brain_regions_large_language_model_CoT = pickle.load(file)
with open(results_path + 'brain_regions_large_language_model_fine-tuning.p', 'rb') as file:
    dict_brain_regions_large_language_model_fine_tuning = pickle.load(file)
with open(results_path + 'brain_regions_large_multimodal_model.p', 'rb') as file:
    dict_brain_regions_large_multimodal_model = pickle.load(file)

# Load the fMRI scans. 
with open(results_path + 'fmri_scans_large_language_models.p', 'rb') as file:
    dict_fmri_scans_large_language_models = pickle.load(file)
with open(results_path + 'fmri_scans_large_language_model_CoT.p', 'rb') as file:
    dict_fmri_scans_large_language_model_CoT = pickle.load(file)
with open(results_path + 'fmri_scans_large_language_model_fine-tuning.p', 'rb') as file:
    dict_fmri_scans_large_language_model_fine_tuning = pickle.load(file)
with open(results_path + 'fmri_scans_large_multimodal_model.p', 'rb') as file:
    dict_fmri_scans_large_multimodal_model = pickle.load(file)

# Load the FLOPs. 
with open(results_path + 'flops_large_language_models.p', 'rb') as file:
    dict_flops_large_language_models = pickle.load(file)
with open(results_path + 'flops_large_language_model_CoT.p', 'rb') as file:
    dict_flops_large_language_model_CoT = pickle.load(file)
with open(results_path + 'flops_large_language_model_fine-tuning.p', 'rb') as file:
    dict_flops_large_language_model_fine_tuning = pickle.load(file)
with open(results_path + 'flops_large_multimodal_model.p', 'rb') as file:
    dict_flops_large_multimodal_model = pickle.load(file)

# %% [code]
# Define the foundation models. 
model_names = ['Gemma', 'Llama', 'Gemma Chain-of-Thought', 'Gemma Without Fine-Tuning', 'Gemma With Fine-Tuning', 'PaliGemma']
list_dict_predictions = [dict_predictions_gemma, 
                         dict_predictions_llama, 
                         dict_predictions_gemma_CoT, 
                         dict_predictions_gemma_no_finetuning, 
                         dict_predictions_gemma_finetuning, 
                         dict_predictions_paligemma]

# Define the brain regions. 
list_brain_regions = [dict_brain_regions_large_language_models['Large Language Models: Gemma'],
                      dict_brain_regions_large_language_models['Large Language Models: Llama'],
                      dict_brain_regions_large_language_model_CoT['Large Language Model Chain-of-Thought'],
                      dict_brain_regions_large_language_model_fine_tuning['Large Language Model Fine-Tuning: Without Fine-Tuning'],
                      dict_brain_regions_large_language_model_fine_tuning['Large Language Model Fine-Tuning: With Fine-Tuning'], 
                      dict_brain_regions_large_multimodal_model['Large Multimodal Model']]

# Define the fMRI scans indexes. 
list_fmri_scans_indexes = [dict_fmri_scans_large_language_models['Large Language Models: Gemma'],
                           dict_fmri_scans_large_language_models['Large Language Models: Llama'],
                           dict_fmri_scans_large_language_model_CoT['Large Language Model Chain-of-Thought'],
                           dict_fmri_scans_large_language_model_fine_tuning['Large Language Model Fine-Tuning: Without Fine-Tuning'],
                           dict_fmri_scans_large_language_model_fine_tuning['Large Language Model Fine-Tuning: With Fine-Tuning'], 
                           dict_fmri_scans_large_multimodal_model['Large Multimodal Model']]

# %% [code]
# Define a function to create a Pandas DataFrame of predictions values. 
def create_df_predictions_values(dict_predictions):

    # Define a list to store the predictions. 
    list_pred = []

    # Iterate through all brain regions and subjects. 
    for brain_region in dict_predictions.keys():
        for subject in subjects:
            list_pred.append(dict_predictions[brain_region][subject].flatten())

    # Flatten the predictions for all brain regions and subjects. 
    list_pred = np.array(list_pred).flatten()

    # Transform into a Pandas DataFrame. 
    df_pred = pd.DataFrame(list_pred, columns = ['predictions'])

    return df_pred['predictions']

# %% [code]
# Create a Pandas DataFrame to store the predictions count values. 
df_predictions_count_values = pd.DataFrame()

# Iterate through all models. 
for dict_predictions in list_dict_predictions:

    # Count the values in the predictions. 
    df_predictions_values = create_df_predictions_values(dict_predictions)
    df_predictions_count_values = pd.concat([df_predictions_count_values, df_predictions_values.value_counts()], axis = 1)

# Change the row names. 0 = decreasing, 1 = increasing, -1 = ambiguous. 
df_predictions_count_values = df_predictions_count_values.rename(index = {0: 'decreasing', 1: 'increasing', -1: 'ambiguous'})

# Change the column names, replacing them by the names of the models. 
df_predictions_count_values.columns = model_names

# Display the DataFrame. 
df_predictions_count_values

# %% [code]
# Compute the proportion of decreasing and increasing predictions, excluding the ambiguous predictions. 
df_predictions_count_values_without_ambiguous = df_predictions_count_values.drop(index = 'ambiguous')
df_predictions_count_values_proportion = df_predictions_count_values_without_ambiguous.div(df_predictions_count_values_without_ambiguous.sum(axis = 0), axis = 1)
df_predictions_count_values_proportion

# %% [code]
# Define a function to create the DataFrame for the statistical tests. 
def create_df_foundation_models(dict_targets, dict_predictions, brain_region_indexes, fmri_scans_indexes):
    
    # Define a Pandas DataFrames and two lists to store the results. 
    df = pd.DataFrame(columns = ['subject', 'brain_region', 'model_accuracy', 'baseline_accuracy'])
    list_targets = []
    list_predictions = []
    counter = 0

    # Iterate through all subjects. 
    for subject in subjects:

        # Iterate through all brain regions. 
        for brain_region_index in brain_region_indexes:
            
            # Fill the DataFrame with the subject and brain region. 
            df.loc[counter, 'subject'] = subject
            df.loc[counter, 'brain_region'] = brain_regions[brain_region_index]

            # Retrieve the targets and predictions values for the current subject and brain region. 
            targets_values = dict_targets[subject]['eegfmriNF'][fmri_scans_indexes, brain_region_index]
            predictions_values = dict_predictions[brain_regions[brain_region_index]][subject]

            # Add the targets and predictions to the lists. 
            list_targets.append(targets_values)
            list_predictions.append(predictions_values)

            # Remove the cases where the prediction is -1. 
            selected_predictions = predictions_values != -1
            targets_values = targets_values[selected_predictions]
            predictions_values = predictions_values[selected_predictions]

            # Compute the accuracy and fill in the DataFrame with the model score. 
            model_score = np.mean(targets_values == predictions_values)
            df.loc[counter, 'model_accuracy'] = model_score

            # Define the values and compute the targets distribution. 
            values = [0, 1]
            targets_distribution = [np.sum(targets_values == 0) / len(targets_values), 
                                    np.sum(targets_values == 1) / len(targets_values)]

            # Define the number of iterations. 
            iterations = 1000
            baseline_score = 0

            # Compute the baseline score using random sampling. 
            for i in range(iterations):
                samples = np.random.choice(values, size = len(targets_values), p = targets_distribution)
                baseline_score += np.mean(targets_values == samples)

            # Compute the mean baseline score. 
            baseline_score /= iterations

            # Fill in the DataFrame with the baseline score. 
            df.loc[counter, 'baseline_accuracy'] = baseline_score

            # Increment. 
            counter += 1

    return df, list_targets, list_predictions

# %% [code]
# Create a function to retrieve foundation model elements. 
def retrieve_foundation_model_elements(model_index, list_dict_predictions, list_brain_regions, list_fmri_scans_indexes):

    # Retrieve the foundation model elements. 
    dict_predictions = list_dict_predictions[model_index]
    brain_region_indexes = [brain_regions.index(region) for region in list_brain_regions[model_index]]
    fmri_scans_indexes = list_fmri_scans_indexes[model_index]

    return dict_predictions, brain_region_indexes, fmri_scans_indexes

# %% [code]
# Example: Create and display the DataFrame for the Gemma model. 
model_index = 0
dict_predictions, brain_region_indexes, fmri_scans_indexes = retrieve_foundation_model_elements(model_index, 
                                                                                                list_dict_predictions,
                                                                                                list_brain_regions,
                                                                                                list_fmri_scans_indexes)

# Create and display the DataFrame. 
df_gemma, _, _ = create_df_foundation_models(dict_targets_foundation_models, 
                                             dict_predictions, 
                                             brain_region_indexes, 
                                             fmri_scans_indexes)
df_gemma

# %% [code]
# Perform Wilcoxon signed-rank test. 
statistics_scipy_gemma, statistics_pingouin_gemma = wilcoxon_test(df_gemma, 'accuracy')

# %% [code]
# Display the SciPy statistics. 
statistics_scipy_gemma

# %% [code]
# Display the Pingouin statistics. 
statistics_pingouin_gemma

# %% [code]
# Define a function to perform a Wilcoxon signed-rank test on a series of models. 
def run_wilcoxon_tests_foundation_models(model_names, dict_targets, list_dict_predictions, list_brain_regions, list_fmri_scans_indexes):

    # Create a summary DataFrame to store the statistics for all models. 
    df_summary = create_summary_df(model_names, 'accuracy')

    # Iterate through all models. 
    for model_index in range(len(model_names)):

        # Retrieve the predictions, brain region indexes, and fMRI scans indexes. 
        dict_predictions, brain_region_indexes, fmri_scans_indexes = retrieve_foundation_model_elements(model_index, 
                                                                                                        list_dict_predictions,
                                                                                                        list_brain_regions,
                                                                                                        list_fmri_scans_indexes)

        # Create the DataFrame and perform the Wilcoxon signed-rank test. 
        df, _, _ = create_df_foundation_models(dict_targets, dict_predictions, brain_region_indexes, fmri_scans_indexes)
        statistics_scipy, statistics_pingouin = wilcoxon_test(df, 'accuracy')

        # Store the statistics in the summary DataFrame. 
        df_summary = store_statistics_in_summary_df(df_summary, model_names, model_index, statistics_scipy, statistics_pingouin, 'accuracy')
        
    # Return the summary DataFrame. 
    return df_summary

# %% [code]
# Run the Wilcoxon signed-rank tests for all models. 
df_summary_foundation_models = run_wilcoxon_tests_foundation_models(model_names, 
                                                                    dict_targets_foundation_models, 
                                                                    list_dict_predictions, 
                                                                    list_brain_regions,
                                                                    list_fmri_scans_indexes)
df_summary_foundation_models

# %% [code]
# Store the FLOPs in a DataFrame. 
df_flops_foundation_models = pd.DataFrame({
    'FLOPs for classification task': {
        'Gemma': dict_flops_large_language_models['Large Language Models: Gemma'],
        'Llama': dict_flops_large_language_models['Large Language Models: Llama'],
        'Gemma Chain-of-Thought (step 1)': dict_flops_large_language_model_CoT['Large Language Model Chain-of-Thought: EEG to Cognitive Functions'], 
        'Gemma Chain-of-Thought (step 2)': dict_flops_large_language_model_CoT['Large Language Model Chain-of-Thought: Cognitive Functions to fMRI'], 
        'Gemma Chain-of-Thought (both steps)': dict_flops_large_language_model_CoT['Large Language Model Chain-of-Thought: EEG to Cognitive Functions'] 
        + dict_flops_large_language_model_CoT['Large Language Model Chain-of-Thought: Cognitive Functions to fMRI'],
        'Gemma Without Fine-Tuning': dict_flops_large_language_model_fine_tuning['Large Language Model Fine-Tuning: Without Fine-Tuning'],
        'Gemma With Fine-Tuning': dict_flops_large_language_model_fine_tuning['Large Language Model Fine-Tuning: With Fine-Tuning'],
        'PaliGemma': dict_flops_large_multimodal_model['Large Multimodal Model']
    }
})

# Display the DataFrame. 
df_flops_foundation_models

# %% [code]
# Retrieve the predictions, brain region indexes, and fMRI scans indexes for the Gemma model without fine-tuning. 
model_index = 3
dict_predictions, brain_region_indexes, fmri_scans_indexes = retrieve_foundation_model_elements(model_index, 
                                                                                                list_dict_predictions,
                                                                                                list_brain_regions,
                                                                                                list_fmri_scans_indexes)

# Create the DataFrame for Gemma without fine-tuning. 
df_gemma_without_finetuning, _, _ = create_df_foundation_models(dict_targets_foundation_models, 
                                                                dict_predictions, 
                                                                brain_region_indexes, 
                                                                fmri_scans_indexes)

# Retrieve the predictions, brain region indexes, and fMRI scans indexes for the Gemma model with fine-tuning. 
model_index = 4
dict_predictions, brain_region_indexes, fmri_scans_indexes = retrieve_foundation_model_elements(model_index, 
                                                                                                list_dict_predictions,
                                                                                                list_brain_regions,
                                                                                                list_fmri_scans_indexes)

# Create the DataFrame for Gemma with fine-tuning. 
df_gemma_with_finetuning, _, _ = create_df_foundation_models(dict_targets_foundation_models, 
                                                             dict_predictions, 
                                                             brain_region_indexes, 
                                                             fmri_scans_indexes)

# Use the model accuracy without fine-tuning as the baseline for the fine-tuned model. 
df_gemma_with_finetuning['baseline_accuracy'] = df_gemma_without_finetuning['model_accuracy']
df_gemma_with_finetuning.head()

# %% [code]
# Perform Wilcoxon signed-rank test. 
statistics_scipy_gemma_with_finetuning, statistics_pingouin_gemma_with_finetuning = wilcoxon_test(df_gemma_with_finetuning, 'accuracy')

# %% [code]
# Create a summary DataFrame to store the statistics. 
df_summary_wilcoxon_finetuning = create_summary_df(['Gemma Without vs. With Fine-Tuning'], 'accuracy')

# Store the statistics in the summary DataFrame. 
df_summary_wilcoxon_finetuning = store_statistics_in_summary_df(df_summary_wilcoxon_finetuning, 
                                                        ['Gemma Without vs. With Fine-Tuning'], 
                                                        0, 
                                                        statistics_scipy_gemma_with_finetuning, 
                                                        statistics_pingouin_gemma_with_finetuning, 
                                                        'accuracy')
df_summary_wilcoxon_finetuning

# %% [markdown]
## **4. Foundation Models (McNemar’s Tests)**

We perform **McNemar’s tests** on the results of the foundation models. 

# %% [code]
# Define a function to perform a McNemar's test on a series of models. 
def run_mcnemars_tests_foundation_models(model_names, dict_targets, list_dict_predictions, list_brain_regions, list_fmri_scans_indexes):
    
    # Create a summary DataFrame and a dictionary to store the statistics for all models. 
    df_summary = pd.DataFrame(columns = ['N', 
                                         'model accuracy', 
                                         'baseline accuracy', 
                                         'combined statistic', 
                                         'combined p-value', 
                                         'median p-value', 
                                         'proportion p-value < 0.05'])
    dict_mcnemars_p_values = dict()

    # Iterate through all models. 
    for model_index in range(len(model_names)):

        # Retrieve the predictions, brain region indexes, and fMRI scans indexes. 
        dict_predictions, brain_region_indexes, fmri_scans_indexes = retrieve_foundation_model_elements(model_index, 
                                                                                                        list_dict_predictions,
                                                                                                        list_brain_regions,
                                                                                                        list_fmri_scans_indexes)

        # Retrieve the lists of targets and predictions. 
        _, list_targets, list_predictions = create_df_foundation_models(dict_targets, dict_predictions, brain_region_indexes, fmri_scans_indexes)

        # Flatten the lists of targets and predictions. 
        targets_values = np.array(list_targets).flatten()
        predictions_values = np.array(list_predictions).flatten()

        # Remove the cases where the prediction is -1. 
        selected_predictions = predictions_values != -1
        targets_values = targets_values[selected_predictions]
        predictions_values = predictions_values[selected_predictions]

        # Compute the accuracy. 
        model_score = np.mean(targets_values == predictions_values)

        # Compute the targets distribution. 
        values = [0, 1]
        targets_distribution = [np.sum(targets_values == 0) / len(targets_values), 
                                np.sum(targets_values == 1) / len(targets_values)]

        # Define the number of iterations and create a list to store the p-values. 
        iterations = 10000
        list_p_values = []
        baseline_score = 0

        # Iterate through the number of iterations. 
        for i in range(iterations):

            # Compute the baseline score using random sampling. 
            samples = np.random.choice(values, size = len(targets_values), p = targets_distribution)
            baseline_score += np.mean(targets_values == samples)

            # Determine the correctness of each model. 
            model_correct = targets_values == predictions_values
            baseline_correct = targets_values == samples

            # Create the 2x2 contingency table. 
            #                baseline correct   baseline wrong
            # model correct        [a]               [b]
            # model wrong          [c]               [d]
            a = np.sum(model_correct & baseline_correct)
            b = np.sum(model_correct & ~baseline_correct)
            c = np.sum(~model_correct & baseline_correct)
            d = np.sum(~model_correct & ~baseline_correct)
            contingency_table = [[a, b],
                                 [c, d]]

            # Perform McNemar's test and store the p-value. 
            result = mcnemar(contingency_table)
            list_p_values.append(result.pvalue)

        # Compute the mean baseline accuracy. 
        baseline_score /= iterations
        
        # Combine the p-values from multiple McNemar iterations. 
        combined_statistic, combined_pvalue = scipy.stats.combine_pvalues(list_p_values, method = 'fisher')

        # Find the median p-value, and the proportion of iterations with a p-value < 0.05. 
        median_pvalue = np.median(list_p_values)
        percentage_significant = np.sum(np.array(list_p_values) < 0.05) / iterations
        
        # Store the statistics in the summary DataFrame and the dictionary. 
        df_summary.loc[model_names[model_index], 'N'] = len(targets_values)
        df_summary.loc[model_names[model_index], 'model accuracy'] = model_score
        df_summary.loc[model_names[model_index], 'baseline accuracy'] = baseline_score
        df_summary.loc[model_names[model_index], 'combined statistic'] = combined_statistic
        df_summary.loc[model_names[model_index], 'combined p-value'] = combined_pvalue
        df_summary.loc[model_names[model_index], 'median p-value'] = median_pvalue
        df_summary.loc[model_names[model_index], 'proportion p-value < 0.05'] = percentage_significant
        dict_mcnemars_p_values[model_names[model_index]] = list_p_values
        
    # Return the summary DataFrame and the dictionary. 
    return df_summary, dict_mcnemars_p_values

# %% [code]
# Run McNemar's tests for all models. 
df_summary_mcnemars, dict_mcnemars_p_values = run_mcnemars_tests_foundation_models(model_names, 
                                                                                   dict_targets_foundation_models, 
                                                                                   list_dict_predictions, 
                                                                                   list_brain_regions,
                                                                                   list_fmri_scans_indexes)

# Display the summary DataFrame. 
df_summary_mcnemars

# %% [code]
# Plot the distribution of p-values for all models. 
fig, axes = plt.subplots(3, 2, figsize = (8, 8))
fig.suptitle('Distribution of p-values for all models', fontsize = 14)

# Iterate through all models. 
for model_index, model_name in enumerate(model_names):
    ax = axes[model_index // 2, model_index % 2]
    ax.hist(dict_mcnemars_p_values[model_name], bins = 40)
    ax.set_title(model_name)
    ax.set_xlabel('p-value')
    ax.set_ylabel('Frequency')

# Adjust the layout of the subplots. 
plt.tight_layout()
plt.subplots_adjust(top = 0.9)

# %% [code]
# Define a function to retrieve fine-tuning vs. no fine-tuning elements. 
def retrieve_finetuning_vs_no_finetuning_elements(model_index, list_dict_predictions, list_brain_regions, list_fmri_scans_indexes):
    
    # Retrieve the predictions, brain region indexes, and fMRI scans indexes. 
    dict_predictions, brain_region_indexes, fmri_scans_indexes = retrieve_foundation_model_elements(model_index, 
                                                                                                    list_dict_predictions,
                                                                                                    list_brain_regions,
                                                                                                    list_fmri_scans_indexes)
    
    # Retrieve the lists of targets and predictions. 
    _, list_targets, list_predictions = create_df_foundation_models(dict_targets_foundation_models, dict_predictions, brain_region_indexes, fmri_scans_indexes)

    # Flatten the lists of targets and predictions. 
    targets_values = np.array(list_targets).flatten()
    predictions_values = np.array(list_predictions).flatten()

    # Detect the cases where the prediction is not equal to -1. 
    selected_predictions = predictions_values != -1

    # Return the targets values, predictions values, and the selected predictions. 
    return targets_values, predictions_values, selected_predictions

# %% [code]
# Return the targets values, predictions values, and the selected predictions for fine-tuning vs. no fine-tuning. 
targets_values, predictions_values_no_finetuning, selected_predictions_no_finetuning = retrieve_finetuning_vs_no_finetuning_elements(3,
                                                                                                                                     list_dict_predictions,
                                                                                                                                     list_brain_regions,
                                                                                                                                     list_fmri_scans_indexes)
targets_values, predictions_values_finetuning, selected_predictions_finetuning = retrieve_finetuning_vs_no_finetuning_elements(4,
                                                                                                                               list_dict_predictions,
                                                                                                                               list_brain_regions,
                                                                                                                               list_fmri_scans_indexes)

# Remove the cases where the prediction is -1 for either one of the two models. 
selected_predictions = selected_predictions_no_finetuning * selected_predictions_finetuning
targets_values = targets_values[selected_predictions]
predictions_values_no_finetuning = predictions_values_no_finetuning[selected_predictions]
predictions_values_finetuning = predictions_values_finetuning[selected_predictions]

# Determine the correctness of each model. 
no_finetuning_correct = predictions_values_no_finetuning == targets_values
finetuning_correct = predictions_values_finetuning == targets_values

# Create the 2x2 contingency table. 
a = np.sum(finetuning_correct & no_finetuning_correct)
b = np.sum(finetuning_correct & ~no_finetuning_correct)
c = np.sum(~finetuning_correct & no_finetuning_correct)
d = np.sum(~finetuning_correct & ~no_finetuning_correct)
contingency_table = [[a, b],
                     [c, d]]

# Perform McNemar's test. 
result = mcnemar(contingency_table)

# Store the results in a DataFrame. 
df_summary_mcnemar_finetuning = pd.DataFrame(columns = ['N', 
                                                 'accuracy with fine-tuning', 
                                                 'accuracy without fine-tuning', 
                                                 'p-value'])
df_summary_mcnemar_finetuning.loc['Gemma Without vs. With Fine-Tuning', 'N'] = len(targets_values)
df_summary_mcnemar_finetuning.loc['Gemma Without vs. With Fine-Tuning', 'accuracy with fine-tuning'] = np.mean(targets_values == predictions_values_finetuning)
df_summary_mcnemar_finetuning.loc['Gemma Without vs. With Fine-Tuning', 'accuracy without fine-tuning'] = np.mean(targets_values == predictions_values_no_finetuning)
df_summary_mcnemar_finetuning.loc['Gemma Without vs. With Fine-Tuning', 'p-value'] = result.pvalue

# Display the summary DataFrame. 
df_summary_mcnemar_finetuning

# %% [code]
# Store the general results in a dictionary. 
general_results = dict()
general_results['df_summary_ml_class'] = df_summary_ml_class
general_results['df_summary_ml_reg'] = df_summary_ml_reg
general_results['df_summary_dl_class'] = df_summary_dl_class
general_results['df_summary_dl_reg'] = df_summary_dl_reg
general_results['df_summary_foundation_models'] = df_summary_foundation_models
general_results['df_summary_wilcoxon_finetuning'] = df_summary_wilcoxon_finetuning
general_results['df_summary_mcnemars'] = df_summary_mcnemars
general_results['df_summary_mcnemar_finetuning'] = df_summary_mcnemar_finetuning
general_results['dict_mcnemars_p_values'] = dict_mcnemars_p_values
general_results['df_flops_deep_learning'] = df_flops_deep_learning
general_results['df_flops_foundation_models'] = df_flops_foundation_models

# Save the general results into a Pickle file. 
with open(results_path + 'general_results.p', 'wb') as file:
    pickle.dump(general_results, file)
```
