## Supplementary material for "Neural Networks and Foundation Models: Two Strategies for EEG-to-fMRI Prediction": 14_Tables_and_Figures.ipynb

```
# %% [markdown]
# **Neural Networks and Foundation Models: Two Strategies for EEG-to-fMRI Prediction**
### **Supplementary Material**

# %% [markdown]
This project includes fourteen Notebooks:

*01: fMRI Preprocessing*

*02: EEG Preprocessing*

*03: Classification Models*

*04: Regression Models*

*05: Neural Networks*

*06: Convolutional Neural Networks*

*07: Recurrent Neural Networks*

*08: Transformers*

*09: Large Language Models*

*10: Large Language Model Chain-of-Thought*

*11: Large Language Model Fine-Tuning*

*12: Large Multimodal Model*

*13: Statistical Tests*

***14: Tables and Figures***

# %% [markdown]
### Notebook 14

# **Tables and Figures**

# %% [markdown]
## **1. Tables**

# %% [code]
import numpy as np
import pandas as pd
import matplotlib.pyplot as plt
%matplotlib inline
import pickle

# %% [code]
# Define the results path. 
results_path = '../results/'

# %% [code]
# Load the general results. 
with open(results_path + 'general_results.p', 'rb') as file:
    general_results = pickle.load(file)

# Store the results into variables. 
df_summary_ml_class = general_results['df_summary_ml_class']
df_summary_ml_reg = general_results['df_summary_ml_reg']
df_summary_dl_class = general_results['df_summary_dl_class']
df_summary_dl_reg = general_results['df_summary_dl_reg']
df_summary_foundation_models = general_results['df_summary_foundation_models']
df_summary_wilcoxon_finetuning = general_results['df_summary_wilcoxon_finetuning']
df_summary_mcnemars = general_results['df_summary_mcnemars']
df_summary_mcnemar_finetuning = general_results['df_summary_mcnemar_finetuning']
dict_mcnemars_p_values = general_results['dict_mcnemars_p_values']
df_flops_deep_learning = general_results['df_flops_deep_learning']
df_flops_foundation_models = general_results['df_flops_foundation_models']

# %% [code]
# Define a function to format the summary DataFrame. 
def format_df_summary(df_summary, score_type):

    # Copy the DataFrame. 
    df_summary_table = df_summary.copy()
    
    # Round values to 3 decimal places. 
    df_summary_table['mean model ' + score_type] = df_summary_table['mean model ' + score_type].astype(float).round(3)
    df_summary_table['mean baseline ' + score_type] = df_summary_table['mean baseline ' + score_type].astype(float).round(3)
    df_summary_table['STD model ' + score_type] = df_summary_table['STD model ' + score_type].astype(float).round(3)
    df_summary_table['STD baseline ' + score_type] = df_summary_table['STD baseline ' + score_type].astype(float).round(3)
    df_summary_table['CI (lower)'] = df_summary_table['CI (lower)'].astype(float).round(3)
    df_summary_table['CI (upper)'] = df_summary_table['CI (upper)'].astype(float).round(3)
    df_summary_table['CLES'] = df_summary_table['CLES'].astype(float).round(3)

    # Create new columns. 
    df_summary_table['model ' + score_type] = df_summary_table['mean model ' + score_type].astype(str) + ' ± ' + df_summary_table['STD model ' + score_type].astype(str)
    df_summary_table['baseline ' + score_type] = df_summary_table['mean baseline ' + score_type].astype(str) + ' ± ' + df_summary_table['STD baseline ' + score_type].astype(str)
    df_summary_table['confidence interval'] = df_summary_table['CI (lower)'].astype(str) + ' - ' + df_summary_table['CI (upper)'].astype(str)

    # Display only the relevant columns.
    df_summary_table = df_summary_table[['N', 'model ' + score_type, 'baseline ' + score_type, 'p-value', 'confidence interval', 'CLES']]
    
    return df_summary_table

# %% [code]
# Concatenate the machine learning, deep learning, and foundation models results for the classification task. 
df_summary_class = pd.concat([df_summary_ml_class, df_summary_dl_class, df_summary_foundation_models], axis = 0)

# Format the DataFrame. 
df_summary_class_table = format_df_summary(df_summary_class, 'accuracy')

# Display the DataFrame. 
print('Classification Task')
df_summary_class_table

# %% [code]
# Concatenate the machine learning and deep learning results for the regression task. 
df_summary_reg = pd.concat([df_summary_ml_reg, df_summary_dl_reg], axis = 0)

# Format the DataFrame. 
df_summary_reg_table = format_df_summary(df_summary_reg, 'MAE')
df_summary_reg_table = df_summary_reg_table.drop(index = ['Ridge Regression', 
                                                          'Lasso Regression'])

# Display the DataFrame. 
print('Regression Task')
df_summary_reg_table

# %% [code]
# Format the DataFrame for the result of the Wilcoxon test for Gemma Without vs. With Fine-Tuning. 
df_summary_wilcoxon_finetuning_table = df_summary_wilcoxon_finetuning.copy()
df_summary_wilcoxon_finetuning_table = format_df_summary(df_summary_wilcoxon_finetuning_table, 'accuracy')

# Display the DataFrame. 
print('Wilcoxon: Gemma Without vs. With Fine-Tuning')
df_summary_wilcoxon_finetuning_table

# %% [code]
# Format the DataFrame for the results of the McNemar's tests. 
df_summary_mcnemars_table = df_summary_mcnemars.copy()

# Round values to 3 decimal places and format the p-values in scientific notation. 
df_summary_mcnemars_table['model accuracy'] = df_summary_mcnemars_table['model accuracy'].astype(float).round(3)
df_summary_mcnemars_table['baseline accuracy'] = df_summary_mcnemars_table['baseline accuracy'].astype(float).round(3)
df_summary_mcnemars_table['combined p-value'] = df_summary_mcnemars_table['combined p-value'].astype(float).round(3).apply(lambda x: f'{x:.2e}')
df_summary_mcnemars_table['median p-value'] = df_summary_mcnemars_table['median p-value'].astype(float).round(3).apply(lambda x: f'{x:.2e}')
df_summary_mcnemars_table['proportion p-value < 0.05'] = df_summary_mcnemars_table['proportion p-value < 0.05'].astype(float).round(3)

# Drop the combined statistics column. 
df_summary_mcnemars_table = df_summary_mcnemars_table.drop(columns = ['combined statistic'])

# Display the DataFrame. 
print('Classification Task: McNemar\'s')
df_summary_mcnemars_table

# %% [code]
# Format the DataFrame for the result of the McNemar's test for Gemma Without vs. With Fine-Tuning. 
df_summary_mcnemar_finetuning_table = df_summary_mcnemar_finetuning.copy()

# Rename the columns. 
new_column_names = {
    'accuracy with fine-tuning': 'model accuracy',
    'accuracy without fine-tuning': 'baseline accuracy',
    'p-value': 'p-value'
}
df_summary_mcnemar_finetuning_table = df_summary_mcnemar_finetuning_table.rename(columns = new_column_names)

# Round values to 3 decimal places and format the p-value in scientific notation. 
df_summary_mcnemar_finetuning_table['model accuracy'] = df_summary_mcnemar_finetuning_table['model accuracy'].astype(float).round(3)
df_summary_mcnemar_finetuning_table['baseline accuracy'] = df_summary_mcnemar_finetuning_table['baseline accuracy'].astype(float).round(3)
df_summary_mcnemar_finetuning_table['p-value'] = df_summary_mcnemar_finetuning_table['p-value'].astype(float).round(3).apply(lambda x: f'{x:.2e}')

# Display the DataFrame. 
print('McNemar\'s: Gemma Without vs. With Fine-Tuning')
df_summary_mcnemar_finetuning_table

# %% [code]
# Display the FLOPs for the deep learning models. 
print('FLOPs for Deep Learning Models')
df_flops_deep_learning_table = df_flops_deep_learning.copy()
df_flops_deep_learning_table['FLOPs for classification task'] = df_flops_deep_learning_table['FLOPs for classification task'].apply(lambda x: '{:,.0f}'.format(x))
df_flops_deep_learning_table['FLOPs for regression task'] = df_flops_deep_learning_table['FLOPs for regression task'].apply(lambda x: '{:,.0f}'.format(x))
df_flops_deep_learning_table

# %% [code]
# Display the FLOPs for the foundation models. 
print('FLOPs for Foundation Models')
df_flops_foundation_models_table = df_flops_foundation_models.copy()
df_flops_foundation_models_table['FLOPs for classification task'] = df_flops_foundation_models_table['FLOPs for classification task'].apply(lambda x: '{:,.0f}'.format(x))
df_flops_foundation_models_table

# %% [markdown]
## **2. Figures**

# %% [code]
# Set the style of the plots. 
plt.rcParams.update({'font.family': 'serif'})

# %% [code]
# Define a function to convert p-values to stars. 
def pvalue_to_stars(p):
    if pd.isna(p):
        return ''
    elif p < 0.001:
        return '***'
    elif p < 0.01:
        return '**'
    elif p < 0.05:
        return '*'
    return ''

# %% [code]
# Define a function to create a bar plot. 
def create_bar_plot(df_summary, score_type, model_abbreviations = []):

    # Determine the task type based on the score type. 
    if score_type == 'accuracy':
        task_type = 'classification'
        height_asterisk = 0.003
    elif score_type == 'MAE':
        task_type = 'regression'
        height_asterisk = 0.012
    
    # Drop the unnecessary results from the DataFrame. 
    df_summary_figure = df_summary.copy()
    if score_type == 'accuracy':
        df_summary_figure = df_summary_figure.drop(index = ['Gemma', 
                                                            'Llama', 
                                                            'Gemma Chain-of-Thought', 
                                                            'Gemma Without Fine-Tuning', 
                                                            'Gemma With Fine-Tuning', 
                                                            'PaliGemma'])
    elif score_type == 'MAE':
        df_summary_figure = df_summary_figure.drop(index = ['Ridge Regression', 
                                                            'Lasso Regression'])
    
    # Compute the standard error (SE) of the mean for the score values. 
    sqrt_N = np.sqrt(pd.to_numeric(df_summary_figure['N']))
    df_summary_figure['SE model ' + score_type] = df_summary_figure['STD model ' + score_type] / sqrt_N
    df_summary_figure['SE baseline ' + score_type] = df_summary_figure['STD baseline ' + score_type] / sqrt_N

    # Define the elements to plot. 
    if model_abbreviations:
        model_names = ['Baseline'] + model_abbreviations
    else:
        model_names = ['Baseline'] + df_summary_figure.index.tolist()
    model_accuracies = [df_summary_figure.iloc[0]['mean baseline ' + score_type]] + df_summary_figure['mean model ' + score_type].tolist()
    model_standard_errors = [df_summary_figure.iloc[0]['SE baseline ' + score_type]] + df_summary_figure['SE model ' + score_type].tolist()
    model_colors = ['green'] + ['blue'] * 6 + ['darkblue'] * 4

    # Retrieve the number of stars corresponding to the p-values. 
    stars = [''] + [pvalue_to_stars(float(p)) for p in df_summary_figure['p-value']]

    # Create the bar plot. 
    fig, ax = plt.subplots(figsize=(8, 4))
    bars = ax.bar(
        model_names,
        model_accuracies,
        yerr = model_standard_errors,
        color = model_colors,
        capsize = 5,
        edgecolor = 'black'
    )

    # Add a horizontal dashed line at the top of the baseline. 
    baseline_top = model_accuracies[0]
    ax.axhline(y = baseline_top, color = 'green', linestyle = '--', linewidth = 1, zorder = 0)

    # Add significance stars above each bar. 
    for bar, star in zip(bars, stars):
        if star:
            height = bar.get_height()
            ax.text(bar.get_x() + bar.get_width() / 2, height + height_asterisk, star,
                    ha = 'center', va = 'bottom', fontsize = 12)

    # Format the axes and adjust the layout. 
    if score_type == 'accuracy':
        ax.set_ylim(0.45, 0.55)
        ax.set_ylabel('mean accuracy', fontsize = 12)
    elif score_type == 'MAE':
        ax.set_ylim(0.6, 0.95)
        ax.set_ylabel('mean MAE', fontsize = 12)
    #fig.suptitle('Machine Learning and Deep Learning Models: ' + task_type.capitalize(), fontsize = 13)
    plt.xticks(rotation = 45, ha = 'right', fontsize = 11)
    plt.tight_layout()

    # Display and save the figure. 
    plt.show()
    fig.savefig(results_path + 'figure_' + task_type + '.pdf', format = 'pdf', bbox_inches = 'tight')

# %% [code]
# Define model abbreviations. 
model_abbreviations_class = ['Logistic', 'KNN', 'DT', 'RF', 'SVM', 'XGBoost', 'MLP', 'CNN', 'RNN', 'Transformer']
model_abbreviations_reg = ['Linear', 'KNN', 'DT', 'RF', 'SVM', 'XGBoost', 'MLP', 'CNN', 'RNN', 'Transformer']
model_abbreviations_foundation_models = ['Gemma', 'Llama', 'Gemma CoT', 'Gemma without fine-tuning', 'Gemma with fine-tuning', 'PaliGemma']

# %% [code]
# Create the bar plot for the classification task. 
create_bar_plot(df_summary_class, 'accuracy', model_abbreviations_class)

# %% [code]
# Create the bar plot for the regression task. 
create_bar_plot(df_summary_reg, 'MAE', model_abbreviations_reg)

# %% [code]
# Plot the distribution of p-values for all foundation models. 
model_names = df_summary_mcnemars_table.index.tolist()
fig, axes = plt.subplots(2, 3, figsize = (8, 5), sharex = True, sharey = True)
#fig.suptitle('Foundation Models: Distribution of p-values', fontsize = 13)

# Iterate through all foundation models. 
for model_index, model_name in enumerate(model_names):
    ax = axes[model_index // 3, model_index % 3]
    ax.hist(dict_mcnemars_p_values[model_name], bins = 40, color = 'blue')
    #ax.set_title(model_name, fontsize = 12)
    ax.set_title(model_abbreviations_foundation_models[model_index], fontsize = 12)
    ax.set_xlim(-0.03, 1.03)
    ax.set_ylim(0, 1800)
    ax.set_xticks([0, 0.5, 1.0], minor = False)
    ax.set_xticks([0.25, 0.75], minor = True)
    ax.tick_params(axis = 'x', which = 'minor', length = 4)
    ax.set_xticklabels(['0', '0.5', '1.0'], minor = False, fontsize = 10)

    ax.axvline(0.05, linestyle = '--', linewidth = 1, color = 'blue')
    ax.text(0.86, 0.95, f'({chr(97 + model_index)})',
            transform = ax.transAxes,
            va = 'top', ha = 'left', fontsize = 12)

# Display outer labels only. 
axes[1, 1].set_xlabel('p-value', fontsize = 12)
axes[0, 0].set_ylabel('frequency', fontsize = 12)
axes[1, 0].set_ylabel('frequency', fontsize = 12)

# Adjust the layout of the subplots. 
plt.tight_layout()
plt.subplots_adjust(top = 0.88)

# Save the figure. 
fig.savefig(results_path + 'figure_foundation_models.pdf', format = 'pdf', bbox_inches = 'tight')
```
