## Supplementary material for "Neural Networks and Foundation Models: Two Strategies for EEG-to-fMRI Prediction": ENV.yml

```
name: eeg-to-fmri-prediction
channels:
  - conda-forge
  - defaults
dependencies:
  - _py-xgboost-mutex=2.0=cpu_0
  - absl-py=2.1.0=pyhd8ed1ab_1
  - accelerate=1.5.2=pyhd8ed1ab_0
  - aiohappyeyeballs=2.4.6=pyhd8ed1ab_0
  - aiohttp=3.11.12=py311h4921393_0
  - aiosignal=1.3.2=pyhd8ed1ab_0
  - antropy=0.1.9=pyhd8ed1ab_0
  - anyio=4.8.0=pyhd8ed1ab_0
  - aom=3.9.1=h7bae524_0
  - appnope=0.1.4=pyhd8ed1ab_1
  - argon2-cffi=23.1.0=pyhd8ed1ab_1
  - argon2-cffi-bindings=21.2.0=py311h460d6c5_5
  - arrow=1.3.0=pyhd8ed1ab_1
  - asttokens=3.0.0=pyhd8ed1ab_1
  - astunparse=1.6.3=pyhd8ed1ab_3
  - async-lru=2.0.4=pyhd8ed1ab_1
  - attrs=25.1.0=pyh71513ae_0
  - aws-c-auth=0.8.6=h660070d_4
  - aws-c-cal=0.8.7=h8f38403_0
  - aws-c-common=0.12.0=h5505292_0
  - aws-c-compression=0.3.1=hd84a0f8_2
  - aws-c-event-stream=0.5.4=h3c33643_2
  - aws-c-http=0.9.4=hedcc1e3_4
  - aws-c-io=0.17.0=ha705ebb_6
  - aws-c-mqtt=0.12.2=h82c6c6a_2
  - aws-c-s3=0.7.13=hb857f95_2
  - aws-c-sdkutils=0.2.3=hd84a0f8_2
  - aws-checksums=0.2.3=hd84a0f8_2
  - aws-crt-cpp=0.31.0=h7378f02_4
  - aws-sdk-cpp=1.11.510=hf067f9e_3
  - azure-core-cpp=1.14.0=hd50102c_0
  - azure-identity-cpp=1.10.0=hc602bab_0
  - azure-storage-blobs-cpp=12.13.0=h7585a09_1
  - azure-storage-common-cpp=12.8.0=h9ca1f76_1
  - azure-storage-files-datalake-cpp=12.12.0=hcdd55da_1
  - babel=2.17.0=pyhd8ed1ab_0
  - beautifulsoup4=4.13.3=pyha770c72_0
  - bleach=6.2.0=pyh29332c3_4
  - bleach-with-css=6.2.0=h82add2a_4
  - blosc=1.21.6=h7dd00d9_1
  - brotli=1.1.0=hd74edd7_2
  - brotli-bin=1.1.0=hd74edd7_2
  - brotli-python=1.1.0=py311h3f08180_2
  - bzip2=1.0.8=h99b78c6_7
  - c-ares=1.34.4=h5505292_0
  - c-blosc2=2.17.1=h9cbb436_0
  - ca-certificates=2025.1.31=hf0a4a13_0
  - cached-property=1.5.2=hd8ed1ab_1
  - cached_property=1.5.2=pyha770c72_1
  - cairo=1.18.2=h6a3b0d2_1
  - certifi=2025.1.31=pyhd8ed1ab_0
  - cffi=1.17.1=py311h3a79f62_0
  - charset-normalizer=3.4.1=pyhd8ed1ab_0
  - click=8.1.8=pyh707e725_0
  - cloudpickle=3.1.1=pyhd8ed1ab_0
  - colorama=0.4.6=pyhd8ed1ab_1
  - comm=0.2.2=pyhd8ed1ab_1
  - contourpy=1.3.1=py311h210dab8_0
  - cpython=3.11.11=py311hd8ed1ab_2
  - cycler=0.12.1=pyhd8ed1ab_1
  - cyrus-sasl=2.1.27=h60b93bd_7
  - cytoolz=1.0.1=py311h917b07b_0
  - darkdetect=0.8.0=pyhd8ed1ab_1
  - dask-core=2025.2.0=pyhd8ed1ab_0
  - dataclasses=0.8=pyhc8e2a94_3
  - datasets=3.5.0=pyhd8ed1ab_0
  - dav1d=1.2.1=hb547adb_0
  - dbus=1.13.6=h3818c69_3
  - debugpy=1.8.12=py311h155a34a_0
  - decorator=5.1.1=pyhd8ed1ab_1
  - deepdiff=8.2.0=pyhd8ed1ab_0
  - defusedxml=0.7.1=pyhd8ed1ab_0
  - deprecated=1.2.18=pyhd8ed1ab_0
  - dill=0.3.7=pyhd8ed1ab_0
  - dipy=1.10.0=py311hca32420_1
  - double-conversion=3.3.1=h286801f_0
  - edfio=0.4.5=pyhd8ed1ab_1
  - eeglabio=0.0.3=pyhd8ed1ab_1
  - eigen=3.4.0=h1995070_0
  - exceptiongroup=1.2.2=pyhd8ed1ab_1
  - executing=2.1.0=pyhd8ed1ab_1
  - expat=2.6.4=h286801f_0
  - ffmpeg=7.1.1=gpl_he2be4f0_103
  - filelock=3.17.0=pyhd8ed1ab_0
  - flatbuffers=24.12.23=h28594ff_0
  - font-ttf-dejavu-sans-mono=2.37=hab24e00_0
  - font-ttf-inconsolata=3.000=h77eed37_0
  - font-ttf-source-code-pro=2.038=h77eed37_0
  - font-ttf-ubuntu=0.83=h77eed37_3
  - fontconfig=2.15.0=h1383a14_1
  - fonts-conda-ecosystem=1=0
  - fonts-conda-forge=1=0
  - fonttools=4.56.0=py311h4921393_0
  - fqdn=1.5.1=pyhd8ed1ab_1
  - freetype=2.13.3=h1d14073_0
  - fribidi=1.0.10=h27ca646_0
  - frozenlist=1.5.0=py311h4921393_1
  - fsspec=2024.12.0=pyhd8ed1ab_0
  - fvcore=0.1.5.post20221221=pyhd8ed1ab_0
  - gast=0.6.0=pyhd8ed1ab_0
  - gdk-pixbuf=2.42.12=h7ddc832_0
  - gettext=0.23.1=h3dcc1bd_0
  - gettext-tools=0.23.1=h493aca8_0
  - gflags=2.2.2=hf9b8971_1005
  - giflib=5.2.2=h93a5062_0
  - gl2ps=1.4.2=hc97c1ff_1
  - glew=2.1.0=h9f76cd9_2
  - glib=2.82.2=heee381b_1
  - glib-tools=2.82.2=h1dc7a0c_1
  - glog=0.7.1=heb240a5_0
  - gmp=6.3.0=h7bae524_2
  - gmpy2=2.1.5=py311hb5d9ff4_3
  - google-pasta=0.2.0=pyhd8ed1ab_2
  - graphite2=1.3.13=hebf3989_1003
  - grpcio=1.67.1=py311ha413f7a_2
  - gst-plugins-base=1.24.7=hb49d354_0
  - gstreamer=1.24.7=hc3f5269_0
  - h11=0.14.0=pyhd8ed1ab_1
  - h2=4.2.0=pyhd8ed1ab_0
  - h5io=0.2.4=pyhecae5ae_1
  - h5py=3.13.0=nompi_py311h20052fc_100
  - harfbuzz=10.4.0=hb72c1af_0
  - hdf4=4.2.15=h2ee6834_7
  - hdf5=1.14.3=nompi_ha698983_109
  - hpack=4.1.0=pyhd8ed1ab_0
  - httpcore=1.0.7=pyh29332c3_1
  - httpx=0.28.1=pyhd8ed1ab_0
  - huggingface_hub=0.30.2=pyhd8ed1ab_0
  - hyperframe=6.1.0=pyhd8ed1ab_0
  - icu=75.1=hfee45f7_0
  - idna=3.10=pyhd8ed1ab_1
  - imagecodecs-lite=2019.12.3=py311h5d790af_8
  - imageio=2.37.0=pyhfb79c49_0
  - imageio-ffmpeg=0.6.0=pyhd8ed1ab_0
  - importlib-metadata=8.6.1=pyha770c72_0
  - importlib_resources=6.5.2=pyhd8ed1ab_0
  - iopath=0.1.10=pyhd8ed1ab_0
  - ipykernel=6.29.5=pyh57ce528_0
  - ipython=8.32.0=pyh907856f_0
  - ipywidgets=8.1.5=pyhd8ed1ab_1
  - isoduration=20.11.0=pyhd8ed1ab_1
  - jedi=0.19.2=pyhd8ed1ab_1
  - jinja2=3.1.5=pyhd8ed1ab_0
  - joblib=1.4.2=pyhd8ed1ab_1
  - json5=0.10.0=pyhd8ed1ab_1
  - jsoncpp=1.9.6=h726d253_1
  - jsonpointer=3.0.0=py311h267d04e_1
  - jsonschema=4.23.0=pyhd8ed1ab_1
  - jsonschema-specifications=2024.10.1=pyhd8ed1ab_1
  - jsonschema-with-format-nongpl=4.23.0=hd8ed1ab_1
  - jupyter=1.1.1=pyhd8ed1ab_1
  - jupyter-lsp=2.2.5=pyhd8ed1ab_1
  - jupyter_client=8.6.3=pyhd8ed1ab_1
  - jupyter_console=6.6.3=pyhd8ed1ab_1
  - jupyter_core=5.7.2=pyh31011fe_1
  - jupyter_events=0.12.0=pyh29332c3_0
  - jupyter_server=2.15.0=pyhd8ed1ab_0
  - jupyter_server_terminals=0.5.3=pyhd8ed1ab_1
  - jupyterlab=4.3.5=pyhd8ed1ab_0
  - jupyterlab_pygments=0.3.0=pyhd8ed1ab_2
  - jupyterlab_server=2.27.3=pyhd8ed1ab_1
  - jupyterlab_widgets=3.0.13=pyhd8ed1ab_1
  - keras=3.8.0=pyh753f3f9_0
  - khronos-opencl-icd-loader=2024.10.24=h5505292_1
  - kiwisolver=1.4.7=py311h2c37856_0
  - krb5=1.21.3=h237132a_0
  - lame=3.100=h1a8c8d9_1003
  - lazy-loader=0.4=pyhd8ed1ab_2
  - lazy_loader=0.4=pyhd8ed1ab_2
  - lcms2=2.17=h7eeda09_0
  - lerc=4.0.0=h9a09cb3_0
  - libabseil=20240722.0=cxx17_h07bc746_4
  - libaec=1.1.3=hebf3989_0
  - libarrow=19.0.1=h9828ad4_3_cpu
  - libarrow-acero=19.0.1=hf07054f_3_cpu
  - libarrow-dataset=19.0.1=hf07054f_3_cpu
  - libarrow-substrait=19.0.1=h4239455_3_cpu
  - libasprintf=0.23.1=h493aca8_0
  - libasprintf-devel=0.23.1=h493aca8_0
  - libass=0.17.3=h16a287c_1
  - libavif16=1.2.1=h3861c80_2
  - libblas=3.9.0=30_h10e41b3_openblas
  - libboost=1.84.0=hc9fb7c5_7
  - libbrotlicommon=1.1.0=hd74edd7_2
  - libbrotlidec=1.1.0=hd74edd7_2
  - libbrotlienc=1.1.0=hd74edd7_2
  - libcblas=3.9.0=30_hb3479ef_openblas
  - libclang-cpp17=17.0.6=default_hf90f093_8
  - libclang-cpp18.1=18.1.8=default_hf90f093_7
  - libclang13=19.1.7=default_h81d93ff_1
  - libcrc32c=1.1.2=hbdafb3b_0
  - libcurl=8.12.1=h73640d1_0
  - libcxx=19.1.7=ha82da77_0
  - libde265=1.0.15=h2ffa867_0
  - libdeflate=1.23=hec38601_0
  - libedit=3.1.20250104=pl5321hafb1f1b_0
  - libev=4.33=h93a5062_2
  - libevent=2.1.12=h2757513_1
  - libexpat=2.6.4=h286801f_0
  - libffi=3.4.2=h3422bc3_5
  - libflac=1.4.3=hb765f3a_0
  - libgettextpo=0.23.1=h493aca8_0
  - libgettextpo-devel=0.23.1=h493aca8_0
  - libgfortran=5.0.0=13_2_0_hd922786_3
  - libgfortran5=13.2.0=hf226fd6_3
  - libglib=2.82.2=hdff4504_1
  - libgoogle-cloud=2.36.0=hdbe95d5_0
  - libgoogle-cloud-storage=2.36.0=h7081f7f_0
  - libgrpc=1.67.1=h0a426d6_2
  - libheif=1.19.7=gpl_h79e6334_100
  - libhwloc=2.11.2=default_hbce5d74_1001
  - libiconv=1.18=hfe07756_0
  - libintl=0.23.1=h493aca8_0
  - libintl-devel=0.23.1=h493aca8_0
  - libjpeg-turbo=3.0.0=hb547adb_1
  - liblapack=3.9.0=30_hc9a63f6_openblas
  - liblightgbm=4.5.0=cpu_h0bc706f_4
  - libllvm15=15.0.7=h4429f82_5
  - libllvm17=17.0.6=hc4b4ae8_3
  - libllvm18=18.1.8=hc4b4ae8_3
  - libllvm19=19.1.7=hc4b4ae8_1
  - liblzma=5.6.4=h39f12f2_0
  - liblzma-devel=5.6.4=h39f12f2_0
  - libmatio=1.5.28=h36f0bf9_2
  - libnetcdf=4.9.2=nompi_h610d594_116
  - libnghttp2=1.64.0=h6d7220d_0
  - libntlm=1.8=h5505292_0
  - libogg=1.3.5=h99b78c6_0
  - libopenblas=0.3.29=openmp_hf332438_0
  - libopentelemetry-cpp=1.18.0=h0c05b2d_1
  - libopentelemetry-cpp-headers=1.18.0=hce30654_1
  - libopenvino=2025.0.0=h3f17238_2
  - libopenvino-arm-cpu-plugin=2025.0.0=h3f17238_2
  - libopenvino-auto-batch-plugin=2025.0.0=h7f72211_2
  - libopenvino-auto-plugin=2025.0.0=h7f72211_2
  - libopenvino-hetero-plugin=2025.0.0=h718ad69_2
  - libopenvino-ir-frontend=2025.0.0=h718ad69_2
  - libopenvino-onnx-frontend=2025.0.0=h76e6831_2
  - libopenvino-paddle-frontend=2025.0.0=h76e6831_2
  - libopenvino-pytorch-frontend=2025.0.0=h286801f_2
  - libopenvino-tensorflow-frontend=2025.0.0=he275e1d_2
  - libopenvino-tensorflow-lite-frontend=2025.0.0=h286801f_2
  - libopus=1.3.1=h27ca646_1
  - libparquet=19.0.1=h636d7b7_3_cpu
  - libpng=1.6.47=h3783ad8_0
  - libpq=17.3=h6896619_0
  - libprotobuf=5.28.3=h3bd63a1_1
  - libre2-11=2024.07.02=h07bc746_2
  - librsvg=2.58.4=h266df6f_2
  - libsndfile=1.2.2=h9739721_1
  - libsodium=1.0.20=h99b78c6_0
  - libsqlite=3.49.1=h3f77e49_2
  - libssh2=1.11.1=h9cc3647_0
  - libtheora=1.1.1=h99b78c6_1006
  - libthrift=0.21.0=h64651cc_0
  - libtiff=4.7.0=h551f018_3
  - libtorch=2.6.0=cpu_generic_hc55bb30_1
  - libusb=1.0.28=h5505292_0
  - libutf8proc=2.10.0=hda25de7_0
  - libuv=1.50.0=h5505292_0
  - libvorbis=1.3.7=h9f76cd9_0
  - libvpx=1.14.1=h7bae524_0
  - libwebp-base=1.5.0=h2471fea_0
  - libxcb=1.17.0=hdb1d25a_0
  - libxgboost=2.1.4=cpu_hf17a1b1_0
  - libxml2=2.13.7=h178c5d8_0
  - libxslt=1.1.39=h223e5b9_0
  - libzip=1.11.2=h1336266_0
  - libzlib=1.3.1=h8359307_2
  - lightgbm=4.5.0=cpu_py_4
  - littleutils=0.2.4=pyhd8ed1ab_1
  - llvm-openmp=19.1.7=hdb05f8b_0
  - llvmlite=0.44.0=py311h55fc170_0
  - locket=1.0.0=pyhd8ed1ab_0
  - loguru=0.7.2=py311h267d04e_2
  - lspopt=1.4.0=pyhd8ed1ab_0
  - lxml=5.3.1=py311h0d6554d_0
  - lz4-c=1.10.0=h286801f_1
  - markdown=3.6=pyhd8ed1ab_0
  - markdown-it-py=3.0.0=pyhd8ed1ab_1
  - markupsafe=3.0.2=py311h4921393_1
  - matplotlib=3.10.0=py311ha1ab1f8_0
  - matplotlib-base=3.10.0=py311h031da69_0
  - matplotlib-inline=0.1.7=pyhd8ed1ab_1
  - mdurl=0.1.2=pyhd8ed1ab_1
  - mffpy=0.10.0=pyhd8ed1ab_1
  - mistune=3.1.1=pyhd8ed1ab_0
  - ml_dtypes=0.4.0=py311h9cb3ce9_2
  - mne=1.9.0=pyqt_h8d421b4_201
  - mne-base=1.9.0=pyh694c41f_201
  - mne-qt-browser=0.6.3=pyh0555025_1
  - more-itertools=10.6.0=pyhd8ed1ab_0
  - mpc=1.3.1=h8f1351a_1
  - mpfr=4.2.1=hb693164_3
  - mpg123=1.32.9=hf642e45_0
  - mpmath=1.3.0=pyhd8ed1ab_1
  - msgpack-python=1.1.0=py311h2c37856_0
  - multidict=6.1.0=py311h30e7462_1
  - multiprocess=0.70.15=py311heffc1b2_1
  - munkres=1.1.4=pyh9f0ad1d_0
  - mysql-common=9.0.1=hd7719f6_4
  - mysql-libs=9.0.1=ha8be5b7_4
  - namex=0.0.8=pyhd8ed1ab_1
  - nbclient=0.10.2=pyhd8ed1ab_0
  - nbconvert-core=7.16.6=pyh29332c3_0
  - nbformat=5.10.4=pyhd8ed1ab_1
  - ncurses=6.5=h5e97a16_3
  - nest-asyncio=1.6.0=pyhd8ed1ab_1
  - networkx=3.4.2=pyh267e887_2
  - nibabel=5.3.2=pyha770c72_1
  - nilearn=0.11.1=pyhd8ed1ab_0
  - nlohmann_json=3.11.3=h00cdb27_1
  - nomkl=1.0=h5ca1d4c_0
  - notebook=7.3.2=pyhd8ed1ab_0
  - notebook-shim=0.2.4=pyhd8ed1ab_1
  - nspr=4.36=h5833ebf_0
  - nss=3.108=ha3c76ea_0
  - numba=0.61.0=py311h74daea0_1
  - numexpr=2.10.2=py311h7469182_0
  - numpy=1.26.4=py311h7125741_0
  - ollama=0.6.5=cpu_hee89a2d_
  - opencl-headers=2024.10.24=h286801f_0
  - openh264=2.6.0=hb5b2745_0
  - openjpeg=2.5.3=h8a3d83b_0
  - openldap=2.6.9=hbe55e7a_0
  - openmeeg=2.5.15=py311h6e48d46_0
  - openssl=3.5.0=h81ee809_0
  - opt_einsum=3.4.0=pyhd8ed1ab_1
  - optree=0.14.0=py311h210dab8_1
  - orc=2.1.1=h74f7c94_0
  - orderly-set=5.3.0=pyh29332c3_0
  - outdated=0.2.2=pyhd8ed1ab_2
  - overrides=7.7.0=pyhd8ed1ab_1
  - packaging=24.2=pyhd8ed1ab_2
  - pandas=2.2.3=py311h9cb3ce9_1
  - pandas-flavor=0.6.0=pyhd8ed1ab_3
  - pandocfilters=1.5.0=pyhd8ed1ab_0
  - pango=1.56.1=h73f1e88_0
  - parso=0.8.4=pyhd8ed1ab_1
  - partd=1.4.2=pyhd8ed1ab_0
  - patsy=1.0.1=pyhd8ed1ab_1
  - pcre2=10.44=h297a79d_2
  - peft=0.15.0=pyhd8ed1ab_0
  - pexpect=4.9.0=pyhd8ed1ab_1
  - pickleshare=0.7.5=pyhd8ed1ab_1004
  - pillow=11.1.0=py311hb9ba9e9_0
  - pingouin=0.5.5=pyhd8ed1ab_1
  - pip=25.0.1=pyh8b19718_0
  - pixman=0.44.2=h2f9eb0b_0
  - pkgutil-resolve-name=1.3.10=pyhd8ed1ab_2
  - platformdirs=4.3.6=pyhd8ed1ab_1
  - ply=3.11=pyhd8ed1ab_3
  - pooch=1.8.2=pyhd8ed1ab_1
  - portalocker=3.0.0=py311h267d04e_0
  - proj=9.5.1=h1318a7e_0
  - prometheus-cpp=1.3.0=h0967b3e_0
  - prometheus_client=0.21.1=pyhd8ed1ab_0
  - prompt-toolkit=3.0.50=pyha770c72_0
  - prompt_toolkit=3.0.50=hd8ed1ab_0
  - propcache=0.2.1=py311h4921393_1
  - protobuf=5.28.3=py311h155a34a_0
  - psutil=6.1.1=py311h917b07b_0
  - pthread-stubs=0.4=hd74edd7_1002
  - ptyprocess=0.7.0=pyhd8ed1ab_1
  - pugixml=1.15=hd3d436d_0
  - pure_eval=0.2.3=pyhd8ed1ab_1
  - py-cpuinfo=9.0.0=pyhd8ed1ab_1
  - py-xgboost=2.1.4=cpu_pyh43b6f25_0
  - pyarrow=19.0.1=py311ha1ab1f8_0
  - pyarrow-core=19.0.1=py311he04fa90_0_cpu
  - pybind11=2.13.6=pyh1ec8472_2
  - pybind11-global=2.13.6=pyh415d2e4_2
  - pybv=0.7.6=pyhd8ed1ab_1
  - pycparser=2.22=pyh29332c3_1
  - pygments=2.19.1=pyhd8ed1ab_0
  - pymatreader=1.0.0=pyhd8ed1ab_1
  - pyobjc-core=11.0=py311hab620ed_0
  - pyobjc-framework-cocoa=11.0=py311hab620ed_0
  - pyopengl=3.1.7=pyhd8ed1ab_0
  - pyparsing=3.2.1=pyhd8ed1ab_0
  - pyqt=5.15.9=py311hc49b008_5
  - pyqt5-sip=12.12.2=py311ha891d26_5
  - pyqtgraph=0.13.7=pyhd8ed1ab_1
  - pyriemann=0.8=pyhd8ed1ab_0
  - pysocks=1.7.1=pyha55dd90_7
  - pysoundfile=0.13.1=pyhd8ed1ab_0
  - pytables=3.10.2=py311hd0690c4_2
  - python=3.11.11=hc22306f_1_cpython
  - python-dateutil=2.9.0.post0=pyhff2d567_1
  - python-fastjsonschema=2.21.1=pyhd8ed1ab_0
  - python-flatbuffers=25.2.10=pyhbc23db3_0
  - python-json-logger=2.0.7=pyhd8ed1ab_0
  - python-picard=0.8=pyha308f57_1
  - python-tzdata=2025.1=pyhd8ed1ab_0
  - python-xxhash=3.5.0=py311h917b07b_2
  - python_abi=3.11=5_cp311
  - pytorch=2.6.0=cpu_generic_py311_h479557c_1
  - pytz=2024.1=pyhd8ed1ab_0
  - pyvista=0.44.2=pyhd8ed1ab_1
  - pyvistaqt=0.11.2=pyhdecd6ff_0
  - pywavelets=1.8.0=py311h18599de_0
  - pyyaml=6.0.2=py311h4921393_2
  - pyzmq=26.2.1=py311h01f2145_0
  - qdarkstyle=3.2.3=pyhd8ed1ab_1
  - qhull=2020.2=h420ef59_5
  - qt-main=5.15.15=h67564f6_2
  - qt6-main=6.8.2=h4464394_0
  - qtpy=2.4.3=pyhd8ed1ab_0
  - rav1e=0.6.6=h69fbcac_2
  - re2=2024.07.02=h6589ca4_2
  - readline=8.2=h92ec313_1
  - referencing=0.36.2=pyh29332c3_0
  - regex=2024.11.6=py311h917b07b_0
  - requests=2.32.3=pyhd8ed1ab_1
  - rfc3339-validator=0.1.4=pyhd8ed1ab_1
  - rfc3986-validator=0.1.1=pyh9f0ad1d_0
  - rich=13.9.4=pyhd8ed1ab_1
  - rpds-py=0.22.3=py311h3ff9189_0
  - safetensors=0.5.3=py311h3ff9189_0
  - scikit-image=0.20.0=py311h4eec4a9_1
  - scikit-learn=1.6.1=py311h47fa2fb_0
  - scipy=1.15.2=py311h0675101_0
  - scooby=0.10.0=pyhd8ed1ab_1
  - sdl2=2.32.54=h994913f_0
  - sdl3=3.2.10=he842692_0
  - seaborn=0.13.2=hd8ed1ab_3
  - seaborn-base=0.13.2=pyhd8ed1ab_3
  - send2trash=1.8.3=pyh31c8845_1
  - setuptools=75.8.0=pyhff2d567_0
  - setuptools-scm=8.1.0=pyhd8ed1ab_1
  - sip=6.8.6=py311h3f08180_1
  - six=1.17.0=pyhd8ed1ab_0
  - sleef=3.8=h8391f65_0
  - sleepecg=0.5.9=py311h18599de_2
  - snappy=1.2.1=h98b9ce2_1
  - sniffio=1.3.1=pyhd8ed1ab_1
  - soupsieve=2.5=pyhd8ed1ab_1
  - sqlite=3.49.1=hd7222ec_2
  - stack_data=0.6.3=pyhd8ed1ab_1
  - statsmodels=0.14.4=py311h0f07fe1_0
  - svt-av1=3.0.2=h8ab69cd_0
  - sympy=1.13.3=pyh2585a3b_105
  - tabulate=0.9.0=pyhd8ed1ab_2
  - tbb=2022.0.0=h0cbf7ec_0
  - tbb-devel=2022.0.0=h6e261d1_0
  - tensorboard=2.18.0=pyhd8ed1ab_1
  - tensorboard-data-server=0.7.0=py311h544b224_2
  - tensorflow=2.18.0=cpu_py311h9d3d1e9_0
  - tensorflow-base=2.18.0=cpu_py311h66b8b4e_0
  - tensorflow-estimator=2.18.0=cpu_py311hfc28eff_0
  - tensorpac=0.6.5=pyhecae5ae_1
  - termcolor=2.5.0=pyhd8ed1ab_1
  - terminado=0.18.1=pyh31c8845_0
  - tf-keras=2.18.0=pyhd8ed1ab_0
  - threadpoolctl=3.5.0=pyhc1e730c_0
  - tifffile=2020.6.3=py_0
  - tinycss2=1.4.0=pyhd8ed1ab_0
  - tk=8.6.13=h5083fa2_1
  - tokenizers=0.21.0=py311h82b0fb8_0
  - toml=0.10.2=pyhd8ed1ab_1
  - tomli=2.2.1=pyhd8ed1ab_1
  - toolz=1.0.0=pyhd8ed1ab_1
  - torchvision=0.21.0=cpu_py311_hccac0c2_1
  - torchvision-extra-decoders=0.0.2=py311h4c72eb5_2
  - tornado=6.4.2=py311h917b07b_0
  - tqdm=4.67.1=pyhd8ed1ab_1
  - traitlets=5.14.3=pyhd8ed1ab_1
  - trame=3.8.0=pyhd8ed1ab_0
  - trame-client=3.5.2=pyhd8ed1ab_0
  - trame-server=3.3.0=pyhd8ed1ab_0
  - trame-vtk=2.8.15=pyhb419c8b_0
  - trame-vuetify=2.8.1=pyhd8ed1ab_0
  - transformers=4.51.3=pyhd8ed1ab_0
  - trx-python=0.3=py311h267d04e_1
  - types-python-dateutil=2.9.0.20241206=pyhd8ed1ab_0
  - typing-extensions=4.12.2=hd8ed1ab_1
  - typing_extensions=4.12.2=pyha770c72_1
  - typing_utils=0.1.0=pyhd8ed1ab_1
  - tzdata=2025a=h78e105d_0
  - unicodedata2=16.0.0=py311h917b07b_0
  - uri-template=1.3.0=pyhd8ed1ab_1
  - urllib3=2.3.0=pyhd8ed1ab_0
  - utfcpp=4.0.6=h54c0426_0
  - vtk=9.3.1=qt_py311he4b582b_216
  - vtk-base=9.3.1=qt_py311hb64ca2f_216
  - vtk-io-ffmpeg=9.3.1=qt_py311he4b582b_216
  - wcwidth=0.2.13=pyhd8ed1ab_1
  - webcolors=24.11.1=pyhd8ed1ab_0
  - webencodings=0.5.1=pyhd8ed1ab_3
  - websocket-client=1.8.0=pyhd8ed1ab_1
  - werkzeug=3.1.3=pyhd8ed1ab_1
  - wfdb=4.2.0=pyhd8ed1ab_1
  - wheel=0.45.1=pyhd8ed1ab_1
  - widgetsnbextension=4.0.13=pyhd8ed1ab_1
  - wrapt=1.17.2=py311h917b07b_0
  - wslink=2.2.2=pyhd8ed1ab_0
  - x264=1!164.3095=h57fd34a_2
  - x265=3.5=hbc6ce65_3
  - xarray=2025.3.1=pyhd8ed1ab_0
  - xgboost=2.1.4=cpu_pyhc1a9e11_0
  - xlrd=2.0.1=pyhd8ed1ab_3
  - xmltodict=0.14.2=pyhd8ed1ab_1
  - xorg-libxau=1.0.12=h5505292_0
  - xorg-libxdmcp=1.1.5=hd74edd7_0
  - xxhash=0.8.3=h5505292_0
  - xz=5.6.4=h9a6d368_0
  - xz-gpl-tools=5.6.4=h9a6d368_0
  - xz-tools=5.6.4=h39f12f2_0
  - yacs=0.1.8=pyh29332c3_2
  - yaml=0.2.5=h3422bc3_2
  - yarl=1.18.3=py311h4921393_1
  - yasa=0.6.5=pyhd8ed1ab_2
  - zeromq=4.3.5=hc1bb282_7
  - zipp=3.21.0=pyhd8ed1ab_1
  - zlib=1.3.1=h8359307_2
  - zlib-ng=2.2.4=h13dfb9a_0
  - zstandard=0.23.0=py311h917b07b_1
  - zstd=1.5.7=h6491c7d_2
prefix: /Users/username/anaconda3/envs/eeg-to-fmri-prediction
```
