## Supplementary material for "Neural Networks and Foundation Models: Two Strategies for EEG-to-fMRI Prediction": README.md

# **Neural Networks and Foundation Models: Two Strategies for EEG-to-fMRI Prediction**

### **Supplementary Material**

***General Note:*** The content of this README file is mostly a less formal, code-oriented description of the experiments reported in the submitted article. For the description of the fMRIPrep preprocessing, go directly to the Data section. For the list of Jupyter Notebooks and a short description of their content, go directly to the Methods section. 

### Abstract

**In this project, we attempt the development of an artificial intelligence system for human brain activity prediction. Our objective is to predict fMRI activity from EEG activity, an endeavor that can be formulated as either a classification task, i.e., predicting whether the fMRI signal increases or decreases, or a regression task, i.e., predicting the value of this signal. To achieve this objective, we follow two distinct strategies: training machine learning and deep learning models (including feedforward neural networks, CNNs, RNNs, and transformers) on an EEG-fMRI dataset, or relying on the capabilities of pre-trained large language models (Gemma, Llama) and large multimodal models (PaliGemma). Our results show that predicting fMRI activity from EEG activity is possible for the brain regions defined by the Harvard-Oxford cortical atlas, even in the challenging context of subjects performing a cognitive task such as neurofeedback. Interestingly, both strategies yield promising results, possibly highlighting two complementary paths for predicting fMRI signals based on EEG signals. Furthermore, a Chain-of-Thought approach demonstrates that large language models can infer the cognitive functions associated with EEG data, and subsequently the fMRI data associated with these cognitive functions. The natural combination of both strategies, i.e., fine-tuning a large language model on an EEG-fMRI dataset, is not straightforward and would certainly require further study. Overall, the methods developed in this project could represent an important step for improving neural interfaces and advancing toward a multimodal foundation model for neuroscience.**

### Introduction

Electroencephalography (EEG) and functional Magnetic Resonance Imaging (fMRI) are two of the major tools used by neuroscientists for human brain activity investigation. While EEG measures the electrical activity of the brain, using electrodes placed on the head, fMRI, on the other hand, measures the activity of the brain indirectly, by detecting the changes in the cerebral blood flow. EEG is typically used to record activity of cortical areas, while fMRI can detect activity in subcortical regions as well. EEG has a good temporal resolution, while fMRI has a good spatial resolution. In EEG-fMRI research, both techniques are used simultaneously, allowing the researchers to obtain two complementary datasets, acquired at the same time on the same subjects. These studies are often particularly insightful, since they provide two different perspectives on human brain activity for the task of interest. 

Most of the time, neuroimaging experiments reported in the scientific literature use only one neuroimaging technique, e.g., EEG or fMRI. Multimodal experiments such as EEG-fMRI studies are rarer, but there is a growing interest in these approaches, and some EEG-fMRI datasets are already accessible in open data repositories. While the human brain can be explored through many different and complementary angles, all neuroimaging techniques, whether they measure the electrical activity on the surface of the head, or the changes in the cerebral blood flow, are ultimately grounded in the same neurophysiological reality. Therefore, a natural research direction is to explore the degree of redundancy, or shared variance, between two types of neuroimaging data acquired at the same time on the same subjects. Specifically, in this project, we train machine learning and deep learning models to make predictions about human brain activity in a particular EEG-fMRI dataset, while also evaluating the predictive capabilities of two pre-trained large language models (LLMs) and one pre-trained large multimodal model (LMM). 

If our general objective is to build an artificial intelligence (AI) system for human brain activity prediction, the most interesting problem, but also the most challenging, is certainly to use EEG data to predict fMRI data. Of course, solving the reverse problem, i.e., to use fMRI data to predict EEG data, could also be attempted. Nevertheless, while fMRI can be used to investigate very specific cognitive processes, this neuroimaging technique is particularly complex and expensive, and requires the subject to remain immobile inside the scanner. By contrast, EEG is a much simpler and more affordable neuroimaging technique, and is better adapted to real-life measurements in a diversity of situations. Hypothetically, if we could achieve near-fMRI precision using only EEG devices, we might be able to obtain the best of both worlds, and this could open entirely new horizons for neuroscience and neurotechnology. Specifically, our focus should be on the fluctuations of the fMRI signal during an experimental condition, since these fluctuations, rather than the absolute value of the signal, can provide important insights into a multiplicity of cognitive processes. 

In this project, we introduce two important modifications compared to more classical fMRI studies. First, instead of focusing on particular voxels, or on small clusters located around these voxels, we rely on the brain regions defined in the Harvard-Oxford cortical atlas, which are both much larger and highly relevant in anatomical and functional terms. Second, instead of focusing only on the regression task, i.e., predicting the value of the fluctuations of the fMRI signal, we add a classification task, i.e., predicting whether the fMRI signal increases or decreases from one scan to the next. Critically, these two modifications allow us to use LLMs and LMMs for the classification task, since we can now rely on brain region keywords, e.g., "Frontal Pole", and fMRI signal evolution keywords, i.e., "Increasing" or "Decreasing". By reframing the prediction problem, we unlock the possibility to leverage large pre-trained models, which would have been much more difficult to do in the initial setting. 

One situation where the human brain might engage in decision-making, learning, reasoning, planning, and strategic thinking, is during a neurofeedback (NF) session. NF consists in providing real-time information to a subject about his/her own brain activity, and asking this subject to try to adapt his/her behavior according to this measure. The objective is to reach a certain brain state, typically by increasing the power of a specific frequency band, in EEG NF, or the activity in a specific brain region, in fMRI NF. Therefore, NF is basically a brain training technology, allowing the subject to learn self-regulation. The brain activity can be measured through a variety of techniques, but EEG and fMRI are the two main choices, and while most NF training protocols rely on a single technique, recent research has shown the potential of EEG-fMRI NF protocols combining both approaches. Furthermore, it should also be noted that NF is one of the technologies that might benefit the most from an AI system for human brain activity prediction. Indeed, if we could use EEG data to predict fMRI data, we might be able to expand the diversity of feedbacks provided to the subjects, while retaining the simplicity and affordability of EEG devices. 

### Data

In this project, we use an EEG-fMRI NF dataset downloaded from OpenNeuro (https://openneuro.org/), an open data repository for neuroimaging data, where researchers can publicly store and share brain files obtained from several neuroimaging techniques, including EEG and fMRI. All brain files are stored in the Brain Imaging Data Structure (BIDS) format, a standard format for neuroimaging and behavioral data (https://bids.neuroimaging.io/). Furthermore, all newly published datasets in OpenNeuro, including the one we selected, are released under CC0 license (https://openneuro.org/faq), which means a public domain dedication and no copyright (https://creativecommons.org/publicdomain/zero/1.0/). 

Specifically, we use the following dataset: **https://openneuro.org/datasets/ds002336/versions/2.0.2**. The corresponding research was conducted at Inria and other institutions: *Lioi, G., Cury, C., Perronnet, L. et al. (2020). Simultaneous EEG-fMRI during a neurofeedback task, a brain imaging dataset for multimodal data integration. Scientific Data, 7, 173, https://doi.org/10.1038/s41597-020-0498-3*. This dataset is the first open access bimodal NF dataset integrating EEG and fMRI, and has several interesting characteristics, notably: 1) It contains EEG data, fMRI data, and NF scores for all subjects. 2) The article contains detailed methodological sections. 3) EEG data has already been preprocessed by the authors. 4) The article presenting this research is publicly available, and can be found here: https://www.nature.com/articles/s41597-020-0498-3. It should be noted that this dataset contains only the data from the first experiment (XP1) reported in the article. The data from the second experiment (XP2) is also accessible on OpenNeuro, but in this project, we focus on XP1. 

The scientific detail of this research is outside the scope of this project, but in summary, the authors conducted a NF experiment where 10 subjects were instructed to use EEG NF scores and fMRI NF scores to perform as well as possible in a motor imagery task, i.e., they needed to execute mentally a movement without any muscle activation. The experiment included six conditions, with alternating rest (20 seconds) and task (20 seconds) blocks within the conditions. In this project, we only use three conditions: the eegfmriNF condition, corresponding to bimodal EEG-fMRI NF, the eegNF condition, corresponding to unimodal EEG NF, and the fmriNF condition, corresponding to unimodal fMRI NF. The three NF conditions were completed in random order by the different subjects. For each condition, the dataset includes the raw fMRI data, the raw EEG data, and the EEG data preprocessed by the authors. It should be noted that EEG acquisition during an fMRI experiment is technically complex, and results in high noise. Therefore, it is extremely useful that the authors included the preprocessed EEG data in the dataset. However, since the fMRI data preprocessed by the authors is not included, we need to perform some fMRI preprocessing steps by ourselves. 

Before running the Notebooks, the user should download the dataset by following the instructions on the OpenNeuro website: https://openneuro.org/datasets/ds002336/versions/2.0.2. Then, the fMRI data needs to be preprocessed with fMRIPrep 23.2.0, a robust preprocessing pipeline which automatically performs a series of fMRI preprocessing steps: https://fmriprep.org/en/23.2.0/index.html. Specifically, the fMRI data needs to be normalized using the MNI152Lin output space, while ignoring slice timing correction. Choosing this output space ensures compatibility with the Harvard-Oxford cortical atlas. If the user chooses, as is recommended, the Docker execution with the fmriprep-docker wrapper (https://fmriprep.org/en/23.2.0/installation.html), running fMRIPrep with these parameters can be done with a command such as this one:

*fmriprep-docker ds002336-download fmriprep_data participant --participant-label sub-xp101 --fs-license-file freesurfer/license.txt --ignore slicetiming --output-spaces MNI152Lin*

This command should be repeated for subjects sub-xp101 to sub-xp110. Depending on the machine, the execution can require several hours per subject to complete. Before running fMRIPrep, the .bidsignore file given in the project directory (.bidsignore) should be added to the dataset directory, in order to ensure BIDS compliance. Furthermore, a FreeSurfer license should be requested here: https://surfer.nmr.mgh.harvard.edu/registration.html. In the command above, this license is located in a freesurfer directory (freesurfer/license.txt). Preprocessing the fMRI data with fMRIPrep is a long step, but one that should absolutely be done before running the Notebooks. If necessary, a complete fMRIPrep tutorial can be found here: https://reproducibility.stanford.edu/fmriprep-tutorial-running-the-docker-image/. 

### Methods

***Terminology Note:*** In this project, the term *machine learning* refers to all non-neural network machine learning models, while the term *deep learning* is used specifically for neural networks. 

In the first Notebook, *fMRI Preprocessing*, we preprocess the fMRI data using the NiBabel and Nilearn libraries, and extract the average voxel values of our brain regions of interest. We remove a systematic bias of the BOLD (Blood Oxygen Level Dependent) signal, which tends to increase during a fMRI session. We also normalize the BOLD signal for the targets, and replace the outliers (STD > 3) by the value of the previous scan. This normalization is applied independently to all fMRI data, whether this data will eventually be used as a train set, a validation set, or a test set. Indeed, our objective is not to predict the absolute value of the BOLD signal, but rather its fluctuations during an experimental condition, and normalization is a straightforward way to achieve this objective. 

In the second Notebook, *EEG Preprocessing*, we preprocess the EEG data using the MNE and YASA libraries. We compute the Power Spectral Density (PSD) for frequencies between 1 and 40 Hz, using both the Welch method and the multitaper method, and the bandpowers for a series of frequency bands of interest: delta (1-4 Hz), theta (4-8 Hz), alpha (8-12 Hz), sigma (12-16 Hz), beta (16-30 Hz), and gamma (30-40 Hz). These frequency bands are widely used in EEG research, although their precise definition can vary. We normalize the bandpower features, and replace the outliers (STD > 4) by the values of the previous scan, while also keeping the bandpowers without normalization as an alternative features set. 

In the third Notebook, *Classification Models*, we train a series of machine learning models for a classification task, using the Scikit-Learn library: logistic regression, k-nearest neighbors (KNN), decision tree (DT), random forest (RF), support vector machine (SVM), and extreme gradient boosting (XGBoost). We select a sequence length of 5 scans, meaning that we consider the EEG bandpowers computed during the fMRI scan of interest, and the ones computed during the 5 previous scans. This sequence length corresponds to 10 seconds, a duration that encompasses the peak of the Hemodynamic Response Function (HRF), improving our chances of capturing, in the EEG data, a trace of the events that influenced the fMRI data. We use the eegfmriNF condition as the train set, the eegNF and fmriNF conditions as the test set, and we evaluate the classification accuracy. When relevant, we fine-tune the model parameters using grid search cross-validation. One subject has a missing eegNF condition and is not included in this and the following Notebooks. 

In the fourth Notebook, *Regression Models*, we train a series of machine learning models for a regression task, using the Scikit-Learn library and the selected sequence length of 5 scans: linear regression, Ridge, Lasso, KNN, DT, RF, SVM, and XGBoost. We use the eegfmriNF condition as the train set, the eegNF and fmriNF conditions as the test set, and we evaluate the regression R^2 metrics. When relevant, we fine-tune the model parameters using grid search cross-validation. 

In the fifth Notebook, *Neural Networks*, sixth Notebook, *Convolutional Neural Networks*, seventh Notebook, *Recurrent Neural Networks*, and eighth Notebook, *Transformers*, we train neural network models with the Adam optimizer, using the TensorFlow library. We use the eegfmriNF condition as the train set, the eegNF condition as the validation set, and the fmriNF condition as the test set. We train our models on the normalized bandpowers, except for the convolutional neural networks, for which we rely directly on the EEG signal. For the classification task, we use the binary cross-entropy loss function and evaluate the classification accuracy, and for the regression task, we use the Mean Squared Error (MSE) loss function and evaluate the regression Mean Absolute Error (MAE). 

In the ninth Notebook, *Large Language Models*, we use two pre-trained LLMs, Gemma-2-2B-IT and Llama-3.2-3B-Instruct, with the same prompts and the same parameters, on the eegfmriNF condition. In order to limit the complexity of the prompts, we associate each fMRI brain region with a single EEG channel, which serves as its sole predictor. This association follows an ad hoc region-channel mapping, which is based, in particular, on electrode proximity to the brain region. The prompts include a general context, the selected EEG channel, the bandpowers for the selected frequency bands, the fMRI brain region of interest, and the description of the prediction task to perform. In this Notebook and the following ones, we use the Hugging Face Transformers library, focus on the classification task only, and select the model parameters in order to obtain a relative variety of responses. We also ensure that our brain regions of interest span at least a reasonable fraction of the brain, and a variety of cognitive functions. 

In the tenth Notebook, *Large Language Model Chain-of-Thought*, we use one pre-trained LLM, Gemma-2-2B-IT, on the eegfmriNF condition. This time, we create a first prompt to infer cognitive functions based on EEG data, and a second, subsequent prompt to infer fMRI data based on cognitive functions. 

In the eleventh Notebook, *Large Language Model Fine-Tuning*, we use one pre-trained LLM, Gemma-2-2B-IT, on the eegfmriNF condition. Then, we fine-tune this model using input-output pairs obtained from the fmriNF condition, and evaluate its performance again on the eegfmriNF condition. 

In the twelfth Notebook, *Large Multimodal Model*, we use one pre-trained LMM, PaliGemma2-3B-Mix-224, on the eegfmriNF condition. Instead of prompting sequences of bandpowers for different frequency bands, we create a topographic map showing the beta bandpower across all EEG channels, and prompt the model with this image, along with a description of the prediction task to perform. 

In the thirteenth Notebook, *Statistical Tests*, we perform Wilcoxon tests and McNemar tests. 

In the fourteenth Noteboook, *Tables and Figures*, we visualize the results. 

### Results

We perform Wilcoxon signed-rank tests on the results of the machine learning and deep learning models, and McNemar's tests on the results of the foundation models. For the machine learning and deep learning models, the baseline is a model selecting the most frequent class for the classification task, and the mean target value for the regression task. For the foundation models, which are applied only on a selection of time steps, the baseline is a model sampling randomly from the target distribution of these time steps. 

All machine learning models for classification, i.e., logistic regression, KNN, DT, RF, SVM, and XGBoost, predict the evolution of the fMRI signal with an accuracy higher than the baseline. Among the machine learning models for regression, i.e., linear regression, Ridge, Lasso, KNN, DT, RF, SVM, and XGBoost, only the RF model and the SVM model predict the evolution of the fMRI signal with a performance better than the baseline, while the other models remain at best at baseline level. 

Among the deep learning models for classification, i.e., feedforward neural networks, convolutional neural networks (CNNs), recurrent neural networks (RNNs), and transformers, all models except CNNs predict the evolution of the fMRI signal with an accuracy higher than the baseline. However, all models except the feedforward neural networks show signs of overfitting, with the validation loss often stagnating or even increasing over the epochs. When these deep learning models are used for regression, none of them generalizes well beyond the train set. Nevertheless, it should be noted that for some subjects and some brain regions, the transformers seem to predict the evolution of the fMRI signal with a performance better than the baseline, and contrary to other neural networks, their prediction values do not tend to fall to the baseline level in general. We do not analyze this effect further, but it suggests a potentially interesting path for future investigation. 

Although the observed effects are much weaker for the foundation models, the Gemma model achieves statistically reliable gains over the baseline, whether it is used with direct prompting or with a Chain-of-Thought approach. With the same prompts and parameters, the Llama model does not reach the baseline. Fine-tuning the Gemma model does not significantly improve its performance. The performance of the PaliGemma model is more ambiguous, with a pattern intermediate between the Gemma and Llama models. 

### Discussion

Using EEG data to predict fMRI data could be more or less challenging, depending on the context, the objective, and the precise definition of the prediction task. For sleeping subjects, making predictions is certainly easier, since the different sleep stages are associated with both distinctive EEG and distinctive fMRI patterns. By contrast, for awake subjects, and particularly for subjects engaged in a demanding cognitive task such as NF, making predictions is certainly much more difficult, since brain patterns are more complex and unpredictable. However, these demanding contexts are arguably the most important ones for expanding our understanding of core executive functions of the human brain, and for developing neurotechnologies targeting these functions. 

Furthermore, using EEG data to predict fMRI data is certainly easier when the prediction task is conducted within a single experimental condition, since the machine learning and deep learning models can be trained, validated, and tested on the same time series. By contrast, in this project, our objective is to develop models that generalize well across conditions, which is significantly more complex. However, once again, this more complex objective is arguably the most important one, since it allows us to envision realistic neurotechnological innovations, such as using an initial EEG-fMRI calibration session to train the models, before leveraging these models to predict fMRI activity in subsequent EEG-only conditions. 

Finally, electric potentials are not always easily accessible to EEG electrodes, even at the cortical level. Moreover, some of our brain regions of interest are primarily associated with high-level, integrated cognitive processes, and not with low-level, sensory mechanisms which are more easily reflected in EEG activity. Overall, in this project, we try to build an AI system for human brain activity prediction in one of the most difficult contexts, but these contexts are precisely the ones where predicting fMRI activity from EEG activity is scientifically and technologically the most relevant. 

This project follows two distinct strategies: directly training machine learning and deep learning models on our EEG-fMRI dataset, or relying on the capabilities of pre-trained LLMs and LMMs. We demonstrate that both strategies yield positive results, which seems to highlight two complementary paths for predicting fMRI signals based on EEG signals. As we mentioned previously, the second strategy is critically dependent on our ability to rely on brain region keywords, e.g., "Frontal Pole", and fMRI signal evolution keywords, i.e., "Increasing" or "Decreasing", which was made possible by using the Harvard-Oxford cortical atlas and adding a classification task. Intuitively, we expect that LLMs might implicitly infer the cognitive functions associated with EEG data, and subsequently the fMRI data associated with these cognitive functions, and our Chain-of-Thought approach points indeed in this direction. 

Based on these promising results, how could we further improve this project, and lay the foundations for an enhanced AI system for human brain activity prediction? We can think of several possibilities for improvement. 

First, given enough time, memory, and computational resources, we could explore additional fMRI and EEG preprocessing pipelines, model architectures and parameters, and prediction strategies. We could also try to include in the training data some characteristics of the subjects, and some elements of the experimental context in which the data was acquired. For LLMs, we could explore additional prompt engineering and parameter tuning techniques, and include the EEG data from several electrodes, instead of a single one. For LMMs, we could also include multiple topographic maps, representing several EEG frequency bands, instead of limiting ourselves to the beta band. Abandoning frequency bands altogether in favor of more complex EEG features is also a possibility, but this would be a double-edged sword, since it would reduce our ability to rely on common EEG keywords. 

Second, we might need to develop methods for integrating EEG-fMRI datasets with different characteristics. Indeed, at a certain point, an AI system for human brain activity prediction will certainly require a significant amount of training data, and a single dataset is unlikely to be sufficient. Nevertheless, such integration could pose a series of complex challenges, since neuroimaging datasets can use different fMRI voxel sizes, different EEG montages, different scan durations, and have many other notable differences. It might be necessary to develop new approaches, and maybe to build new multimodal templates, in order to achieve this integration. Of course, we would certainly benefit from larger datasets, and ideally more balanced in terms of sex and age, if they become available in the future. 

Third, instead of relying on simple and general machine learning and deep learning models, we might need to develop more complex and specific models, based on our knowledge of the human brain. For example, it might be necessary to take into account the brain anatomical and functional connectivity, the dynamics of the fMRI BOLD signal, and the sources of the EEG signal. At this point, we might even consider expanding the scope of the project, and trying to integrate other neuroimaging modalities, such as Magnetoencephalography (MEG). This research direction might imply a multidisciplinary scientific collaboration, and would take us closer to the creation of a multimodal foundation model for neuroscience, integrating EEG, fMRI, and other neuroimaging modalities. 

Finally, as new, more powerful LLMs and LMMs become available, they could become increasingly important tools for predicting fMRI activity based on EEG activity, particularly if these models are pre-trained on a significant corpus of neuroscience knowledge. Combining them with machine learning and deep learning models directly trained on EEG-fMRI datasets, whether through fine-tuning or other methods, could become an important path for further advances. 

### Conclusion

Building an AI system for human brain activity prediction should most probably be considered as a long-term endeavor. Nevertheless, the methods developed in this project could already represent an important step, by demonstrating the feasibility of predicting fMRI activity from EEG activity, even in the challenging context of subjects performing a cognitive task such as NF. Overall, it is difficult to overstate the importance of this research direction, and the impact that it could have for neuroscience and neurotechnology. While fMRI is a complex and expensive neuroimaging technique, EEG is much simpler and more affordable, and better adapted to a diversity of real-life situations. A future iteration of this project, or another similar initiative, could accelerate neuroscience research, connect the scientific results from several neuroimaging realms, and improve promising brain technologies, such as NF systems and brain-computer interfaces. 

Working toward the creation of a multimodal foundation model for neuroscience, integrating EEG, fMRI, and other neuroimaging modalities, seems to be a natural horizon for scientific research on the human brain. If such a foundation model can be built, it could become an important tool for neuroscience laboratories, neurotechnology enterprises, and even AI laboratories willing to replicate more closely the processes of the human brain. From an AI-powered neuroscience platform to a tool for neuroscience-inspired AI, the possibilities of an AI system for human brain activity prediction seem endless, and would naturally require careful reflection and evaluation, considering the sensitive nature of human brain data. Hopefully, this project will contribute to this objective. 

### Structure and Environment

This project includes fourteen Notebooks, to be run in order:

1. fMRI Preprocessing
2. EEG Preprocessing
3. Classification Models
4. Regression Models
5. Neural Networks
6. Convolutional Neural Networks
7. Recurrent Neural Networks
8. Transformers
9. Large Language Models
10. Large Language Model Chain-of-Thought
11. Large Language Model Fine-Tuning
12. Large Multimodal Model
13. Statistical Tests
14. Tables and Figures

The Conda environment for this project, including in particular the NiBabel, Nilearn, MNE, YASA, Scikit-Learn, TensorFlow, and Hugging Face Transformers libraries, can be recreated with the .yml file given in the project directory (ENV.yml). 
